# Multi-Omic Profiling of the Liver Across Diets and Age in a Diverse Mouse Population

**DOI:** 10.1101/2020.08.20.222968

**Authors:** Evan G. Williams, Niklas Pfister, Suheeta Roy, Cyril Statzer, Jack Haverty, Jesse Ingels, Casey Bohl, Moaraj Hasan, Jelena Čuklina, Peter Bühlmann, Nicola Zamboni, Lu Lu, Collin Y. Ewald, Robert W. Williams, Ruedi Aebersold

## Abstract

Systems biology approaches often use inferred networks of gene expression and metabolite data to identify regulatory factors and pathways connected with phenotypic variance. Generally, study-specific multi-layer “Omics” datasets are used to contextualize generic molecular networks. In this regard separating upstream causal mechanisms, downstream biomarkers, and incidental correlations remains a significant challenge, yet it is essential for designing mechanistic experiments. To address this, we designed a study following a population of 2157 individuals from 89 isogenic BXD mouse strains across their lifespan to identify molecular interactions among genotype, environment, age (GxExA) and metabolic fitness. Each strain was separated into two cohorts, one fed low fat (6% cal/fat) and the other high fat (60% cal/fat) diets. Tissues were collected for 662 individuals (309 cohorts) diverging across age (7, 12, 18, and 24 months), diet, sex, and strain. Transcriptome, proteome, and metabolome data were generated for liver. Of these we identified linear relations among these molecular data with lifespan for the same genomes of mice (Roy et al. 2020), and we defined ∼1100 novel protein-coding genes associated with longevity. We knocked down the ortholog of *Ctsd* in *C. elegans*. The treatment reduced longevity both in wildtype and in mutant long-lived strains, thus validating the prediction. Next, to assess the molecular impact of GxExA on gene expression, the multi-omics data was parsed into metabolic networks where connectivity varied due to the independent variables. Differences in edge strengths connecting nodes in these molecular networks according to each variable enabled causal inference by using stability selection, with roughly 21% of novel gene–pathway connections being causally affected by diet and/or age. For instance, *Chchd2* is activated by aging and drives changes in the proteasome, oxidative phosphorylation, and mitochondrial translation transcriptional networks. Together, we have developed a large multi-omics resource for studying aging in the liver, and a resource for turning standard associations into causal networks.

## INTRODUCTION

Aging is a dynamic and multi-faceted process driven over a lifetime of interactions among genetic variants, environmental factors, and stochastic processes. Despite its complexity, lifespan is a heritable trait, with genotype explaining 30–50% of its variation across laboratory mice [1, 2] and ∼25% in humans [1]. Age is the most prominent “risk factor” for a wide range of diseases, such as metabolic syndrome, diabetes, heart disease, neurodegeneration, and most cancers [3, 4]. Cells and tissues display common perturbations with increasing age such as a diminished capacity for proteostasis [5, 6] and the accumulation of mitochondrial defects [7]. These and other common endpoints are recognized, but there is substantial diversity in the mechanisms and timelines connecting chronological age (“lifespan”) to biological age (“healthspan”) across individuals and tissues [8], let alone across organisms. It is now possible to measure biomarkers of biological age, such as by DNA methylation signatures [9]. However, it is unclear whether interventions that directly affect the dynamics of aging biomarkers, such as methylation, would causally improve either lifespan or healthspan [10, 11]. Given the inherent challenges of obtaining tissue biopsies longitudinally in human clinical cohorts, replicable but genetically diverse model organisms take a unique position in the biomolecular analysis of aging. In particular, isogenic cohorts permit “paired” tissue biopsies to be collected across multiple times and environments. This allows the creation of resources such as the *Tabula Muris Senis* project [12], which have established a baseline resource for how transcript expression changes across tissues and time for one particular genome of mouse—C57BL/6. Further research is necessary to test how genetic variation and environmental interactions (GxE) influence molecular aging, the extent to which relation are congruent between cognate mRNA and protein, how changes in molecular levels link to aging phenotypes. Getting at the causality of these relations is critical in developing more sophisticated interventions to reduce disease burden and enhance health and lifespan. However, causal inference requires multiple simultaneous axes of variation and/or longitudinal sample acquisition [13, 14], a relative rarity for population-scale studies of gene expression, which tend to focus on cross-sectional analysis for single independent variables.

In this study, we have generated transcriptome, proteome and metabolome profiles from liver samples from animals of the large and genetically diverse BXD mouse family across time and over two diets. These data were generated in livers from 300 distinct genotype, age, diet, and sex-matched cohorts and combined with phenotypes collected across the entire family, including blood biomarkers, organ weights, longitudinal body weight, and of greatest importance—lifespan. We used this unique dataset to examine the relations between genetics, dietary environments, and age (GxExA). We show that the data resource provides a platform for detecting, evaluating, and testing how biomolecular processes vary as a function of GxExA, and the extent that each tier of molecular data reveals changes in metabolic gene networks linked to key outcomes and phenotypes. Moreover, we use the dataset to highlight an approach to develop extensive and testable causal models in biomedical research. This step is essential to facilitate a shift towards an integrative data analysis strategy that takes advantage of increased complexity in study designs, advances in measurement technologies, and greater sample sizes emerging in life science studies.

The multiple independent variables segregating in this study (diet, age, genotype) combined with molecular profiles allowed us to broadly apply a causal inference method we recently developed [15, 16] called stabilized regression. This method starts with supervised learning, using as input a target of interest (e.g. gene expression or a phenotype), searches for any measurements which covary with the target, then evaluates how these associations change as a function of each independent variable. Regression coefficients for each independent variable are combined with a stability score that estimates whether the novel target is more likely to be upstream of a canonical pathway (i.e. causal), downstream (i.e. a biomarker), or ambiguous (i.e. a connection not affected by the secondary independent variable). We have performed causal inference analysis for 23 gene sets which are known to vary as a function of diet or age, searching for modifier genes outside the canonical gene sets which explain differences in gene expression networks as a function of genotype, age, and diet. Roughly 20% of the detected novel gene–pathway associations were specific to an age or dietary environment, indicating a causal relationship between these genes, the target pathway, and an independent variable (i.e. age or diet).

Here we have generated the largest and most extensive set of replicable multi-omics aging tissue data in a multivariate study design. It provides two key resources for the study of aging, metabolism, and complex trait analysis. First, these data were generated in the inbred BXD population, and thus provide an extensible reference for the range of effects of GxExA on gene expression, metabolites, and core physiological phenotypes such as lifespan and body weight. Second, this multivariate study design (and data) demonstrates the capacity for new advances in statistics for the study of complex networks: stabilized regression can calculate the causality for associations which are impacted by two or more independent causal variables.

## RESULTS

### Clinical Analysis of Lifespan as a Function of Genotype and Diet

In this study, we followed 2157 mice from 89 strains of the BXD family across their natural range of lifespan. These individuals were placed in the colony around 5 months of age, after which cohorts were evenly segregated into two dietary cohorts, one fed a standard chow diet (CD; Harlan Teklad 2018, 6% calories from fat) and the other a high fat diet (HFD; Harlan 06414, 60% calories from fat). For 60 strains, selected pairs of individuals from each cohort (i.e. strain and diet matched) were sacrificed at 7, 12, 18, and if possible 24 months of age to collect a biobank consisting of 662 individuals with tissues across time, diet, and genotype (**Figure 1A**, **Table S1**). These individuals belong to 309 distinct cohorts (i.e. matched for age, sex, strain, and diet), which are age and diet-balanced (**Figure S1A**). The liver was selected as the primary organ of interest due to its central role in metabolism and the wide range of liver-related clinical and molecular phenotypes known to vary across the BXDs as a function of diet, sex, and genotype [17, 18]. The liver was pulverized in liquid nitrogen and aliquoted for transcriptomic, proteomic, and metabolomic preparations which were performed in parallel (**Figure 1B**).

**Figure 1.**
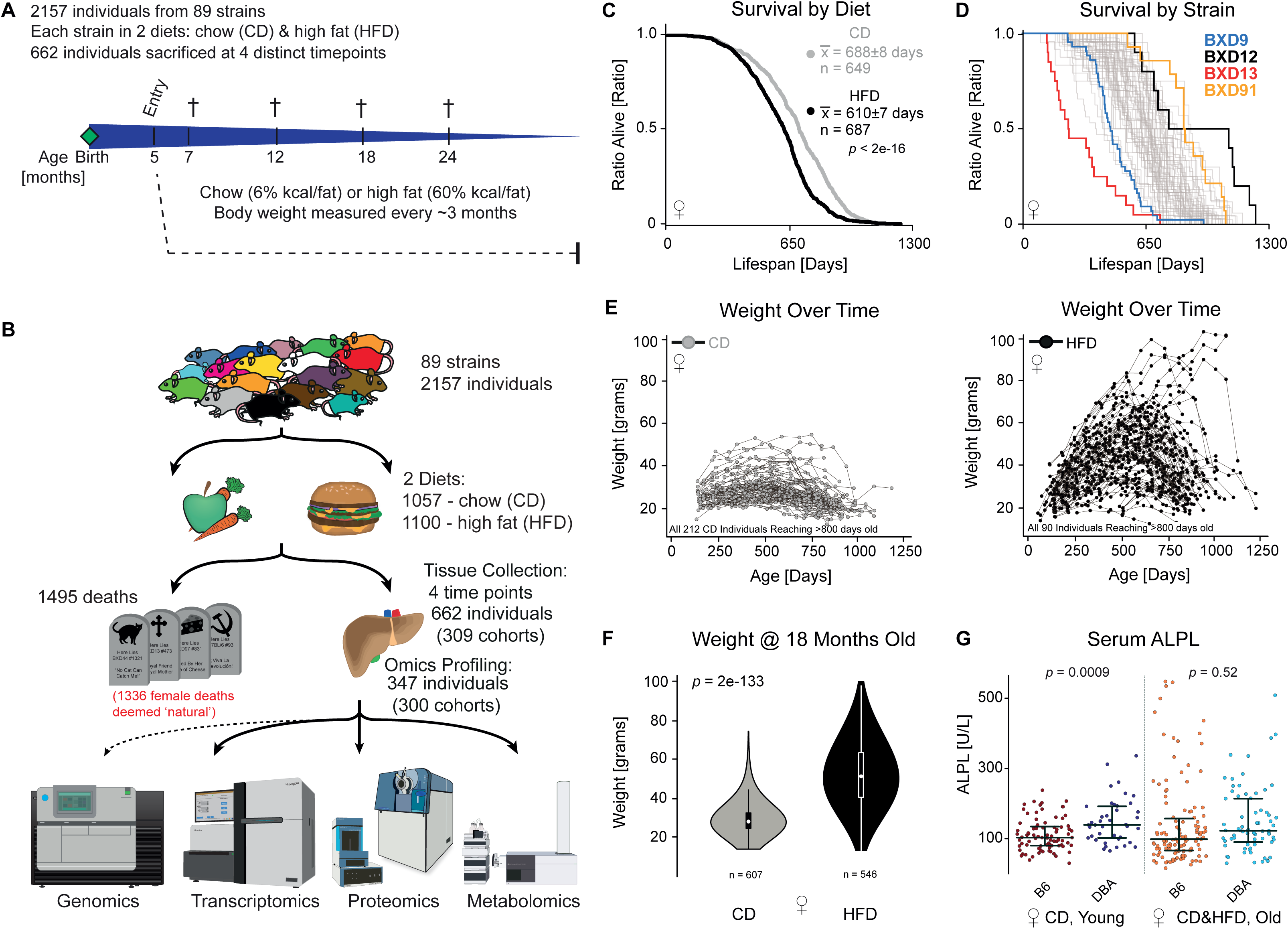
Overview of Aging Colony (**A**) Study overview. Animals entered the aging program at around 151 days of age and were set into dietary cohorts. 662 individuals were selected for sacrifice at 7, 12, 18, and/or 24 months of age for sacrifice. (**B**) Workflow of the lifespan and tissue collection cohorts. The 662 individuals are from 309 cohorts according to diet, age, sex, and strain. 343 individuals were selected for omics profiling, corresponding to 300 distinct cohorts out of the 309 originally acquired. Of the 1495 natural deaths, 1336 were used for lifespan calculations (see Methods). (**C**) Kaplan–Meier survival curves for CD and HFD females, irrespective of strain. (**D**) Kaplan–Meier survival curves for all 60 BXD strains with at least 8 natural deaths in the female cohorts, irrespective of diet. (**E**) Weight-over-time for 212 CD (left) and 90 HFD (right) individuals which reached ≥ 800 days of age. All animals were weighed bimonthly. (**F**) Violin plots of body weight at 18±1.8 months of age in each diet. (**G**) ALPL serum metabolite levels across BXD strains as a function of several cofactors. The canonical allelic effect is clear for young, CD cohorts but it is masked by GxExA in other cohorts.

Earlier studies have demonstrated that lifespan across the BXD family of mice varies by ∼3-fold—from 11 to 32 months [19–21]. These differences among strains are consistent across studies, even twenty years apart (r = 0.77, **Figure S1B** [1, 20, 21]). Likewise, we find significant correlations between our CD lifespans [22] and those of the previous studies (r = 0.53 and = 0.69, for the 1988 and 2010 studies, respectively, **Figure S1B**). In this study, we considered 1336 individual female mice which lived out their natural lifespans for the calculation of longevity, permitting comparisons across diets (**Figure 1C**) and strain (**Figure 1D**). 48 strains had sufficient data in both diets for this analysis (≥ 6 natural deaths in both dietary cohorts; **Table S1**). Genetic variation across the population explained 73% of variation in mean lifespan, versus 12% by diet and 15% by gene–diet interactions, with an unexplained residual of only 0.5% (**Figure S1C**). Mean strain lifespans, with both dietary cohorts mixed together, vary from 314±37 days (mean ± SEM for BXD13, n = 20) to 870±39 days (BXD91, n = 14) (**Figure 1D, Figure S1D**). While HFD feeding causes a mean 10% decrease in longevity, the magnitude of decrease varied by strain: BXD9’s lifespan is unaffected, while BXD65 lives nearly an additional year longer on CD (p = 3e-6, **Figure S1E**). HFD leads to a significant decrease in lifespan in 40% of the strains (p ≤ 0.05, Kaplan–Meier curves), and 64% have at least a tendency to live shorter on HFD (p ≤ 0.10). These differences notwithstanding, the HFD effect is relatively consistent across strains, with the mean lifespan of strains between diets correlating at r = 0.65 (**Figure S1F**). Although diet has a relatively modest effect on longevity, it has a substantial impact on weight (**Figure 1E–F**), explaining 40% of variance (versus 23% explained by genotype, 5% by genotype-diet interaction, and a 23% unexplained residual, **Figure S1C**). By 18 months of age, individuals had an average 78% increase in body mass and 89% of strains gained weight significantly upon HFD feeding (p < 0.05, AUC). As with lifespans, the effect of HFD on body weight varied depending on genetic background: BXD16s gained the least with an average increase of 11%, while BXD100s gained an average of 133% (**Figure S1G–H**).

In addition to body weight and longevity, we also measured 18 plasma metabolites commonly used in clinical settings (e.g. cholesterol, iron, glucose, and alkaline phosphatase (ALPL) levels; **Table S2**). As a function of diet and genotype, we observed that HFD reduces the circulating serum level of ALPL, as reported previously in the BXDs [18] along with an increase in circulating ALPL in old mice (**Figure S1I**), as has been observed in humans [23]. Strains with the B6 allele of *Alpl* are known to have lower ALPL levels than those with the D2 allele [17], which we again observed in typical control conditions (i.e. CD, young) (*p* = 0.004, **Figure 1G**). However, this effect is not independent of environment: for old females, the effect caused by *Alpl* sequence variants is masked by environmental interactions between age and diet (p = 0.54, **Figure 1G**). These interactions between age, genotype, and diet on ALPL are known [17, 18], but this illustrates the essential challenge of causal discovery: a single circulating metabolite is affected by genotype (*Alpl* allelic variants), diet, and age. Even with such a large sample size, the p-values of traditional statistical analyses would be non-significant after correction for multiple testing without this prior biological knowledge or a more targeted statistical approach.

### Multi-Layer Molecular Analysis of the Aging Liver

Gene expression of mRNA and protein makes up the most comprehensively quantifiable estimate of gene activity. We can quantify to what extent these gene products are influenced by GxExA and use this understanding to model how gene networks respond to differences in age, diet, and genotype, resulting in diverging complex clinical phenotypes. While an mRNA and its protein both result from the same gene, the expression of these two paired gene products tends to diverge unpredictably in response to stimuli, with only ∼20-30% of the variance for an average protein being explained by variance in its mRNA [24]. Furthermore, studies on aging have found that only a small minority of transcripts vary substantially across age, e.g. only 0.4% of transcripts varied by ≥ 2-fold in muscle tissue between young and old animals as a function of caloric restriction (CR) [25]. Similar effect sizes have been seen for proteomics, with as little as 1% of protein assays changing by more than 2-fold in mice between 5 and 26 months-of-age [26]. Conversely, comparatively sized (i.e. ∼2-fold) effect sizes have been observed for overall changes in pathway activity across lifespans, e.g. ribosome translation rates drop by as much as 2-fold between young and old mice [27]. Differential responses of mRNA and protein expression as a function of GxExA for entire pathways led us to search for patterns in these relationships, and the consequences this would have on hypothesis generation and validation. By comprehensively quantifying these molecular traits in the liver— mRNA, protein, metabolites—we hypothesized that we could identify key genes and molecular networks involved in the etiology of hepatic aging, dietary response, and their interactions with genetic variants across the population.

We selected livers from 347 individuals for multi-omic gene expression analysis, representing 300 of the 309 original cohorts according to strain, sex, age, and diet (**Table S1**, sheet “Cohorts_Harvested”). After sample quality control (QC; see Methods), this resulted in RNA-seq data from 291 individuals (255 cohorts) and proteomics data from 315 individuals (278 cohorts), with 275 individuals overlapping in both datasets (240 cohorts). Furthermore, untargeted metabolomics data were generated for livers from 624 individuals (298 cohorts), resulting in a total of 274 individuals (239 cohorts) with full data in all three layers. RNA-seq data were generated with 20 million reads per sample on a HiSeq PE150, with 25394 distinct transcripts quantified, of which 20827 are annotated as protein-coding. Proteomics data were generated using SWATH-MS on a Sciex 6600 instrument, with 3940 proteins quantified after QC. The metabolomics data were generated on an Agilent 6550 instrument, with 464 uniquely-detected metabolites remaining after QC. The processed and normalized set of all omics data are available in **Table S2** (for raw data, see **Data Availability**). We first considered the 3772 genes which were measured at both the mRNA and protein level to get a comparable overview of the expression data. These genes include some highly represented ontologies (e.g. mitochondria, cytoplasm, and ribosomal proteins) while others are depleted (e.g. membrane proteins and secreted proteins) (**Table S3**, sheet 1). Some functional categories are fundamentally absent due to tissue type (e.g. olfactory receptors) or selection time (e.g. developmental proteins), while other depletions are due to technical reasons, e.g. membrane-bound proteins are difficult to extract, separate, and digest in proteomics [28, 29].

Diet had a significant impact on the expression of 893 transcripts and 1352 proteins (**Figure 2A**, adjusted t-test between discrete groups) while 1562 transcripts and 998 proteins significantly covaried with age (**Figure S2A**, correlation coefficient, adjusted p-value). An average of 65% of observed variation in transcripts and proteins was explained by the three independent variables—genotype (“strain”), diet, age—and their GxExA interaction (**Figure 2B**). While strain had the largest individual effect (∼28%), the GxExA term accounted for the plurality of explained variance (∼33%), with a similarly-sized unexplained residual (∼35%). Conversely, for metabolites only 45% of observed variance was explainable by the independent variables, where “strain” had again the strongest average (∼16%) but with a reduced interaction term (∼24%) and much larger residual (∼55%) (**Figure S2B**). For an individual gene, its transcript and protein were correspondingly affected by diet or age (r = 0.36 and r = 0.33 respectively; **Figure 2C**), while the effects of diet and age on gene expression were independent overall (**Figure S2C**). We next examined the relationships between mRNA and protein levels. Across all samples, mRNA is reasonably predictive for the abundance of a protein (r = 0.44, **Figure 2D**)—that is, more abundant mRNAs tend to be the more abundant proteins and vice-versa. However, we are generally interested in how genes and pathways respond to perturbations (i.e. genotype, diet, or age). The average correlation of all 3772 mRNA with their protein as a function of GxExA was rho = 0.14, with 33% of mRNA–protein pairs covarying significantly across all measurements (adj.p < 0.05; **Figure 2E**). That is, knowing the variation in mRNA expression across genotype, diet, and age provides only a weak predictor for variance in its corresponding protein.

**Figure 2.**
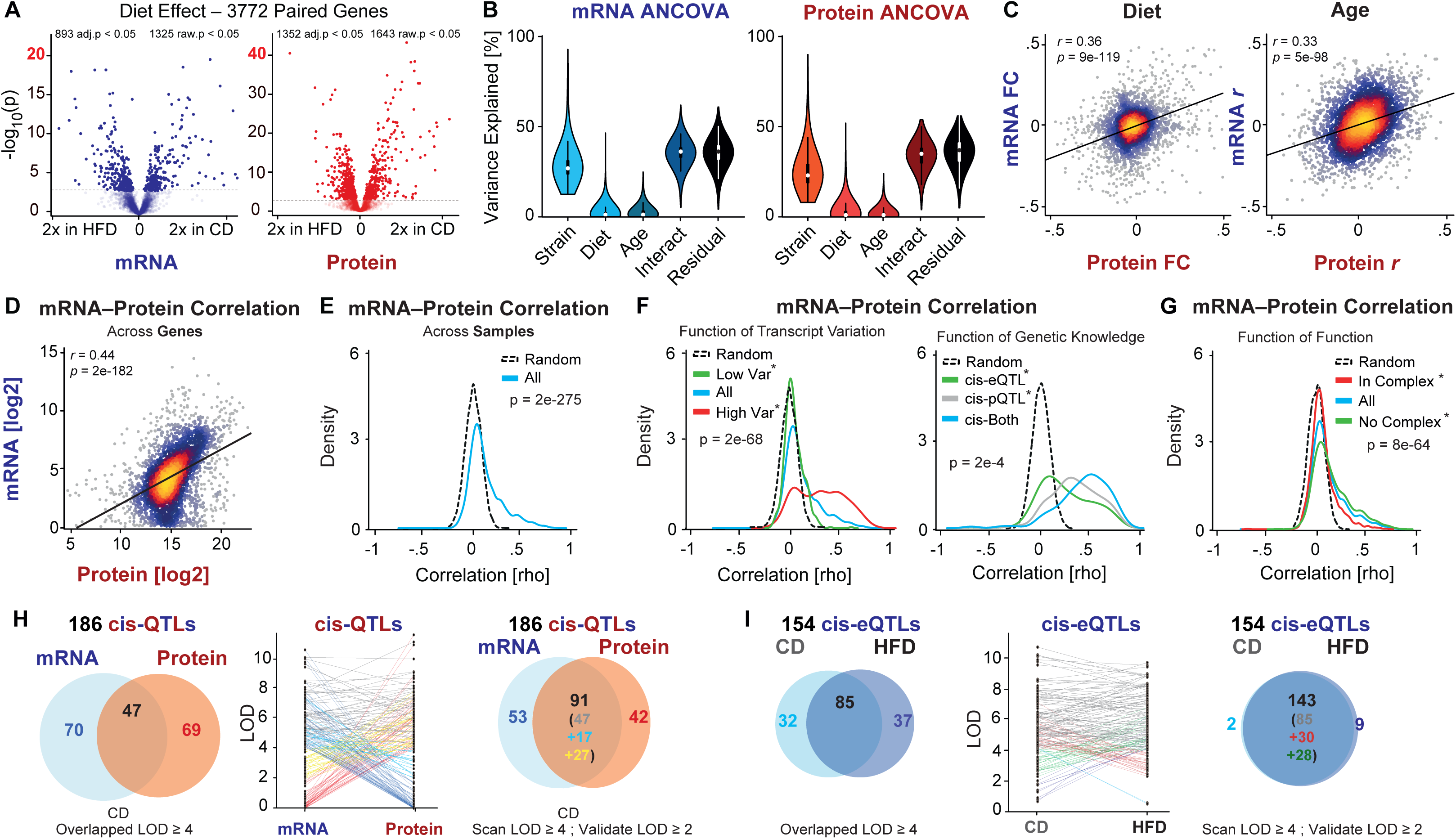
**Multi-omics Overview of mRNA, Protein, and Metabolite Liver Expression** (**A**) Volcano plot for all 3772 transcripts and proteins measured in both expression types. 22% of transcripts and 35% of proteins were significantly affected by diet. (**B**) Variation explained (ANCOVA) as a function of the independent variables, their interactions, and the residual. The “interact” term is the sum of all interactions between diet*age, diet*strain, age*strain, age*diet, and age*diet*strain. (**C**) Correlation density plot of the relationship between the effects of diet (left) and age (right) on genes’ product mRNAs and proteins. Brighter colors represent higher density data. This is the data from the X-axes in the Volcano plots in (A); transcripts that are affected strongly by diet or age tend to be proteins which are significantly affected. (**D**) Correlation of average mRNA and protein levels across all samples and all genes, i.e. lowly-expressed transcripts tend to be lowly-expressed proteins and vice-versa. (**E**) Density plot showing ∼33% of transcripts significantly covary with their protein (i.e. area under the blue curve but above the black curve of randomized data). (**F**) (Left) Density plot of mRNA–protein correlations as a function of the mRNA expression variance; (right) density plot showing correlation as a function of cis-QTL presence; (center-right) as a function of diet (**G**) Density plot of correlation as a function of the gene existing in a complex. Reported p-values are for the comparisons marked with an asterisk (*). (**H**) (Left) Genes with highly-significant cis-QTLs (LOD ≥ 4) in CD cohorts at the mRNA and protein level. (Middle) Slopegraph showing the change in LOD score between mRNA and protein; (Right) Venn diagram showing ∼50% cis-QTL overlap at more permissive cutoffs (LOD ≥ 4 in discovery cohort, ≥ 2 in validation cohort). (**I**) (Left) Across diet at the mRNA level, roughly half of cis-eQTLs are found congruently. (Middle) Slopegraph showing the change in LOD score across diet. (Right) Venn diagram accounting for less strict alignment cutoffs, showing ≥ 90% of cis-eQTLs align across diet.

Despite this low average correlation, additional data can be used to improve the predictive capacity of mRNA for its protein product in some cases. Independent variables with a large effect size on a transcript’s expression are far more likely to have a corresponding effect on the protein’s expression. More highly variable transcripts tended to correlate better with their proteins—71% of the most abundant decile of transcripts covary with their protein, versus only 6% of the least-abundant decile (**Figure 2F**, **Figure S2D)**. More abundant transcripts also tended to correlate better: only 12% of the least-abundant decile of transcripts covaried with their protein, compared to 63% of the most-abundant. While this could indicate higher levels of noise in low-abundance transcripts (and proteins, given abundance correlates at r = 0.44), it is worth noting that abundance and variability are only weakly correlated (rho = 0.07, **Figure S2F**). Thus, measurement knowledge of these variation within an omics layer indicates some edge cases where protein and transcript measurements can be used as reasonable proxies. For the 60 transcripts that are in the top decile of abundance and variability, 87% correlate with their protein significantly and with an average rho = 0.51—versus an average 33% at an average rho = 0.14 for an average pair across the full set.

Other factors stemming from prior knowledge can also be used to predict mRNA–protein covariation. For instance, genes with significant quantitative trait loci (QTLs) mapping to their own location—i.e. cis-QTLs—are more likely to have more significant transcript–protein relationships (**Figure 2F**). QTLs indicate a causal impact of DNA sequence variants inside (or adjacent to) a gene which cause varying transcript expression levels (cis-eQTL) or protein levels (cis-pQTL), and these tend to be highly robust and reproducible [30]. The knowledge of which genes have cis-QTLs can also provide predictive information across expression type: transcripts that have strong cis-eQTLs (logarithm of the odds (LOD) ≥ 4) correlate substantially better with their protein (rho ∼ 0.27). Other predictive patterns can be observed for particular gene categories. For instance, genes that are involved in all complexes annotated by CORUM [31] are substantially less likely to have significant mRNA–protein covariance (average rho ∼ 0.06, **Figure 2G**). This is despite that complex-member mRNAs tend to be slightly more abundant than average (p = 2e-6) and have no difference in their variation (p = 0.08). More importantly, the size of the complex impacts the expected correlation: at an adjusted p < 0.05, 34% of the 360 quantified genes in dimers have significant mRNA–protein correlations, against only 4% of the 431 genes in complexes of ≥ 20 subunits (i.e. not more significant than expected by chance) (**Figure S2G**). Together, these findings indicate that the highest-variance genes selected through differential gene expression analysis will *mostly* validate across-layer. However, caution must be taken for smaller effects. This is crucial for the study of complex traits and GxExA, as we recall that expression variation is relatively subdued: < 1% of genes vary by >2-fold as a function of age. That is, overlapping mRNA–protein results are helpful for increasing confidence in the mechanism, but these will drive a small percentage of the overall connectivity in networks. Critical molecular changes may only be evident at the transcriptome or the proteome level, and thus the discrepancies must be retained (e.g. [18, 32]).

Finally, we examined the relationships between gene expression and the varying genetics across the BXD population via QTL mapping on all 3772 transcript–protein pairs. 216 genes mapped to a significant cis-eQTL or cis-pQTL at LOD ≥ 4 (**Figure 2H**; >99.9% true positive rate using *discovery* cutoffs, **Figure S2H**). While only 25% of cis-QTLs were observed at this threshold for both mRNA and protein levels concurrently (i.e. 53 out of 216), an additional 24% were observed at a secondary threshold when followed-up with a specific QTL hypothesis (LOD ≥ 2, corresponding to a 99.7% true positive rate when used as a *validation* cutoff, **Figure S2H**). The observation that nearly half of cis-QTLs (49%, i.e. rightmost panel of **Figure 2H**) are unique to transcript or protein levels is in line with previous results (e.g. [33]) and further underscores the utility of separately analyzing both types of gene product. We next examined the reproducibility of cis-QTLs as a consequence of diet. At discovery cutoffs (i.e. LOD ≥ 4), just over half of cis-eQTLs (**Figure 2I**) and cis-pQTLs (**Figure S2I**) were observed in common across diets, while at validation cutoffs, more than 90% of cis-QTLs—for both transcripts and proteins, separately—were observed in both dietary conditions. However, it is worth noting that some genes only yield cis-QTLs under certain environmental states, such as *Cyp3a11* and *Cyp3a16*, which map to robust *cis*-pQTLs, but exclusively in HFD, or *Akt2* which maps to a robust cis-pQTL but only in aged animals (**Table S3**, sheet 2). This indicates that in general the interactions of diet with genetic background have a modulating interaction combined with genotype, but not a uniform effect across all strains, a parallel analysis leading to the same conclusion as ANCOVA (i.e. **Figure 2B**). Similar general trends were observed when comparing cis-QTLs across age instead of diet; 45% of cis-QTLs were found to concordantly affect both transcript and protein within age group, while 92% of cis-pQTLs were found in common across age groups (**Figure S2J**). Together, these results indicate that while genes’ mRNA and protein products have many common responses to GxE variables, they cannot be *ad hoc* assumed to be proxies for one-another. Hypothesis discovery and molecular analyses are thus expected to yield unique results when the different measurement layers are used separately, or better yet in tandem.

### Metabolic Characteristics of Age

Next, we correlated all gene expression data with the measured age of the animals to look for molecular signatures of aging (**Table S4**). Dietary cohorts were correlated separately and we selected the 100 proteins and transcripts from each diet with the highest longevity correlation for DAVID analysis [34] (corresponding to p < 5e-4 in both CD and HFD, equivalent to rho > ∼|0.34|). This combined list of 158 mRNAs and 176 proteins was mined for enrichment in KEGG pathways and in GO cellular compartments. The extracellular exosome was by far the most enriched compartment in both mRNA (45% of genes, p = 2e-20) and protein (60% of genes, p = 8e-44), with the mitochondria coming in a distant second for both (mRNA: 29% of genes, p = 7e-12, protein: 23% of genes, p = 7e-7) (Figure 3A). Few functional pathways were enriched, with only steroid hormone biosynthesis highlighted with mRNA (p = 1e-8) and the lysosome pathway for protein (p = 1e-5). The tremendous enrichment of extracellular genes with age highlights recent research into the aging-related decline of homeostasis within the extracellular matrix (ECM, or “matrisome”) [35]. We quantitatively examined this particular association in more detail using Gene Set Enrichment Analysis (GSEA) [36]. We observed that not only are ECM genes disproportionately associated with aging, but they are *directionally* associated, with an age-associated increase in ECM transcripts and proteins (**Figure 3B**). Furthermore, research in collagens (a key ECM component) have shown that COL1A1 mutant mice have decreased lifespans—showing that a disbalance in the ECM can in fact *drive* aging [37], rather than being a simple bystander. The DAVID analysis highlighted additional associations in the BXD liver data between aging and other pathways, such as the lysosome and mitochondria. Loss of proteostasis has been recognized as one of the hallmarks of aging, a mechanism with which the lysosomes have a direct relationship [38], and mitochondrial function is likewise known to decrease with age [39].

**Figure 3.**
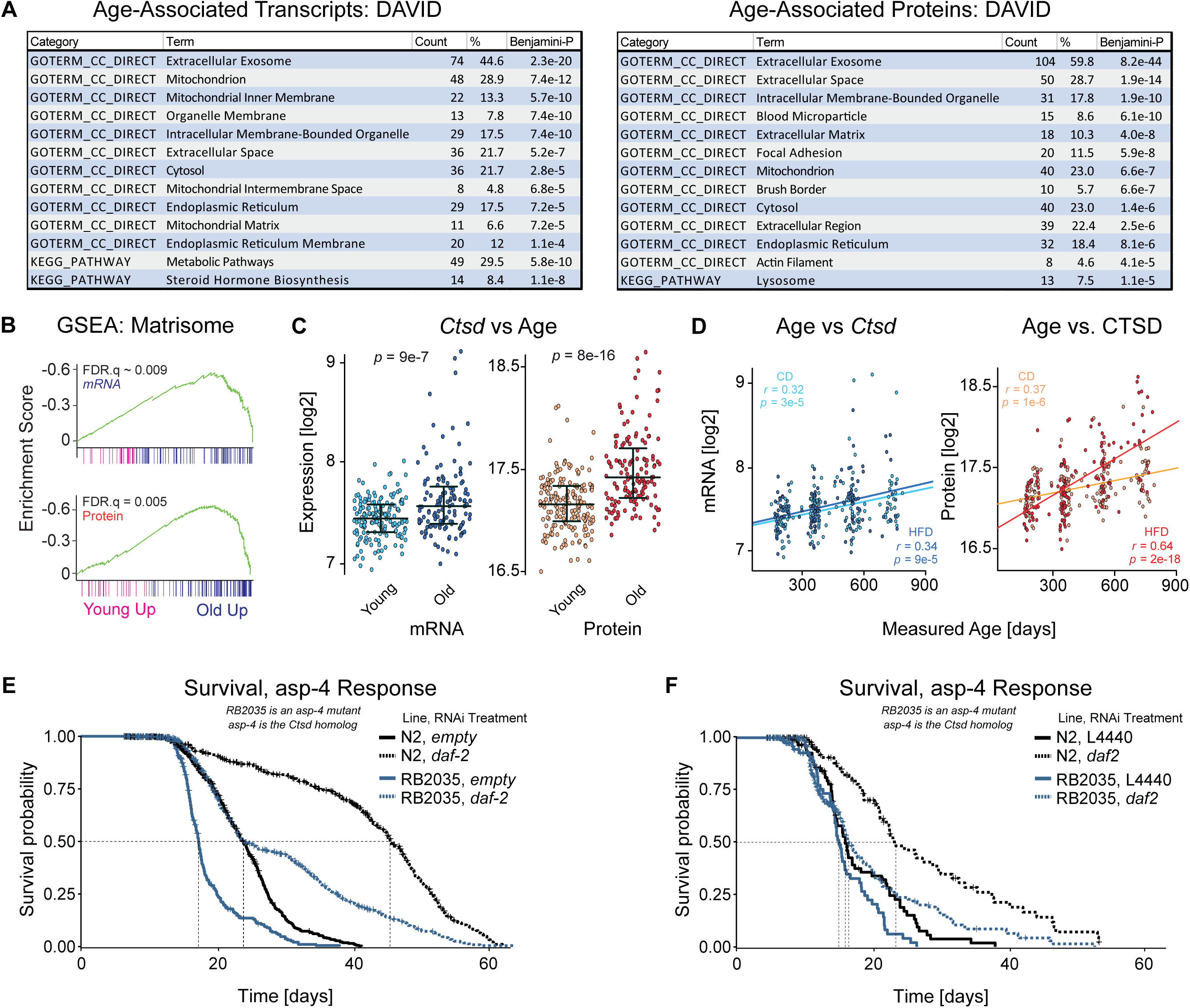
Aging Candidate Discovery & *C. elegans* (**A**) DAVID analysis of the top transcripts and proteins which correlate with the measured age of the mouse when the tissue was taken. (**B**) GSEA for the matrisome gene set (a superset of ECM genes) showing a large enrichment with age for both mRNA and protein. (**C**) *Ctsd* gene expression as a function of age, using a bimodal cutoff. Note that the age groups are relatively discrete. (**D**) Correlation plots of age versus *Ctsd* and CTSD, showing a significant correlation with age regardless of dietary conditions. (**E**) Longevity analysis of RB2035, a mutant *C. elegans* with the removal of the *Ctsd* homolog *asp-4*, compared to wildtype N2 *C. elegans*. (**F**) Repeat of experiment in panel E.

With these gene candidates and the enriched gene sets in mind, we looked for candidate longevity genes which could be tested for causality in *C. elegans* (**Table S4**). Approximately 52 of the top aging-associated genes (or 17%) had a clear *C. elegans* ortholog according to WormBase [40]. Among these 52, *Ctsd* was highlighted as a gene of interest due to its dual involvement in both the lysosome process (i.e. protein degradation) and ECM (it specifically targets ECM proteins, [41]). The single *Ctsd* ortholog, *asp-4* (BLAST p = 1e-107, score=386), has not been examined for longevity in the *C. elegans* literature. The *Ctsb* ortholog *W07B8.4* has been shown to increase in expression with age and to affect reproductive aging, but not lifespan, in both wildtype worms and long-lived *daf-2* mutants [42, 43]. Literature in mammals has also shown an increase in *Ctsd* expression with age, although such studies have been largely focused on brain tissues [44]. In our study, we observed significant increases in *Ctsd* as a function of age in mRNA and protein data both categorically (**Figure 3C**) and in a quantitative correlation with age (**Figure 3D**). We hypothesized *asp-4* may affect longevity, and that like *W07B8.4* it could have an interactive effect with *daf-2*, a gene whose knockdown leads to decreased protein turnover (around 30%) and large increases in lifespan (more than 50%) [45]. We found that RB2035, the *asp-4* mutant *C. elegans* line, has a significant decrease in lifespan compared to wildtype when both were on empty vector treatment (L4440), with a median lifespan of 17.2 days, versus 23.7 days for control (p = 7e-27, **Figure 3E**). Reducing insulin/IGF-1 receptor signalling via adulthood-specific *daf-2* treatment resulted in the expected doubling of the lifespan (45.1 days, p = 2e-95) in wildtype animals. In the RB2035 background, this effect was much reduced, with the same median lifespan as controls (23.7 days) although with a far longer lifespan tail, resulting in an overall lifespan extension (p = 1e-15). These patterns were again confirmed in a second experimental replicate (**Figure 3F**; details in **Table S5**). Together, these experiments indicate that molecular signatures of aging can be used to identify both biomarkers and causal mechanisms that influence longevity. With this concept in mind, we set out to establish how we can generate targeted hypotheses about how diet, age, and genetic interactions drive divergences in key metabolic pathways by using the large multi-omics dataset.

### Using gene-environment-age interactions to understand liver physiology

In addition to the pathways detected by DAVID of top age-associated correlates, we hypothesized that other core metabolic pathways may have modifier genes which are GxExA-dependent and which can be used to understand the molecular basis behind physiological changes in the population. To identify these modifier genes, and their causal relationships with target pathways, some prior hypotheses are necessary to limit the possible search space for our recently-developed causal inference system [15]. Thus, we selected 23 gene sets from GSEA [36] which literature reports link to at least one of our independent study variables of diet, age, or BXD genotype (**Table S6**). A further 2 “false” gene sets were also selected: one of entirely random genes, and one of random metabolic genes. Prior hypotheses are detailed in **Table S6**, including e.g. that CYP450 gene family is downregulated in HFD-fed individuals due to a reduction in plant-based xenobiotics [46], oxidative phosphorylation (OXPHOS) subunits are downregulated in aged individuals [47], and DBA/2J genetic variants upregulate supercomplex assembly in the electron transport chain [48]. Additional pathways could be selected, but we limited this initial input set to 25 to reduce multiple testing. To reach the causal step, we initially focused on two avenues of association analyses: (1) identify molecular coexpression networks which are significant for our data types (i.e. mRNA, protein) and independent variables (e.g. for both dietary cohorts), and (2) examine molecular signatures that diverge across data type or independent variable. All pathways had significant levels of genotypically-induced expression variation as expected due to the selection criteria applied (i.e. **Figure 2B**). 22 pathways had significant protein coexpression networks, and 17 pathways had significant coexpression at the mRNA level (**Table S6**).

Given the discrepancies between mRNA and protein behavior for complexes (**Figure 2G**), we first examined the OXPHOS pathway (called “Respiratory_ Electron_ Transport”) as it is composed of large protein complexes, is known to decrease as a function of age [49], and has variant complex assembly across the BXDs due to genetic variants in *Cox7a2l* [18]. Gene expression of both OXPHOS mRNA and protein corresponded to strong correlation networks (p < 1e-4, **Figure 4A**), but no correlation was observed between the mRNA and protein expression networks (p = 0.54, **Figure 4B** and **Figure S3A**). Furthermore, two key substructural elements of OXPHOS were evident. Uniquely at the mRNA level, the mitochondrially-encoded OXPHOS subunits (red-highlighted triangles, **Figure 4A**) were distinct from the nuclear-encoded primary cluster. Uniquely at the protein level, each complex of OXPHOS formed a distinct subnetwork within the overall structure (**Figure S3B**), and with no difference observed between nuclear and mitochondrially-encoded genes (**Figure 4A**).

**Figure 4.**
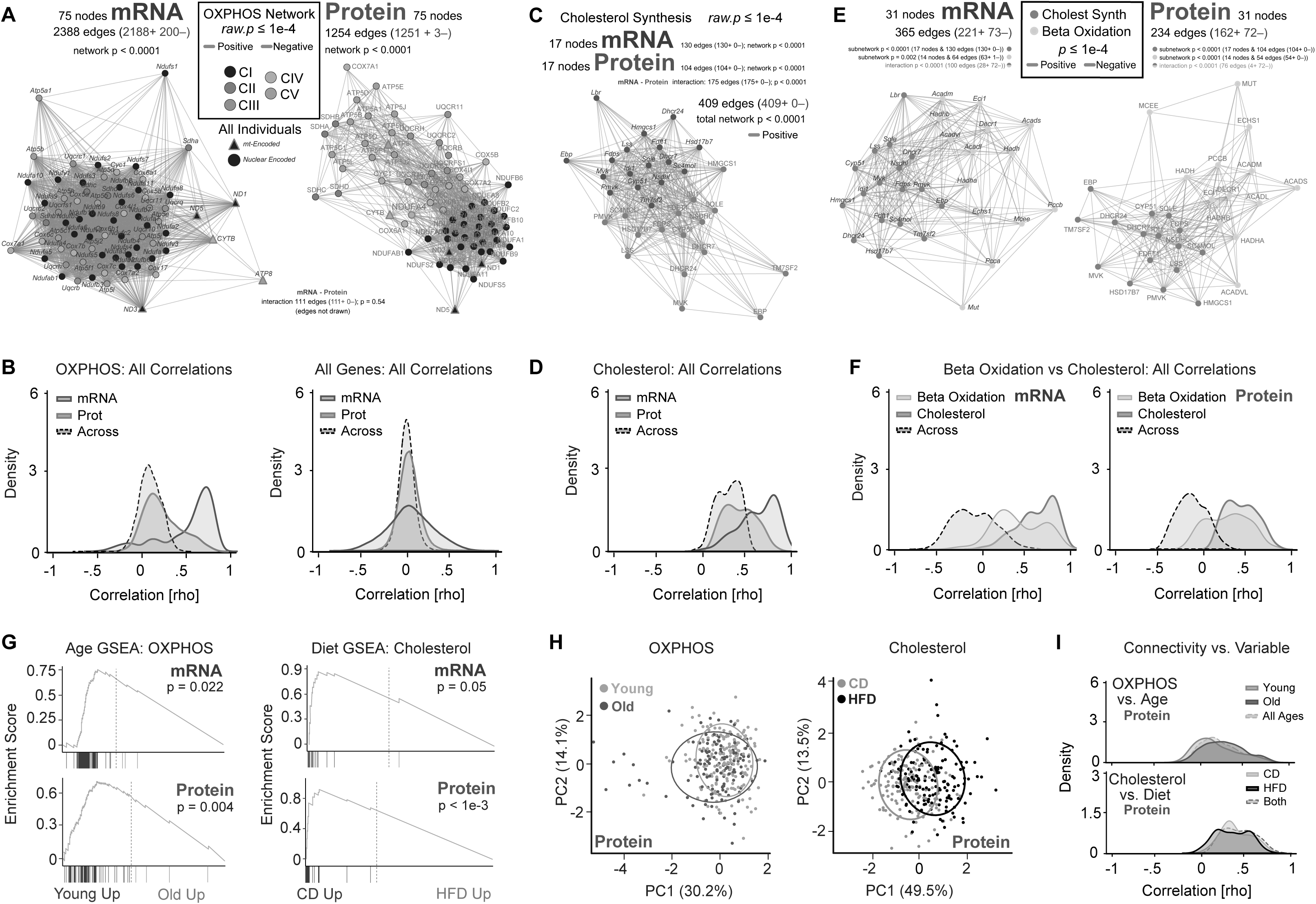
**Functional Gene Networks of Transcripts and Their Proteins** (**A**) OXPHOS Spearman correlation networks for the 75 genes with both mRNA (left) and protein (right) measurements. Node color represents to which component of OXPHOS the gene belongs. Red-highlighted nodes are mitochondrially-encoded. NDUFA4 is a Complex IV member, despite its gene symbol [76]. (**B**) Spearman correlation density plot corresponding to (A), now showing all 5550 correlations for mRNA and protein networks (i.e. 75∧2 minus identity) and “Across” for the 11000 correlations in the mRNA–protein correlation network (i.e. all possible connections between nodes where one node is an mRNA and the other is a protein). (**C**) Spearman correlation network for both mRNA (blue) and protein (red) in the cholesterol biosynthesis network. (**D**) Density plot corresponding to panel C. Of all 1088 possible edges (272 within mRNA and within protein, and 544 across), 409 are significant at p < 0.0001 (i.e. the edges drawn in panel C). (**E**) Spearman correlation network for the cholesterol biosynthesis and beta oxidation gene networks for mRNA (left) and protein (right) drawn together. (**F**) Spearman correlation density plot for the above graphs. “Across” means within mRNA and protein, but across beta oxidation to cholesterol nodes. (**G**) GSEA enrichment for the OXPHOS and cholesterol gene sets as a function of age for mRNA and protein levels. (**H**) PCA plot of the first two principal components of the OXPHOS and cholesterol protein pathways as a function of age and diet, respectively, visualizing the moderate, significant, separation by these two variables. (**I**) Correlation density plot of OXPHOS versus age and cholesterol versus diet.

We next examined the cholesterol biosynthesis process, a comparatively linear molecular pathway of enzymatic reactions driven by individual genes, rather than protein complexes like OXPHOS. Here, the two layers of gene expression were strongly correlated both within expression type, and across from mRNA to protein (**Figure 4C–D**). When we added the beta oxidation pathway to this same analysis, we observed a strong negative correlation between it and cholesterol biosynthesis for both mRNA and protein (**Figure 4E**), indicative of their complementary underlying functions [50]. As for cholesterol, beta oxidation yielded analogous networks for both its mRNA and protein (**Figure 4F**). While gene sets of metabolic pathways that are not predominantly protein complexes tend to have significant correlation between mRNA and protein (i.e. **Figure 2G**), this relationship is always weaker than the within-layer correlation (e.g. beta oxidation, TCA cycle, **Figure S3C**). That is, transcripts in any pathway are closer to other transcripts in the same pathway than they are to their matching proteins. To examine the consequences of divergence between mRNA and protein on subsequent hypothesis discovery and deconvolution of the causes of variation in phenotypes and gene expression, we examined the impact of our independent variables of diet and age on each of the selected functional gene networks. 17 gene sets were affected by diet or age at the mRNA and/or protein level, a predictably high overall enrichment given that the sets were selected with diet and age hypotheses in mind (**Figure 4G, Table S6**). While age and diet had major impact on these pathways, genetic variation across strain still played the largest role, which precluded a simple categorization of any given animal into an age or diet purely based on PCA of a single gene set (**Figure 4H**). Furthermore, even with a nearly uniform downregulation in some gene sets, e.g. cholesterol biosynthesis induced by HFD, the overall network connectivity remained similar (**Figure 4I**). This suggests that the genetic mechanism driving the network’s response to the causal study variables (e.g. diet) may lie outside of the canonical gene sets. We thus set out to expand the search space of these gene sets to identify interactors and effectors using stability selection.

### Data-Driven Approaches to Non-Consensus Networks & Causal Inference

Functional gene ontologies provide a crucial platform for moving between data-driven hypothesis generation and mechanistic molecular studies. However, gene set annotations necessitate arbitrary cutoffs for categorization, as metabolic pathways are subsets of larger sets of interconnected genetic mechanisms. The majority of the genome still remains relatively unexplored in the literature [51], and data-driven approaches can identify gene functions and their associated pathways or diseases, including by building off of reference gene sets [52]. Confirmation of data-driven results depends on painstaking molecular biology validation and systems biology approaches that cross-validate associations across different conditions and/or independent studies. Meta-analyses tend to be consensus-driven [53], which is critical to reduce the perennial issue of false discovery in systems biology, where the number of response variables *p* (e.g. all transcripts measured by RNA-seq) inevitably exceeds the measured number of samples *n* (i.e. p ≫ n). Additionally, many gene functions may only be evident in a certain tissue, or under certain experimental conditions. Thus, we hypothesized that the variations observed in our canonical functional networks may be affected by “unstable” gene partners which are only evident in particular study conditions (e.g. aged or HFD-fed individuals). Instability in these relationships can be causally driven by our independent variables of diet, age, and genotype, which would in turn permit their causal relationship with the canonical metabolic pathway. To determine the relationship of unstable genes to our pathways, we applied a machine learning technique which we recently developed for gene expression studies which compares the effects of multiple independent variables on a target network [15]. This method allows for a linear regression-based variable selection (similar to lasso regression [54]) and is combined with stability selection [55] which uses resampling to control for false discoveries. This significantly reduces false discovery compared to correlation networks or hierarchical clustering. Furthermore, the method performs a causal analysis, similar to one we developed previously [16], to assess if strongly-associated candidate genes mediate the effect of a secondary independent (e.g. if strain is the primary independent, diet or age is the secondary). Together, the regression analysis should ensure that discovered target genes are proximal to the target pathway, while the stability analysis can indicate whether non-consensus connections are due to the nodes being upstream, downstream, or inside the target network.

Here, we processed our same 25 gene sets to identify potential novel associating genes for all 3772 genes measured at both the mRNA and protein level. The analysis first uncovers novel genes which associate (regress) with a target pathway across genotypes, such as cholesterol biosynthesis (**Figure 5A** for protein, **Figure S4A** for mRNA). The stability selection analysis then analyzes how the connectivity between the newly-identified genes changes according to a secondary segregating—and causal—independent variable such as diet (**Figure 5A**, left vs. right networks). Novel nodes that are similarly positioned in both networks are retained (e.g. *Rdh11*), but causal directionality with the target network cannot be inferred as it does not diverge according to the secondary variable (in this case, diet). Genes with diverging connectivity, such as *Obscn*, can have causal association inferred. Both RDH11 and OBSCN proteins are significantly *and causally* downregulated by HFD (**Figure 5B**), but only OBSCN has diverging connectivity. Causal analysis then informs us that OBSCN is statistically upstream of the effects of HFD on cholesterol biosynthesis (as are other genes in the upper-left quadrant like FABP2, ACAT1, CYP2C29, MAVS, CYP2C39, HMGCS2, and ALDH1L1, **Figure 5C**). That is, an intervention directly affecting cholesterol biosynthesis should not impact these proteins, but targeting these proteins should have an effect on genes in the cholesterol biosynthesis pathway, possibly only under certain conditions (e.g. HFD conditions for OBSCN). In this case, no genes were identified that were proximal to and statistically downstream of cholesterol biosynthesis (genes in the empty lower-righthand quadrant).

**Figure 5.**
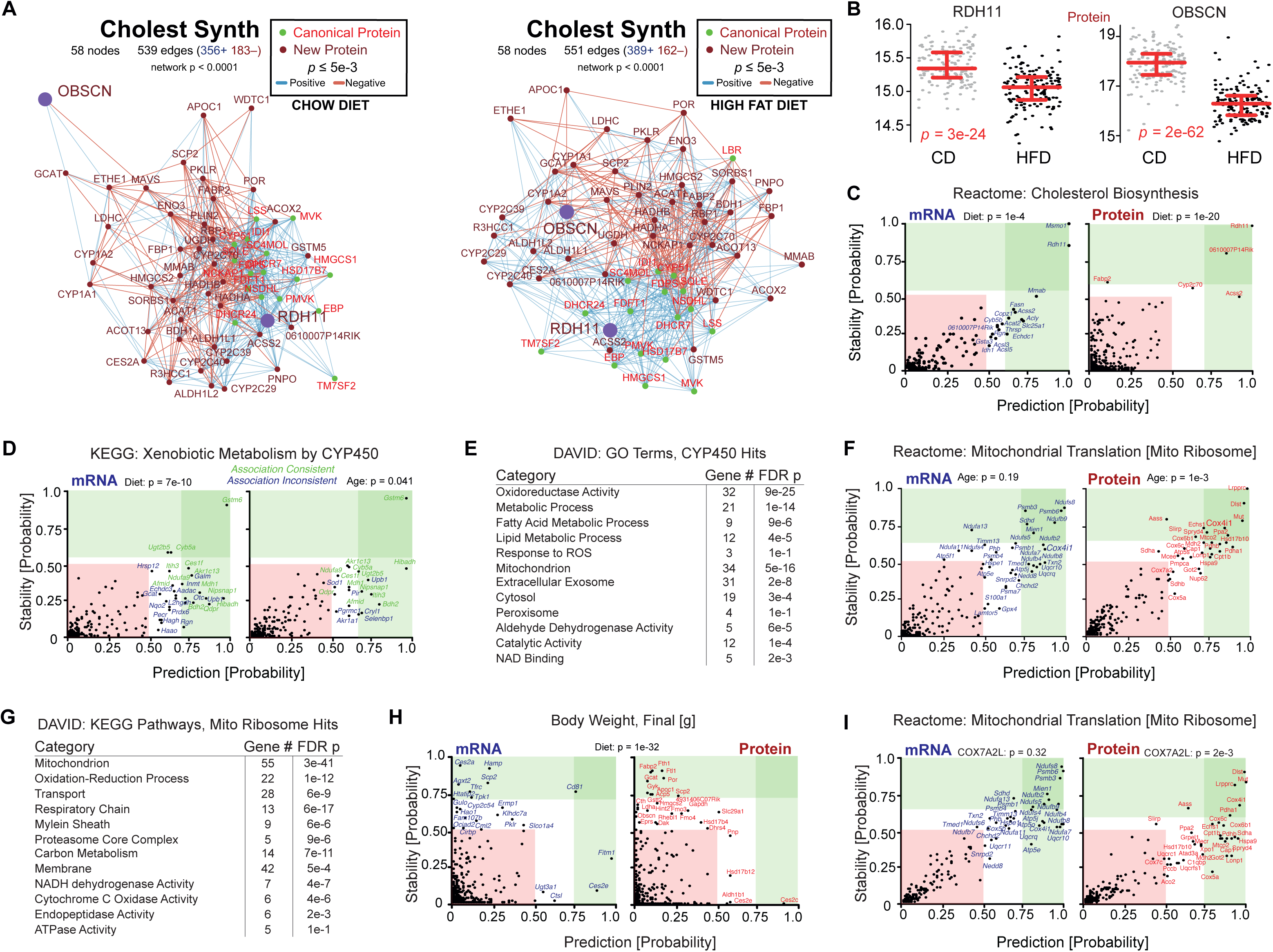
**Network Expansion and Functional Gene Discovery** (**A**) Spearman correlation plot of the core cholesterol biosynthesis network (17 red nodes) along with 37 candidates selected by regression analysis in protein data, performed separately for CD (left) and HFD (right) cohorts (dark red nodes). (**B**) Dot plot of RDH11 and OBSCN as a function of diet; note that although both have large diet effects, RDH11 is centrally connected in both dietary networks, but OBSCN is peripheral in CD. (**C**) Prediction–Stability plot for the cholesterol biosynthesis pathway in mRNA and protein as a function of diet. (**D**) Prediction–Stability plot for the CYP450 pathway showing transcript hits as a function of diet or age. (**E**) DAVID enrichment analysis of the top candidate genes found through stability analysis of the CYP450 pathway. (**E**) Prediction–Stability plot for the mitochondrial ribosomal pathway for mRNA and protein as a function of age. (**F**) Prediction–Stability plot for the mitochondrial translation pathway for mRNA and protein as a function of age. (**G**) DAVID enrichment analysis of the top candidate genes found through stability analysis of the mitochondrial ribosome pathway, showing that most hits are already known to be mitochondrially localized. (**H**) Prediction–Stability plot for the effect of HFD on body weight, showing a massive skew away from the X=Y axis, due to the causal impact of HFD on body weight. (**I**) Prediction–Stability plot for the mitochondrial translation gene set as a function of the COX7A2L allele, which is expected to have a downstream impact but only in protein data [18]. The average hit falls within |0.21| units of distance for mRNA, versus |0.33| units for protein, which is expected as the *Cox7a2l* allele is known to have a much stronger effect on OXPHOS protein organization than on the equivalent mRNA networks.

Causal analysis was done on all 25 pathways (**Table S7**) according to both diet and age and for both mRNA and protein. Candidates were retained as potential “hits” if they had either prediction or stability scores ≥ 0.50, which corresponds to a permutation-based false discovery of approximately 1 node per gene set and test (e.g. **Figure S4B**, each plot is the overlaid sum of 100 tests using randomized labels). At these cutoffs, we detected an average of 23 gene candidate hits per mRNA network and 19 per protein network, for a total of 2101 associations belonging to 748 distinct genes. Candidate genes appear multiple times, as they can be associated with multiple independent gene sets and as a function diet, age, and mRNA/protein measurement type. 450 of the 2101 total associations—around 21%—had clear directional relationships to their target pathway (prediction or stability score of ≥ 0.50 and at least 0.30 units away from the X=Y linear axis). Roughly 44% of hits were found consistently whether segregating by diet or age, but this varied substantially between pathways. 47% of the 34 novel CYP450-related mRNAs and of the 62% of the 21 novel CYP450-related proteins were consistently identified across age and diet (**Figure 5D** **& S4D**) versus 69% of the 29 novel peroxisome-associated mRNAs (**Figure S4C**). Gene associations identified using mRNA and protein diverged more substantially, with no more than 18% of discovered genes consistently associated between the gene and target pathway in both mRNA and protein data (**Table S7**). For instance, only 2 novel genes (4%) associated with the CYP450 pathway as a function of diet consistently in both mRNA and protein (*Ugt2b5* and *Gstm6*, **Figure 5D** **& S4D**).

To determine the relevance of these findings, we used DAVID [34] to determine general relationships between hit genes and the canonical target pathway. For CYP450, 70 distinct genes were picked up across mRNA, protein, and diet/age networks, of which 32 were associated with the “oxidoreductase activity” (adj.p = 9e-25, **Figure 5E**). Of these target genes, many have clear functional interactions with CYP450, such as two genes in glutathione metabolism (*Gstm6* and *Gstm7*) and four carboxylesterase genes (e.g. *Ces1e*). However, at least two dozen candidate genes have no clear known connection with CYP450 or any proximal pathway, such as *Nipsap1*, *Afmid*, or *Tmem205* (all hits are detailed in **Table S7**). Similar patterns were seen with other gene sets. For instance, the mitochondrial translation gene set (Reactome, M27446) yielded 71 candidate genes. Only one was found in common to mRNA and protein searches: *Cox4i1* (**Figure 5F** and **Table S7**). Among those 71 candidate genes, 55 were associated to the mitochondria and 13 to the respiratory chain (**Figure 5G**). The remaining 16 candidate genes, including 5 proteasomal genes, had no established functional or positional connection to the mitochondria (**Table S7**), such as *Mien1, Nedd8*, and *Tmed1*.

We next considered two potential analytical extensions to stability inference. First, the target variable does not have to be a gene set; it can be any variable(s) expected to have common causal and response factors. We first examined overall body weight as a function of diet (**Figure 5H**). As gene expression cannot affect diet in this study (i.e. taste preferences are not a factor), diet has a large unidirectional and causal impact on body weight (p = 1e-32), and diet drives nearly half of all variance in body weight (**Figure S1C**). Thus, we should expect to see body weight-associated genes lie far away from the X=Y axis as a function of diet, but not age. Indeed, we observe that among the 94 candidate transcripts or proteins related to weight as a function of diet, 95% have predictions of causal direction. Conversely, only 41 transcripts or proteins were associated with body weight as a function of age between 6 to 24 months (which has no uniform on body weight, p = 0.32), with 23% yielding causal information (**Figure S4E, Table S7**, sheet “BodyWeight”). The second additional possibility that we considered is to use a causal independent variable other than diet and age. The genotype axis can be subdivided: every gene has two possible alleles (i.e. “B” or “D” from the **B**-X-**D** population). Due to the huge size of such a search space, we cannot check how every gene network diverges as a consequence of every variant allele. As with selecting gene sets of interest using prior knowledge, we must select genetic targets of interest on which to separate using starting hypotheses. The BXDs have two diverging alleles of *Cox7a2l,* where a *cis-*acting variant leads to major changes in protein organization of OXPHOS complexes [18]. We examined causal associations with the mitochondrial translation (ribosome) gene set, which should be generally upstream of OXPHOS, particularly as a result of this specific *Cox7a2l* variant which affects supercomplex stability. As expected, analyses discovered many logical interacting genes with the mitochondrial ribosome: predominantly OXPHOS subunits and UPR^mt^ regulators (**Figure 5I**). Moreover, the mRNA hits are found near a line where X=Y, whereas protein hits are shifted further away. This is expected, as variant alleles of *Cox7a2l* are known to affect gene networks and organization of oxidative phosphorylation complexes—but only at the protein level [18]. That is, the *Cox7a2l* allele only provides causal effects on protein-level data. If the same input data are used, but strains are instead segregated by their allele of a functionally unrelated gene, such as *Hmgcs2*, gene hits are instead near the X=Y axis (**Figure S4F**). Thus, as with the associations between body weight and gene expression, the same input data yield different hit genes and differing estimates of causal associations depending on the separating variable—and causal information is only yielded in cases where the separating variable (e.g. diet) has an impact on the target trait (e.g. body weight).

## DISCUSSION

Aging is a dynamic process driven by a complex longitudinal mixture of genetic predestination, environmental effects, stochastic processes, and their interactions (GxE). Despite the relatively high heritability of longevity and wealth of knowledge about aging, much remains unknown about molecular causality even for well-studied aging processes such as mitochondrial stress, telomere shortening, and DNA methylation. Greying hair and shortened telomeres have strong, clear associations with age, but it remains a challenge to causally determine whether a hypothetical telomere-lengthening treatment would improve lifespan any more than does black hair dye. Even when causality with lifespan has been shown, such as the effect of CR on lifespan, it is essential to deconvolute the effects of genetic background. Indeed, in mammals, CR has been causally shown to both shorten *and extend* lifespan, depending on genetic background [56, 57]. These phenotypic effects are highly reproducible, indicating that variant molecular mechanisms activate only be evident under certain genotypes and environments. Here, we provide a large, novel multi-omics aging dataset and demonstrate how multivariate experimental designs can be combined with causal data analysis strategies to examine longstanding questions in how molecular factors vary and cause complex traits as a function of GxExA.

We measured the transcriptional, proteomic, and metabolomic landscapes of livers from 300 different cohorts of the BXD mouse population as a function of age, sex, strain, or diet. Genetic differences alone explained ∼28% of variation for all molecular measurement layers, dwarfing genetically-independent effects of diet and age (< 5%). However, diet and age do have a substantial effect: interactions between genotype, diet, and age explained a further ∼34% of variance. Rather, diet and age have a striking impact on molecular expression, but they are highly dependent upon their interactions with the genome. This suggests that the unitary effect of any single factor will progressively diminish with additional environmental variables while interaction effects will increase—an inevitable complication for the study of aging in natural populations. That is, once a population is diverse enough for its genes and its environment, even genes that are strongly modulated by environmental factors (e.g. CYP450 genes by specific dietary compounds) will have their effect linked to any one single factor gradually diluted. This observation raises a critical converse: studies using single, fixed environmental variables will struggle to extrapolate their findings to a general population—especially if this is compounded by the presence of additional hidden variables as is inevitable for the study of age (as we have seen for CR). In order to demystify the heritable molecular factors driving complex traits, we must consider study designs with multiple, simultaneous, independent variables to measure, and then decipher, the interactions.

Over the past decade, it has become increasingly recognized that analyses using mRNA or protein measurements often yield substantially different conclusions, even when measured at the same time in the same samples. That is, transcripts are not merely an intermediate proxy for protein. We observe here that a typical transcript and protein respond across complex GxExA stimuli with only moderate concomitance (rho ∼ 0.14). This general conclusion has numerous exceptions, e.g. factors leading to large effect sizes changes at the transcriptional level are far more likely to replicate at the protein level. Conversely, genes in complexes are far less likely to have shared differentiation in response to a stimulus even if highly variable according to GxExA. We have examined our multi-omic dataset to uncover candidate genes related to age and uncovered a few novel aging candidates from different pathways. One gene, *Ctsd*, has a clear *C. elegans* ortholog, *asp-4*, which we discovered reduces lifespan when knocked out in both control worms and in long-lived *daf-2* worms. However, relatively few candidate genes stood out as varying strongly with age. Indeed, numerous aging studies have now shown that only a few genes have large effect sizes (e.g. > 2-fold) as a consequence of lifespan [25, 26], and these may not be the most important genes in terms of causality. A 2-fold change is relatively “moderate” for variant expression in a single gene, but we must consider changes in entire pathways: a 2-fold change in expression of the entire OXPHOS pathway is a striking impact, as is a 2-fold change in a phenotype such as exercise capacity, insulin response, or lifespan. Consequently, we must design studies keeping in mind that 99% of gene variants will vary by less than 2-fold across age. Fortunately, it is now possible to quantify the transcriptome and proteome across hundreds of samples with sufficient precision to detect small fold changes [58]. This improves our ability to detect smaller effect sizes, but more importantly opens new avenues in biostatistics.

Differential expression analysis benefits from larger sample sizes, but in practical terms the returns quickly diminish for now-traditional bioinformatic approaches like volcano plots or GSEA. While an *n* of 10,000 transcriptomes or proteomics would allow tiny effect sizes to be determined “significant”, such effects are of limited translational utility. Scientists are obligated to find new statistical rationales for generating these larger datasets. This requires fundamentally adapting experimental design, rather than simply upscaling studies focusing on single axes of variation. Recent translations of machine learning advances to biology have become popular, especially for unsupervised analysis of genome data and imaging, but these algorithms tend to require sample sizes that are still orders of magnitude larger than are realistic for gene expression studies (e.g. n > 100,000), resulting in a “p ≫ n” (or “big-p, little-n”) problem. Furthermore, such approaches tend to search for consensus across input datasets [59]. However, differences in molecular networks as a consequence of different independent study variables can also indicate the causality behind pathway variance; a true association connecting a node (e.g. a transcript) to body weight gain in animals fed HFD may not be evident in animals fed CD. These inconsistent, or unstable, edges need to be retained, examined, and ranked, as we have shown in this study. Addressing these issues requires substantial knowledge of biological priors. Due to the difficulty of collecting tens of thousands of tissue samples, let alone processing multi-omics on them, even in the next decade, expression studies will remain far below of the sample size necessary for the “classical” unsupervised machine learning algorithms. Here, we have applied a stability inference algorithm that we recently developed [15] which takes advantages of two aspects of this study design. First, the study’s three independent variables (i.e. genetics, diet, age) permit stability analysis. That is, correlation networks and regression analyses can be performed across the primary axis of genetics, and discrepancies in network connectivity as a function of diet or age can be not only quantified as by node centrality analysis, but also it can be inferred whether the discrepancy is *due to* the intervention, or if it is a response. Second, the study’s acquisition of both mRNA and protein data provides for a second “type” of consensus: results that are consistent across mRNA and protein gain improved confidence, while results that are inconsistent can be flagged according to certain criteria (e.g. presence of target gene in a protein complex). We have used this uniquely large and complex aging dataset to identify novel associations between molecular pathways, genes, and age and diet, and for a significant fraction of these candidates (∼21%), their causal associations.

Altogether, this dataset and method demonstrate how simultaneous applications of multiple independent variables can be used for hypothesis discovery. Concurrent results from a multivariate study provide increased confidence in consensus. More critically, if the segregating variables are known to be causal, associations which are condition-dependent can in some cases be statistically shown as upstream or downstream, rather than results considered ephemeral and inconsistent, then discarded. This requires a study design which is at present fairly unusual and which we aim to popularize with this work: designing studies with multiple independent variables in a full (or nearly-full) fractional design [60], and with one axis sufficiently deep to allow the calculation of significant correlation networks. With these developments, we can move hypothesis discovery in data-driven studies from correlation networks to causal pathways.

## METHODS

Supplementary figures, tables, code, and further methods are available in the online version of the paper.

### Mouse Care and Handling

All animal care was handled according to the NIH’s *Guidelines for the Care and Use of Laboratory Animals* and was also approved by the Animal Care and Use Committee of the University of Tennessee Health Science Center (UTHSC). 2157 mice from 89 strains of the BXD family (including parents and both F1s) were followed across their lifespan. 159 animals were males and 1998 animals were females. Animals were maintained in the UTHSC vivarium in Specific Pathogen-Free (SPF) housing throughout the longevity experiment. The housing environment was a 12-hour day/night cycle in 20–24°C temperature with housing cages of 145 in^2^ with up to 10 animals per cage. Diets were either Harlan Teklad 2018 (CD; 18.6% protein, 6.2% fat, 75.2% carbohydrates) or Harlan Teklad 06414 (HFD; 18.4% protein, 60.3% fat, 21.3% carbohydrates). Water was Memphis city municipal tap water. Food and water were *ad libitum*. All animals were followed from their point of entry into the colony (typically around 4 months of age) until death. Animals were checked daily for morbidity and were weighed approximately every 2-3 months throughout their lives. 662 animals were sacrificed at specific ages for tissue collection across cohort (i.e. diet, strain, sex, and age) while all other animals lived out their natural lifespans. For animals living out their natural lifespan, ∼90% died naturally while the remaining ∼10% were euthanized according to AAALAC guidelines and made by an independent veterinarian at the UTHSC facility. Euthanized animals were retained for lifespan calculations, with the expectation that they would have otherwise died shortly thereafter.

Of note: on April 28, 2016 all mice were moved from the study’s major housing facility (“Nash”, which was slated for demolition) to a new building (“TSRB”). By this point, 94% of sacrificed individuals had been born, raised, and sacrificed in the Nash facility, so only 6% of individuals processed for omics analysis were moved, all of which were sacrificed between 1 September and 26 October 2016, i.e. after 4 to 5 months of acclimatization.

### Aging Calculations

Lifespan calculations and significance tests were made using the “survival” package on R with the survfit and survdiff functions. 1495 deaths were recorded, of which 1386 were female. 40 of these female deaths were suppressed prior to lifespan calculations for various reasons, e.g. 7 mice died due to flooded cages, 2 animals were accidentally entered at far too old an age (>1.5 years), 2 mice were found with broken limbs, 6 were sacrificed for an urgent revision for an unrelated paper, 3 mice died before the average age of entry into the colony (7 months), and the rest were removed by the veterinarian for non-definitively-aging related reasons (e.g. significant seizures noted during body weighings). The 662 animals which were sacrificed for this study’s tissue collection aim were not used for lifespan calculations.

### Cohort Sacrifice Selection

Animals were selected for tissue harvest with the following aims: 2 animals per strain, diet, and age, for a target of 4 age points, i.e. up to a target maximum of 16 sacrificed animals per strain (2 replicates * 2 diets * 4 ages). In the final sample collection database, an average of 11 animals were available per strain (60 strains, 662 animals). The target ages were 7, 12, 18, and 24 months of age. Roughly every 3 months for the duration of the experiment, ∼40 animals were selected for sacrifice, with approximately 15 animals sacrificed per day over the course of 3 or 4 continuous days. Animals were removed from the aging colony the night prior to sacrifice, but retained access to food and water. Sacrifices started at approximately 9am with the anesthetic Avertin used via intraperitoneal injection of 0.2 mL per 10 g of body weight. Animals were perfused with ice cold phosphate-buffered saline. The liver was the first organ harvested. The gall bladder was removed, the liver weighed, and then immediately frozen in liquid nitrogen in 20 mL scintillator vials. When reporting the number of strains analyzed for each part of this study, we count the two F1s—B6D2 and D2B6—and the parental strains. Although F1 hybrids are not “inbred strains”, they can be reliably and reproducibly generated to provide biological replicates, and thus can be used as reliably as inbred strains for studies on gene-by-environment interactions. C57BL/6J and DBA/2J are counted as “BXD strains” for simplicity, although like the F1s, they do not help with QTL in the context of this study. A more detailed breakdown of BXD genetics has been published recently [61].

### Time Points for Aging Calculations

While CD and HFD comparisons were binary, comparisons across age were somewhat more challenging. For some analyses, linear regression was used for time at sacrifice (or measurement) against the target variable. However, for other analyses, particularly GSEA, age-QTLs, and causal inference, discrete bins were used. For binning animals based on age group, animals were binned by absolute age—i.e. not normalized by strain lifespan—with young mice considered those sacrificed before 419 days of age, and old mice considered those beyond 431 days of age, with a mean±σ of 293±80 vs 615±97, respectively. Note that the bins are more distinct than the standard deviations suggest as the age distribution for sacrificed individuals is not normal; e.g. only 12 animals were sacrificed between the ages of 400 and 500 days; see **Table S1** or **Figure S1D** for more details.

### Transcriptomics

RNA was prepared using TRIzol reagent and cleaned up with RNEasy MinElute kits (Qiagen). RNA-seq and RNA integrity (RIN) checks were performed by NovoGene on a HiSeq PE150. Samples with RIN ≥ 6.0 were retained for RNA-seq. All samples were run with 20 million reads. Normalization was performed by ComBat. All RNA-seq data were scaled by adding 1 to the normalized counts and then taking the log2. 127 genes were removed from paired analysis due to the measurements being > 50% “0” at the mRNA level. Transcripts with more than half zeroes were not considered for mRNA–protein correlation due to this high number of matching values which throw off Spearman correlations. A further 38 genes had up to 100 counts of 0 which have lower than average (rho = 0.05) correlation with their transcript. Of these 163 genes, twelve of these genes were significantly affected by diet, and one by age, while all genes had significantly lower than average correlation with their protein level (rho = 0.05). Good data could potentially be contained for these transcripts, but given the preponderance of noise, we have discounted all mRNAs with more than half read counts of zero. All measurements, including those with high “0” counts, are included in **Table S2**. Note that for multi-omics analyses we only used the 3772 genes with overlapping mRNA and protein data, but all RNA-seq data for all transcripts is included in **Table S2**.

### Proteomics

A detailed step-by-step protocol has been published for the sample [62]. In brief: liver samples were first entirely pulverized by mortar and pestle in liquid nitrogen, proteins were then extracted from ∼20-50 mg of powered liver in 750 µL of RIPA-M buffer. The remaining cell pellet was then lysed fully in 8M of urea. The fractions were combined and 100 µg of each sample precipitated with acetone overnight at −20°C. The precipitated sample was resuspended in urea, reduced with dithiothreitol, and alkylated with iodoacetamide. Samples were diluted to 1.5M urea and then digested overnight (22 hours) using modified porcine trypsin. Peptides were cleaned C18 MACROSpin plates (Nest Group). Roughly 1.5 µg of each peptide sample was loaded onto a PicoFrit emitter on an Eksigent LC system coupled to an AB Sciex 6600 TripleTOF mass spectrometer and acquired in SWATH data-independent acquisition mode (DIA) [63] with 100 variable windows in a 60 minute gradient. A recent review also provides more detail on the full DIA pipeline [64]. In brief, the resulting .wiff files were converted to mzXML using Proteowizard 3.0.5533 before being run through the OpenSWATH pipeline v2.4.0 [65]. The library used was merged from our prior mouse library [28] together with part of the PanHuman library [66]. All peptides in the PanHuman library were BLASTed against the canonical mouse proteome from UniProt, version downloaded July 2017. Peptides which were found to be proteotypic in mouse, and not already extant in the mouse library, were then merged with the mouse library. Note that all runs for both human and mouse library generation were acquired with on the same machine (a TripleTOF 5600+) with the same settings (detailed in [66]). This merged “PanMouse” library contains 103,644 proteotypic peptides corresponding to 8219 unique proteins. This library was used to search the mzXML files for the OpenSWATH pipeline using the msproteomicstools package available on GitHub. Scoring and filtering were done by PyProphet at 1% peptide FDR, followed by cross-run alignment with TRIC using a max retention time difference of 60 seconds and a target 1% FDR, 29935 proteotypic peptides were identified which corresponding to 3694 unique proteins, were quantified across the 375 retained MS injections. Proteome data were segregated into batches based on noted changes during mass spectrometry (LC column change, MS tuning and cleaning). Within batch, samples were normalized by LOESS followed by across-batch normalization with ComBat, a synthetic approach based off of similar concerns with other large omics datasets. Due to the unexpected complexity of this normalization, we turned this approach and sample QC into a separate publication [67].

### Metabolomics

Pre-homogenized liver samples of were precisely weighed (to a target of ∼20 mg) and then extracted in ∼7 mL of a solution of 40% acetonitrile, 40% methanol, and 20% water, incubated for 24 hours at −20°C. The suspension was then centrifuged, the supernatant transferred to a new tube, and then lyophilized. Dried samples were kept at −80°C until ready to be injected on the mass spectrometer, when they were resuspended in water according to the weight of the input tissue sample to a target of 5 mg/mL. The same extraction of all 621 samples was done in duplicate, starting from the same pre-homogenized liver sample, approximately one month apart, which is detailed by the “Run1” and “Run2” suffix in the data (**Table S2**). Untargeted metabolomics analysis was performed on an Agilent 6550 QTOF instrument in negative mode at 4 GHz, for a mass range of 50–1000 Da using a previously-described protocol [68]. All samples were injected in technical duplicates in both experiments, and nearly all samples were injected in biological duplicates, i.e. each liver sample was injected 4 times to allow measurement stability to be calculated, so most cohorts had 8 measurements (2 full-process replicates * 2 technical replicates * 2 biological replicates (usually) per age-strain-diet cohort). An average of 19,000 features were detected in the runs, of which about 400 could be tentatively annotated as deprotonated metabolites listed in the Human Metabolome Database. Both experimental runs were run approximately one month apart, with all samples in each run being extracted and run in technical duplicate the same sequence both times.

### Hypothesis Discovery, Causal Inference, and Machine Learning Methods

For our network expansion analysis we employed a novel regression and variable selection technique [15], which is optimized for gene expression studies and explicitly allows incorporating causal reasoning similar to a method we recently published [16]. Our preliminary goal was to determine which genes are functionally related to a given pathway. The secondary goal was to determine if the gene was varying as a function of a secondary independent variable (e.g. diet, age), and if it was, then the causal directionality of this association with the target pathway and its sign (positive or negative, i.e. promotive or inhibitory). To this end, we compute the average expression level of the given pathway and use it as a response variable. Then, we randomly sample subsets of the pre-selected predictors and regress each subset onto the response resulting in a single regression coefficient. The individual regression coefficients are finally combined by a weighted average, where the weights are selected to ensure both good predictive performance and stability across the independent variable (diet, age, or a Mendelian QTL). By assessing the predictive performance of these regressions, we then rank the genes by their functional relation to the response (large values indicate a strong functional relation, small values a weak functional relation). For Mendelian separation using COX7A2L and HMGCS2, the 20 heterozygous F1 animals were removed. The selection procedure also uses stability selection [55] in order to control false discoveries and improve reliability. Furthermore, to empirically benchmark the false discovery rate, we perform a permutation based analysis as follows (**Figure S4B**): We apply the entire selection procedure 100 times by permuting the observations of the response variable in each iteration and keeping everything else fixed (and hence preserving the correlation structure between the predictors). This analysis was performed for all the pathways we considered. Note that the gene set “Random75” finds significant correlations—this is expected as there is 75 random genes are using their true expression data, thus true correlations are expected, just they are not expected to be related to any specific pathway.

#### C. elegans Testing

For selecting orthologs, each of the top 100 genes from CD and HFD for mRNA and protein—a total of 300 unique genes as roughly 25% of the top 100 were picked up in at least two conditions—was checked in WormBase for orthologs. Genes with a single annotated ortholog were considered for further analysis. Genes such as *Cyp4f18* matched to dozens of orthologs—essentially the entire family of cytochrome P450 genes—and were thus considered too non-specific for reasonable cross-species analysis, but could potentially be of further use for checking the associations between pathways and aging. To test *Ctsd*, *C. elegans* populations were maintained on NGM plates seeded with *Escherichia coli* bacteria at room temperature, with the exception that temperature-sensitive mutants were incubated at 15°C. For this work the *C. elegans* strains wildtype N2 and RB2035 *asp-4(ok2693)* were used as well as the RNAi clones *L4440* (empty vector control) and *daf-2* (RNAi, Vidal library [69]). Animals were age-synchronized using population lysis [70] and then transferred at the late L4 stage to plates seeded with the selected RNAi clone and containing 50 μM 5-Fluoro-2’deoxyuridine (FUDR). Depending on the genotype, animals were placed at 15°C (N2 and RB2035) to complete their development. After the L4 stage all animals were shifted permanently to 20°C and their survival was quantified. Manual survival scoring (by hand) was conducted as described previously [71]. Briefly, individuals which did not move in response to being prodded were classified as dead. Automated lifespan measurements were conducted using air-cooled Epson V800 flatbed scanners at scanning intervals of 30 minutes as described [72]. For the automated measurements the animals were transferred to fresh plates (BD Falcon Petri Dishes, 50x9mm) at day four of adulthood to facilitate the image detection process by removing as many eggs as possible. The survival data was analyzed using R in combination with the survival (v3.1-12) and survminer (v0.3.1) libraries. In the analysis, all animals which were observed to burrow, undergo bagging, explode or have escaped the agar surface were censored, and the L4 stage was set as the timepoint zero. The survival function was estimated utilizing the product-limit (Kaplan-Meier) approach and the null hypothesis was tested using the log-rank (Mantel-Cox) method.

### Miscellaneous Bioinformatics and Statistical Tests

All functions should work in R 4.0.4. QTL calculations were performed with r/QTL2 v0.22-11 (2020-07-09) [73] using a linear mixed model and a kinship matrix generated by the “leave one chromosome out” (loco) method. Genotypes used were from the 2019 build of the BXD genotypes from GeneNetwork [74]. Transcript QTLs are referred to as eQTLs (“expression” QTLs). Protein QTLs are referred to as pQTLs, and metabolite QTLs as mQTLs. The blood serum measurements are considered as clinical phenotypes rather than as a part of metabolomics or proteomics due partly for historical categorization standards, and also due to differences in the measurement technologies and (in this particular study) source tissue. QTLs were declared as *cis* if their peak LOD region was within ±10 MB of the gene location. All *cis*-QTLs are included in a supplemental data file as their generation is computationally intensive (**Table S3**, sheet 2). The code for generating these QTLs is also included (attached code files, filename b_Figure2_HelperFile_QTLs.r).

Distributions (e.g. **Figure 4B**) were compared using a chi-squared test or Kalmogorov–Smirnov, as indicated. The R package “corrgram” v1.13 was used for generating correlation matrices. Two group comparisons were made by Welch’s t-test. Multiple group comparisons were made by ANOVA with Tukey post-hoc tests. The contribution of each independent variable to trait expression was calculated using ANCOVA. Correlations were performed using Pearson (r) or Spearman (rho), as indicated. To determine community structure in correlation networks, the package ggbiplot v0.55 was used. Adjusted p-values were calculated using the Benjamini-Hochberg method of the p.adjust function in the R library *stats* (v4.0.4). Spearman correlation networks were plotted using the imsbInfer v0.2.4 library in R (itself based off of the R library called iGraph). Lifespan calculations for mice and significance tests were made using the “survival” v3.2-7 package on R with the survfit and survdiff functions. Lifespan calculations for *C. elegans* are detailed in the *C. elegans* section. The output longevity data were retained for strains with ≥ 6 recorded lifespans within a cohort. To compare lifespan across diet, a minimum of 12 natural deaths were thus necessary. Outliers were removed with the R library “outliers” (v0.14) using the “rm.outlier” function.

Reference gene sets for the functional analyses in Figures 3–5 were taken from the “C2” curated gene set lists on the GSEA website using version 4.0.0. The exact reference names of all 25 gene sets are in **Table S6**. Note that this analysis only considers the 3772 genes with both mRNA and protein data. Canonical functional gene assignments were curated either from GSEA (for pathway membership) or for CORUM (for complex membership). To determine the significance of correlation networks, random networks were permuted using the same number of nodes as in the target network, but randomly selected from amongst the 3772 other proteins and mRNA. For networks with multiple categories of gene sets, the target and random networks were computed both on a per-set basis as well as the total set and the interaction between the sets. 10,000 random networks were permuted for each comparison, and networks were assigned p-values based on this permutation, according to how many random gene sets had at least as many edges as the target set at the given cutoff, or assigned p < 0.0001 if no random network had as many edges as the input network at the cutoff. All figures were generated either in R and refined in Adobe Illustrator, or were hand-drawn entirely in Adobe Illustrator (e.g. **Figure 1B**, **Figure 3A**).

## Supporting information

All Supplemental Tables

All code used to generate figures

## SUPPLEMENTARY INFORMATION & DATA AVAILABILITY

Supplementary figures and tables are included. All code required for generating the figures, along with some helper files, are also included. Raw mass spectrometry data for metabolomics are available on the MassIVE resource under accession ID MSV000081441 with reviewer access ftp://MSV000081441@massive.ucsd.edu (login is “Guest” with no password). Raw mass spectrometry data for proteomics are available on ProteomeXchange [75] under accession ID PXD009160 with reviewer login and password **<u>MuFdMZtp</u>**. The full transcriptomics data are available on GeneNetwork.org under Species: mouse, Group: BXD NIA Longevity Study, Type: Liver mRNA, and Dataset: UTHSC BXD Harvested Liver RNA-Seq. All processed data, for the omics layers and the phenotypes, are available in a “ready to use” **Table S2**, which is the version of the data that was used to generate the figures.

## ACKNOWLEDGEMENTS

Research was further supported by the EPFL, ETHZ, the NIH (R01AG043930 to RW), the European Research Council (Proteomics4D (AdvG grant 670821 and Proteomics v3.0; AdvG-233226 to RA), and SNSF (31003A-140780, 31003A-143914 and CSRII3-136201 to RA, PP00P3_163898 to CS and CYE). NP and PB were supported by ERC No 786461 (CausalStats - ERC-2017-AdvG). EGW was supported by an NIH F32 Ruth Kirchstein Fellowship (F32GM119190). Thanks to Lorne Rose at the University of Tennessee Center of Excellence Sequencing Facility for transcriptomics, and to Sebastien Lamy at the EPFL phenotyping unit (UDP) for clinical blood analysis, to Casey Chapman for BXD colony maintenance, and to Ludovic Gillet, Patrick Pedrioli, Özgen Eren, Chloe Lee, and Yansheng Liu for proteomics discussions.

## AUTHOR CONTRIBUTIONS

EGW and RW established the BXD aging project and secured funding. SR, EGW, CB, JI, RWW, and LL managed the colony and the tissue collections. JI and CB managed the daily needs of the BXD colony. JI, SR, and RWW performed the transcriptomics. RA and EGW performed the proteomics. JH calculated the QTLs. MH, NZ, and EGW performed the metabolomics. JC normalized the proteomics data. NP and PB developed and performed the causality/stability analyses. EGW performed all other statistical analyses on the omics data. CS and CYE performed the *C. elegans* experiments. The paper was written by EGW, with assistance from the other authors for their specific contributions.

---

**Figure S1.**
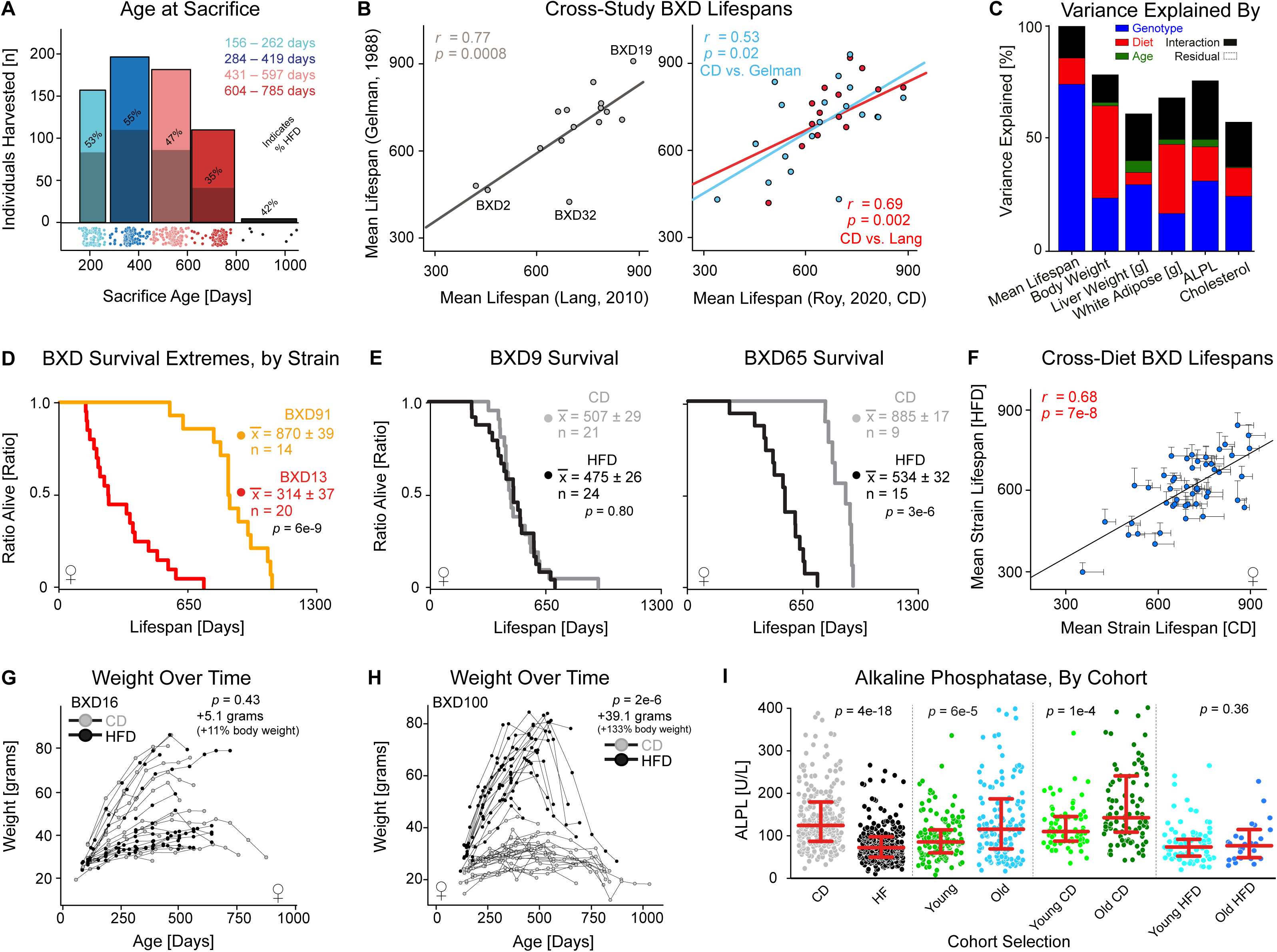
**Supplemental Overview of Phenotype Data** (**A**) Histogram and dot plot of the animals with tissues collected at each time and diet, with the fraction of CD and HFD noted. Due to decreased lifespan in HFD, there is a disbalance in the CD/HFD ratio for the oldest timepoint at 24 months. (**B**) Significant lifespan correlations are observed between BXD strains in all three lifespan studies. (**C**) Proportion of variance explained by genotype, diet, age, or the interactions between these three variables, for key phenotypes, as calculated by ANCOVA. (**D**) Lifespan calculations as a function of genotype for two strains with extreme differences in lifespan: BXD13 (very short lived) and BXD91 (very long lived). (**E**) Kaplan–Meier curves for two strains with extreme effects of diet on lifespan; BXD9 is unaffected, while BXD65 lives almost a year shorter on HFD. (**F**) Lifespan correlation by strain between CD and HFD fed cohorts in this study. (**G–H**) Body weight over time chart for two strains with extreme effects of diet on weight; BXD16 is obese in either case, while BXD100 more than doubles its weight on HFD. (**I**) Serum alkaline phosphatase (ALPL) levels, segregated by different independent variables.

**Figure S2.**
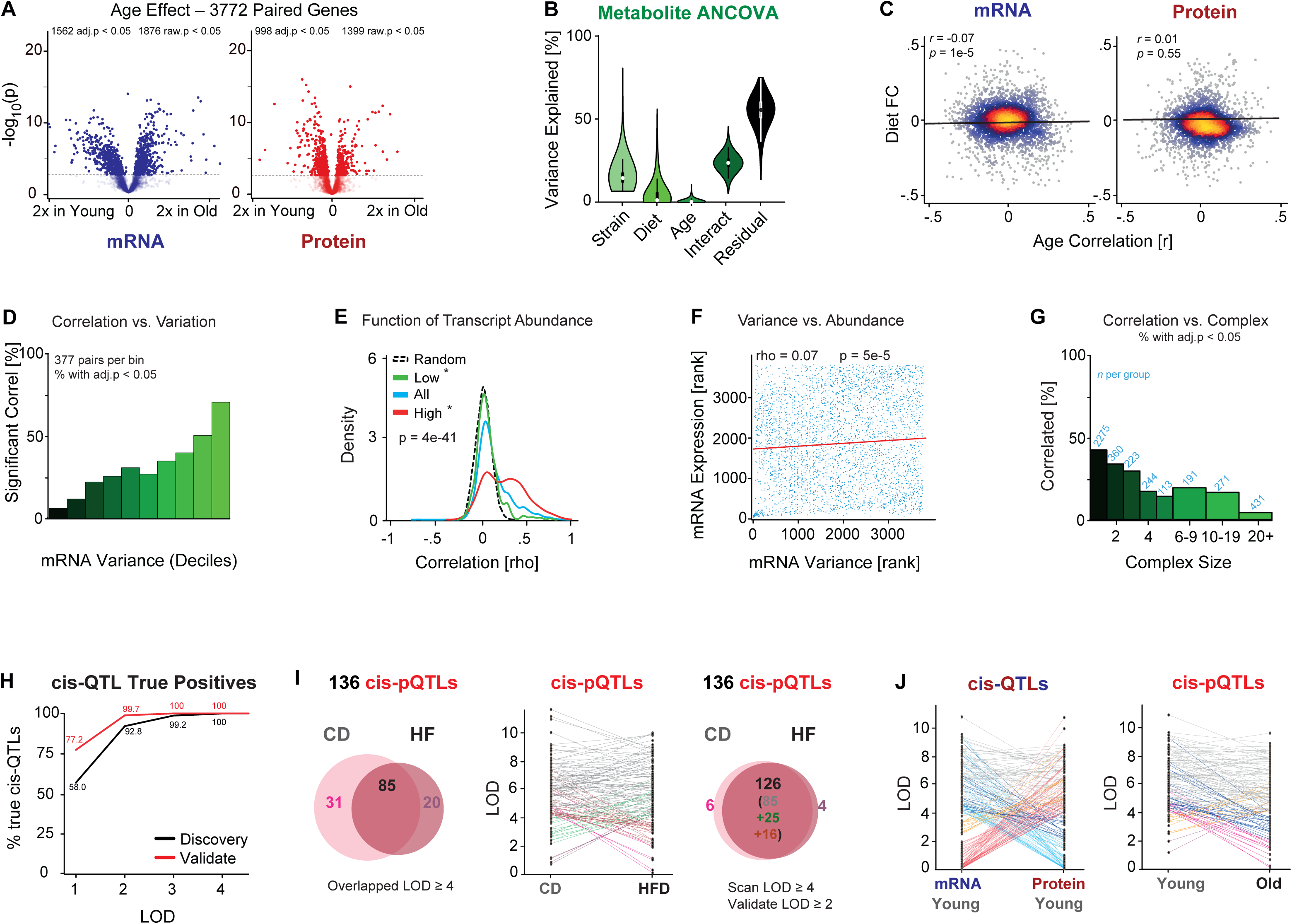
**Supplemental Patterns in Multifactorial, Multi-omic Analysis** (**A**) Volcano plot showing the effect of age on mRNA and protein levels between old and young individuals. (**B**) Variation explained for all metabolites a function of the independent variables and their interactions, and the residual. (**C**) Correlation density plot of the impact of diet and age on mRNA and protein levels. (**D**) Histogram of the percentage of significantly correlated mRNA–protein pairs as a function of the variance of mRNA expression across the population. (**E**) Density plot of mRNA–protein correlation as a function of transcript abundance. (**F**) Spearman correlation plot between expression variance and abundance for mRNA. Higher-expressed genes have more variance and are slightly more likely to correlate with their protein. (**G**) Histogram of the percentage of significantly correlated mRNA–protein pairs as a function of the size of the protein complex to which the gene belongs, using CORUM annotation; genes in larger complexes are much less likely to correlate between their mRNA and protein levels. (**H**) Empirical calculations of the false discovery rate of cis-eQTLs as a function of LOD score depending on if the gene is a novel target (“discovery”) or the validation of expectations from independent cis-QTL data. (**I**) Left: Venn diagram of the overlap of cis-pQTLs for CD or HFD cohorts using a strict cutoff of LOD ≥ 4. Middle: Slopegraph of LOD scores of all 165 genes with significant cis-pQTLs. Right: The same Venn diagram, but now using more flexible cutoffs. (**J**) cis-QTL consistency on a gene-level basis between mRNA and protein levels for just young individuals (left) and as a function across age for protein levels (right). The same patterns are observed here as for across-diet comparisons.

**Figure S3.**
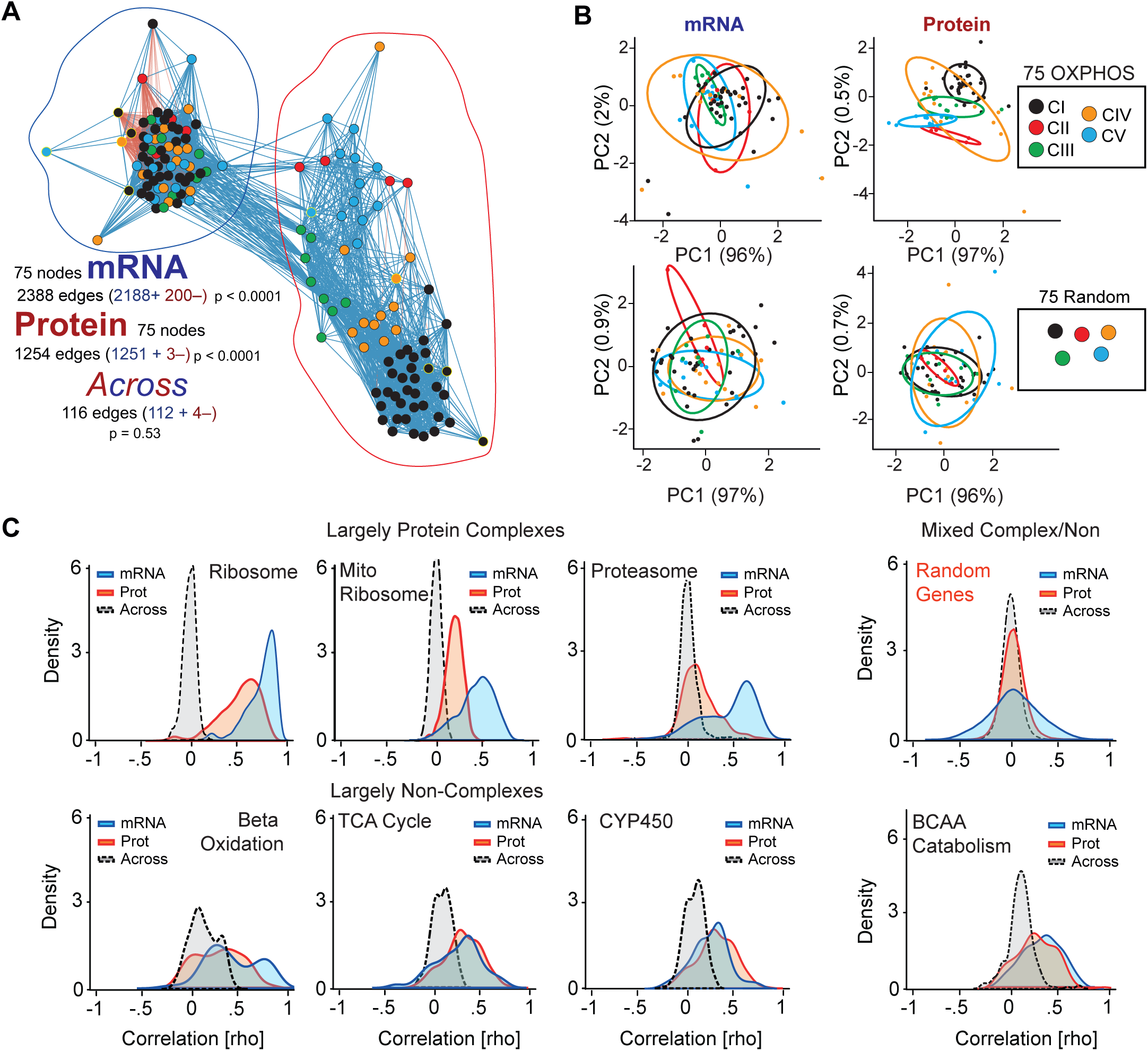
**Supplemental Functional Network Analysis and False Discovery** (**A**) Spearman correlation network for the 75 OXPHOS genes measured at both the mRNA and protein level, showing the low level of connectivity between the two layers (∼3% of edges are across mRNA to protein). (**B**) Top: PCA biplots for the 75 OXPHOS genes as a function of complex membership. mRNA has no differentiation between complexes, while the proteins are distinct per complex, except for CIV. Bottom: Negative control PCA plots for a random selection of 75 genes, showing no differentiation between random sets of the same size. (**C**) Correlation density plots of correlation networks for the ribosome, mitochondrial ribosome, beta oxidation, and TCA cycle as a function of mRNA level, protein level, or across the two. Even in cases where mRNA and protein both have strong correlation networks (e.g. ribosome, TCA cycle), there is little or no correlation between mRNA and protein. The correlations of random genes within mRNA/protein or across layer is shown at the top-right. While the average correlation of two mRNAs is similar to the average of two random proteins (+0.037 for mRNA, +0.027 for protein), the standard deviation is highly different: ±0.252 for mRNA, ±0.122 for protein. Consequently, network significance is always permuted.

**Figure S4.**
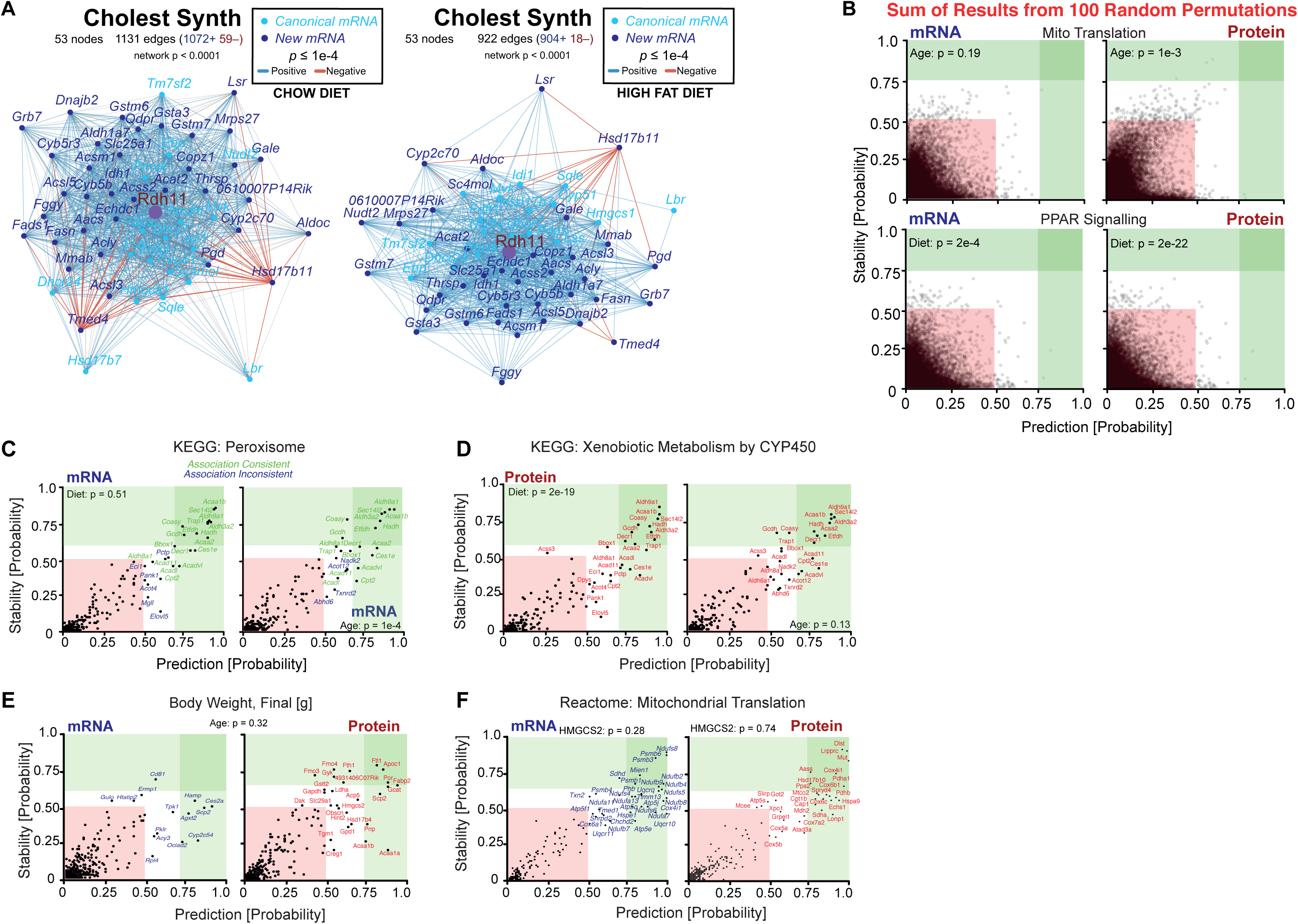
**Supplemental Stability Inference and False Discovery** (**A**) Correlation networks of cholesterol synthesis core membership plus the top significant candidates as a function of CD and HFD, for mRNA. (**B**) The overlap of 100 negative control permutation networks using the 68-member mitochondrial translation gene set (top) and the 42-member PPAR signalling pathway gene set. The theoretical upper-bound false discovery in this area is 1 discovery per test in the green area regardless of gene set. Here, for this randomized gene set, the sum of 200 permutation tests discovers only 4 nodes in the mRNA and 2 in the protein network, empirically indicating a false positive of approximately 0.03 per test averaged across all runs. Permuted false discovery in the white area is approximately 1 node per test. Note that the exact cutoff values change slightly depending on input. (**C**) Prediction–stability plot for the peroxisome network in mRNA, separated by either diet or age. (**D**) Prediction–stability plot for the CYP450 network discovery at the mRNA (left) and protein (right) level. Significant hits are labeled. (**E**) Prediction–stability plot for body weight as a function of age. Compared to body weight (Figure 5H), where the exact same raw data are used, age provides minimal causal information for how gene expression varies according to body weight. (**F**) Prediction–stability plot examining the mitochondrial ribosome expanded network as a function of *Hmgcs2* allele. For the 31 protein hits, the average distance from the X=Y line is 0.22 units according to *Hmgcs2* (compared to 0.33 by *Cox7a2l*).

**Table S1. Processed Phenotype, mRNA, Protein, and Metabolite Data** This table contains all raw longevity and body weight data, and a list of which harvested cohorts were selected and used for the omics experiments.

**Table S2. Processed Phenotype, mRNA, Protein, and Metabolite Data** This table contains a full list of normalized measurements for every sacrificed individual at the mRNA, protein, and metabolite level. Note that the same metabolomics extraction was done twice for every liver in complete experimental replicate about two months apart. Each individual liver was thus run 4 times: two separate extractions on the same liver tissue, and then each of those extractions had two technical injection replicates. The two separate extractions are *both* included in the output table, prefaced with either “Metab_Run1” or “Metab_Run2”. The two technical injection replicates for each were averaged to provide the data in the table (i.e. “Run1-TechRep1” was averaged with “Run1-TechRep2” and “Run2-TechRep1” with “Run2-TechRep2”). Furthermore, as this metabolomics profiling was relatively inexpensive, we analyzed around 620 livers—so a larger set than for mRNA and protein—thus many more cohorts have biological replicates run in addition to the two different “types” of technical replicate (i.e. injection technical replicate and extraction technical replicate).

**Table S3. GO Enrichment Categories of 3772 Paired mRNA & Protein Quantified & cis-QTLs** Sheet 1 contains a list of the enriched GO terms for all 3772 genes that were measured at the mRNA and protein level, showing that certain categories are well-covered (e.g. 37% of cytoplasm genes, p = 4e-180) while other categories are relatively poorly represented (e.g. only 4% of lipoprotein genes, p = 2e-6). Sheet 2 contains a list of all LOD scores for all 3772 genes at their cis-locus as a function of mRNA/protein, dietary status, and age.

**Table S4. Aging Biomarkers** This is a list of all genes that were highly correlated with age, their ontological sets, and for top hits, a breakdown of their associated *C. elegans* gene(s).

**Table S5. C. elegans Ctsd Longevity Data** This contains the summary data and all individual worm data for both experiments on *asp-4* knockdown (i.e. *Ctsd*) in *C. elegans*.

**Table S6. Candidate Gene Networks for Functional Analysis** This table contains a list of all 25 functional gene networks analyzed at the mRNA and protein level. The gene sets were pre-selected by the authors for the core role they play in energy metabolism, and with prior hypotheses in mind that gene sets may be affected by diet (based on our prior experience [18]) or age (based on literature searches; see table). Basic summary data about each gene set are available, e.g. number of members, GSEA results, network significance, and summary of stability analysis.

**Table S7. Stability Analysis: Gene Hits** This table contains all of the results from the stability analysis on all 25 candidate gene sets as a function of diet, age, or expression type (protein/mRNA). The gene sets that were highlighted in specific figure panels get a second sheet inside the table containing only the subset of genes that matched the stricter criteria (i.e. x- or y- axis values of ≥ 0.50) along with basic information about each gene (e.g. description, Entrez ID). Genes that had differences of ≥ 0.30 in the x and y axes were considered to yield causal information (e.g. such genes will correlate with the target pathway but only—or much more strongly—in one diet or age than the other).

