## Supplementary material for "Multi-Omic Profiling of the Liver Across Diets and Age in a Diverse Mouse Population": All code used to generate figures: b_Figure1_Plotting_Code_vF.r

```
################################################################################  
# First priority is loading the multi-omics dataset, which will be used throughout
## This is the INDIVIDUAL data
################################################################################# 

setwd("~/Dropbox/Aging\ BXD\ Study/1\ _\ Manuscript\ -\ Evan\ Multi-Omic/")

AllData_header=read.table("aTableS2_ALL_DATA.csv", nrows=8, row.names=1, header=T, sep=",")
AllData_header=AllData_header[,3:ncol(AllData_header)]
AllData=read.table("aTableS2_ALL_DATA.csv", row.names=1, header=T,  na.strings="", sep=",", skip=8, stringsAsFactors=FALSE)
AllData=AllData[1:nrow(AllData),3:ncol(AllData)]

AllData_CD=matrix(NA, nrow=nrow(AllData), ncol=length(AllData))
for(i in 1:length(AllData)) {
	if(as.character(AllData_header["Diet",i])=="CD") {   AllData_CD[,i]=as.numeric(as.character(AllData[,i]))	}
} 
rownames(AllData_CD)=rownames(AllData)

AllData_HF=matrix(NA, nrow=nrow(AllData), ncol=length(AllData))
for(i in 1:length(AllData)) {
	if(as.character(AllData_header["Diet",i])=="HF") {   AllData_HF[,i]=as.numeric(as.character(AllData[,i]))	}
}
rownames(AllData_HF)=rownames(AllData)

AllData_young=matrix(NA, nrow=nrow(AllData), ncol=length(AllData))
for(i in 1:length(AllData)) {
	if(as.character(AllData_header["Age",i])<=420) {	AllData_young[,i]=as.numeric(as.character(AllData[,i]))	}
}
rownames(AllData_young)=rownames(AllData)

AllData_old=matrix(NA, nrow=nrow(AllData), ncol=length(AllData))
for(i in 1:length(AllData)) {
	if(as.character(AllData_header["Age",i])>=420) {	AllData_old[,i]=as.numeric(as.character(AllData[,i]))	}
}
rownames(AllData_old)=rownames(AllData)

AllData_YoungCD=matrix(NA, nrow=nrow(AllData), ncol=length(AllData)) # 627 - sum(is.na(AllData_YoungCD["Prot_7873_Sept1",]), na.rm=T)
for(i in 1:length(AllData)) {
	if(as.character(AllData_header["Age",i])<=420 & as.character(AllData_header["Diet",i])=="CD") {	AllData_YoungCD[,i]=as.numeric(as.character(AllData[,i]))	}
}
rownames(AllData_YoungCD)=rownames(AllData)

AllData_YoungHF=matrix(NA, nrow=nrow(AllData), ncol=length(AllData))
for(i in 1:length(AllData)) {
	if(as.character(AllData_header["Age",i])<=420 & as.character(AllData_header["Diet",i])=="HF") {	AllData_YoungHF[,i]=as.numeric(as.character(AllData[,i]))	}
}
rownames(AllData_YoungHF)=rownames(AllData)

AllData_OldCD=matrix(NA, nrow=nrow(AllData), ncol=length(AllData))
for(i in 1:length(AllData)) {
	if(as.character(AllData_header["Age",i])>=420 & as.character(AllData_header["Diet",i])=="CD") {	AllData_OldCD[,i]=as.numeric(as.character(AllData[,i]))	}
}
rownames(AllData_OldCD)=rownames(AllData)

AllData_OldHF=matrix(NA, nrow=nrow(AllData), ncol=length(AllData))
for(i in 1:length(AllData)) {
	if(as.character(AllData_header["Age",i])>=420 & as.character(AllData_header["Diet",i])=="HF") {	AllData_OldHF[,i]=as.numeric(as.character(AllData[,i]))	}
}
rownames(AllData_OldHF)=rownames(AllData)

### We can also segregate on other factors, shown here as an example
AllData_COX7A2L_B6Allele=matrix(NA, nrow=nrow(AllData), ncol=length(AllData))
for(i in 1:length(AllData)) {
	if(as.character(AllData["COX7Allele",i])==0) {   AllData_COX7A2L_B6Allele[,i]=as.numeric(as.character(AllData[,i]))	}
} 
rownames(AllData_COX7A2L_B6Allele)=rownames(AllData)

AllData_COX7A2L_D2Allele=matrix(NA, nrow=nrow(AllData), ncol=length(AllData))
for(i in 1:length(AllData)) {
	if(as.character(AllData["COX7Allele",i])==1) {   AllData_COX7A2L_D2Allele[,i]=as.numeric(as.character(AllData[,i]))	}
}
rownames(AllData_COX7A2L_D2Allele)=rownames(AllData)

###### Now we can also separate data based on other variables:

## Yields proteomics data for 107 animals with lifespans ≥ 720 days
# and 108 animals with lifespans ≤ 663 days
# count with "627 - sum(is.na(AllData_LongLived["Prot_7873_Sept1",]), na.rm=T)
AllData_LongLived=matrix(NA, nrow=nrow(AllData), ncol=length(AllData)) # 627 - sum(is.na(AllData_LongLived["Prot_7873_Sept1",]), na.rm=T)
for(i in 1:length(AllData)) {
	if(is.na(as.character(AllData["Pheno_StrainMedianLifespan",i]))) {	AllData_LongLived[,i]=NA }
	else if(as.character(AllData["Pheno_StrainMedianLifespan",i])>=720) {	AllData_LongLived[,i]=as.numeric(as.character(AllData[,i]))	}
}
rownames(AllData_LongLived)=rownames(AllData)

## Yields proteomics data for 103 animals that have expected lifespans below 648 days, count with 627 - sum(is.na(AllData_ShortLived["Prot_7873_Sept1",]), na.rm=T)
AllData_ShortLived=matrix(NA, nrow=nrow(AllData), ncol=length(AllData))
for(i in 1:length(AllData)) {
	if(is.na(as.character(AllData["Pheno_StrainMedianLifespan",i]))) {	AllData_ShortLived[,i]=NA }
	else if(as.character(AllData["Pheno_StrainMedianLifespan",i])<=663) {	AllData_ShortLived[,i]=as.numeric(as.character(AllData[,i]))	}
}
rownames(AllData_ShortLived)=rownames(AllData)

################################################################################  
# FIGURE ONE
################################################################################# 

#$$$$$$ #$$$$$$ #$$$$$$ #$$$$$$ #$$$$$$ #$$$$$$ #$$$$$$ #$$$$$$
#$$$$$$ #$$$$$$ #$$$$$$ #$$$$$$ #$$$$$$ #$$$$$$ #$$$$$$ #$$$$$$
# PANEL 1A & PANEL 1B

# Entirely Hand-Drawn

#$$$$$$ #$$$$$$ #$$$$$$ #$$$$$$ #$$$$$$ #$$$$$$ #$$$$$$ #$$$$$$
#$$$$$$ #$$$$$$ #$$$$$$ #$$$$$$ #$$$$$$ #$$$$$$ #$$$$$$ #$$$$$$
# PANEL 1C 

library(survival) # For Kaplan-Meier graphs
library(openxlsx) # This is WAY faster than the xlsx library.
survive_raw=read.xlsx("~/Dropbox/Aging\ BXD\ Study/1\ _\ Manuscript\ -\ Evan\ Multi-Omic/aTableS1_AgingDB.xlsx", na.strings="") # Loads sheet 1 automatically
survive=subset(survive_raw, survive_raw$CauseOfDeath=="Natural" | survive_raw$CauseOfDeath=="Euthanized" & survive_raw$Sex=="F")

### THIS CODE SETS ANIMALS ALIVE OR DEAD FOR THE GRAPHING AND LIFESPAN CALCULATIONS
lifestatus = survive$AgeIfStillAlive # tells if animal is dead (1) or alive (0)
i=1
for(i in 1:length(lifestatus)) {
	if(lifestatus[i]==0)  {
		lifestatus[i]=1 		# sets animal as dead if its "AgeIfStillAlive" is zero (UTHSC's dead marker)
	}
	else {lifestatus[i]=0}		# sets animal as alive if it has an AgeIfStillAlive (UTHSC's "still alive" marker)
}

### Now this calculates the lifespans and plots them, here for all CD and HFD females, separated by diet but not by strain.
time=c(subset(survive$AgeIfStillAlive2, survive$DietCode == "CD" & survive$Sex=="F"), subset(survive$AgeIfStillAlive2, survive$DietCode == "HF" & survive$Sex=="F"))
time2=c(subset(survive$AgeAtDeath, survive$DietCode == "CD" & survive$Sex=="F"), subset(survive$AgeAtDeath, survive$DietCode == "HF" & survive$Sex=="F"))
event=c(subset(lifestatus, survive$DietCode == "CD" & survive$Sex=="F"), subset(lifestatus, survive$DietCode == "HF" & survive$Sex=="F"))
strains=c(subset(survive$DietCode, survive$DietCode == "CD" & survive$Sex=="F"), subset(survive$DietCode, survive$DietCode == "HF" & survive$Sex=="F"))
xy=survfit(Surv(time, time2, event)~strains)
plot(xy, lwd=3, xlab="Days", ylab="Ratio Alive", main="BXD Aging Colony", bty="L", col=c("grey","black"), xlim=c(0,1300), xaxp=c(0, 1300,2), ylim=c(0,1), yaxp=c(0,1,2), las=1)
#legend("topright", lty=1, lwd=3, bty="n", col=c("black","grey"), c("HFD", "CD"))
print(survfit(Surv(time, time2, event)~strains), print.rmean=T)


#$$$$$$ #$$$$$$ #$$$$$$ #$$$$$$ #$$$$$$ #$$$$$$ #$$$$$$ #$$$$$$
#$$$$$$ #$$$$$$ #$$$$$$ #$$$$$$ #$$$$$$ #$$$$$$ #$$$$$$ #$$$$$$
# PANEL 1D
library(survival) # For Kaplan-Meier graphs
library(openxlsx) # This is WAY faster than the xlsx library.
survive_raw=read.xlsx("~/Dropbox/Aging\ BXD\ Study/1\ _\ Manuscript\ -\ Evan\ Multi-Omic/aTableS1_AgingDB.xlsx", na.strings="") # Loads sheet 1 automatically
survive=subset(survive_raw, survive_raw$CauseOfDeath=="Natural" | survive_raw$CauseOfDeath=="Euthanized" & survive_raw$Sex=="F")

### THIS CODE SETS ANIMALS ALIVE OR DEAD FOR THE GRAPHING AND LIFESPAN CALCULATIONS
lifestatus = survive$AgeIfStillAlive # tells if animal is dead (1) or alive (0)
i=1
for(i in 1:length(lifestatus)) {
	if(lifestatus[i]==0)  {
		lifestatus[i]=1 		# sets animal as dead if its "AgeIfStillAlive" is zero
	}
	else {lifestatus[i]=0}		# sets animal as alive if it has an AgeIfStillAlive of nonzero
}

time=survive$AgeIfStillAlive2
time2=survive$AgeAtDeath
event=lifestatus
strains=survive$StrainNameCurrent
xy=survfit(Surv(time, time2, event)~strains)

dev.new(width=8, height=7)
plot(xy, lwd=3, xlab="Days", ylab="Ratio Alive", main="BXD Aging Colony", bty="L", col=colors()[1:50], xlim=c(0,1300), xaxp=c(0, 1300,2), ylim=c(0,1), yaxp=c(0,1,2), las=1)
#legend("topright", lty=1, lwd=3, bty="n", col=c("red","blue","green","purple"), c("BXD13", "BXD91", "BXD9", "BXD73")) # Drawn in Illustrator
print(survfit(Surv(time, time2, event)~strains), print.rmean=T)
survdiff(Surv(time2, event)~strains)

# Strains with n.events < 6 were removed in Illustrator based on the output table from survdiff, and colors were assigned for the four target strains in Illustrator.

#$$$$$$ #$$$$$$ #$$$$$$ #$$$$$$ #$$$$$$ #$$$$$$ #$$$$$$ #$$$$$$
#$$$$$$ #$$$$$$ #$$$$$$ #$$$$$$ #$$$$$$ #$$$$$$ #$$$$$$ #$$$$$$
# PANEL 1E

library(gplots)

# This loads the same file but now loads sheet 2
allweight=read.xlsx("~/Dropbox/Aging\ BXD\ Study/1\ _\ Manuscript\ -\ Evan\ Multi-Omic/aTableS1_AgingDB.xlsx", sheet=2, na.strings="")

AgeOverTime=cbind(allweight$AgeAtWeight01, allweight$AgeAtWeight02, allweight$AgeAtWeight03, allweight$AgeAtWeight04, allweight$AgeAtWeight05, allweight$AgeAtWeight06, allweight$AgeAtWeight07, allweight$AgeAtWeight08, allweight$AgeAtWeight09, allweight$AgeAtWeight10, allweight$AgeAtWeight11, allweight$AgeAtWeight12, allweight$AgeAtWeight13, allweight$AgeAtWeight14, allweight$AgeAtWeight15)
rownames(AgeOverTime)=allweight$EarTagNumberCurrent
WeightOverTime=cbind(allweight$Weight01, allweight$Weight02, allweight$Weight03, allweight$Weight04, allweight$Weight05, allweight$Weight06, allweight$Weight07, allweight$Weight08, allweight$Weight09, allweight$Weight10, allweight$Weight11, allweight$Weight12, allweight$Weight13, allweight$Weight14, allweight$Weight15)
rownames(WeightOverTime)=allweight$EarTagNumberCurrent
TimeOnDietOverTime=cbind(allweight$DaysOnDietAtWeight01, allweight$DaysOnDietAtWeight02, allweight$DaysOnDietAtWeight03, allweight$DaysOnDietAtWeight04, allweight$DaysOnDietAtWeight05, allweight$DaysOnDietAtWeight06, allweight$DaysOnDietAtWeight07, allweight$DaysOnDietAtWeight08, allweight$DaysOnDietAtWeight09, allweight$DaysOnDietAtWeight10, allweight$DaysOnDietAtWeight11, allweight$DaysOnDietAtWeight12, allweight$DaysOnDietAtWeight13, allweight$DaysOnDietAtWeight14, allweight$DaysOnDietAtWeight15)
rownames(TimeOnDietOverTime)=allweight$EarTagNumberCurrent

CohortTags = subset(allweight$EarTagNumberCurrent,allweight$AgeAtDeath>=800)
CohortTagsCD = subset(allweight$EarTagNumberCurrent, allweight$AgeAtDeath>=800 & allweight$DietCode=="CD" & allweight$Sex=="F")
CohortTagsHF = subset(allweight$EarTagNumberCurrent, allweight$AgeAtDeath>=800 & allweight$DietCode=="HF" & allweight$Sex=="F")

# Initialize the graph and then plot (LEFT) for CD
plotCI(c(NA), c(NA), ylim=c(10, 105), main="All_CD", xlim=c(0, 1250), bty="L", xlab="Age [days]", ylab="Weight [grams]", las=1, yaxp=c(0, 100, 4), xaxp=c(0, 1250, 4))
for(i in 1:length(CohortTagsHF)) {
plotCI(subset(AgeOverTime[,1:15], allweight$EarTagNumberCurrent==CohortTagsCD[i]), subset(WeightOverTime[,1:15], allweight$EarTagNumberCurrent==CohortTagsCD[i]), pch=21, pt.bg="grey", col="grey", type="o", cex.main=2, add=T)
}

# Initialize the graph and then plot (RIGHT) for HFD
plotCI(c(NA), c(NA), ylim=c(10, 105), main="All_HFD", xlim=c(0, 1250), bty="L", xlab="Age [days]", ylab="Weight [grams]", las=1, yaxp=c(0, 100, 4), xaxp=c(0, 1250, 4))
for(i in 1:length(CohortTagsHF)) {
plotCI(subset(AgeOverTime[,1:15], allweight$EarTagNumberCurrent==CohortTagsHF[i]), subset(WeightOverTime[,1:15], allweight$EarTagNumberCurrent==CohortTagsHF[i]), pch=21, pt.bg="black", col="grey", type="o", cex.main=2, add=T)
}

# Note that the final plots are cleaned up in Illustrator (e.g. black outline added, legend added)

#$$$$$$ #$$$$$$ #$$$$$$ #$$$$$$ #$$$$$$ #$$$$$$ #$$$$$$ #$$$$$$
#$$$$$$ #$$$$$$ #$$$$$$ #$$$$$$ #$$$$$$ #$$$$$$ #$$$$$$ #$$$$$$
# PANEL 1F

# Pulled out from the above data, taking for the point closest to 18 months, but within the maximum boundaries of between 490±599 days of age (545 days ± 10%)

library(vioplot)
CDweights=c(13.7,15.9,17.3,17.7,17.8,18,18.2,18.6,18.7,19,19.1,19.3,19.5,19.5,19.7,19.7,19.8,19.9,19.9,19.9,19.9,19.9,20,20,20.1,20.1,20.2,20.2,20.3,20.4,20.4,20.5,20.7,20.7,20.7,20.7,20.7,20.8,20.8,20.9,21,21,21,21.1,21.1,21.2,21.2,21.2,21.2,21.3,21.3,21.4,21.4,21.4,21.5,21.6,21.6,21.6,21.6,21.6,21.7,21.7,21.9,22,22.1,22.1,22.1,22.1,22.2,22.2,22.3,22.3,22.4,22.4,22.4,22.5,22.5,22.5,22.6,22.6,22.6,22.6,22.6,22.6,22.7,22.7,22.7,22.8,22.8,22.9,22.9,22.9,23,23,23,23,23.1,23.1,23.1,23.1,23.2,23.2,23.2,23.3,23.3,23.4,23.4,23.4,23.4,23.5,23.5,23.5,23.5,23.6,23.6,23.6,23.6,23.6,23.7,23.7,23.8,23.8,23.8,23.9,23.9,23.9,23.9,23.9,24,24,24,24,24,24.1,24.1,24.1,24.1,24.1,24.2,24.2,24.2,24.2,24.2,24.3,24.3,24.3,24.3,24.3,24.4,24.4,24.4,24.4,24.4,24.4,24.5,24.5,24.5,24.5,24.6,24.7,24.7,24.7,24.7,24.7,24.7,24.7,24.7,24.8,24.8,24.8,24.8,24.8,24.8,24.9,24.9,24.9,24.9,24.9,25,25,25,25,25.1,25.1,25.2,25.2,25.2,25.2,25.2,25.2,25.2,25.3,25.3,25.3,25.3,25.3,25.3,25.4,25.4,25.4,25.5,25.5,25.5,25.5,25.5,25.6,25.6,25.6,25.6,25.6,25.6,25.7,25.8,25.8,25.8,25.8,25.8,25.8,25.9,25.9,25.9,25.9,25.9,25.9,25.9,25.9,26,26,26,26,26.1,26.1,26.1,26.1,26.1,26.1,26.1,26.2,26.2,26.3,26.3,26.3,26.3,26.3,26.3,26.4,26.4,26.4,26.5,26.5,26.5,26.5,26.5,26.5,26.6,26.6,26.6,26.6,26.6,26.6,26.6,26.6,26.7,26.7,26.7,26.7,26.7,26.7,26.7,26.7,26.8,26.8,26.8,26.8,26.9,26.9,26.9,26.9,26.9,27,27,27,27,27,27.1,27.1,27.2,27.2,27.2,27.2,27.3,27.3,27.3,27.4,27.4,27.5,27.5,27.5,27.6,27.6,27.6,27.6,27.7,27.8,27.8,27.8,27.8,27.8,27.8,27.9,27.9,27.9,27.9,27.9,28,28,28,28,28,28.1,28.2,28.2,28.2,28.2,28.2,28.3,28.3,28.3,28.3,28.3,28.4,28.4,28.4,28.4,28.4,28.4,28.5,28.5,28.5,28.5,28.5,28.5,28.6,28.6,28.6,28.6,28.6,28.6,28.7,28.7,28.7,28.7,28.8,28.8,28.9,28.9,29,29,29,29.1,29.2,29.2,29.2,29.2,29.2,29.3,29.3,29.3,29.4,29.4,29.4,29.4,29.4,29.5,29.5,29.6,29.6,29.6,29.6,29.6,29.6,29.6,29.6,29.7,29.7,29.7,29.7,29.9,29.9,29.9,29.9,30,30,30,30,30.1,30.1,30.1,30.1,30.1,30.2,30.2,30.3,30.3,30.3,30.4,30.4,30.4,30.5,30.5,30.5,30.5,30.6,30.6,30.6,30.6,30.8,30.8,30.8,30.8,30.8,30.8,30.9,31,31,31,31,31.1,31.2,31.2,31.3,31.3,31.3,31.3,31.4,31.5,31.5,31.5,31.6,31.6,31.6,31.7,31.7,31.8,31.9,31.9,31.9,32,32,32.1,32.1,32.1,32.1,32.1,32.2,32.2,32.2,32.2,32.4,32.4,32.4,32.5,32.6,32.7,32.7,32.8,32.8,32.9,33,33.1,33.1,33.2,33.2,33.4,33.6,33.6,33.6,33.7,33.7,33.7,33.7,33.7,33.7,33.8,33.8,33.8,33.8,33.9,33.9,33.9,34,34,34.1,34.1,34.1,34.2,34.2,34.2,34.5,34.6,34.6,34.6,34.6,34.6,34.8,34.8,35.4,35.4,35.5,35.6,35.7,35.7,35.7,35.9,36,36,36.1,36.2,36.3,36.3,36.3,36.3,36.3,36.4,36.5,36.7,36.7,36.7,36.7,36.8,36.9,37,37.1,37.1,37.2,37.3,37.5,37.5,37.8,37.8,37.8,37.9,38,38.1,38.1,38.1,38.2,38.3,38.4,38.4,38.5,38.5,38.6,39,39.4,39.4,39.5,39.6,39.6,39.7,40.1,40.2,40.6,40.7,40.9,41.2,41.4,41.4,42,42.1,42.1,42.2,42.7,43.1,43.4,43.7,43.9,43.9,44.1,44.1,44.2,44.9,45.2,45.7,46.6,47,47,47.1,47.2,47.7,48.6,49,49.5,49.6,49.9,51.2,52.4,53.4,53.9,54.4,54.5,58,58.8,59.3,61.3,67.7,73.8)
HFDweights=c(13.1,18,19.2,20.4,21.3,21.6,21.8,21.8,21.8,22.7,22.7,23,23,23,23,23.1,23.1,23.1,23.3,23.8,23.9,25.2,25.5,25.6,26,26.6,27.3,27.4,27.6,28.1,28.4,28.8,28.9,29.1,29.1,29.2,29.2,29.8,29.9,30,30,30.2,30.2,30.3,30.3,30.4,30.4,30.6,30.6,30.9,31,31.4,31.5,31.7,31.7,31.8,31.9,32,32.1,32.2,32.2,32.3,32.7,32.7,32.7,32.8,32.9,32.9,33,33.2,33.3,33.6,34.1,34.2,34.4,34.4,34.6,34.7,35.2,35.2,35.3,35.3,35.4,35.4,35.5,35.5,35.5,35.6,35.7,35.7,35.7,35.8,36,36,36.2,36.4,36.5,36.6,36.9,37,37.1,37.2,37.4,37.5,37.5,37.6,37.8,38,38,38,38.2,38.3,38.4,38.4,38.4,38.4,38.4,38.6,38.7,38.7,38.9,38.9,39,39,39.1,39.2,39.2,39.2,39.3,39.4,39.5,39.6,40,40,40,40,40.1,40.2,40.3,40.4,40.4,40.8,40.9,40.9,41.1,41.1,41.1,41.2,41.2,41.3,41.4,41.4,41.6,41.6,41.8,41.8,41.9,41.9,42,42.1,42.2,42.3,42.3,42.6,42.6,42.7,42.7,42.8,42.9,42.9,43,43.1,43.1,43.2,43.3,43.3,43.3,43.4,43.7,43.7,43.8,43.9,44,44,44.1,44.2,44.2,44.5,44.7,44.7,44.8,44.8,44.9,44.9,44.9,45.2,45.3,45.3,45.3,45.4,45.4,45.4,45.6,45.6,45.6,45.7,46,46.1,46.1,46.1,46.2,46.2,46.3,46.6,46.6,46.6,46.8,46.9,47,47.1,47.1,47.2,47.2,47.3,47.3,47.3,47.3,47.3,47.4,47.5,47.8,47.8,47.8,47.9,48,48.1,48.1,48.2,48.2,48.3,48.4,48.4,48.5,48.5,48.7,48.8,48.8,48.8,48.8,48.9,49,49.1,49.1,49.2,49.2,49.3,49.3,49.4,49.6,49.6,49.7,49.8,49.9,50,50.1,50.2,50.2,50.3,50.6,50.6,50.6,50.7,50.8,50.9,51,51.2,51.5,51.6,51.7,51.9,52,52,52,52.2,52.2,52.2,52.4,52.4,52.5,52.5,52.6,52.6,52.7,52.7,52.9,53,53.1,53.2,53.3,53.5,53.6,53.7,53.9,54.1,54.2,54.2,54.3,54.4,54.4,54.5,54.6,54.7,54.7,54.7,54.7,54.7,54.7,54.8,54.9,55,55.1,55.1,55.1,55.3,55.6,55.7,55.7,55.7,55.8,55.8,55.8,56,56.3,56.4,56.6,56.6,56.7,56.8,57,57.3,57.5,57.7,57.8,57.8,57.9,57.9,58,58,58,58.2,58.2,58.3,58.3,58.4,58.6,58.6,58.6,58.6,58.6,58.7,58.7,58.9,58.9,59,59,59.2,59.2,59.2,59.3,59.5,59.5,59.6,59.6,59.7,59.9,60.1,60.2,60.2,60.2,60.4,60.4,60.4,60.5,60.5,60.5,60.5,60.6,60.7,60.8,60.8,61.1,61.2,61.3,61.5,61.6,61.7,61.7,61.8,61.8,61.9,62.1,62.2,62.2,62.5,62.5,62.7,62.7,62.8,62.8,62.9,63,63.2,63.3,63.3,63.5,63.5,63.5,63.6,63.6,64.2,64.4,64.5,64.5,64.7,64.7,64.8,64.8,64.9,64.9,65,65.1,65.2,65.3,65.4,65.5,65.6,65.6,65.7,65.7,65.8,65.8,66.2,66.6,66.7,66.8,67.4,67.5,67.5,67.6,67.7,68,68.2,68.2,68.3,68.5,68.6,68.8,69,69,69.4,69.6,69.6,69.6,69.7,69.8,69.8,70.1,70.3,70.8,70.9,71.2,71.2,71.4,71.6,71.6,71.6,71.9,72.1,72.2,72.3,72.4,72.8,72.9,72.9,73.1,73.1,73.1,73.2,73.4,73.6,73.7,73.7,73.8,74,74.9,74.9,75,75,75.4,75.5,75.6,75.7,75.7,76.8,76.9,77.2,77.2,77.5,78.4,79.9,80.1,80.2,80.4,80.4,80.6,80.6,80.7,81,81.1,81.1,81.2,81.3,81.3,81.8,81.8,82.1,82.1,82.3,82.3,82.6,82.7,82.8,82.9,83.4,83.5,83.8,84,84.7,84.9,87.4,88,89.3,90.4,94.4,95.7,99.2)

vioplot(CDweights,HFDweights)

t.test(CDweights,HFDweights)$p.value


#$$$$$$ #$$$$$$ #$$$$$$ #$$$$$$ #$$$$$$ #$$$$$$ #$$$$$$ #$$$$$$
#$$$$$$ #$$$$$$ #$$$$$$ #$$$$$$ #$$$$$$ #$$$$$$ #$$$$$$ #$$$$$$
# PANEL 1G
# Young is defined as < 420 days, and old as the contrary (> 420 days)
# Data are pulled out of the "Table S2_All_Data" for ALPL metabolite, simply by sorting the table in Excel for each group
# Colors and formatting applied in Illustrator, and the two plots were combined in Illustrator (dubbed "LEFT SIDE" and "RIGHT SIDE" below in the code)

## LEFT SIDE
library(outliers)
b6_f_cd_young_alpl_metabolite=c(30,114,101,87,137,82,104,82,92,94,202,NA,75,82,82,78,NA,113,96,86,81,65,NA,207,110,93,139,106,191,136,98,172,158,102,57,144,92,133,79,107,110,110,NA,NA,48,80,80,175,61,63,67,56,72,63,122,206,50,123,116,166,145,96,59,67,NA,NA,128,131,119,103,103,196,54,NA,127,NA,51,NA,96,39,79,126,NA,191,113,184,88,155,161,238,129,184,198,132,392)
d2_f_cd_young_alpl_metabolite=c(160,229,336,139,109,140,176,87,146,162,114,NA,77,99,193,96,133,152,134,222,201,65,79,503,197,165,NA,NA,102,104,129,75,70,76,89,250,261,107,312,201,192,135,150,181,139,NA,NA,NA)

groupA=rm.outlier(b6_f_cd_young_alpl_metabolite)
groupB=rm.outlier(d2_f_cd_young_alpl_metabolite)
stripchart(groupA, at=0.7, pch=21, cex=1.2, bg="grey", method="jitter", xlim=c(0.6, 1.3), las=1, ylim=c(0,500), vertical=T)
stripchart(groupB, at=1.0, pch=21, cex=1.2, bg="black", method="jitter", add=T, vertical=T)
	segments(0.65, quantile(groupA, na.rm=T)[2], 0.75, quantile(groupA, na.rm=T)[2], col="red", lwd=3)	# first quantile bar for first cohort
	segments(0.6, median(groupA, na.rm=T), 0.8, median(groupA, na.rm=T), col="red", lwd=3)	# median bar for first cohort
	segments(0.65, quantile(groupA, na.rm=T)[4], 0.75, quantile(groupA, na.rm=T)[4], col="red", lwd=3)	# first quantile bar for first cohort
	segments(0.7,quantile(groupA, na.rm=T)[2],0.7,quantile(groupA, na.rm=T)[4], col="red", lwd=3)
	segments(.95, quantile(groupB, na.rm=T)[2], 1.05, quantile(groupB, na.rm=T)[2], col="red", lwd=3)	# first quantile bar for second cohort	
	segments(.9, median(groupB, na.rm=T), 1.1, median(groupB, na.rm=T), col="red", lwd=3)	# median bar for second cohort, etc
	segments(.95, quantile(groupB, na.rm=T)[4], 1.05, quantile(groupB, na.rm=T)[4], col="red", lwd=3)	# first quantile bar for second cohort
	segments(1.0,quantile(groupB, na.rm=T)[2],1.0,quantile(groupB, na.rm=T)[4], col="red", lwd=3)
# legend	
pr = signif(t.test(groupA,groupB, paired=FALSE)$p.value,digits=2)			# This outputs the p-value of the correlation
pletter=substitute(paste("   ", italic("p"), " = ", pr), list(pr=signif(pr[1], digits=2)))	# Makes the r-value with italicized r

legend(legendposition, bg='white', bty="n", cex=1, text.col="red", legend=c(as.expression(pletter)))

## RIGHT SIDE
b6_f_old_bothdiets_alpl_metabolite=c(NA,NA,104,126,122,NA,NA,NA,NA,NA,NA,NA,101,137,NA,266,377,161,148,399,98,113,NA,78,NA,99,181,220,119,80,45,NA,435,546,134,84,64,68,133,132,80,544,NA,NA,NA,NA,92,289,38,262,640,46,17,40,386,37,34,201,189,99,70,332,255,459,533,114,88,309,236,189,107,115,73,NA,67,33,165,37,72,42,51,109,52,85,93,38,94,93,72,66,NA,NA,NA,NA,NA,111,52,178,148,36,48,NA,NA,83,143,NA,NA,NA,NA,NA,NA,NA,86,118,118,65,NA,NA,112,239,83,160,68,80,118,98,498,105,56,62,27,150,83,118,67,370,396,67,51,292,73)
d2_f_old_bothdiets_alpl_metabolite=c(NA,NA,NA,NA,NA,NA,235,175,126,113,732,507,NA,NA,NA,NA,72,160,338,145,102,50,69,81,94,45,181,124,141,NA,NA,123,252,61,122,221,396,NA,118,48,162,91,215,84,250,114,335,227,138,145,128,214,239,100,251,NA,73,104,116,117,124,110,229,84,39,101,37,273,386,284,NA,NA,47,NA,NA,NA,NA,NA,NA,NA,NA,NA,NA,NA,NA,134,174,167,270,NA,NA,121,98,99,103,150)

groupA=rm.outlier(b6_f_old_bothdiets_alpl_metabolite)
groupB=rm.outlier(d2_f_old_bothdiets_alpl_metabolite)
stripchart(groupA, at=0.7, pch=21, cex=1.2, bg="grey", method="jitter", xlim=c(0.6, 1.3), las=1, ylim=c(0,500), vertical=T)
stripchart(groupB, at=1.0, pch=21, cex=1.2, bg="black", method="jitter", add=T, vertical=T)
	segments(0.65, quantile(groupA, na.rm=T)[2], 0.75, quantile(groupA, na.rm=T)[2], col="red", lwd=3)	# first quantile bar for first cohort
	segments(0.6, median(groupA, na.rm=T), 0.8, median(groupA, na.rm=T), col="red", lwd=3)	# median bar for first cohort
	segments(0.65, quantile(groupA, na.rm=T)[4], 0.75, quantile(groupA, na.rm=T)[4], col="red", lwd=3)	# first quantile bar for first cohort
	segments(0.7,quantile(groupA, na.rm=T)[2],0.7,quantile(groupA, na.rm=T)[4], col="red", lwd=3)
	segments(.95, quantile(groupB, na.rm=T)[2], 1.05, quantile(groupB, na.rm=T)[2], col="red", lwd=3)	# first quantile bar for second cohort	
	segments(.9, median(groupB, na.rm=T), 1.1, median(groupB, na.rm=T), col="red", lwd=3)	# median bar for second cohort, etc
	segments(.95, quantile(groupB, na.rm=T)[4], 1.05, quantile(groupB, na.rm=T)[4], col="red", lwd=3)	# first quantile bar for second cohort
	segments(1.0,quantile(groupB, na.rm=T)[2],1.0,quantile(groupB, na.rm=T)[4], col="red", lwd=3)
# legend	
pr = signif(t.test(groupA,groupB, paired=FALSE)$p.value,digits=2)			# This outputs the p-value of the SPEARMAN correlation but does not go below e-16 and I can't figure out the fix
pletter=substitute(paste("   ", italic("p"), " = ", pr), list(pr=signif(pr[1], digits=2)))	# Makes the r-value with italicized r

legend(legendposition, bg='white', bty="n", cex=1, text.col="red", legend=c(as.expression(pletter)))

################################################################################  
# SUPPLEMENTAL FIGURE ONE
################################################################################# 

#$$$$$$ #$$$$$$ #$$$$$$ #$$$$$$ #$$$$$$ #$$$$$$ #$$$$$$ #$$$$$$
#$$$$$$ #$$$$$$ #$$$$$$ #$$$$$$ #$$$$$$ #$$$$$$ #$$$$$$ #$$$$$$
# PANEL S1A

# NOTE: Need to load "histo.dot.plot" function at the end of this code

library(openxlsx)
survive_raw=read.xlsx("~/Dropbox/Aging\ BXD\ Study/1\ _\ Manuscript\ -\ Evan\ Multi-Omic/aTableS1_AgingDB.xlsx", na.strings="")
survive=subset(survive_raw, survive_raw$CauseOfDeath=="Harvest" & survive_raw$Sex=="F") 
SacAge = survive$AgeAtDeath # gets the animal's age at sacrifice
Histo.Dot.Plot(SacAge) 

# Histograms were merged in Illustrator, and dimensions were shifted to get them to align and colors to match.

#$$$$$$ #$$$$$$ #$$$$$$ #$$$$$$ #$$$$$$ #$$$$$$ #$$$$$$ #$$$$$$
#$$$$$$ #$$$$$$ #$$$$$$ #$$$$$$ #$$$$$$ #$$$$$$ #$$$$$$ #$$$$$$
# PANEL S1B

# NOTE: Need to load "XY.Corr" function at the end of this code (search for "XYCORR.FIND")

strains=c("C57","DBA","B6D2","D2B6","BXD1","BXD2","BXD5","BXD6","BXD8","BXD9","BXD11","BXD12","BXD13","BXD14","BXD15","BXD16","BXD18","BXD19","BXD20","BXD21","BXD22","BXD23","BXD24","BXD24a","BXD25","BXD27","BXD28","BXD29","BXD30","BXD31","BXD32","BXD33","BXD34","BXD35","BXD36","BXD37","BXD38","BXD39","BXD40","BXD41","BXD42","BXD43","BXD44","BXD45","BXD48","BXD48a","BXD49","BXD50","BXD51","BXD52","BXD53","BXD54","BXD55","BXD56","BXD59","BXD60","BXD61","BXD62","BXD63","BXD64","BXD65","BXD65a","BXD65b","BXD66","BXD67","BXD68","BXD69","BXD70","BXD71","BXD72","BXD73","BXD73a","BXD73b","BXD74","BXD75","BXD76","BXD77","BXD78","BXD79","BXD81","BXD83","BXD84","BXD85","BXD86","BXD87","BXD88","BXD89","BXD90","BXD91","BXD93","BXD94","BXD95","BXD98","BXD99","BXD100","BXD101","BXD102")
Gelman=c(NA,NA,NA,NA,594,479,528,687,617,816,750,738,439,493,798,642,742,904,NA,NA,743,763,835,NA,NA,711,765,703,737,577,440,NA,NA,NA,NA,NA,NA,NA,NA,NA,NA,NA,NA,NA,NA,NA,NA,NA,NA,NA,NA,NA,NA,NA,NA,NA,NA,NA,NA,NA,NA,NA,NA,NA,NA,NA,NA,NA,NA,NA,NA,NA,NA,NA,NA,NA,NA,NA,NA,NA,NA,NA,NA,NA,NA,NA,NA,NA,NA,NA,NA,NA,NA,NA,NA,NA,NA)
Lang=c(704,645,NA,NA,NA,463,NA,717,618,NA,797,671,NA,428,NA,681,NA,893,NA,NA,697,NA,775,NA,NA,860,798,792,815,NA,704,821,942,NA,NA,NA,689,747,610,NA,575,NA,NA,NA,NA,NA,NA,NA,NA,NA,NA,NA,NA,NA,NA,NA,NA,NA,NA,NA,NA,NA,NA,NA,NA,NA,NA,NA,NA,NA,NA,NA,NA,NA,NA,NA,NA,NA,NA,NA,NA,NA,NA,NA,NA,NA,NA,NA,NA,NA,NA,NA,NA,NA,NA,NA,NA)
Us_CD=c(802,630,933,771,NA,NA,553,636,452,507,650,712,342,489,723,614,527,723,NA,NA,NA,NA,690,NA,NA,758,876,804,NA,537,691,NA,NA,NA,NA,NA,NA,614,585,NA,NA,728,551,756,695,645,NA,NA,862,NA,763,NA,NA,645,NA,651,756,723,640,NA,824,NA,726,516,NA,NA,615,721,NA,NA,702,NA,820,NA,769,NA,802,NA,417,NA,523,769,658,804,811,NA,709,706,925,NA,NA,602,623,NA,698,688,861)
Us_HFD=c(699,646,NA,838,477,NA,350,604,722,469,602,872,292,406,727,604,435,603,NA,NA,563,NA,541,NA,NA,574,660,697,NA,590,612,NA,579,NA,NA,NA,NA,NA,397,NA,NA,707,616,695,547,635,NA,NA,561,NA,591,NA,NA,539,NA,546,722,547,727,NA,534,NA,606,616,730,NA,654,642,NA,NA,699,NA,770,524,619,NA,676,NA,480,NA,NA,500,563,717,583,NA,731,540,816,NA,NA,438,551,NA,600,482,760)

# Left Plot
XY.Corr(FileName=NA, Lang, Gelman, addlegend=T, addlabels=T, strains=strains, xlimits=c(300,1000), ylimits=c(300,1000),  ptbgcolor="grey", legendalign="bottomright", type="Pearson", cex.legend=2)

# Right Plot
XY.Corr(FileName=NA, Us_CD, Lang, addlegend=T, strains=strains, xlimits=c(300,1000), ylimits=c(300,1000),  ptbgcolor="red", legendalign="bottomright", type="Pearson", cex.legend=2)
XY.Corr(FileName=NA, Us_CD, Gelman, add=T, addlegend=T, strains=strains, xlimits=c(300,1000), ylimits=c(300,1000),  ptbgcolor="blue", legendalign="topleft", type="Pearson", cex.legend=2)

#$$$$$$ #$$$$$$ #$$$$$$ #$$$$$$ #$$$$$$ #$$$$$$ #$$$$$$ #$$$$$$
#$$$$$$ #$$$$$$ #$$$$$$ #$$$$$$ #$$$$$$ #$$$$$$ #$$$$$$ #$$$$$$
# PANEL S1C

# Histogram drawn by hand to scale using the ANCOVA results in Table S1, sheet "Var_Explained_of_All_Data"

#$$$$$$ #$$$$$$ #$$$$$$ #$$$$$$ #$$$$$$ #$$$$$$ #$$$$$$ #$$$$$$
#$$$$$$ #$$$$$$ #$$$$$$ #$$$$$$ #$$$$$$ #$$$$$$ #$$$$$$ #$$$$$$
# PANEL S1D

library(survival) # For Kaplan-Meier graphs
library(openxlsx)

survive_raw=read.xlsx("~/Dropbox/Aging\ BXD\ Study/1\ _\ Manuscript\ -\ Evan\ Multi-Omic/aTableS1_AgingDB.xlsx", na.strings="")
survive=subset(survive_raw, survive_raw$CauseOfDeath=="Natural" | survive_raw$CauseOfDeath=="Euthanized") 
survive=subset(survive_raw, survive_raw$CauseOfDeath=="Natural" | survive_raw$CauseOfDeath=="Euthanized" & survive_raw$Sex=="F")
# This sets the animals as "alive" or "dead" for graphing & lifespan calculations
lifestatus = survive$AgeIfStillAlive # tells if animal is dead (1) or alive (0)
i=1
for(i in 1:length(lifestatus)) {
	if(lifestatus[i]==0)  {
		lifestatus[i]=1 		# sets animal as dead if its "AgeIfStillAlive" is zero (UTHSC's dead marker)
	}
	else {lifestatus[i]=0}		# sets animal as alive if it has an AgeIfStillAlive (UTHSC's "still alive" marker)
}
xlim=c(0, 1300)
xaxp=c(0, 1300, 2)
ylim=c(0, 1)
yaxp=c(0, 1, 2)

### BXD13 vs BXD91 lifespan, diets combined
time=subset(survive$AgeIfStillAlive2, survive$StrainNameCurrent == "BXD13" | survive$StrainNameCurrent == "BXD91" & survive$Sex== "F")
time2=subset(survive$AgeAtDeath, survive$StrainNameCurrent == "BXD13" | survive$StrainNameCurrent == "BXD91" & survive$Sex== "F")
event=subset(lifestatus, survive$StrainNameCurrent == "BXD13" | survive$StrainNameCurrent == "BXD91" & survive$Sex== "F")
strains=subset(survive$StrainNameCurrent, survive$StrainNameCurrent == "BXD13" | survive$StrainNameCurrent == "BXD91" & survive$Sex== "F")
group=subset(survive$DietCode, survive$StrainNameCurrent == "BXD13" | survive$StrainNameCurrent == "BXD91" & survive$Sex== "F")
xy=survfit(Surv(time, time2, event)~strains)
plot(xy, lwd=3, xlab="Days", ylab="Ratio Alive", main="BXD Aging Colony", bty="L", col=c("red","orange","yellow","green","blue","purple"), xlim=xlim, xaxp=xaxp, ylim=ylim, yaxp=yaxp, las=1)

print(survfit(Surv(time, time2, event)~strains), print.rmean=T)
survdiff(Surv(time2, event)~strains)


#$$$$$$ #$$$$$$ #$$$$$$ #$$$$$$ #$$$$$$ #$$$$$$ #$$$$$$ #$$$$$$
#$$$$$$ #$$$$$$ #$$$$$$ #$$$$$$ #$$$$$$ #$$$$$$ #$$$$$$ #$$$$$$
# PANEL S1E

library(survival) # For Kaplan-Meier graphs
library(openxlsx)

survive_raw=read.xlsx("~/Dropbox/Aging\ BXD\ Study/1\ _\ Manuscript\ -\ Evan\ Multi-Omic/aTableS1_AgingDB.xlsx", na.strings="")
survive=subset(survive_raw, survive_raw$CauseOfDeath=="Natural" | survive_raw$CauseOfDeath=="Euthanized") 
survive=subset(survive_raw, survive_raw$CauseOfDeath=="Natural" | survive_raw$CauseOfDeath=="Euthanized" & survive_raw$Sex=="F")
# This sets the animals as "alive" or "dead" for graphing & lifespan calculations
lifestatus = survive$AgeIfStillAlive # tells if animal is dead (1) or alive (0)
i=1
for(i in 1:length(lifestatus)) {
	if(lifestatus[i]==0)  {
		lifestatus[i]=1 		# sets animal as dead if its "AgeIfStillAlive" is zero (UTHSC's dead marker)
	}
	else {lifestatus[i]=0}		# sets animal as alive if it has an AgeIfStillAlive (UTHSC's "still alive" marker)
}
xlim=c(0, 1300)
xaxp=c(0, 1300, 2)
ylim=c(0, 1)
yaxp=c(0, 1, 2)

### LEFT PANEL: BXD9 lifespan, split by diet; legend hand-drawn
time=c(subset(survive$AgeIfStillAlive2, survive$DietCode == "HF" & survive$StrainNameCurrent == "BXD9"), subset(survive$AgeIfStillAlive2, survive$DietCode == "CD" & survive$StrainNameCurrent == "BXD9"))
time2=c(subset(survive$AgeAtDeath, survive$DietCode == "HF" & survive$StrainNameCurrent == "BXD9"), subset(survive$AgeAtDeath, survive$DietCode == "CD" & survive$StrainNameCurrent == "BXD9"))
event=c(subset(lifestatus, survive$DietCode == "HF" & survive$StrainNameCurrent == "BXD9"), subset(lifestatus, survive$DietCode == "CD" & survive$StrainNameCurrent == "BXD9"))
strains=c(subset(survive$DietCode, survive$DietCode == "HF" & survive$StrainNameCurrent == "BXD9"), subset(survive$DietCode, survive$DietCode == "CD" & survive$StrainNameCurrent == "BXD9"))
xy=survfit(Surv(time, time2, event)~strains)
plot(xy, lwd=3, xlab="Days", ylab="Ratio Alive", main="BXD Aging Colony", bty="L", col=c("grey","black"), xlim=xlim, xaxp=xaxp, ylim=ylim, yaxp=yaxp, las=1)
print(survfit(Surv(time, time2, event)~strains), print.rmean=T)
survdiff(Surv(time2, event)~strains)


### RIGHT PANEL: BXD65 lifespan, split by diet; legend hand-drawn
time=c(subset(survive$AgeIfStillAlive2, survive$DietCode == "HF" & survive$StrainNameCurrent == "BXD65"), subset(survive$AgeIfStillAlive2, survive$DietCode == "CD" & survive$StrainNameCurrent == "BXD65"))
time2=c(subset(survive$AgeAtDeath, survive$DietCode == "HF" & survive$StrainNameCurrent == "BXD65"), subset(survive$AgeAtDeath, survive$DietCode == "CD" & survive$StrainNameCurrent == "BXD65"))
event=c(subset(lifestatus, survive$DietCode == "HF" & survive$StrainNameCurrent == "BXD65"), subset(lifestatus, survive$DietCode == "CD" & survive$StrainNameCurrent == "BXD65"))
strains=c(subset(survive$DietCode, survive$DietCode == "HF" & survive$StrainNameCurrent == "BXD65"), subset(survive$DietCode, survive$DietCode == "CD" & survive$StrainNameCurrent == "BXD65"))
xy=survfit(Surv(time, time2, event)~strains)
plot(xy, lwd=3, xlab="Days", ylab="Ratio Alive", main="BXD Aging Colony", bty="L", col=c("grey","black"), xlim=xlim, xaxp=xaxp, ylim=ylim, yaxp=yaxp, las=1)
print(survfit(Surv(time, time2, event)~strains), print.rmean=T)
survdiff(Surv(time2, event)~strains)


#$$$$$$ #$$$$$$ #$$$$$$ #$$$$$$ #$$$$$$ #$$$$$$ #$$$$$$ #$$$$$$
#$$$$$$ #$$$$$$ #$$$$$$ #$$$$$$ #$$$$$$ #$$$$$$ #$$$$$$ #$$$$$$
# PANEL S1F
# First we need to calculate the diet-specific lifespans for all strains, as so:
survive=subset(survive_raw, survive_raw$CauseOfDeath=="Natural" | survive_raw$CauseOfDeath=="Euthanized" & survive_raw$Sex=="F")
time=survive$AgeIfStillAlive2
time2=survive$AgeAtDeath
event=lifestatus
strains=survive$StrainNameCurrent
xy=survfit(Surv(time, time2, event)~strains)

time=subset(survive$AgeIfStillAlive2, survive$DietCode == "CD")
time2=subset(survive$AgeAtDeath, survive$DietCode == "CD")
event=subset(lifestatus, survive$DietCode == "CD")
strains=subset(survive$StrainNameCurrent, survive$DietCode == "CD")
xy=survfit(Surv(time, time2, event)~strains)
plot(xy, lwd=3, xlab="Days", ylab="Ratio Alive", main="BXD Aging Colony", bty="L", col=c("red","orange","yellow","green","blue","purple"), xlim=xlim, xaxp=xaxp, ylim=ylim, yaxp=yaxp, las=1)
#legend("topright", lty=1, lwd=3, bty="n", col=c("red","blue","green","purple"), c("BXD13", "BXD91", "BXD9", "BXD73"))
print(survfit(Surv(time, time2, event)~strains), print.rmean=T)

time=subset(survive$AgeIfStillAlive2, survive$DietCode == "HF")
time2=subset(survive$AgeAtDeath, survive$DietCode == "HF")
event=subset(lifestatus, survive$DietCode == "HF")
strains=subset(survive$StrainNameCurrent, survive$DietCode == "HF")
xy=survfit(Surv(time, time2, event)~strains)
plot(xy, lwd=3, xlab="Days", ylab="Ratio Alive", main="BXD Aging Colony", bty="L", col=c("red","orange","yellow","green","blue","purple"), xlim=xlim, xaxp=xaxp, ylim=ylim, yaxp=yaxp, las=1)
#legend("topright", lty=1, lwd=3, bty="n", col=c("red","blue","green","purple"), c("BXD13", "BXD91", "BXD9", "BXD73"))
print(survfit(Surv(time, time2, event)~strains), print.rmean=T)

# Now  we take the CD and HFD averages for each strain, remove those with n < 6, and put them in the following vectors:

library(gplots)
strains=c("B6D2F1","BXD100","BXD101","BXD102","BXD12","BXD13","BXD16","BXD161","BXD168","BXD169","BXD170","BXD171","BXD18","BXD19","BXD24","BXD27","BXD28","BXD29","BXD32","BXD34","BXD39","BXD40","BXD43","BXD45","BXD48","BXD48a","BXD51","BXD53","BXD56","BXD6","BXD60","BXD61","BXD62","BXD63","BXD65","BXD65b","BXD66","BXD68","BXD69","BXD70","BXD73","BXD73b","BXD77","BXD79","BXD8","BXD84","BXD85","BXD86","BXD87","BXD89","BXD9","BXD90","BXD91","BXD95","BXD98","B6","D2B6F1","D2")
CD_mean=c(902,728,688,843,NA,342,563,720,862,NA,496,NA,527,723,690,758,876,774,691,NA,614,585,728,756,695,645,862,763,645,636,651,756,723,640,885,726,516,NA,615,711,710,820,784,417,452,746,658,692,NA,709,507,687,899,602,623,800,801,650)
CD_se=c(3.22E+01,4.02E+01,4.69E+01,5.70E+01,NA,7.03E+01,4.06E+01,8.51E+01,5.41E+01,NA,5.42E+01,NA,7.14E+01,4.98E+01,5.15E+01,5.45E+01,3.62E+01,4.18E+01,5.01E+01,NA,5.96E+01,6.37E+01,4.82E+01,3.08E+01,4.09E+01,3.91E+01,2.94E+01,4.98E+01,6.73E+01,3.91E+01,5.00E+01,3.52E+01,3.72E+01,8.40E+01,1.67E+01,2.52E+01,5.97E+01,NA,5.47E+01,3.18E+01,2.76E+01,3.21E+01,5.39E+01,3.64E+01,5.56E+01,7.09E+01,5.42E+01,1.24E+02,NA,4.58E+01,2.87E+01,3.78E+01,5.58E+01,3.60E+01,3.97E+01,3.87E+01,4.36E+01,2.30E+01)
HFD_mean=c(755,600,492,729,972,292,608,NA,842,620,431,624,435,603,541,574,650,697,612,579,NA,397,707,695,546,635,561,591,539,604,546,722,547,727,534,606,616,543,654,585,682,770,676,480,NA,500,563,717,583,731,475,540,804,438,551,666,752,644)
HFD_se=c(56.1,38.2,34.8,34,97.1,35,28,NA,46.9,45,33.3,24.5,33.4,39.4,35,54,38.2,31.4,22.7,60.8,NA,42.3,42.7,69,30.4,28.2,72.8,55.2,35.5,18.5,24.8,39.5,40.5,32.7,32,31.1,65.6,117.6,28,34.3,34.9,30.9,49.8,47.6,NA,29.7,34.8,38.7,59.9,38.2,26.2,50,40.4,38.2,41.9,23.3,62.5,19.1)

## Now to plot the data, we need to load "XY.Corr" function written at the end of this document (search for XYCORR.FIND)!
XY.Corr(FileName=NA, CD_mean, HFD_mean, CD_se, HFD_se, addlegend=T, strains=strains, xlimits=c(200,950), ylimits=c(200,950),  ptbgcolor="grey", legendalign="bottomright", type="Pearson", cex.legend=2)


#$$$$$$ #$$$$$$ #$$$$$$ #$$$$$$ #$$$$$$ #$$$$$$ #$$$$$$ #$$$$$$
#$$$$$$ #$$$$$$ #$$$$$$ #$$$$$$ #$$$$$$ #$$$$$$ #$$$$$$ #$$$$$$
# PANEL S1G
library(gplots)
library(openxlsx) 
allweight=read.xlsx("~/Dropbox/Aging\ BXD\ Study/1\ _\ Manuscript\ -\ Evan\ Multi-Omic/aTableS1_AgingDB.xlsx", sheet=2, na.strings="")

AgeOverTime=cbind(allweight$AgeAtWeight01, allweight$AgeAtWeight02, allweight$AgeAtWeight03, allweight$AgeAtWeight04, allweight$AgeAtWeight05, allweight$AgeAtWeight06, allweight$AgeAtWeight07, allweight$AgeAtWeight08, allweight$AgeAtWeight09, allweight$AgeAtWeight10, allweight$AgeAtWeight11, allweight$AgeAtWeight12, allweight$AgeAtWeight13, allweight$AgeAtWeight14, allweight$AgeAtWeight15)
rownames(AgeOverTime)=allweight$EarTagNumberCurrent
WeightOverTime=cbind(allweight$Weight01, allweight$Weight02, allweight$Weight03, allweight$Weight04, allweight$Weight05, allweight$Weight06, allweight$Weight07, allweight$Weight08, allweight$Weight09, allweight$Weight10, allweight$Weight11, allweight$Weight12, allweight$Weight13, allweight$Weight14, allweight$Weight15)
rownames(WeightOverTime)=allweight$EarTagNumberCurrent
TimeOnDietOverTime=cbind(allweight$DaysOnDietAtWeight01, allweight$DaysOnDietAtWeight02, allweight$DaysOnDietAtWeight03, allweight$DaysOnDietAtWeight04, allweight$DaysOnDietAtWeight05, allweight$DaysOnDietAtWeight06, allweight$DaysOnDietAtWeight07, allweight$DaysOnDietAtWeight08, allweight$DaysOnDietAtWeight09, allweight$DaysOnDietAtWeight10, allweight$DaysOnDietAtWeight11, allweight$DaysOnDietAtWeight12, allweight$DaysOnDietAtWeight13, allweight$DaysOnDietAtWeight14, allweight$DaysOnDietAtWeight15)
rownames(TimeOnDietOverTime)=allweight$EarTagNumberCurrent
CohortTags = subset(allweight$EarTagNumberCurrent, allweight$StrainNameCurrent=="BXD16")
CohortTagsCD = subset(allweight$EarTagNumberCurrent, allweight$StrainNameCurrent=="BXD16" & allweight$DietCode=="CD" & allweight$Sex=="F")
CohortTagsHF = subset(allweight$EarTagNumberCurrent, allweight$StrainNameCurrent=="BXD16" & allweight$DietCode=="HF" & allweight$Sex=="F")

# Initialize the graph and then plot for CD
plotCI(c(NA), c(NA), ylim=c(10, 105), main="BXD16", xlim=c(0, 1000), bty="L", xlab="Age [days]", ylab="Weight [grams]", las=1, yaxp=c(0, 100, 4), xaxp=c(0, 1000, 4))
for(i in 1:length(CohortTagsHF)) {
plotCI(subset(AgeOverTime[,1:15], allweight$EarTagNumberCurrent==CohortTagsCD[i]), subset(WeightOverTime[,1:15], allweight$EarTagNumberCurrent==CohortTagsCD[i]), pch=21, pt.bg="grey", col="grey", type="o", cex.main=2, add=T)
}
legend("topleft", c("CD","HFD"), cex=2, pt.bg=c("grey","black"), box.lwd=0, pch=21, col=c("black","black"))

# Initialize the graph and then plot for HFD
for(i in 1:length(CohortTagsHF)) {
plotCI(subset(AgeOverTime[,1:15], allweight$EarTagNumberCurrent==CohortTagsHF[i]), subset(WeightOverTime[,1:15], allweight$EarTagNumberCurrent==CohortTagsHF[i]), pch=21, pt.bg="black", col="black", type="o", cex.main=2, add=T)
}


#$$$$$$ #$$$$$$ #$$$$$$ #$$$$$$ #$$$$$$ #$$$$$$ #$$$$$$ #$$$$$$
#$$$$$$ #$$$$$$ #$$$$$$ #$$$$$$ #$$$$$$ #$$$$$$ #$$$$$$ #$$$$$$
# PANEL S1H
library(gplots)
library(openxlsx) 
allweight=read.xlsx("~/Dropbox/Aging\ BXD\ Study/1\ _\ Manuscript\ -\ Evan\ Multi-Omic/aTableS1_AgingDB.xlsx", sheet=2, na.strings="")

AgeOverTime=cbind(allweight$AgeAtWeight01, allweight$AgeAtWeight02, allweight$AgeAtWeight03, allweight$AgeAtWeight04, allweight$AgeAtWeight05, allweight$AgeAtWeight06, allweight$AgeAtWeight07, allweight$AgeAtWeight08, allweight$AgeAtWeight09, allweight$AgeAtWeight10, allweight$AgeAtWeight11, allweight$AgeAtWeight12, allweight$AgeAtWeight13, allweight$AgeAtWeight14, allweight$AgeAtWeight15)
rownames(AgeOverTime)=allweight$EarTagNumberCurrent
WeightOverTime=cbind(allweight$Weight01, allweight$Weight02, allweight$Weight03, allweight$Weight04, allweight$Weight05, allweight$Weight06, allweight$Weight07, allweight$Weight08, allweight$Weight09, allweight$Weight10, allweight$Weight11, allweight$Weight12, allweight$Weight13, allweight$Weight14, allweight$Weight15)
rownames(WeightOverTime)=allweight$EarTagNumberCurrent
TimeOnDietOverTime=cbind(allweight$DaysOnDietAtWeight01, allweight$DaysOnDietAtWeight02, allweight$DaysOnDietAtWeight03, allweight$DaysOnDietAtWeight04, allweight$DaysOnDietAtWeight05, allweight$DaysOnDietAtWeight06, allweight$DaysOnDietAtWeight07, allweight$DaysOnDietAtWeight08, allweight$DaysOnDietAtWeight09, allweight$DaysOnDietAtWeight10, allweight$DaysOnDietAtWeight11, allweight$DaysOnDietAtWeight12, allweight$DaysOnDietAtWeight13, allweight$DaysOnDietAtWeight14, allweight$DaysOnDietAtWeight15)
rownames(TimeOnDietOverTime)=allweight$EarTagNumberCurrent
CohortTags = subset(allweight$EarTagNumberCurrent, allweight$StrainNameCurrent=="BXD100")
CohortTagsCD = subset(allweight$EarTagNumberCurrent, allweight$StrainNameCurrent=="BXD100" & allweight$DietCode=="CD" & allweight$Sex=="F")
CohortTagsHF = subset(allweight$EarTagNumberCurrent, allweight$StrainNameCurrent=="BXD100" & allweight$DietCode=="HF" & allweight$Sex=="F")

# Initialize the graph and then plot for CD
plotCI(c(NA), c(NA), ylim=c(10, 105), main="BXD16", xlim=c(0, 1000), bty="L", xlab="Age [days]", ylab="Weight [grams]", las=1, yaxp=c(0, 100, 4), xaxp=c(0, 1000, 4))
for(i in 1:length(CohortTagsHF)) {
plotCI(subset(AgeOverTime[,1:15], allweight$EarTagNumberCurrent==CohortTagsCD[i]), subset(WeightOverTime[,1:15], allweight$EarTagNumberCurrent==CohortTagsCD[i]), pch=21, pt.bg="grey", col="grey", type="o", cex.main=2, add=T)
}
legend("topleft", c("CD","HFD"), cex=2, pt.bg=c("grey","black"), box.lwd=0, pch=21, col=c("black","black"))

# Initialize the graph and then plot for HFD
for(i in 1:length(CohortTagsHF)) {
plotCI(subset(AgeOverTime[,1:15], allweight$EarTagNumberCurrent==CohortTagsHF[i]), subset(WeightOverTime[,1:15], allweight$EarTagNumberCurrent==CohortTagsHF[i]), pch=21, pt.bg="black", col="black", type="o", cex.main=2, add=T)
}


#$$$$$$ #$$$$$$ #$$$$$$ #$$$$$$ #$$$$$$ #$$$$$$ #$$$$$$ #$$$$$$
#$$$$$$ #$$$$$$ #$$$$$$ #$$$$$$ #$$$$$$ #$$$$$$ #$$$$$$ #$$$$$$
# PANEL S1I

# Need to also load the "Dot.Plot" function at the end of this script (search for "DOTPLOT.FIND")
library(openxlsx)
library(outliers)

bloodmetabs=read.xlsx("~/Dropbox/Aging\ BXD\ Study/1\ _\ Manuscript\ -\ Evan\ Multi-Omic/aTableS1_AgingDB.xlsx", sheet=3, na.strings="")

stripchart(subset(bloodmetabs$ALPL, bloodmetabs$Diet=="CD"), at=0.8, pch=21, cex=2, bg="grey", method="jitter", xlim=c(0.6, 1.4), ylim=c(0, 400), vertical=T)
stripchart(subset(bloodmetabs$ALPL, bloodmetabs$Diet=="HF"), at=1.2, pch=21, cex=2, bg="black", method="jitter", add=T, vertical=T)
CatA=subset(bloodmetabs$ALPL, bloodmetabs$Diet=="CD")
CatB=subset(bloodmetabs$ALPL, bloodmetabs$Diet=="HF")
t.test(CatA, CatB)$p.value

stripchart(subset(bloodmetabs$ALPL, bloodmetabs$AgeCohort == 1), at=0.8, pch=21, cex=2, bg="green", method="jitter", xlim=c(0.6, 1.4), ylim=c(0,400), vertical=T)
stripchart(subset(bloodmetabs$ALPL, bloodmetabs$AgeCohort == 4), at=1.2, pch=21, cex=2, bg="blue", method="jitter", add=T, vertical=T)
CatA=subset(bloodmetabs$ALPL, bloodmetabs$AgeCohort == 1)
CatB=subset(bloodmetabs$ALPL, bloodmetabs$AgeCohort == 4)
t.test(CatA, CatB)$p.value

stripchart(subset(bloodmetabs$ALPL, bloodmetabs$AgeCohort == 1 & bloodmetabs$Diet=="CD"), at=0.8, pch=21, cex=2, bg="lightgreen", method="jitter", xlim=c(0.6, 1.4), ylim=c(0,400), vertical=T)
stripchart(subset(bloodmetabs$ALPL, bloodmetabs$AgeCohort == 4 & bloodmetabs$Diet=="CD"), at=1.2, pch=21, cex=2, bg="darkgreen", method="jitter", add=T, vertical=T)
CatA=subset(bloodmetabs$ALPL, bloodmetabs$AgeCohort == 1 & bloodmetabs$Diet=="CD")
CatB=subset(bloodmetabs$ALPL, bloodmetabs$AgeCohort == 4 & bloodmetabs$Diet=="CD")
t.test(CatA, CatB)$p.value

stripchart(subset(bloodmetabs$ALPL, bloodmetabs$AgeCohort == 1 & bloodmetabs$Diet=="HF"), at=0.8, pch=21, cex=2, bg="lightblue", method="jitter", xlim=c(0.6, 1.4), ylim=c(0,400), vertical=T)
stripchart(subset(bloodmetabs$ALPL, bloodmetabs$AgeCohort == 4 & bloodmetabs$Diet=="HF"), at=1.2, pch=21, cex=2, bg="darkblue", method="jitter", add=T, vertical=T)
CatA=subset(bloodmetabs$ALPL, bloodmetabs$AgeCohort == 1 & bloodmetabs$Diet=="HF")
CatB=subset(bloodmetabs$ALPL, bloodmetabs$AgeCohort == 4 & bloodmetabs$Diet=="HF")
t.test(CatA, CatB)$p.value


######################################################################################################################################################
######################################################################################################################################################
######################################################################################################################################################
######################################################################################################################################################
######################################################################################################################################################


#### BELOW IS FUNCTIONS THAT WERE WRITTEN FOR THE PLOTTING CODE ####

######################################################################################################################################################
######################################################################################################################################################
# Histogram with dot plot function
######################################################################################################################################################


Histo.Dot.Plot <- function (x, chart.title, breaks, axisexpansion, ylimits, xlimits, xlabel, ylabel, colors, addlegend, legendalign, figurewidth, figureheight, cex.pt) {
if(missing(chart.title)) { chart.title="" }
if(missing(axisexpansion)) { axisexpansion=0.02 } # For increasing the axis boundaries
if(missing(xlimits)) { xlimits=c(min(x, na.rm=T)-axisexpansion*min(x, na.rm=T), max(x, na.rm=T)+axisexpansion*max(x, na.rm=T)) }
if(missing(ylimits)) { ylimits=c(0, length(x)/2) }
if(missing(xlabel)) { xlabel="" }
if(missing(ylabel)) { ylabel="" }
if(missing(colors)) { colors=c("brown3","brown2", "grey", "grey","chartreuse3","chartreuse2") } # Colors for the dot plot and colors for the histogram
if(missing(addlegend)) { addlegend=T }
if(missing(legendalign)) { legendalign="topleft" }
if(missing(breaks)) { breaks=8 }
if(missing(figurewidth)) { figurewidth=7 }
if(missing(figureheight)) { figureheight=7 }
if(missing(cex.pt)) { cex.pt=1 }

dev.new(width=figurewidth, height=figureheight)			# Draws a new window with both the histogram area and the dot plot area
par(mfrow=c(2,1), mar=c(2,3,0,1)+1)
mat <- matrix(c(1,2), 2, 1, byrow = TRUE)
nf <- layout(mat, widths = c(8,8), height = c(6,2), TRUE)


hist(x, breaks=breaks, xlim=xlimits, ylim=ylimits,col=colors, main=NA, las=1)
par(bty="l")

x=sort(x)
x_len=length(x)
x1=x[1:((x_len)/4)]
x2=x[((x_len)/4):((2*x_len)/4)]
x3=x[((2*x_len)/4):((3*x_len)/4)]
x4=x[((3*x_len)/4):((4*x_len)/4)]

stripchart(x1, vertical=F, cex=cex.pt, pch=19, col=c("red"), method="jitter", ylim=c(.8, 1.2), xlim=xlimits)
stripchart(x2, vertical=F, cex=cex.pt, pch=19, col=c("yellow3"), method="jitter", add=T)
stripchart(x3, vertical=F, cex=cex.pt, pch=19, col=c("chartreuse2"), method="jitter", add=T)
stripchart(x4, vertical=F, cex=cex.pt, pch=19, col=c("deepskyblue2"), method="jitter", add=T)
}


############################################################################################
######################## **** DOT PLOT, FUNCTION SEPARATOR ***** #######################
############################################################################################
## "DOTPLOT.FIND" ##
# Up to four input variables are allowed
Dot.Plot <- function (x, y, chart.title, axisexpansion, ylimits, xlimits, xlabel, ylabel, colors, addlegend, stackmethod, legendalign, addquantiles, datatype, group, digits, cex.axis, cex.legend, cex.lab, cex.pt, pch, pch2, bg, bty) {
if(missing(y)) { y=c(NA) }
if(missing(chart.title)) { chart.title="" }
if(missing(axisexpansion)) { axisexpansion=0.02 } # For increasing the axis boundaries
if(missing(ylimits)) { ylimits=c(min(x, y, na.rm=T)-axisexpansion*min(x, y, na.rm=T), max(x, y, na.rm=T)+axisexpansion*max(x, y, na.rm=T)) }
if(missing(xlabel)) { xlabel="" }
if(missing(ylabel)) { ylabel="" }
if(missing(colors)) { colors=c("grey","black", "red", "red","green","blue") } # First two colors are for OUTSIDE dot colors of x and y, next two colors are for the quantile colors for x and y, then for w and u
if(missing(addlegend)) { addlegend=T }
if(missing(stackmethod)) { stackmethod="jitter" }
if(missing(legendalign)) { legendalign="topleft" }
if(missing(addquantiles)) { addquantiles="All" }
if(missing(datatype)) { datatype="pheno" } # For calculating fold change
if(missing(group)) { group=c("CD","HFD") }
if(missing(digits)) { digits=1 } # for how far to round for numbers on the y-axis
if(missing(cex.axis)) { cex.axis=1 }
if(missing(cex.legend)) { cex.legend=1 }
if(missing(cex.lab)) { cex.lab=1 }
if(missing(cex.pt)) { cex.pt=2 }
if(missing(xlimits)) { xlimits=c(0.6,1.0) }
if(missing(pch)) { pch=21 }	# This is the point style (now: circle with inside and outside colors)
if(missing(pch2)) { pch2=21 }	
if(missing(bg)) { bg=c("white", "white") }	# This is the outside color of the object (inside is specified in "colors" above)
if(missing(bty)) { bty="n" }	# This is the default for the box type (L; o, l, 7, c, u, ], or n allowed)

x=x[!is.na(x)]
y=y[!is.na(y)]

xq=quantile(x, na.rm=T)
yq=quantile(y, na.rm=T)

minimum = min(x,y)
maximum = max(x,y)

par(bty=bty)
stripchart(x, vertical=T, at=0.8, pch=pch, cex=cex.pt, col=bg[1], bty="l", bg=colors[1], method=stackmethod, offset=0.5, ylim=ylimits, xlim=xlimits, las=1)
stripchart(y, add=T, vertical=T, cex=cex.pt, pch=pch2, at=1.2, col=bg[2], bg=colors[2], method=stackmethod, offset=0.5)
#stripchart(w, add=T, vertical=T, cex=2, pch=16, at=1.6, col=colors[5], method=stackmethod)
#stripchart(u, add=T, vertical=T, cex=2, pch=16, at=2.0, col=colors[6], method=stackmethod)

avgZ = median(x, na.rm=T)
avgY = median(y, na.rm=T)

if(datatype=="pheno") {
	foldchange=signif(max(avgZ, avgY)/min(avgZ, avgY), digits=3)
	totalrange=signif(maximum/minimum, digits=2)
}
if(datatype=="RNA") {
	foldchange=signif(2^(max(avgZ, avgY, na.rm=T)-min(avgZ, avgY, na.rm=T)), digits=3)
	totalrange=signif(2^(maximum-minimum), digits=2)
}

if(addquantiles=="All") {
	segments(0.75, xq[2], 0.85, xq[2], col=colors[3], lwd=3)	# first quantile bar for first cohort
	segments(1.15, yq[2], 1.25, yq[2], col=colors[4], lwd=3)	# first quantile bar for second cohort
	segments(0.7, xq[3], 0.9, xq[3], col=colors[3], lwd=3)	# median bar for first cohort
	segments(1.1, yq[3], 1.3, yq[3], col=colors[4], lwd=3)	# median bar for second cohort
	segments(0.75, xq[4], 0.85, xq[4], col=colors[3], lwd=3)	# third quantile bar for first cohort
	segments(1.15, yq[4], 1.25, yq[4], col=colors[4], lwd=3)	# third quantile bar for second cohort
	
	segments(0.8,xq[2],0.8,xq[4], col=colors[3], lwd=3)
	segments(1.2,yq[2],1.2,yq[4], col=colors[4], lwd=3)
	}

if(addquantiles=="Median") {
	segments(0.7, xq[3], 0.9, xq[3], col=colors[3], lwd=3)	# median bar for first cohort
	segments(1.1, yq[3], 1.3, yq[3], col=colors[4], lwd=3)	# median bar for second cohort
	}
	
if(addlegend==T) {
	significance = signif(t.test(x,y, paired=FALSE)$p.value,digits=2) # TWO TAILED PAIRED WELCH'S T-TEST!
	groups=c(group[1],group[2])
	pvalue = substitute(paste(italic(p), " = ", significance))
	foldchange = paste("Fold Change = ", foldchange)
	totalrange = paste("Total Range = ", totalrange)

	legend(legendalign, legend=c(groups, as.expression(pvalue), foldchange), cex=cex.legend, col=c(colors[1],colors[2],"white", "white"), box.lwd=0, pch=c(19,19,NA,NA))
	}
}


######################################################################################################################################################
######################################################################################################################################################
# XY.Corr function "XYCORR.FIND"
######################################################################################################################################################
XY.Corr <- function (FileName, x, y, xSE, ySE, chart.title, ylimits, xlimits, xaxp, yaxp, xlabel, ylabel, addlegend, ptbgcolor, ptcolor, addlabels, legendalign, type,strains, add, pch, digits, axisexpansion, cex.legend, cex.main, cex.pt) {
if(missing(xSE)) { xSE=c(NA) }
if(missing(ySE)) { ySE=c(NA) }
if(missing(chart.title)) { chart.title="" }
if(missing(xlabel)) { xlabel="" }
if(missing(ylabel)) {	ylabel="" }
if(missing(addlegend)) { addlegend=F }
if(missing(ptcolor)) { ptcolor="black" }
if(missing(ptbgcolor)) { ptbgcolor=ptcolor }
if(missing(addlabels)) { addlabels=F }
if(missing(legendalign)) { legendalign="topleft" }
if(missing(type)) { type="Pearson" } # Default is Pearson
if(missing(add)) { add=F } # Default is Pearson
if(missing(pch)) { pch=21 } # Default is with black ring around; switch to 19 if you just want large pure color, 20 for small pure color
if(missing(strains)) { strains=NA }
if(missing(digits)) { digits=1 } # Use to change # significant decimal points on x and y axis.
if(missing(axisexpansion)) { axisexpansion=0.02 } # For increasing the axis boundaries
if(missing(cex.legend)) { cex.legend=1 }
if(missing(cex.main)) { cex.main=1.5 }
if(missing(cex.pt)) { cex.pt=1.6 }

if(type=="Pearson") {

if(missing(xlimits)) { xlimits=c(min(x, na.rm=T)-axisexpansion*min(x, na.rm=T), max(x, na.rm=T)+axisexpansion*max(x, na.rm=T)) }
if(missing(ylimits)) { ylimits=c(min(y, na.rm=T)-axisexpansion*min(y, na.rm=T), max(y, na.rm=T)+axisexpansion*max(y, na.rm=T)) }
if(missing(xaxp)) { xaxp=c(round(min(x, na.rm=T), digits), round(max(x, na.rm=T), digits=digits), 2) }
if(missing(yaxp)) { yaxp=c(round(min(y, na.rm=T), digits=digits), round(max(y, na.rm=T), digits), 2) }

plotCI(x=x, y=y, uiw=xSE, err=T, liw=NA, gap=0, pch=pch, pt.bg=ptbgcolor, lwd=3, cex=cex.pt, cex.main=cex.main, xlab=xlabel, ylab=ylabel, bty="L", main=chart.title, barcol="black", col=ptcolor, ylim=ylimits, xlim=xlimits, add=add, xaxp=xaxp, yaxp=yaxp)
plotCI(x=x, y=y, uiw=ySE, gap=0, liw=NA, lwd=3, add=T, pch=pch, cex=(cex.pt-.1), pt.bg=ptbgcolor, barcol="black", col=ptcolor) # Adds Y (vertical) error bars
}

if(type=="Spearman" || type=="Spearman-Kendall") {
xy=cbind(x,y)
x=x[complete.cases(xy)]
y=y[complete.cases(xy)]

x=rank(x, na.last="keep")
y=rank(y, na.last="keep")

x_len=rank(x, na.last=NA)
y_len=rank(y, na.last=NA)

if(missing(xlimits)) { xlimits=c(0, length(x_len)+1) }
if(missing(ylimits)) { ylimits=c(0, length(y_len)+1) }

plotCI(x=x, y=y, err=F, liw=NA, gap=0, pch=pch, pt.bg=ptbgcolor, lwd=3, cex=cex.pt, xlab=xlabel, ylab=ylabel, bty="L", main=chart.title, barcol="black", col=ptcolor, ylim=ylimits, xlim=xlimits, add=add, xaxp=c(0, length(x_len)+1, 2), yaxp=c(0,length(y_len)+1, 2))
}

abline(lm(y~x), lty=1, lwd=cex.legend, col=ptcolor) # Linear regression
if(addlegend==T) { 
	if(type=="Pearson") {
		correl=cor(x, y, method=c("pearson"), use="complete.obs")
		pval=signif(summary(lm(y~x))$coefficients[8], digits=2)
		rletter=substitute(paste(italic(r), "  = ", correl), list(correl=signif(correl[1], digits=2)))	# Makes the r-value with italicized r
		pletter=substitute(paste(italic(p), " = ", pval))												# Makes the p-value with italicized p
		totallegend=c(as.expression(rletter), as.expression(pletter))
		
		if(pval < 0.05) { legend(legendalign, bg='white', bty="n", cex=cex.legend, text.col="red", legend=totallegend) }
		if(pval >= 0.05) { legend(legendalign, bg='white', bty="n", cex=cex.legend, text.col="black", legend=totallegend) }
		
		}
	if(type=="Spearman") {
		rhoc = cor(x, y, method=c("spearman"), use="complete.obs")
		pr = signif(summary(lm(y~x))$coefficients[8], digits=2)
		rletter=substitute(paste(italic("rho"), " = ", rhoc), list(rhoc=signif(rhoc[1], digits=2)))	# Makes the r-value with italicized r
		pletter=substitute(paste("   ", italic(p), " = ", pr))												# Makes the p-value with italicized p
		totallegend=c(as.expression(rletter), as.expression(pletter))

		if(pr < 0.05) { legend(legendalign, bg='white', bty="n", cex=cex.legend, text.col="red", legend=totallegend) }
		if(pr >= 0.05) { legend(legendalign, bg='white', bty="n", cex=cex.legend, text.col="black", legend=totallegend) }
	}	
	if(type=="Spearman-Kendall") {
		rhoc = sqrt(summary(lm(y~x))$r.squared)
		pr = signif(summary(lm(y~x))$coefficients[8], digits=2)
		taub = cor.test(x,y,method="kendall")$estimate[1]
		ptau = signif(cor.test(x,y,method="kendall")$p.value[1], digits=2)
		rletter=substitute(paste(italic("rho"), " = ", rhoc), list(rhoc=signif(rhoc[1], digits=2)))	# Makes the r-value with italicized r
		pletter=substitute(paste("   ", italic(p), " = ", pr))												# Makes the p-value with italicized p
		taubletter=substitute(paste("   ", italic(tau), "  = ", taub), list(taub=signif(taub[1], digits=2)))	# Makes the tau-b value with italicized tau
		ptauletter=substitute(paste("   ", italic(p), " = ", ptau))												# Makes the p-value with italicized p
		
		totallegend=c(as.expression(rletter), as.expression(pletter), as.expression(taubletter), as.expression(ptauletter))
		
		if(pr < 0.05) { legend(legendalign, bg='white', bty="n", cex=cex.legend, text.col="red", legend=totallegend) }
		if(pr >= 0.05) { legend(legendalign, bg='white', bty="n", cex=cex.legend, text.col="red", legend=totallegend) }	
	}
}
	
if(addlabels==T) text(x,y+0.05*abs(y),labels=paste(strains, "  "), col="black", pos=1) #Adds datapoint labels. Only does this for Pearson.
}
```
