## Supplementary material for "Multi-Omic Profiling of the Liver Across Diets and Age in a Diverse Mouse Population": All code used to generate figures: b_Figure2_HelperFile_QTLs.r

```
library(qtl2)

##To perform QTL, need to creatae yaml file - collection of various files like genotype and phenotype files:
#if you want to run QTL on p traits, then make a pheno file in the same format as "p_bxd_pheno.csv", using a n x p format
#Also supply a geno file, like "p_bxd_geno.csv" with the genotype of each of the n mice.
#We have supplied those files in this case. Because this code takes a very long time to run (particularly the permutation testing), we have included the output .RDS files as well.


# First need to generate pheno and geno files
# The geno file must have the full genotypes for every individual in the BXD.pheno file

setwd("b_Figure2_HelperFiles_QTLs/")
write_control_file("p_bxd_2.yaml", crosstype = "risib", gmap_file = "bxd_gmap.csv",
pheno_file = "p_bxd_pheno.csv", # HAVE TO GENERATE THIS FILE
geno_file = "p_bxd_geno.csv",  # ALSO HAVE TO GENERATE THIS FILE
alleles = c("B", "D"),sep = ",",xchr = "X", overwrite = TRUE)

#Time to run QTL LOD SCORES, prepare the input files and run:
bd <- read_cross2("p_bxd_2.yaml")
map <- insert_pseudomarkers(bd$gmap, step=1)
pr <- calc_genoprob(bd, map, error_prob=0.002, cores = 4)
kinship_loco <- calc_kinship(pr, "loco", cores=4)
Xcovar <- get_x_covar(bd)
out_pg <- scan1(pr, bd$pheno, kinship_loco, Xcovar=Xcovar) ## If it takes too long to run, can split up the pheno e.g. bd$pheno[,1:100] and parallelize that way
saveRDS(out_pg, file = paste("Block_All_LODS.rds", sep=""))

## Now for calculating QTL P-values

setwd("b_Figure2_HelperFiles_QTLs/")

bd <- read_cross2("p_bxd_2.yaml")
map <- insert_pseudomarkers(bd$gmap, step=1)
pr <- calc_genoprob(bd, map, error_prob=0.002)
apr <- genoprob_to_alleleprob(pr)
kinship <- calc_kinship(pr)
grid <- calc_grid(bd$gmap, step=1)
pr_grid <- probs_to_grid(pr, grid)
kinship_loco <- calc_kinship(pr, "loco")
Xcovar <- get_x_covar(bd)
#out_pg <- scan1(pr, bd$pheno, kinship, Xcovar=Xcovar) # Could also calculate LODS together with p-values
k<-1
if(!(file.exists("Block_1_pvals.rds"))){
operm <- scan1perm(pr, bd$pheno[,1:5], kinship_loco, Xcovar=Xcovar, n_perm=100) #1342 (v near start ):
saveRDS(operm, "Block_All_pvals.rds")
}
```
