## Supplementary material for "Multi-Omic Profiling of the Liver Across Diets and Age in a Diverse Mouse Population": All code used to generate figures: b_Figure2_Plotting_Code_vF.r

```
################################################################################  
# First priority is loading the multi-omics dataset, which will be used throughout
## This is the INDIVIDUAL data
################################################################################# 

setwd("~/Dropbox/Aging\ BXD\ Study/1\ _\ Manuscript\ -\ Evan\ Multi-Omic/")

AllData_header=read.table("aTableS2_ALL_DATA.csv", nrows=8, row.names=1, header=T, sep=",")
AllData_header=AllData_header[,3:ncol(AllData_header)]
AllData=read.table("aTableS2_ALL_DATA.csv", row.names=1, header=T,  na.strings="", sep=",", skip=8, stringsAsFactors=FALSE)
AllData=AllData[1:nrow(AllData),3:ncol(AllData)]

AllData_CD=matrix(NA, nrow=nrow(AllData), ncol=length(AllData))
for(i in 1:length(AllData)) {
	if(as.character(AllData_header["Diet",i])=="CD") {   AllData_CD[,i]=as.numeric(as.character(AllData[,i]))	}
} 
rownames(AllData_CD)=rownames(AllData)

AllData_HF=matrix(NA, nrow=nrow(AllData), ncol=length(AllData))
for(i in 1:length(AllData)) {
	if(as.character(AllData_header["Diet",i])=="HF") {   AllData_HF[,i]=as.numeric(as.character(AllData[,i]))	}
}
rownames(AllData_HF)=rownames(AllData)

AllData_young=matrix(NA, nrow=nrow(AllData), ncol=length(AllData))
for(i in 1:length(AllData)) {
	if(as.character(AllData_header["Age",i])<=420) {	AllData_young[,i]=as.numeric(as.character(AllData[,i]))	}
}
rownames(AllData_young)=rownames(AllData)

AllData_old=matrix(NA, nrow=nrow(AllData), ncol=length(AllData))
for(i in 1:length(AllData)) {
	if(as.character(AllData_header["Age",i])>=420) {	AllData_old[,i]=as.numeric(as.character(AllData[,i]))	}
}
rownames(AllData_old)=rownames(AllData)

#  This loads a datafile with just the top 3772 matching mRNAs and proteins 
# Note that the gene name in column 1 and column 3 might vary, this is due to aliases/synonyms, but they are all matched.
setwd("~/Dropbox/Aging\ BXD\ Study/1\ _\ Manuscript\ -\ Evan\ Multi-Omic/")

Top3772Data_header=read.table("b_Figure2_HelperFile_Matching3772_v2.csv", nrows=8, row.names=1, header=T, sep=",")
Top3772Data_header=Top3772Data_header[,3:ncol(Top3772Data_header)]
Top3772Data=read.table("b_Figure2_HelperFile_Matching3772_v2.csv", row.names=1, header=T,  na.strings="", sep=",", skip=8, stringsAsFactors=FALSE)
Top3772Data=Top3772Data[1:nrow(Top3772Data),3:ncol(Top3772Data)]

Top3772Data_CD=matrix(NA, nrow=nrow(Top3772Data), ncol=length(Top3772Data))
for(i in 1:length(Top3772Data)) {
	if(as.character(Top3772Data_header["Diet",i])=="CD") {   Top3772Data_CD[,i]=as.numeric(as.character(Top3772Data[,i]))	}
} 
rownames(Top3772Data_CD)=rownames(Top3772Data)

Top3772Data_HF=matrix(NA, nrow=nrow(Top3772Data), ncol=length(Top3772Data))
for(i in 1:length(Top3772Data)) {
	if(as.character(Top3772Data_header["Diet",i])=="HF") {   Top3772Data_HF[,i]=as.numeric(as.character(Top3772Data[,i]))	}
}
rownames(Top3772Data_HF)=rownames(Top3772Data)

Top3772Data_young=matrix(NA, nrow=nrow(Top3772Data), ncol=length(Top3772Data))
for(i in 1:length(Top3772Data)) {
	if(as.character(Top3772Data_header["Age",i])<=420) {	Top3772Data_young[,i]=as.numeric(as.character(Top3772Data[,i]))	}
}
rownames(Top3772Data_young)=rownames(Top3772Data)

Top3772Data_old=matrix(NA, nrow=nrow(Top3772Data), ncol=length(Top3772Data))
for(i in 1:length(Top3772Data)) {
	if(as.character(Top3772Data_header["Age",i])>=420) {	Top3772Data_old[,i]=as.numeric(as.character(Top3772Data[,i]))	}
}
rownames(Top3772Data_old)=rownames(Top3772Data)

################################################################################  
# FIGURE TWO
################################################################################# 

#$$$$$$ #$$$$$$ #$$$$$$ #$$$$$$ #$$$$$$ #$$$$$$ #$$$$$$ #$$$$$$
#$$$$$$ #$$$$$$ #$$$$$$ #$$$$$$ #$$$$$$ #$$$$$$ #$$$$$$ #$$$$$$
# PANEL 2A
library(outliers)
library(ggplot2)
library(gplots)

setwd("~/Dropbox/Aging\ BXD\ Study/1\ _\ Manuscript\ -\ Evan\ Multi-Omic/")

mRNAdat_CD=Top3772Data_CD[1:3772,]
mRNAdat_HF=Top3772Data_HF[1:3772,]

Protdat_CD=Top3772Data_CD[3773:dim(Top3772Data_CD)[1],]
Protdat_HF=Top3772Data_HF[3773:dim(Top3772Data_HF)[1],]

## Plots LEFT panel
Diet_FC_mRNA_All=rowMeans(mRNAdat_CD,na.rm=T)-rowMeans(mRNAdat_HF,na.rm=T)
Diet_p_mRNA_All=NA

for(i in 1:length(Diet_FC_mRNA_All)) {
	Diet_p_mRNA_All[i]=t.test(mRNAdat_CD[i,], mRNAdat_HF[i,])$p.value
}
Diet_padjust_mRNA_All=p.adjust(Diet_p_mRNA_All,method="BH")

log2FC_rm=rm.outlier(Diet_FC_mRNA_All, fill=T)	# Replace huge outliers with NA
pvalue=rm.outlier(Diet_p_mRNA_All,fill=T) # Replace huge outliers with NA
neglog10p=-log(pvalue, 10)
plotCI(log2FC_rm, neglog10p, bty="L", pch=21, col="blue", pt.bg="black", ylim=c(0,22), xlim=c(-1.1,1.1), las=1)

length(which(Diet_padjust_mRNA_All<0.05))		# Sets Bonferroni-corrected p-value as cutoff
length(which(Diet_p_mRNA_All<0.05)) # Counts number of peptides affected at the nominal p-value

## Plots RIGHT panel
Diet_FC_Protein_All=rowMeans(Protdat_CD,na.rm=T)-rowMeans(Protdat_HF,na.rm=T)
Diet_p_Protein_All=NA

for(i in 1:length(Diet_FC_Protein_All)) {
	Diet_p_Protein_All[i]=t.test(Protdat_CD[i,], Protdat_HF[i,])$p.value
}
Diet_padjust_Protein_All=p.adjust(Diet_p_Protein_All,method="BH")

log2FC_rm=rm.outlier(Diet_FC_Protein_All, fill=T)	# Replace huge outliers with NA
pvalue=rm.outlier(Diet_p_Protein_All,fill=T) # Replace huge outliers with NA
neglog10p=-log(pvalue, 10)
plotCI(log2FC_rm, neglog10p, bty="L", pch=21, col="red", pt.bg="black", ylim=c(0,42), xlim=c(-1.1,1.1), las=1)

length(which(Diet_padjust_Protein_All<0.05))		# Sets Bonferroni-corrected p-value as cutoff
length(which(Diet_p_Protein_All<0.05)) # Counts number of peptides affected at the nominal p-value


#$$$$$$ #$$$$$$ #$$$$$$ #$$$$$$ #$$$$$$ #$$$$$$ #$$$$$$ #$$$$$$
#$$$$$$ #$$$$$$ #$$$$$$ #$$$$$$ #$$$$$$ #$$$$$$ #$$$$$$ #$$$$$$
# PANEL 2B 
# These data are taken from Table S1, sheet "Var_Explained_of_All_Data", separated by data type

library(vioplot)

# LEFT plot
VarExplained_mRNA_Genotype=c(18,23,26,31,28,27,27,14,34,19,25,21,22,23,26,30,26,31,25,16,42,37,24,38,28,19,20,27,36,36,19,27,27,27,29,56,36,27,23,25,25,24,28,20,30,30,19,26,38,31,29,42,19,29,32,27,44,27,31,22,21,16,35,31,20,25,26,27,33,20,22,24,32,25,28,31,23,21,16,19,31,21,24,27,39,26,27,37,30,34,25,34,21,33,34,27,33,26,22,25,35,29,18,32,26,17,24,28,30,29,22,18,25,28,29,27,29,41,34,34,29,30,22,22,26,22,29,21,32,34,29,25,23,23,32,37,26,23,15,22,23,25,35,23,35,33,33,34,25,32,30,27,31,29,36,27,23,25,41,21,30,35,29,29,22,19,27,24,24,21,27,26,24,23,31,26,22,27,28,29,32,20,25,30,30,23,18,21,32,23,25,25,21,34,31,27,20,26,29,36,27,26,24,24,54,37,25,19,27,25,24,33,24,51,39,30,21,24,27,36,26,21,23,32,24,25,28,20,17,44,27,25,25,23,28,49,23,31,27,29,38,36,32,22,26,29,25,30,28,20,41,35,28,43,32,29,58,31,23,23,19,20,32,27,25,26,23,22,26,30,25,26,39,22,29,26,21,40,29,33,34,21,28,23,24,31,28,28,32,25,27,30,24,29,32,36,30,25,24,33,27,25,40,30,23,26,26,22,24,23,23,20,28,26,21,23,26,32,23,23,25,26,42,24,40,39,45,18,24,24,34,17,34,21,34,22,29,26,73,25,25,26,37,28,40,33,22,21,28,26,24,23,25,54,18,25,20,23,27,27,26,38,21,26,42,21,23,24,34,23,26,27,24,38,21,18,23,24,21,22,23,22,23,42,32,34,29,27,22,24,18,23,31,25,27,23,36,28,35,28,30,43,31,26,33,24,25,27,27,53,29,23,24,28,34,25,40,22,35,26,26,45,34,28,27,52,30,23,28,27,33,18,29,65,31,30,21,26,24,24,29,24,19,37,34,29,30,22,26,22,25,28,27,26,24,33,27,30,22,36,25,37,31,24,26,28,28,27,29,25,29,28,26,25,26,28,29,27,33,31,46,30,26,33,28,28,23,24,22,26,31,27,20,28,27,27,20,26,25,24,27,31,41,27,27,22,28,33,26,24,35,24,29,27,28,25,28,28,30,25,27,26,29,30,27,28,28,20,32,26,33,19,35,26,29,27,26,21,22,25,32,21,58,24,19,29,33,29,28,29,19,25,26,20,31,24,65,22,28,26,27,32,33,26,46,29,31,32,21,21,63,24,25,30,23,25,22,28,23,25,28,22,28,30,25,22,31,23,21,21,74,34,26,28,23,46,31,22,25,31,20,27,30,18,21,28,25,25,26,36,20,25,25,36,35,31,22,36,42,27,30,23,24,30,29,38,32,21,35,25,28,41,29,24,30,32,25,26,19,25,27,23,22,25,26,30,26,28,37,27,40,24,26,33,26,29,28,21,31,27,23,18,31,20,29,24,16,26,27,23,52,32,23,32,25,20,22,24,17,21,28,21,26,20,22,22,26,43,29,27,24,25,27,33,33,21,36,30,27,29,23,53,24,41,20,27,32,25,46,28,27,21,28,21,25,21,33,27,29,26,24,27,46,28,23,21,25,29,29,31,31,31,39,26,18,25,42,28,15,23,21,26,59,41,32,31,14,27,27,27,26,26,24,23,27,18,43,34,21,32,26,17,29,33,44,23,28,31,31,25,27,30,50,21,26,24,20,27,33,29,29,31,22,19,22,21,24,46,24,29,18,23,28,25,28,28,29,23,21,28,38,46,24,42,24,30,32,19,21,46,22,34,23,22,24,31,30,23,32,26,33,27,21,36,63,15,22,23,22,32,19,25,28,33,22,25,27,31,21,25,40,27,26,40,29,21,27,31,41,25,21,27,26,21,30,30,29,33,27,28,19,26,22,24,26,21,19,22,25,21,24,30,20,42,25,27,29,23,47,27,25,30,23,22,25,22,21,30,36,21,22,19,21,27,24,28,32,18,58,26,27,41,17,23,25,26,23,17,30,23,26,26,28,25,19,27,23,30,24,23,24,26,33,27,22,28,30,19,27,21,29,24,26,28,17,38,26,31,41,27,28,28,28,25,34,51,28,32,30,21,31,32,23,23,32,27,29,21,28,33,29,66,25,29,36,43,23,26,19,26,21,32,25,24,24,41,29,39,52,45,34,27,25,25,33,20,21,26,27,32,27,29,27,22,38,33,19,66,27,28,31,29,38,23,23,17,24,30,33,27,20,47,29,35,24,24,29,27,37,25,29,30,22,26,29,31,16,32,21,22,28,25,25,27,33,18,35,28,26,41,39,27,22,70,29,17,26,34,32,26,19,21,30,23,28,25,35,23,20,28,44,18,46,27,30,31,28,23,27,20,50,30,21,33,27,28,25,20,22,25,28,20,31,52,27,24,29,22,29,19,57,30,28,34,25,25,23,28,23,23,22,29,27,34,28,39,46,28,24,33,27,27,30,64,22,36,38,22,22,24,25,31,22,22,32,23,22,27,26,17,26,31,26,24,27,32,34,44,34,45,27,29,23,23,24,32,22,23,42,24,25,30,27,43,24,23,33,44,32,26,44,23,30,27,42,22,18,19,24,25,23,30,31,30,30,30,26,27,24,18,19,26,26,16,35,32,30,23,31,32,31,24,25,22,24,36,25,36,29,45,30,25,25,31,19,24,16,19,27,20,22,35,33,55,25,29,32,26,20,25,24,21,33,24,24,23,30,31,22,37,18,23,29,20,24,32,21,29,24,25,18,31,30,22,30,23,31,37,28,29,27,21,22,13,30,28,26,25,37,26,24,35,28,26,28,25,20,22,66,28,27,24,43,18,30,25,31,21,27,26,20,30,21,28,21,23,26,22,29,20,23,17,21,20,26,31,26,62,28,28,25,15,23,43,18,22,30,57,29,27,25,25,31,25,22,30,32,33,30,32,26,21,31,28,26,24,39,18,27,24,22,41,22,29,35,31,28,21,19,28,25,27,24,22,26,28,29,19,23,26,28,28,44,30,19,24,31,21,22,27,27,17,35,34,35,27,26,32,39,20,25,24,26,23,26,26,21,28,23,19,29,29,19,21,22,25,21,24,24,29,22,19,24,25,28,31,23,60,25,23,64,28,28,28,23,31,42,27,34,27,21,24,24,30,25,26,35,29,24,21,23,26,26,26,30,27,23,38,29,22,38,32,27,30,23,26,29,28,27,26,30,21,26,29,34,23,32,29,34,23,24,35,22,25,32,28,28,24,80,18,24,23,29,25,30,26,19,25,21,22,31,19,32,30,25,22,32,26,31,27,19,27,22,26,23,34,24,32,18,24,28,33,20,39,28,25,21,27,33,23,26,30,21,28,36,22,31,23,23,31,49,23,23,26,32,31,22,27,25,31,23,25,22,28,28,28,24,26,20,23,24,28,27,28,25,37,32,29,27,27,36,30,23,32,32,12,27,26,35,29,31,57,22,54,20,26,30,20,23,56,31,29,27,30,26,18,26,31,37,30,29,27,27,24,33,31,25,43,23,27,23,29,34,22,32,21,27,21,59,27,71,48,24,35,23,24,25,26,20,30,26,23,23,19,19,31,19,30,35,26,22,22,52,21,23,14,29,34,85,25,32,24,30,20,40,19,34,29,20,18,34,30,25,17,26,24,28,29,25,27,23,29,20,31,21,27,30,15,72,36,28,37,27,29,18,20,32,19,23,23,29,30,25,22,28,20,24,30,32,22,25,39,22,27,36,20,32,26,26,26,25,25,30,23,40,32,38,23,23,32,25,24,27,24,22,29,37,87,36,28,22,47,23,27,25,31,21,49,39,36,85,25,36,24,21,54,27,30,20,26,27,28,20,22,30,28,24,26,23,23,28,23,58,29,21,23,23,25,31,24,22,42,27,26,22,30,25,29,28,26,47,28,28,31,24,26,45,24,27,28,26,29,23,26,44,23,25,22,32,17,50,24,25,27,37,20,25,35,24,31,25,20,19,27,30,29,22,27,27,30,28,32,53,35,34,33,31,34,22,28,25,22,24,19,33,19,25,24,27,41,33,23,32,21,31,26,21,28,27,27,18,19,22,26,16,21,21,22,19,33,45,29,27,31,25,34,24,25,30,23,23,20,29,21,58,20,29,21,27,28,30,27,26,24,23,20,31,30,20,35,25,18,28,22,34,36,29,21,30,33,26,28,33,24,27,21,25,23,25,21,24,29,18,58,28,24,27,55,21,26,26,28,22,26,24,35,20,28,19,32,20,26,28,24,22,72,27,28,27,37,27,31,30,28,22,19,34,18,22,47,70,26,25,21,34,20,28,20,35,19,27,35,23,29,27,23,22,25,25,32,28,22,26,18,18,25,26,28,21,20,27,22,58,30,17,29,20,24,86,39,32,21,38,25,38,44,18,29,25,21,23,18,26,41,27,27,53,18,27,60,34,41,43,26,32,20,23,29,24,30,22,27,34,28,25,22,29,30,30,61,31,45,25,29,26,27,25,25,75,28,18,24,19,20,22,27,28,31,21,25,30,34,23,25,21,25,45,35,20,32,31,23,32,30,12,31,27,26,21,63,29,27,31,31,19,24,28,28,19,66,23,29,27,27,30,32,37,79,26,26,33,67,33,20,25,18,42,26,38,26,29,23,57,29,22,26,34,26,43,26,26,24,21,30,28,28,23,21,24,23,23,31,22,31,25,20,22,20,27,33,30,21,28,28,68,20,29,23,44,23,21,24,24,44,21,26,25,24,34,25,34,28,30,29,20,38,29,32,24,29,27,22,26,26,29,25,35,30,29,19,29,22,21,25,25,16,30,20,25,19,28,31,26,20,25,29,35,21,34,47,24,21,61,24,22,25,27,27,26,35,26,27,26,32,24,30,26,36,33,22,63,19,25,23,25,27,22,23,20,22,28,27,23,28,26,32,47,27,33,59,31,25,15,29,22,21,22,26,33,24,28,86,27,26,19,33,24,29,24,23,22,28,21,25,65,26,29,20,31,25,25,26,29,27,14,31,24,30,24,29,29,27,30,77,23,28,21,32,24,27,23,22,27,24,20,28,29,27,35,17,30,24,27,23,23,27,26,24,25,31,27,27,26,28,25,32,29,25,30,21,22,31,24,24,21,26,25,27,27,28,28,32,26,80,29,42,21,29,27,22,30,36,29,35,34,42,27,29,66,37,45,21,78,24,21,16,27,22,30,31,32,30,25,30,26,32,30,20,33,31,25,24,25,26,30,21,36,23,55,19,23,28,28,22,22,23,25,22,26,23,29,23,26,28,25,26,31,30,24,24,34,29,28,26,22,29,29,31,22,24,28,20,26,29,24,31,24,33,24,21,27,30,28,24,28,24,35,37,33,24,38,19,23,23,27,31,25,15,29,30,21,23,28,28,25,18,20,27,23,33,29,27,25,24,19,29,31,32,38,22,30,20,34,23,28,31,24,23,21,24,21,26,32,24,27,25,26,21,22,21,22,24,21,24,29,28,21,32,32,23,29,33,19,27,21,29,22,53,27,24,27,90,24,29,33,25,25,25,75,25,31,28,31,18,41,26,29,27,28,31,29,25,23,25,26,19,27,20,26,27,21,28,32,25,33,25,27,18,25,23,24,43,25,32,20,25,23,29,25,28,76,24,28,24,26,35,26,23,26,23,28,29,55,27,23,26,20,24,36,22,27,24,23,26,18,23,26,28,28,22,18,30,18,32,41,25,30,27,33,19,29,18,24,23,22,22,46,35,27,25,25,20,19,34,32,34,22,32,25,22,21,26,26,25,32,33,24,35,22,31,22,23,23,24,27,24,27,22,24,22,20,23,19,26,38,23,30,25,33,27,20,26,20,28,27,32,27,26,28,71,74,27,34,26,24,23,26,22,31,26,28,27,28,24,28,32,20,26,27,26,31,21,24,29,35,23,18,25,47,29,27,27,27,29,21,30,30,32,25,27,24,33,30,25,52,19,27,28,30,20,28,43,21,27,30,22,35,19,22,23,27,35,27,35,24,25,26,23,30,27,24,23,25,30,30,24,26,81,28,26,29,24,19,25,28,16,34,21,28,23,26,28,35,46,38,26,28,26,35,31,23,30,21,33,21,25,35,29,24,21,24,22,20,29,28,32,28,32,23,44,17,26,35,24,24,26,83,28,31,31,30,24,25,31,25,21,69,25,19,29,30,27,39,26,25,20,19,31,26,23,30,48,21,19,29,28,27,33,26,25,19,56,24,34,63,29,28,26,26,58,33,16,23,28,36,37,38,19,26,29,23,27,32,39,27,21,31,21,35,35,25,29,40,27,39,25,28,43,21,44,21,26,28,35,27,23,24,28,27,22,25,33,27,19,26,25,32,26,24,22,24,23,23,45,27,28,25,22,29,23,30,20,20,29,28,25,36,29,28,25,24,28,19,31,32,17,25,20,23,35,82,34,26,13,27,23,33,24,24,22,29,20,22,26,23,27,22,23,26,25,20,24,26,25,30,26,29,34,44,29,20,32,26,16,16,23,23,30,38,20,26,22,23,23,22,19,31,24,27,33,28,61,23,22,50,25,36,22,33,27,23,20,48,26,44,35,31,15,26,30,24,31,25,25,27,29,36,37,24,20,26,25,34,35,36,27,22,26,25,17,24,22,28,36,17,29,32,24,20,31,23,27,23,24,23,23,24,24,45,23,28,29,86,31,23,30,31,32,24,18,26,26,25,24,25,28,19,19,27,30,35,31,28,31,30,25,24,21,36,23,20,17,29,21,26,27,35,21,25,20,39,67,31,30,20,21,29,35,16,30,23,24,24,26,32,28,25,29,20,33,22,35,21,29,33,28,23,27,70,18,26,26,29,24,27,27,24,21,25,27,24,30,31,27,27,22,24,23,25,37,28,30,15,28,23,23,33,22,27,27,28,18,28,30,31,29,24,26,28,41,32,34,21,29,24,61,19,35,26,20,25,23,67,32,19,26,23,25,25,27,26,20,30,27,34,31,42,23,31,78,25,22,24,25,29,22,23,21,28,22,29,23,22,26,34,21,28,18,28,36,20,25,25,16,16,28,27,19,23,21,29,27,26,23,26,24,34,27,33,29,26,22,25,27,28,27,22,32,29,32,54,34,30,27,22,24,23,25,29,28,50,23,28,22,30,23,18,27,18,41,20,24,25,25,37,18,27,33,33,25,16,26,25,31,28,24,19,24,26,24,28,19,29,23,25,22,43,28,23,26,25,34,27,31,37,32,29,18,21,42,26,19,24,24,24,32,27,26,26,25,27,19,31,33,21,18,27,23,25,21,27,29,22,24,26,53,22,21,25,31,23,22,20,25,20,17,26,27,37,27,25,22,35,26,34,29,27,27,23,25,24,28,21,27,23,22,20,22,31,24,24,31,22,27,28,27,24,24,25,22,20,60,25,21,20,24,22,25,30,26,21,26,23,16,21,22,28,34,21,22,36,28,24,19,23,23,28,23,25,21,25,27,27,81,21,27,23,24,28,25,28,20,31,30,35,22,22,53,34,27,26,20,27,24,27,64,32,28,41,25,27,34,23,32,27,26,23,52,41,30,29,31,19,25,18,22,28,25,28,25,19,32,31,22,19,30,28,28,18,16,25,27,34,26,78,24,28,25,18,55,34,26,27,29,22,25,24,26,34,25,20,23,20,23,24,20,28,20,21,25,29,26,19,19,30,35,28,27,30,21,28,29,20,29,22,30,32,29,29,21,25,37,71,28,26,22,24,24,19,30,24,23,17,25,32,29,23,26,29,31,23,24,25,39,29,17,31,24,22,72,32,29,39,22,29,21,13,30,27,24,27,24,26,25,24,23,22,35,21,32,35,14,26,17,26,30,27,25,25,23,23,21,35,38,33,29,29,23,21,46,27,26,36,25,27,26,26,29,24,22,32,31,24,21,29,21,22,37,26,25,23,20,30,27,26,20,18,21,25,29,23,30,25,24,56,27,26,36,23,52,29,21,38,28,28,45,36,33,25,19,23,25,24,27,52,24,37,27,25,22,23,37,25,27,21,24,19,24,18,26,23,20,27,24,31,24,21,23,21,20,26,33,33,28,25,23,27,22,30,31,27,23,26,27,23,22,31,29,29,28,37,39,28,24,18,19,27,26,21,28,33,79,30,24,22,21,76,21,26,25,24,28,22,34,35,19,23,23,31,27,28,17,22,25,30,24,25,26,25,22,25,29,30,23,22,25,27,22,21,15,27,45,45,45,22,25,25,25,25,23,33,20,34,22,25,25,20,19,26,28,26,32,25,58,23,30,27,32,26,33,20,26,23,22,21,39,29,26,24,26,29,24,21,25,31,32,30,29,35,32,22,29,31,22,21,31,27,23,26,42,20,31,29,37,21,47,64,26,24,31,70,20,26,27,20,26,19,32,28,23,21,30,39,25,28,19,28,29,32,32,36,22,37,30,33,32,24,48,22,26,43,66,28,38,28,24,24,25,32,36,65,35,21,24,34,30,24,25,34,25,37,31)
VarExplained_mRNA_Diet=c(45,29,25,23,23,20,20,24,18,21,19,18,18,19,17,17,15,15,16,19,15,14,20,18,19,13,16,14,13,16,16,14,15,11,16,14,11,13,15,12,13,12,12,12,11,11,14,13,9,14,11,9,10,13,9,10,14,10,11,9,10,9,9,11,10,11,10,11,8,12,8,13,12,9,6,11,12,11,10,15,11,9,11,11,9,9,8,12,8,6,10,9,10,8,7,7,12,6,8,8,9,10,8,8,7,9,8,9,9,7,9,8,9,8,7,6,10,5,11,7,7,11,9,7,8,8,9,8,8,8,10,7,9,9,11,10,8,8,7,7,7,8,6,10,10,8,8,8,7,7,8,7,9,9,7,7,7,10,8,9,8,5,8,9,7,7,5,9,8,7,7,9,6,10,7,7,4,7,6,5,7,7,10,10,7,9,6,7,7,7,10,8,6,8,6,5,7,6,8,6,7,8,5,7,7,5,7,6,7,4,7,7,8,5,8,8,9,4,6,8,8,6,7,5,8,5,5,6,6,6,6,8,5,5,5,4,6,6,8,6,5,5,4,6,6,5,6,7,6,5,6,4,5,5,6,6,3,5,6,8,4,5,5,5,8,6,4,5,6,6,5,5,7,4,4,7,5,8,5,6,5,4,4,6,6,6,4,3,6,6,5,8,4,4,4,6,5,7,6,3,6,5,4,4,6,5,8,4,6,5,4,4,8,6,7,5,5,4,5,4,5,5,3,5,5,3,5,4,6,5,5,5,6,5,5,7,4,5,4,4,7,4,5,4,3,4,5,5,4,4,5,5,4,3,5,3,5,4,4,5,6,5,5,5,4,6,5,6,3,3,4,5,3,5,6,3,4,4,5,6,3,4,3,4,4,5,4,3,5,6,4,4,6,5,4,5,4,3,5,3,4,3,4,4,4,4,4,4,3,3,4,4,5,4,3,6,5,6,4,4,4,2,5,5,5,4,5,4,4,4,4,6,2,3,6,4,3,4,4,5,4,3,5,3,5,6,2,4,3,3,3,4,4,5,4,5,6,4,3,3,4,5,4,4,3,4,4,4,4,3,3,2,3,4,3,4,5,3,3,4,2,6,4,3,3,5,5,4,3,5,3,4,4,4,5,3,4,5,3,2,4,4,3,3,2,4,2,3,3,5,4,3,5,3,4,2,5,3,5,4,3,3,4,2,4,2,3,4,4,4,3,4,3,3,2,4,4,4,2,2,1,4,4,4,3,4,3,4,3,4,4,2,5,5,2,6,4,3,4,3,2,2,2,2,4,3,4,2,4,5,3,3,3,4,3,3,2,2,2,4,4,4,3,2,3,5,4,2,3,3,1,2,4,4,4,5,3,2,3,4,2,3,2,3,2,2,4,1,2,3,3,4,3,4,2,3,3,3,2,3,4,3,2,2,3,2,4,2,4,4,4,3,4,3,3,4,4,3,3,2,2,3,5,3,4,3,4,3,2,5,4,5,4,4,3,2,3,2,2,2,3,2,4,5,4,2,3,4,2,2,3,5,3,2,2,3,3,3,2,3,2,3,3,3,2,1,2,2,2,2,2,3,3,3,3,4,2,3,2,2,2,3,4,2,3,3,5,2,3,4,2,3,3,3,5,2,3,3,3,2,3,2,3,2,4,3,3,3,3,2,2,2,2,2,2,3,1,2,2,2,2,2,1,3,4,2,2,3,4,3,5,1,1,2,3,2,3,1,3,4,1,2,4,4,2,3,4,2,4,4,2,4,3,2,3,2,3,4,4,4,2,3,3,3,3,2,2,2,3,2,3,3,3,3,2,1,3,2,3,2,1,2,3,2,4,3,2,3,3,2,2,2,3,2,4,2,1,1,3,2,3,3,2,3,3,3,3,2,2,4,2,3,4,1,2,4,2,2,3,2,2,3,3,1,3,2,3,1,4,4,2,2,2,2,3,2,3,3,2,3,4,2,1,2,4,3,3,1,1,2,1,3,2,3,3,2,3,2,2,3,2,2,1,2,3,3,2,2,1,2,2,1,2,3,3,2,2,2,2,4,2,2,2,2,2,3,3,3,2,3,2,1,2,4,2,2,1,3,2,2,1,2,2,2,3,2,1,2,2,3,3,3,2,2,2,1,2,2,2,1,2,2,2,1,2,1,1,2,2,3,1,2,2,1,2,2,2,2,1,2,3,2,2,2,1,1,3,2,2,2,3,3,1,2,3,1,1,1,2,2,3,2,4,1,2,2,2,2,3,3,2,1,2,1,4,3,2,1,2,1,3,2,2,2,2,1,1,2,2,3,2,2,4,3,3,3,2,2,2,3,2,2,1,2,2,2,3,2,1,2,1,2,2,2,2,1,1,2,1,1,2,2,2,2,1,2,2,2,1,2,2,2,1,2,2,2,3,1,3,1,1,2,1,3,1,1,1,1,2,2,1,1,3,1,2,2,2,1,2,2,2,2,1,1,2,1,1,1,2,2,3,1,2,1,1,1,2,2,1,1,2,3,2,3,2,2,3,1,3,2,1,2,1,2,2,1,2,2,2,1,2,1,1,1,2,1,1,3,2,1,2,0,2,1,2,2,1,1,3,1,0,1,2,1,0,1,2,1,1,2,1,1,2,2,2,1,1,0,1,2,2,1,1,1,1,1,1,1,2,2,2,3,1,0,3,2,2,2,1,2,2,1,2,2,1,2,2,2,2,2,2,1,2,2,2,1,2,2,1,1,1,1,0,2,2,1,2,1,2,1,2,1,1,1,1,1,1,1,1,2,3,2,1,1,2,1,1,0,1,3,1,2,1,1,3,1,2,2,1,2,2,2,1,2,1,3,2,0,1,2,1,1,1,1,2,2,3,1,2,2,2,2,1,2,2,1,1,2,1,1,2,1,2,1,0,1,1,2,2,0,1,2,2,1,1,1,2,2,1,2,1,1,1,1,1,2,1,1,2,2,1,1,2,2,1,0,2,0,1,1,1,2,2,2,2,1,1,0,1,1,1,0,1,2,2,2,1,3,2,1,1,1,2,2,1,2,2,1,1,2,1,2,1,2,2,2,1,1,2,2,2,1,1,2,1,2,1,1,1,1,2,0,1,2,2,1,1,0,1,3,1,1,1,1,2,1,2,2,2,1,2,1,1,1,1,3,1,1,1,2,2,0,2,1,2,1,3,0,1,1,1,1,2,0,2,2,2,2,2,1,1,1,0,0,1,2,2,2,1,1,1,1,0,0,1,1,1,0,1,1,0,1,1,1,1,1,1,1,3,2,3,0,1,1,1,1,0,1,1,2,2,1,1,1,1,1,1,0,0,1,1,1,1,1,1,1,2,0,2,1,1,1,1,1,2,3,1,2,1,1,2,0,2,2,1,1,1,1,1,1,2,1,1,2,0,1,1,2,1,2,1,1,1,2,0,1,1,0,1,1,0,0,1,0,2,2,2,1,2,0,1,1,1,1,1,1,1,1,1,1,2,2,2,2,1,1,1,2,2,0,1,1,1,1,0,1,1,2,1,1,2,2,1,2,1,1,2,2,1,1,2,1,1,1,0,1,1,1,1,0,1,0,1,1,1,2,0,1,1,1,1,1,2,1,1,1,2,1,2,2,1,0,1,1,0,1,1,2,1,1,1,1,0,1,1,0,1,1,1,1,1,1,1,1,2,1,0,1,0,1,1,2,1,1,1,1,1,1,1,1,1,1,1,1,1,1,1,0,1,1,1,1,1,0,1,0,0,1,1,1,1,2,1,1,2,2,1,2,0,1,0,1,1,0,0,1,0,1,0,1,1,1,1,1,1,1,1,1,1,1,2,1,1,1,0,1,1,1,1,1,1,0,1,1,1,1,1,0,0,1,1,1,0,1,1,1,0,0,1,2,0,2,1,1,2,1,2,2,1,0,1,1,1,1,0,1,2,1,0,2,0,0,1,2,1,1,0,2,1,0,0,0,0,1,1,1,1,1,1,0,0,2,1,1,0,1,1,1,1,1,0,0,0,2,1,1,0,1,1,0,1,1,1,1,0,1,1,1,1,0,1,1,1,2,1,1,1,1,0,0,1,1,1,1,1,1,0,1,1,1,1,2,1,0,0,0,1,1,1,1,0,1,0,1,1,0,1,0,1,0,0,0,1,1,2,0,1,1,0,1,1,1,0,1,1,1,1,1,2,1,1,1,0,1,0,1,1,2,0,1,0,1,0,2,1,1,1,1,0,2,1,0,1,1,1,1,1,1,1,0,1,0,1,1,1,0,1,1,2,1,0,1,1,1,1,1,0,1,1,2,1,1,1,1,1,1,0,0,1,0,1,1,1,1,1,0,1,1,1,1,0,1,1,1,0,0,1,0,0,0,1,0,0,1,1,1,0,0,1,1,0,1,1,1,1,1,1,1,0,1,1,1,0,2,0,1,1,1,1,1,1,1,0,1,0,0,0,0,1,0,1,1,0,0,0,0,0,0,0,1,1,1,0,0,0,0,1,1,1,0,0,1,0,1,1,1,1,0,0,1,0,2,0,0,1,1,0,1,0,1,1,1,1,1,1,0,0,0,1,1,1,0,0,1,1,1,1,2,0,1,0,2,0,1,1,0,0,0,1,1,0,0,1,0,1,0,0,0,0,1,1,0,0,0,0,0,0,1,0,0,0,0,0,0,1,1,0,0,1,1,0,0,1,1,0,1,0,0,0,0,0,0,1,1,1,0,0,0,1,1,1,1,0,0,1,0,1,1,1,1,0,1,1,1,1,1,0,0,1,0,0,0,1,0,0,1,0,0,1,1,0,0,0,0,1,0,1,1,0,0,0,0,0,0,1,1,1,1,1,1,1,0,0,0,1,0,0,0,1,0,1,0,0,0,0,0,0,0,0,0,0,0,0,0,0,0,0,1,0,1,0,1,1,1,0,1,1,1,1,0,0,0,0,0,0,0,1,0,0,0,0,0,0,1,0,0,1,1,0,1,0,0,0,1,0,0,1,0,0,0,0,1,0,1,1,0,1,1,0,0,1,0,0,0,0,0,1,1,1,0,0,0,0,0,1,1,0,0,1,1,1,0,1,0,1,0,0,1,0,1,0,0,0,1,0,0,1,0,0,0,0,0,1,0,1,1,0,0,1,0,1,0,0,0,0,1,0,0,0,0,0,0,0,1,0,0,1,1,0,0,0,0,0,0,0,0,0,0,0,0,0,0,0,0,0,1,0,0,0,0,1,0,0,1,1,1,0,0,1,0,0,0,0,0,0,2,0,0,1,0,1,0,1,1,0,0,0,0,0,0,0,0,0,1,0,0,0,0,0,0,0,0,0,0,0,1,0,1,0,1,0,0,1,0,0,0,1,1,1,0,1,0,0,0,1,0,0,1,0,0,0,1,0,1,1,0,0,1,1,0,0,0,0,1,0,0,0,1,1,0,1,1,0,1,0,0,1,0,0,0,0,0,1,0,0,0,0,0,0,0,0,1,0,0,1,0,0,1,0,0,0,0,1,0,0,1,0,0,0,1,0,0,0,0,1,0,0,1,0,0,0,1,0,1,0,1,1,0,1,0,1,0,0,0,0,0,0,0,0,0,0,0,0,0,1,0,1,0,0,0,0,0,1,0,0,1,0,0,0,0,0,0,0,1,0,0,0,0,0,0,0,0,1,0,0,0,1,0,0,0,0,0,0,0,0,0,0,0,1,0,0,0,1,1,0,0,0,0,1,0,1,0,0,0,0,0,0,0,1,0,1,0,0,0,0,0,0,1,0,0,0,1,0,1,1,0,0,0,1,0,0,0,0,1,0,0,0,0,0,0,0,0,0,0,0,0,0,0,0,0,0,1,0,0,0,0,0,0,0,0,0,1,0,0,1,0,0,1,0,0,0,0,0,0,0,0,0,0,0,0,0,0,1,0,0,0,0,0,0,0,0,0,0,0,0,0,0,0,0,0,0,1,0,0,0,0,0,0,0,0,0,0,0,0,0,0,0,0,0,0,0,0,0,0,0,1,0,0,0,0,0,0,0,0,0,0,1,0,0,0,1,0,0,1,0,0,0,0,0,1,0,0,1,0,0,0,0,0,1,0,0,0,0,0,0,0,1,0,0,0,0,0,1,0,0,0,0,0,0,0,0,1,1,0,1,0,1,0,0,0,0,0,0,0,0,1,0,0,0,0,0,0,0,1,1,0,0,1,0,0,0,0,0,0,0,0,0,0,0,0,0,0,1,0,1,0,0,0,0,0,0,0,0,0,0,0,1,0,1,0,0,0,0,0,0,0,0,0,0,0,0,0,0,0,0,0,0,0,1,0,0,0,0,1,0,0,0,0,0,0,1,0,0,1,0,0,0,0,0,0,0,0,0,0,0,0,1,0,0,0,1,0,0,0,0,0,1,0,1,0,0,0,0,0,0,0,0,0,0,1,0,0,0,0,0,0,0,0,0,1,0,0,0,0,0,0,0,0,1,0,0,0,0,0,0,0,0,0,0,1,0,0,0,0,0,0,0,0,0,0,0,0,0,0,0,0,0,0,0,0,1,0,0,0,0,0,0,0,0,0,0,0,1,0,1,0,0,0,0,1,0,0,0,0,0,0,0,0,1,0,0,0,0,0,0,0,0,0,0,0,0,0,0,0,1,0,0,0,0,0,0,0,0,0,0,0,0,0,0,0,0,0,0,0,0,0,0,0,0,0,0,0,0,0,0,0,1,0,0,0,0,1,0,0,0,0,0,0,0,0,0,0,0,0,0,0,0,0,0,0,0,0,0,1,0,0,0,0,0,0,0,0,0,0,0,0,0,0,0,0,0,0,0,0,0,0,0,0,0,0,0,0,0,1,0,1,0,0,0,0,0,0,0,0,0,0,0,0,0,0,1,0,0,0,0,0,0,0,0,0,0,0,0,0,0,0,0,0,0,1,0,0,1,0,0,1,0,0,0,0,0,0,0,0,0,0,0,0,0,0,0,0,0,0,0,0,0,0,0,0,0,0,1,0,0,0,0,0,0,0,0,0,0,0,0,0,0,0,0,0,0,0,0,0,0,0,0,0,0,0,0,0,0,0,0,0,0,0,0,0,0,0,0,0,0,0,0,0,0,0,0,0,0,0,0,1,1,0,0,0,0,0,0,0,0,0,0,0,0,0,0,0,0,0,0,0,0,1,0,0,1,0,0,0,0,0,0,0,0,0,0,0,0,0,0,0,0,0,0,1,0,0,0,0,0,0,0,0,0,0,0,0,0,0,0,0,0,0,1,0,0,0,0,0,0,0,0,0,0,0,0,0,0,0,0,0,0,0,0,0,0,0,0,0,0,0,0,0,0,0,0,0,0,0,0,0,0,0,0,0,0,0,0,0,0,0,0,0,0,0,0,0,0,0,0,0,0,0,1,0,0,0,0,0,0,0,0,0,0,0,0,0,0,0,0,0,0,0,0,0,0,0,0,0,0,0,0,0,0,0,0,0,0,0,0,0,0,0,0,0,0,0,0,0,0,0,0,0,0,0,0,0,1,0,0,0,0,0,0,0,0,0,0,0,0,0,0,0,0,0,0,0,0,0,0,0,0,0,0,0,0,0,0,0,0,0,0,0,0,0,0,0,0,0,0,0,0,0,0,0,0,0,0,0,0,0,0,0,0,0,0,0,0,0,0,0,0,0,0,0,0,0,0,0,0,0,0,0,0,0,0,0,0,0,0,0,0,0,0,0,0,0,0,0,0,0,0,0,0,0,0,0,0,0,0,0,0,0,0,1,0,0,0,0,0,0,0,0,0,0,0,0,0,0,0,0,0,0,0,0,0,0,0,0,0,0,0,0,0,0,0,0,0,0,0,0,0,0,0,0,0,0,0,0,0,0,0,0,0,0,0,0,0,0,0,0,0,0,0,0,0,0,0,0,0,0,0,0,0,0,0,0,0,0,0,0,0,0,0,0,0,0,0,0,0,0,0,0,0,0,0,0,0,0,0,0,0,0,0,0,0,0,0,0,0,0,0,0,0,0,0,0,0,0,0,0,0,0,0,0,0,0,0,0,0,0,0,0,0,0,0,0,0,0,0,0,0,0,0,0,0,0,0,0,0,0,0,0,0,0,0,0,0,0,0,0,0,0,0,0,0,0,0,0,0,0,0,0,0,0,0,0,0,0,0,0,0,0,0,0,0,0,0,0,0,0,0,0,0,0,0,0,0,0,0,0,0,0,0,0,0,0,0,0,0,0,0,0,0,0,0,0,0,0,0,0,0,0,0,0,0,0,0,0,0,0,0,0,0,11,11,10,10,10,9,7,7,6,4,4,4,4,4,3,3,3,3,3,2,2,2,2,2,2,2,2,2,2,2,2,2,2,2,2,1,1,1,1,1,1,1,1,1,1,1,1,1,1,1,1,1,1,1,1,1,1,1,1,1,1,1,1,1,1,1,0,0,0,0,0,0,0,0,0,0,0,0,0,0,0,0,0,0,0,0,0,0,0,0,0,0,0,0,0,0,0,0,0,0,0,0,0,0,0,0,0,0,0,0,0,0,0,0,0,0,0,0,0,0,0,0,0)
VarExplained_mRNA_Age=c(7,0,2,2,1,2,10,7,10,0,1,0,4,5,0,0,0,2,7,9,3,6,12,2,1,2,0,6,0,0,1,4,2,0,0,0,2,4,2,0,3,2,1,3,0,4,6,6,0,1,10,1,14,3,6,5,2,1,0,0,3,1,0,3,3,0,4,0,0,6,4,0,2,0,1,2,1,3,5,0,0,0,4,2,0,4,0,0,0,3,1,1,2,1,3,0,1,0,5,0,3,3,3,1,2,1,1,0,1,7,4,13,10,2,1,0,3,1,3,0,3,0,0,5,4,7,4,11,2,0,4,0,5,0,8,0,0,5,12,0,0,1,0,0,0,1,0,2,0,0,6,5,4,1,1,1,0,5,3,4,3,1,2,0,7,2,0,1,6,0,0,0,4,6,2,2,0,0,0,1,10,2,5,0,3,6,0,1,11,4,4,7,1,0,1,2,0,2,2,2,0,1,0,3,0,1,3,0,4,2,0,0,3,3,7,10,2,2,1,5,0,0,4,2,7,1,2,0,0,0,6,0,0,4,0,0,2,1,0,7,3,2,1,2,1,3,1,2,2,4,2,3,0,2,5,2,1,1,0,4,12,9,0,15,6,7,3,4,0,1,7,7,1,2,1,2,2,0,4,3,7,0,2,2,2,0,0,2,5,1,1,2,6,6,1,2,13,2,0,5,0,4,0,1,2,1,4,6,5,0,1,7,6,0,0,1,1,1,0,4,1,1,0,3,1,0,6,0,5,6,4,2,1,5,1,0,4,0,1,2,2,2,6,0,2,1,0,2,3,2,4,3,4,3,3,1,2,8,2,2,3,1,4,5,0,1,1,4,2,0,10,2,0,3,4,1,1,0,1,6,0,0,2,4,3,0,0,0,0,10,4,4,5,1,0,2,0,6,5,0,1,0,11,0,4,0,2,0,3,0,2,2,4,5,1,7,0,2,0,2,0,1,2,8,0,0,3,1,0,1,2,2,2,1,5,0,0,3,0,0,8,0,0,4,2,2,1,5,10,2,0,4,0,1,5,1,4,2,0,1,1,0,1,3,2,2,1,0,1,0,1,5,1,3,1,2,0,0,0,1,0,7,1,4,2,1,5,6,0,0,0,1,4,1,0,0,0,2,2,1,4,5,4,2,1,8,0,1,3,2,0,1,0,5,0,2,1,3,8,24,7,1,1,3,2,0,0,3,7,0,0,0,0,0,0,1,3,13,0,0,0,1,1,2,8,1,1,1,4,0,0,5,0,1,2,3,0,0,4,1,0,3,2,0,3,1,1,0,5,4,1,0,2,2,1,2,0,1,2,3,1,1,3,0,7,1,0,0,1,6,0,0,3,0,7,0,1,1,0,4,2,2,1,1,0,5,3,1,0,1,3,1,13,1,0,1,4,1,2,4,3,0,4,0,0,4,4,5,0,1,8,1,6,1,1,2,15,1,0,4,0,3,7,0,9,1,0,0,5,5,2,3,3,4,1,1,0,11,0,1,0,1,0,11,2,4,2,1,10,0,9,1,2,6,0,1,3,5,1,4,0,0,0,2,0,0,1,2,0,13,1,0,3,1,1,1,5,1,1,0,2,0,2,1,1,3,3,3,4,2,0,1,2,5,3,0,0,1,0,0,0,13,2,2,3,1,3,1,1,1,0,7,1,1,1,4,3,0,4,1,3,0,3,2,5,0,0,1,2,8,3,2,5,11,3,7,2,2,1,7,1,3,0,0,2,1,0,0,3,4,2,5,0,0,0,24,2,1,0,0,2,0,1,2,14,1,2,0,2,7,0,3,0,0,0,2,2,14,0,4,1,0,0,0,10,6,0,0,13,2,14,0,1,1,5,1,1,0,3,0,2,0,0,1,0,7,2,0,0,7,2,2,3,0,12,7,0,2,11,7,2,0,0,0,3,4,1,17,0,1,1,1,0,1,0,0,0,0,0,4,7,8,0,2,0,3,0,0,1,7,1,1,1,1,2,3,2,7,1,2,3,11,11,1,0,1,1,5,1,0,1,0,5,6,7,3,0,0,1,5,0,1,0,0,1,0,6,0,6,9,7,6,1,1,9,7,7,1,1,0,0,1,0,0,3,5,3,2,0,9,0,0,8,9,0,1,4,5,1,2,1,0,0,0,0,2,0,1,0,0,1,2,11,8,0,1,0,1,0,1,2,0,0,0,1,3,0,2,2,1,6,1,0,0,0,0,2,0,1,0,0,7,0,2,0,4,1,1,6,2,3,0,0,3,1,0,1,2,2,0,6,5,2,6,3,1,4,3,0,3,1,0,2,2,2,2,2,0,1,0,0,1,1,1,4,0,0,1,0,3,0,2,0,8,0,1,0,3,0,1,12,1,0,1,0,3,1,8,1,0,1,0,8,1,0,2,3,0,0,2,3,1,0,2,1,0,2,0,1,1,0,3,7,1,2,0,3,3,2,4,1,0,0,0,1,3,2,0,1,1,0,0,0,0,0,0,1,1,0,1,2,0,0,0,0,6,0,2,0,7,0,0,1,1,12,0,1,2,0,0,3,4,8,1,0,0,6,3,1,0,5,2,1,1,1,1,1,4,0,1,2,0,7,2,0,6,3,6,4,17,1,1,8,8,4,5,3,3,3,3,0,0,0,0,0,1,2,5,5,0,5,1,1,0,2,0,0,4,10,3,7,0,0,11,2,8,0,1,0,6,6,1,0,14,1,2,0,0,1,2,2,1,0,1,5,2,12,4,0,6,10,0,0,0,0,0,1,1,1,0,10,4,3,0,0,0,0,5,12,1,3,0,14,11,7,0,1,0,1,7,0,0,1,3,1,0,0,3,0,1,0,1,0,12,0,2,1,0,2,3,9,3,0,11,1,5,2,0,1,1,1,0,0,0,1,2,5,0,8,0,2,0,2,0,1,7,4,1,3,1,0,2,0,3,2,0,2,0,6,0,3,1,4,4,0,0,2,0,0,2,0,3,1,1,6,8,1,7,6,1,2,1,0,10,1,1,5,5,2,0,0,1,2,1,0,2,2,0,3,10,0,10,0,3,3,0,0,2,0,9,1,8,3,0,2,2,0,4,1,3,2,2,2,1,0,0,1,0,2,3,1,0,5,0,11,1,0,6,8,0,5,2,1,2,0,1,0,0,0,11,0,11,16,1,3,0,0,2,4,1,0,5,3,1,1,8,0,10,2,1,0,3,0,0,0,5,3,5,6,0,0,2,2,0,0,0,11,2,0,2,0,22,0,6,2,7,0,1,3,11,3,3,1,0,3,3,0,1,1,6,10,1,0,0,0,0,0,0,0,2,0,3,1,3,7,1,8,1,3,2,6,4,1,2,0,3,0,6,1,0,1,6,5,5,0,0,0,2,0,0,0,6,2,4,1,1,1,0,1,1,1,0,1,0,0,0,0,2,0,0,1,0,3,1,1,1,10,4,0,5,2,2,5,12,0,0,2,2,2,12,0,0,2,0,5,0,2,1,6,2,0,2,1,0,3,0,6,7,2,0,7,2,11,0,4,7,0,3,8,0,5,0,6,5,0,1,7,16,4,0,0,2,1,0,4,0,3,2,0,2,3,0,0,1,7,0,0,0,2,11,0,5,0,0,0,1,0,0,1,2,2,0,14,2,4,1,0,1,2,0,4,4,1,3,5,5,0,3,0,0,2,1,3,6,1,2,1,8,0,3,0,0,0,0,1,9,1,0,1,1,2,2,14,1,0,6,0,1,1,0,2,0,1,1,1,6,0,1,3,5,1,2,1,0,1,0,1,0,3,2,4,1,0,0,4,3,1,1,9,9,4,1,0,0,3,0,0,0,1,0,7,3,14,1,6,3,10,0,1,0,1,0,0,0,0,5,0,0,2,1,0,1,3,4,0,0,6,0,1,1,0,2,2,0,1,1,7,2,1,0,0,1,1,0,0,12,1,11,0,5,2,5,0,0,2,0,0,3,3,0,1,5,2,1,1,0,2,1,2,0,0,0,0,0,0,5,5,4,0,9,0,0,0,1,1,9,7,0,0,1,11,5,5,2,2,0,2,1,4,4,10,0,0,0,1,0,0,0,3,0,0,0,0,0,0,0,2,0,0,0,0,0,5,4,0,1,5,0,2,0,0,3,0,3,1,2,0,0,5,0,1,0,0,1,0,2,4,0,0,1,0,3,0,0,0,0,4,13,13,2,1,7,1,0,11,0,0,2,4,0,0,9,1,4,0,0,1,0,2,6,1,0,1,0,12,0,2,2,3,1,0,0,0,1,0,0,0,0,1,2,0,0,1,4,1,5,4,5,0,2,2,1,0,1,1,2,1,5,1,3,5,0,1,5,4,8,0,1,1,1,2,1,0,2,4,1,0,4,0,1,1,5,2,0,1,0,0,0,0,0,2,0,0,3,1,0,0,0,0,4,0,0,0,3,6,1,1,1,0,1,0,0,1,0,2,2,0,0,1,8,3,4,1,3,3,2,0,1,1,1,0,0,0,0,2,0,1,0,1,2,1,4,0,0,1,2,1,1,0,4,0,0,0,0,2,0,0,0,0,1,6,0,14,4,0,0,0,0,1,2,4,0,5,1,0,0,1,1,5,1,11,0,4,2,2,6,1,6,1,0,1,1,0,0,1,0,2,1,1,1,0,3,2,1,0,0,2,0,4,0,1,0,0,0,1,0,1,3,5,3,0,0,1,2,1,0,1,0,0,0,0,1,0,1,0,2,15,0,1,4,1,0,1,0,0,1,0,1,1,1,1,1,0,1,0,4,1,2,6,8,0,2,0,3,1,0,0,0,1,0,2,0,0,1,4,4,1,6,0,0,0,5,0,2,0,2,0,5,0,1,0,5,2,0,0,0,0,1,0,0,0,2,0,2,1,0,5,10,1,0,3,0,0,3,1,0,9,0,0,0,3,1,0,1,2,0,0,6,1,2,0,3,1,4,0,1,0,1,6,1,0,3,0,0,2,0,1,2,0,2,1,1,1,2,5,0,6,1,0,0,0,0,0,1,3,0,1,1,4,1,1,0,0,10,1,0,1,2,12,2,5,12,6,0,1,0,2,5,0,1,0,0,2,0,2,0,0,0,5,0,0,0,1,2,1,3,0,0,1,1,0,0,0,0,13,0,0,0,0,2,1,3,5,0,1,1,1,0,0,1,0,0,0,11,2,1,0,1,1,0,0,2,0,0,0,4,1,0,3,5,0,0,1,1,0,3,1,0,2,0,0,0,1,3,1,5,0,1,2,0,0,0,1,1,4,0,1,1,0,0,0,3,0,1,1,0,1,0,0,2,14,0,1,5,4,1,1,1,1,1,6,3,1,0,0,1,0,7,0,3,2,0,0,0,1,2,3,0,0,8,1,2,4,2,2,0,2,6,2,0,1,0,1,0,0,9,5,2,1,0,4,2,1,9,1,9,8,0,2,0,0,1,6,0,0,1,2,0,1,4,0,0,6,0,9,1,1,2,3,8,0,0,5,0,1,3,1,6,0,1,1,3,4,0,0,6,0,1,3,0,5,0,1,0,1,0,1,2,0,2,4,9,0,0,2,0,0,0,0,3,2,0,1,1,7,2,0,0,3,0,0,0,0,0,6,6,1,0,0,2,0,0,1,1,2,1,0,5,6,0,1,1,0,0,0,0,3,0,1,0,5,0,1,0,1,13,3,6,1,3,0,18,0,0,2,3,0,2,10,4,0,3,2,1,1,0,1,5,3,0,1,0,2,1,1,2,0,1,0,0,1,1,8,0,1,0,7,6,1,5,4,2,0,0,0,2,1,0,2,1,7,1,1,0,0,1,2,0,0,2,0,0,6,3,0,1,6,1,0,3,1,1,2,0,0,0,1,6,0,4,9,3,0,0,1,2,3,1,0,0,1,5,1,0,0,0,0,0,0,0,0,0,0,2,0,1,6,4,2,1,4,1,0,0,0,0,15,5,0,2,9,1,0,0,4,1,0,0,1,0,6,1,1,0,1,0,0,0,7,0,3,1,0,5,0,0,2,9,0,2,13,1,0,1,0,2,6,1,0,0,0,0,0,0,0,0,3,4,0,1,3,0,1,0,1,1,0,0,1,0,5,0,0,4,0,0,0,2,1,2,3,0,1,3,0,0,1,3,0,1,0,0,1,6,5,3,4,0,0,2,0,0,2,0,1,0,5,0,0,9,0,0,0,0,1,0,0,1,1,3,1,0,0,0,3,4,2,2,0,2,0,0,0,0,0,3,2,0,0,7,11,1,6,8,1,2,1,0,5,0,0,2,2,2,0,1,3,0,1,1,0,4,0,2,3,0,1,6,0,0,0,1,6,1,6,1,0,2,1,0,2,0,3,2,0,2,2,0,14,6,8,0,15,1,1,2,0,0,1,7,2,4,0,6,1,3,4,1,1,1,0,19,0,0,3,0,1,0,0,0,2,1,3,1,1,0,7,0,1,1,0,0,10,7,4,0,1,1,2,6,0,3,0,2,0,2,1,4,5,0,0,1,0,0,0,0,7,9,0,0,11,1,7,7,2,11,1,0,0,1,2,1,4,4,0,1,0,1,1,0,0,0,3,0,6,1,1,1,8,0,1,0,12,0,2,1,1,0,2,2,0,1,0,0,0,4,2,2,5,0,0,1,0,4,0,1,0,2,0,0,0,0,6,0,0,2,0,0,0,5,1,3,0,0,11,0,0,0,1,12,2,1,1,0,0,0,1,0,0,2,0,6,0,0,0,3,3,4,0,0,1,1,0,0,0,1,1,1,3,0,2,2,2,0,0,1,0,10,2,0,0,2,1,1,0,7,1,13,1,11,3,1,0,1,6,0,16,3,3,0,0,0,3,0,2,1,7,0,0,2,5,1,0,3,5,0,0,0,1,4,0,0,0,0,0,2,0,12,0,0,1,0,0,1,0,5,3,0,0,7,0,8,3,1,0,2,0,0,0,7,0,4,0,2,0,3,1,11,0,0,0,3,3,3,0,0,0,0,1,1,0,0,0,0,0,0,0,0,0,0,2,4,0,1,0,1,0,3,1,1,0,1,2,13,1,15,5,0,0,1,0,6,13,0,0,0,0,0,0,0,3,0,2,0,1,0,1,1,0,0,0,0,0,0,3,7,1,0,0,0,0,1,0,4,0,0,2,2,0,1,0,0,1,1,0,1,4,0,5,6,2,0,1,1,0,1,0,2,5,0,12,0,0,0,4,0,0,2,4,1,4,0,1,5,4,3,15,1,0,2,1,0,2,0,0,0,0,0,0,0,16,2,5,0,3,3,0,9,0,0,1,1,7,1,0,0,2,3,0,1,0,0,0,0,0,1,2,1,4,11,1,0,1,1,0,2,6,1,0,0,0,2,4,0,4,0,0,1,0,2,0,0,0,0,1,1,1,2,0,0,0,4,7,0,2,9,5,1,0,0,0,2,1,0,0,0,1,2,3,2,7,0,7,3,3,0,0,6,0,0,1,4,7,0,0,3,3,0,0,0,7,0,1,0,1,4,1,0,0,5,2,0,2,1,0,3,0,0,1,0,0,1,0,1,1,8,3,1,10,1,4,0,1,2,1,5,1,6,0,0,0,0,1,1,0,0,0,2,0,3,3,0,1,1,2,8,1,0,1,1,0,1,1,0,0,0,2,1,7,0,0,0,0,7,0,1,0,2,0,0,1,5,0,0,1,1,3,1,0,2,4,2,5,1,0,2,1,0,1,6,7,0,0,0,0,3,1,1,3,1,0,0,1,0,5,0,1,1,0,0,1,0,0,0,1,0,0,0,0,2,2,1,1,0,1,7,0,5,1,0,1,0,0,0,0,1,0,0,1,0,0,0,1,1,0,8,1,1,0,1,4,0,0,0,0,4,2,0,0,1,0,0,0,0,1,1,0,8,0,8,0,0,1,2,1,1,0,1,2,0,0,10,0,1,2,0,0,0,0,0,1,0,8,1,0,0,0,1,0,0,0,1,0,2,3,1,1,0,0,10,1,1,0,0,1,3,0,1,2,0,2,4,1,2,1,0,9,1,4,0,8,4,4,0,4,0,1,6,3,3,0,2,0,14,0,1,16,1,1,1,1,9,1,0,0,3,1,0,0,2,2,0,2,0,8,4,0,0,0,0,0,4,0,0,0,1,4,0,3,1,2,0,1,2,2,1,0,0,2,0,3,1,13,1,2,0,1,0,1,0,3,2,1,0,1,1,0,1,0,5,0,1,2,7,1,0,4,3,1,0,0,1,4,0,7,0,1,0,1,0,6,3,0,0,0,1,0,6,0,1,4,11,1,6,6,0,7,0,0,2,1,3,0,3,0,0,2,0,3,0,0,1,0,1,0,2,2,0,1,1,0,1,14,0,0,1,0,0,1,2,1,0,1,0,0,0,1,4,0,0,1,6,1,12,2,0,0,1,0,0,3,1,2,0,0,1,16,0,1,0,0,0,1,9,0,0,0,0,0,2,0,0,0,0,0,2,0,5,1,0,9,0,0,1,1,5,1,0,0,0,0,0,3,1,3,1,0,2,1,1,1,1)
VarExplained_mRNA_Interaction=c(19,29,22,23,22,22,23,31,21,29,27,31,26,27,33,27,28,26,24,29,18,25,29,22,27,33,31,26,27,25,31,31,26,38,32,18,22,29,31,32,27,31,28,32,26,32,32,29,27,21,22,25,30,26,28,32,21,28,29,34,34,35,31,31,35,30,38,34,32,29,35,29,33,30,36,31,30,30,36,32,28,33,30,30,29,28,39,30,32,32,31,25,33,32,29,36,27,33,33,43,29,30,33,34,28,35,33,36,30,35,31,30,30,33,28,37,31,29,28,36,34,31,27,37,33,32,30,34,29,32,32,30,40,36,27,27,37,32,39,32,30,34,34,33,32,34,30,32,37,32,25,37,29,34,26,33,34,30,29,32,26,25,34,34,35,38,33,35,33,38,33,32,33,33,32,35,35,32,34,36,24,34,30,30,31,38,38,34,26,32,31,28,39,27,34,29,40,33,33,27,33,30,32,30,21,24,36,38,31,35,31,31,38,23,25,35,32,33,38,22,36,32,40,33,35,38,30,31,37,29,31,34,34,36,36,24,36,35,30,30,23,31,30,38,38,30,35,30,33,36,20,33,34,23,30,33,19,33,32,33,34,34,32,29,30,32,28,29,33,32,34,30,25,28,34,33,38,31,34,31,30,33,36,31,31,33,37,32,26,35,33,30,33,30,36,29,18,34,38,32,31,31,29,36,33,35,35,31,38,31,41,29,30,36,37,28,38,30,30,35,37,30,32,31,30,28,25,37,30,31,31,36,34,33,31,46,31,38,11,36,35,35,23,32,24,34,36,37,31,31,35,38,34,24,33,41,39,40,36,35,37,27,29,28,20,34,38,35,32,37,25,32,35,21,40,37,34,35,33,35,39,44,38,29,29,32,33,36,25,35,36,31,30,31,33,37,29,32,27,35,35,31,27,34,27,40,33,36,36,25,33,36,34,29,36,31,27,37,32,32,35,25,32,34,30,22,27,34,34,35,31,38,37,15,32,33,38,33,39,38,24,39,39,31,33,36,42,33,33,38,36,35,33,33,28,23,28,35,40,22,35,34,33,31,38,36,36,28,32,32,39,36,26,36,44,34,32,35,34,34,26,30,31,36,34,28,32,35,35,37,32,31,33,34,40,34,39,37,41,36,36,31,25,35,30,36,41,33,37,33,33,38,35,30,32,40,39,31,33,26,37,36,34,29,32,35,36,41,29,28,29,35,32,37,40,34,37,31,31,36,35,36,20,34,39,38,33,31,28,33,32,33,35,38,38,41,20,35,33,37,29,35,31,37,29,32,31,31,36,35,20,34,34,34,35,33,35,37,33,36,33,38,36,39,29,39,35,30,40,31,15,37,35,33,32,29,32,35,37,35,32,33,35,39,37,40,35,38,31,29,36,33,28,30,30,33,33,31,24,32,30,42,33,42,36,30,34,37,30,30,27,33,33,35,32,32,28,40,37,35,35,37,32,30,30,30,36,37,31,35,28,36,35,25,41,38,34,36,30,40,37,40,35,28,33,42,36,33,32,36,19,30,34,25,34,35,37,40,40,40,34,40,37,40,38,38,40,24,35,28,40,34,35,32,32,45,33,33,36,32,36,21,36,27,39,29,32,35,22,31,32,39,32,34,36,39,32,36,35,31,33,28,29,40,39,34,39,33,28,30,35,33,29,36,32,35,26,33,35,34,39,35,19,27,37,33,38,35,30,29,28,35,34,33,30,37,31,32,41,30,36,35,32,31,29,36,38,34,30,37,40,30,25,44,33,28,36,33,34,39,32,38,42,35,33,32,34,27,36,34,37,36,33,40,35,33,30,36,37,37,31,27,38,31,30,33,35,36,34,22,33,35,40,38,38,35,34,34,33,35,33,33,41,36,17,37,30,40,36,29,33,40,37,41,28,32,35,33,42,30,29,32,31,33,33,36,38,27,31,27,38,33,37,33,37,32,33,34,35,31,38,33,40,33,36,42,30,35,36,28,37,30,38,27,38,31,33,30,26,36,34,25,35,35,41,38,35,34,35,45,36,41,37,31,34,35,30,41,20,34,39,28,41,38,36,35,37,41,31,40,29,30,36,38,32,38,35,31,34,35,39,37,35,34,36,34,27,36,35,42,37,40,29,36,37,31,36,28,26,36,29,25,33,31,30,19,38,32,33,41,31,32,32,44,35,35,36,38,33,32,28,18,35,28,33,25,36,36,37,33,40,33,34,42,36,28,41,34,21,25,30,35,33,37,37,39,38,35,36,32,33,35,35,37,27,32,38,13,33,35,31,28,32,34,37,29,36,40,37,34,37,27,33,32,34,32,34,32,33,35,36,32,37,36,38,34,39,29,37,35,36,37,37,33,36,40,35,33,39,25,24,34,35,15,31,42,36,28,36,34,37,35,37,36,30,42,30,39,34,34,27,35,30,33,32,31,32,37,30,34,21,39,40,32,40,33,39,40,35,36,38,36,33,23,39,37,40,37,34,43,19,28,33,38,37,35,39,35,37,35,32,35,29,30,36,18,29,35,37,30,31,36,33,19,32,34,27,36,33,39,42,31,35,43,33,36,40,38,39,34,34,37,32,37,34,32,37,31,30,25,32,36,33,34,36,31,35,35,29,36,26,28,35,25,38,33,37,26,25,32,27,40,34,33,34,39,37,46,37,32,39,35,35,33,36,33,35,39,32,42,29,39,34,39,32,33,40,36,34,31,34,37,37,30,43,34,32,33,31,26,35,40,41,37,38,38,29,37,35,39,44,33,35,25,40,37,31,31,40,45,32,41,36,38,40,29,39,34,36,25,44,38,26,35,34,35,37,24,36,32,37,33,38,37,33,36,38,35,38,33,37,36,36,32,34,38,36,30,32,29,28,33,34,27,30,31,39,39,18,34,38,35,27,37,34,31,29,40,36,42,41,35,30,39,43,33,31,34,37,38,36,46,30,41,33,32,40,17,33,35,34,36,38,28,31,41,38,24,31,36,33,40,36,34,30,27,31,46,32,37,33,33,35,30,36,41,29,36,38,38,44,28,37,33,34,35,32,38,34,34,36,36,35,39,37,37,37,40,29,33,34,33,33,32,37,39,36,42,33,36,39,41,27,27,32,33,35,32,29,40,36,33,35,41,31,30,43,38,30,39,33,28,34,41,35,37,34,38,40,27,34,38,32,33,32,33,35,18,34,35,19,30,30,39,35,31,23,31,26,30,33,39,38,32,39,35,26,33,34,43,36,33,33,42,34,35,32,32,35,31,31,20,37,32,39,29,36,34,38,30,37,41,26,36,30,36,42,32,31,30,35,35,41,39,32,33,34,35,11,37,39,32,33,36,28,34,37,37,41,37,31,33,31,37,38,35,32,36,32,35,37,29,32,34,35,32,34,33,37,39,33,33,41,36,38,31,37,36,37,37,38,33,37,35,31,38,33,37,36,33,23,36,35,40,38,35,32,31,34,34,34,32,39,28,34,30,40,34,34,31,34,35,33,33,33,28,33,34,38,44,32,32,34,32,31,28,33,29,26,37,31,18,31,22,39,35,33,37,34,21,30,35,35,31,37,37,33,29,29,36,22,40,36,37,31,36,33,14,37,33,33,38,35,31,32,38,34,40,21,29,15,24,33,27,41,36,44,35,42,33,36,43,35,38,41,35,43,41,25,38,35,35,24,41,40,30,37,31,7,40,39,37,41,39,27,38,30,30,38,45,32,30,36,33,36,33,37,39,34,32,35,39,33,32,46,34,35,42,13,27,39,32,40,35,39,34,35,38,35,36,40,31,38,37,31,39,39,29,34,38,37,29,37,35,32,41,32,29,37,32,38,35,36,37,34,30,31,37,45,27,28,42,30,36,35,36,25,7,30,36,43,30,35,37,35,33,35,28,31,30,10,31,33,35,40,24,31,34,34,28,36,38,38,37,34,39,34,33,41,41,36,33,20,37,38,31,38,27,31,37,35,28,34,39,40,35,38,35,35,36,24,31,28,33,29,37,24,36,36,38,40,33,39,37,32,36,38,39,37,37,28,37,37,40,32,35,35,36,38,29,33,38,35,39,35,34,33,32,29,34,33,37,25,31,36,38,35,34,39,42,37,36,38,34,35,40,33,42,37,29,33,36,28,37,33,35,43,39,35,35,46,38,39,34,42,39,41,38,40,32,25,33,40,38,33,29,32,37,27,38,31,37,35,42,20,37,26,33,28,35,34,34,33,35,30,40,29,33,39,33,45,36,35,38,29,28,31,39,34,37,33,32,32,37,28,38,37,38,41,40,33,35,34,21,29,38,34,20,34,40,29,33,34,41,34,27,40,41,37,37,38,30,34,32,34,14,36,33,37,28,30,30,39,34,37,48,29,39,39,23,13,34,40,39,23,37,30,38,36,42,34,29,36,35,36,42,39,36,37,32,39,39,38,50,42,38,33,38,37,31,36,40,25,35,35,30,31,37,6,29,33,45,33,33,31,20,35,36,32,35,34,34,42,26,37,31,24,43,36,21,32,27,27,44,37,33,41,34,43,41,42,36,31,40,40,38,38,31,38,20,36,23,37,34,35,34,36,35,15,33,39,36,34,39,37,39,38,36,30,37,32,37,33,35,38,35,30,33,39,32,28,39,34,36,39,30,34,27,35,19,34,37,34,24,36,37,34,35,37,19,42,36,34,34,27,27,31,11,41,32,39,17,31,40,33,38,24,36,24,41,43,37,23,38,39,36,32,38,28,37,35,41,38,33,32,41,30,35,36,35,34,36,38,33,40,34,41,32,43,36,30,34,33,32,19,40,35,32,26,36,33,36,37,30,41,37,39,36,38,40,34,34,38,31,40,24,32,33,36,30,32,43,36,36,35,35,29,35,32,40,36,36,38,34,38,36,43,36,40,41,38,34,36,43,33,34,29,36,35,25,34,36,19,42,32,37,34,33,34,31,40,34,34,34,35,36,38,34,32,40,19,38,36,39,31,36,54,36,38,37,21,43,36,36,50,33,24,36,29,22,35,37,43,33,39,40,36,40,32,33,38,8,35,42,29,33,36,41,35,40,41,38,38,32,17,36,36,45,29,40,37,29,29,32,42,31,38,34,39,30,27,32,37,14,38,38,39,36,35,40,38,42,34,37,33,39,34,34,31,33,33,36,38,37,39,34,41,35,37,34,36,35,37,36,39,36,34,39,35,32,38,41,33,36,45,36,35,31,32,35,40,43,39,10,35,25,30,33,34,41,33,35,34,35,36,26,35,30,15,27,29,40,11,38,39,60,37,35,27,33,29,35,34,33,35,39,34,38,32,32,46,35,33,36,35,39,27,38,26,38,35,32,31,35,37,37,40,38,35,36,36,37,34,36,38,39,32,34,42,35,32,33,37,37,43,32,31,25,35,33,34,37,37,33,39,28,37,36,36,31,34,35,36,33,39,39,33,27,31,39,30,35,41,38,35,30,36,38,35,34,31,38,34,32,36,43,27,43,36,31,32,34,30,34,46,32,37,33,27,33,36,34,32,35,33,39,38,44,39,39,32,33,27,34,37,38,32,30,45,42,38,36,41,34,36,35,38,34,33,39,35,33,37,35,39,31,37,22,35,35,34,5,44,33,29,41,37,37,13,39,33,32,35,35,26,35,38,33,41,31,35,39,34,32,33,40,30,37,37,39,39,32,31,31,39,35,34,44,39,40,36,29,39,35,32,34,32,31,34,33,11,31,39,41,42,30,37,35,32,31,36,36,21,46,42,37,36,29,32,37,35,39,35,42,43,42,35,39,36,36,40,39,38,36,30,36,27,40,34,42,34,37,36,39,42,43,27,33,33,33,34,42,37,32,31,30,36,33,33,42,37,37,38,34,35,31,37,35,30,35,32,36,41,33,34,31,34,35,36,40,37,36,41,40,29,37,32,42,35,33,43,32,38,35,35,34,31,38,36,12,16,30,28,32,40,41,36,35,36,36,38,41,38,34,33,33,35,36,30,40,30,35,36,34,32,35,37,38,24,29,33,40,32,36,33,33,39,30,33,37,37,33,35,33,23,37,29,35,39,42,29,24,37,41,37,36,26,36,34,34,36,26,36,32,31,36,41,36,34,33,41,38,33,28,33,39,34,9,36,37,36,34,32,36,32,40,35,43,38,37,43,34,37,26,34,37,29,35,33,29,36,33,40,30,39,43,34,34,35,44,36,35,38,35,37,35,37,32,37,27,41,40,36,36,36,37,7,36,31,35,32,37,38,32,42,36,13,32,36,35,32,35,32,36,36,38,39,38,30,33,30,26,35,29,40,26,38,32,35,30,37,22,35,33,20,38,30,35,41,22,39,31,37,35,25,31,35,38,36,31,41,35,30,32,34,40,37,42,33,22,35,32,23,32,31,36,36,24,37,26,41,32,34,33,37,32,37,32,32,36,45,38,41,37,31,37,31,37,36,32,34,31,42,27,36,34,33,37,33,40,37,38,38,33,39,35,27,30,34,40,42,29,35,23,35,40,38,41,33,34,9,34,39,40,40,37,30,36,37,37,34,37,40,39,37,33,37,34,39,38,29,41,36,30,40,33,39,30,27,40,38,34,37,47,30,41,36,32,33,42,34,39,35,36,46,35,33,41,37,32,33,20,35,42,27,37,32,37,30,32,33,33,21,32,27,35,36,37,43,35,34,34,33,39,32,35,31,27,48,38,36,38,35,30,32,35,41,36,37,41,44,34,33,31,39,38,33,37,41,31,36,26,36,37,42,39,34,37,31,39,27,32,8,36,37,34,36,37,38,40,35,32,42,44,24,35,35,44,35,31,33,31,33,37,33,36,32,38,29,39,40,29,37,43,35,31,28,31,38,43,27,17,31,33,40,40,25,31,48,34,38,30,34,37,31,35,35,33,39,34,35,35,34,33,32,37,42,36,16,39,35,36,37,43,30,34,33,38,40,37,38,33,34,42,31,34,38,38,39,31,36,35,44,37,36,35,34,43,32,35,36,49,33,32,36,33,37,41,35,26,33,34,36,40,39,19,34,33,36,34,35,36,15,35,39,39,36,36,34,42,36,39,33,34,30,33,24,38,36,12,45,31,36,32,32,41,39,35,34,36,28,39,37,38,35,35,34,39,34,29,36,35,37,38,43,36,36,38,39,36,39,38,34,41,38,32,28,38,29,35,39,37,37,35,38,27,34,31,35,31,23,33,30,35,35,36,35,35,33,35,20,39,35,35,35,39,39,35,36,25,38,40,33,33,33,40,37,25,33,34,41,33,34,35,34,33,37,32,31,39,42,40,35,36,39,36,21,36,35,29,30,32,34,34,29,29,35,40,33,26,37,44,33,39,33,32,42,32,41,31,38,34,33,29,35,33,33,32,34,35,34,35,36,37,28,22,35,38,34,33,36,40,45,44,42,43,36,36,33,33,41,42,33,38,35,33,35,31,36,39,34,35,38,37,40,35,38,43,35,35,40,43,32,35,39,35,34,40,34,35,34,20,44,39,37,42,34,38,40,35,36,34,37,40,38,32,26,34,36,35,23,39,39,45,38,35,31,39,36,33,34,37,33,10,43,33,37,39,31,35,33,32,33,34,30,37,46,20,31,37,34,35,35,38,37,17,38,31,30,36,34,29,34,36,38,35,36,24,32,38,33,32,34,33,39,28,32,30,33,39,37,34,35,33,29,34,33,36,42,37,35,34,34,34,13,33,35,35,39,23,35,30,36,33,31,36,33,34,32,34,36,33,35,35,38,37,39,40,35,40,33,37,38,36,33,27,34,38,27,40,31,35,41,36,28,35,36,32,37,42,42,32,15,35,35,38,31,38,38,36,38,33,37,34,33,31,40,38,31,32,38,33,36,27,33,37,34,39,33,14,33,31,32,37,31,32,48,35,35,37,36,34,36,35,37,51,34,30,39,33,31,40,37,40,34,39,38,34,35,37,43,35,33,36,34,34,37,34,39,22,33,36,31,36,37,32,38,40,35,33,34,33,43,38,33,42,31,29,35,34,42,37,34,39,38,34,43,38,34,33,36,33,36,40,21,36,33,31,36,23,41,41,30,35,34,27,30,32,41,39,36,37,36,40,22,32,35,34,28,35,36,29,39,24,38,36,43,38,36,38,38,34,30,35,39,38,38,34,41,36,45,30,32,32,43,40,34,36,36,32,34,35,35,35,34,36,33,30,33,36,36,35,41,45,39,39,34,34,40,33,35,9,30,42,34,36,10,32,35,35,35,34,37,29,33,39,44,38,31,33,35,39,39,37,33,30,39,35,35,31,34,32,34,37,39,39,31,34,37,40,28,23,25,21,33,32,33,31,34,36,32,35,34,37,30,36,41,38,32,36,39,37,31,21,40,37,36,27,32,34,42,37,40,36,38,26,30,33,44,38,34,39,38,37,40,33,29,36,30,33,36,34,34,38,35,33,41,30,37,30,37,34,30,23,38,25,18,36,32,31,14,40,33,31,38,37,33,28,32,32,29,31,29,32,39,38,34,35,32,33,32,38,38,40,29,27,36,23,38,36,26,18,44,34,37,33,32,40,37,30,18,30,38,34,33,31,38,36,34,35,35,31)
VarExplained_mRNA_Residual=c(11,19,25,20,26,30,21,24,17,30,29,30,31,26,24,25,30,27,28,27,21,18,15,19,26,32,33,27,24,23,33,24,30,24,23,12,30,27,29,30,32,30,31,33,33,23,29,26,26,33,29,23,27,30,25,25,20,34,29,35,33,38,25,23,33,35,22,28,26,33,31,34,22,35,29,26,34,35,32,35,31,37,32,31,23,32,25,22,29,25,32,31,35,27,27,30,27,35,32,23,25,28,38,26,36,38,34,26,30,22,36,32,26,29,34,30,28,24,24,23,27,29,41,29,30,30,28,26,29,26,26,38,24,32,22,26,28,32,27,39,40,32,24,34,23,25,28,24,31,29,31,25,28,27,31,32,35,31,19,35,32,33,27,28,28,33,35,32,29,34,33,32,33,28,27,30,38,33,31,29,27,38,30,30,30,24,37,37,24,34,30,32,34,30,29,36,33,34,29,30,33,36,38,35,18,33,28,37,31,35,38,30,27,17,22,18,37,36,28,29,30,40,26,27,25,31,34,42,39,21,30,33,35,32,30,22,33,27,36,28,32,25,34,31,29,34,33,31,31,36,31,25,33,27,27,31,19,31,38,32,31,33,30,23,31,28,41,40,35,30,29,32,29,43,32,33,34,20,28,27,23,41,30,38,36,30,31,34,31,34,34,31,32,31,27,27,34,32,32,27,36,34,27,29,36,33,26,37,28,41,31,40,28,32,35,43,30,33,41,34,32,38,23,36,24,30,19,40,35,34,26,39,25,36,28,25,32,31,11,33,31,34,29,35,31,27,38,35,35,38,32,31,33,16,41,30,35,25,30,31,28,29,41,36,33,38,32,32,29,36,35,34,38,33,30,41,37,37,40,31,36,30,35,21,33,28,33,34,49,25,37,37,27,38,35,33,32,31,27,34,31,22,28,36,32,32,36,33,31,20,32,35,33,33,27,31,28,33,29,37,35,28,27,25,39,21,35,37,34,33,30,37,30,16,25,34,37,34,32,34,35,34,37,25,26,28,25,35,27,35,36,30,35,34,40,38,35,28,34,37,34,24,31,39,31,30,31,40,34,39,28,28,45,32,26,33,33,34,30,30,26,27,39,24,33,38,36,31,39,32,34,37,39,33,29,35,37,31,30,37,29,30,27,32,40,29,29,30,31,35,27,34,32,34,35,31,26,35,25,21,26,34,33,37,36,34,33,32,27,41,35,42,30,34,29,35,31,30,44,37,32,39,16,37,32,28,30,34,38,34,44,35,34,35,27,27,11,41,30,33,41,27,31,35,18,36,32,34,34,35,13,39,35,31,38,37,40,32,39,33,34,36,30,28,35,33,30,45,34,39,9,27,33,35,35,21,33,39,35,27,44,35,33,39,41,26,34,34,41,31,38,38,31,29,33,33,38,28,30,34,34,31,36,26,32,25,25,35,31,41,33,23,28,37,34,30,28,31,40,33,36,35,35,41,31,37,34,33,25,28,26,32,33,35,30,30,35,30,37,30,37,39,31,36,33,29,43,37,28,38,17,31,38,34,39,41,35,28,39,32,36,37,33,35,38,38,31,29,34,30,33,37,32,32,31,30,23,32,33,37,38,22,35,28,38,38,28,34,25,36,38,37,35,37,31,38,31,32,33,41,40,30,21,28,32,42,31,35,38,36,32,27,29,35,47,33,28,37,43,40,34,36,18,26,22,34,46,34,37,33,38,35,35,31,37,36,21,31,35,28,35,43,35,33,23,37,30,33,31,29,29,31,21,33,38,22,39,36,29,28,36,29,33,41,27,44,38,26,35,28,42,34,35,32,35,35,36,25,39,29,28,25,35,24,33,29,31,42,28,28,28,28,33,37,29,31,34,41,29,37,29,37,35,24,16,39,42,34,40,29,44,30,27,24,35,33,35,33,23,36,26,38,40,25,32,36,31,24,24,43,38,37,35,42,30,36,34,30,36,34,30,31,36,38,34,32,47,41,37,42,38,36,39,26,31,37,32,38,24,32,36,32,31,40,31,37,42,28,27,32,38,39,35,33,33,32,36,38,20,32,32,28,40,37,36,35,31,39,31,25,37,36,32,32,38,27,34,35,38,40,36,34,30,37,35,31,39,40,35,26,30,34,34,25,44,28,32,35,29,33,39,45,37,43,34,27,33,32,35,35,37,32,32,24,31,35,32,38,38,32,38,14,38,41,30,28,38,34,39,38,30,31,40,32,37,31,27,25,24,28,33,29,38,35,29,36,37,36,28,31,35,34,36,37,29,31,40,18,36,36,29,35,27,34,35,51,35,26,29,34,40,25,33,29,37,39,33,39,27,36,33,36,40,36,27,32,42,36,41,38,34,35,35,30,28,41,28,35,34,32,22,36,41,13,39,36,35,28,30,38,41,41,24,38,39,29,29,38,43,35,25,44,23,36,36,38,36,39,41,44,28,27,30,32,30,37,31,36,39,32,32,41,35,24,32,34,28,39,34,35,21,41,37,26,36,38,34,34,39,40,42,34,42,33,34,35,22,34,35,29,41,36,34,15,32,29,31,37,43,36,28,33,34,33,33,40,31,29,33,47,33,30,40,36,36,34,26,20,33,28,38,34,36,41,39,30,38,34,24,21,47,41,27,22,33,38,27,26,40,37,26,36,35,39,23,37,42,29,32,41,31,31,32,36,29,35,38,28,32,35,43,34,38,31,30,26,28,39,33,28,27,37,36,32,31,27,41,28,38,26,31,33,34,30,36,35,42,40,35,34,21,31,32,18,34,33,36,41,38,28,31,33,27,36,35,46,30,29,30,34,34,38,30,33,32,31,39,45,38,34,43,34,29,36,36,38,28,26,33,35,35,40,41,42,34,30,34,44,28,40,37,27,37,33,40,38,38,37,14,37,33,40,27,43,35,40,34,37,27,31,36,32,45,32,34,36,38,42,31,39,40,34,48,35,38,35,32,19,32,35,36,47,32,23,49,36,29,18,39,34,40,32,31,37,40,34,35,14,31,29,39,45,33,31,35,31,27,38,32,38,33,29,36,34,31,31,37,40,43,26,38,26,39,34,32,33,33,38,46,29,36,30,20,37,41,34,32,32,44,33,30,40,35,35,31,39,37,36,29,34,37,42,33,35,30,42,34,26,37,41,32,39,44,34,42,34,44,37,35,31,43,31,26,41,36,35,40,20,37,40,16,36,38,32,39,29,32,30,36,41,46,34,37,37,35,32,34,31,35,34,40,38,39,32,34,37,33,28,34,44,30,25,34,32,35,36,34,35,29,30,27,34,46,34,33,37,25,37,33,39,29,28,36,36,34,38,36,40,9,42,35,41,36,35,34,38,35,37,33,37,31,43,35,30,35,38,35,31,34,38,40,38,38,33,41,33,41,32,43,36,37,26,36,20,32,41,40,36,28,38,35,36,39,35,32,39,34,37,40,34,27,40,37,33,27,32,34,36,40,30,40,39,34,30,38,40,33,38,42,33,40,36,35,37,37,34,32,34,27,26,31,35,42,35,34,59,32,36,36,33,30,21,36,22,36,31,35,39,34,23,33,35,32,32,37,43,33,23,28,33,47,31,35,38,30,32,38,40,39,37,39,32,30,45,27,40,37,38,16,31,14,23,42,36,35,38,30,37,36,35,34,34,28,40,35,33,37,27,36,35,38,38,21,34,29,51,33,33,8,35,26,37,26,34,31,40,34,32,40,33,34,39,38,48,35,33,33,31,39,39,39,30,33,36,33,33,34,40,13,36,30,30,30,33,41,38,32,41,38,36,29,36,36,41,40,40,37,40,31,37,33,30,40,38,26,36,34,43,27,32,31,37,33,39,23,37,30,39,31,39,39,30,28,39,36,31,27,6,32,35,33,22,40,35,39,30,43,22,27,31,5,42,27,36,37,22,34,35,43,44,35,31,39,39,35,30,35,38,35,34,35,42,20,32,39,32,37,36,36,33,41,25,38,34,35,33,36,32,35,38,28,33,41,34,45,36,27,39,33,32,33,38,38,35,22,36,32,34,31,37,21,38,36,31,29,34,33,27,37,39,30,36,41,31,33,36,42,39,40,32,27,31,21,33,29,27,34,31,36,30,36,40,37,46,32,40,38,33,36,30,33,41,34,38,34,38,30,33,35,38,36,39,38,35,40,38,36,39,36,34,28,37,33,29,41,35,40,37,41,37,45,40,35,36,21,41,41,32,32,34,34,31,41,39,35,40,38,34,36,30,29,36,35,34,37,35,38,39,33,24,39,39,34,38,31,41,36,36,29,37,43,34,46,20,42,37,39,24,43,30,44,38,43,28,40,33,36,26,44,29,39,42,37,43,41,12,34,33,35,31,38,38,30,32,36,25,35,41,37,28,15,39,35,38,39,41,41,38,27,38,38,30,39,35,36,34,37,38,37,35,31,38,35,27,39,36,40,34,40,44,36,37,17,31,41,40,48,37,7,31,34,33,28,41,29,33,47,34,42,34,39,43,31,31,33,40,22,37,35,18,34,32,30,29,29,46,34,37,32,26,35,31,35,32,33,37,32,36,32,15,32,31,38,37,37,39,39,40,10,38,35,40,32,36,40,33,34,31,48,35,33,28,39,38,40,40,24,31,35,34,29,37,30,31,46,32,38,41,43,16,36,34,35,45,43,38,35,35,42,14,34,32,37,37,41,39,31,10,28,41,28,16,35,39,40,43,31,34,32,30,27,40,18,30,37,38,33,36,28,36,38,32,40,36,40,29,31,44,39,37,42,32,39,36,34,45,36,46,28,29,39,43,37,40,13,36,34,42,24,31,46,37,39,22,36,37,35,39,27,35,30,38,32,39,34,32,38,29,38,40,40,30,38,36,36,38,36,30,38,40,34,38,39,40,36,48,27,44,35,40,35,33,38,34,40,36,32,32,30,28,38,42,19,31,46,36,30,39,38,34,31,38,40,33,39,34,35,24,34,35,18,39,38,34,43,37,23,39,35,39,50,26,40,34,23,35,27,36,37,16,33,37,41,36,33,38,34,33,35,42,33,5,38,32,50,34,38,29,36,36,37,34,40,33,17,37,33,33,28,33,33,32,36,41,42,38,36,30,37,40,43,40,31,8,37,34,40,33,35,33,38,36,38,38,45,30,36,38,32,49,36,40,35,40,24,37,32,41,38,33,36,34,32,35,35,31,35,35,35,46,40,29,43,29,32,37,40,41,40,37,32,23,35,10,36,28,48,38,36,31,36,29,35,28,29,28,37,41,16,37,25,38,10,35,39,17,36,41,40,35,38,35,40,35,35,29,34,40,34,37,29,37,41,37,33,39,36,39,19,41,28,40,40,37,37,39,34,38,37,40,29,37,39,36,37,35,37,28,33,37,32,36,34,37,32,36,37,44,42,34,36,40,33,36,34,40,36,25,37,48,38,35,36,42,33,28,26,33,35,37,28,44,35,29,37,31,30,46,34,35,47,38,31,40,39,39,51,30,40,32,38,38,39,41,26,37,30,32,32,37,34,46,29,42,38,26,37,26,40,36,46,38,36,41,36,32,41,46,30,37,34,39,37,41,33,37,39,32,35,36,31,24,43,38,38,39,41,24,38,38,38,4,31,38,30,32,37,38,9,36,35,39,33,46,27,33,32,40,30,36,36,36,42,42,39,40,41,36,30,33,39,39,37,43,28,39,34,37,35,37,36,27,36,33,47,28,42,33,40,36,13,27,32,34,29,32,37,39,31,42,36,32,22,26,34,36,43,42,28,40,37,36,40,30,38,32,39,31,36,42,42,30,37,32,28,39,36,26,32,34,33,42,39,37,36,33,26,32,38,40,35,36,43,34,36,35,41,34,42,34,41,36,30,38,32,34,32,28,48,31,44,39,35,42,38,45,38,36,39,33,33,38,39,34,32,38,35,33,32,39,36,37,41,37,37,33,41,36,36,16,10,42,37,40,36,35,31,39,31,37,30,30,34,43,39,34,30,34,42,32,30,43,39,37,30,41,45,37,27,42,34,32,40,35,46,36,31,37,34,36,36,33,34,36,24,43,42,27,31,36,29,32,41,30,32,40,33,44,44,42,36,38,36,33,45,38,30,37,36,40,32,39,41,41,36,35,38,10,36,37,30,42,49,34,40,43,31,34,32,39,28,37,27,25,28,36,40,36,32,39,42,37,39,31,34,29,26,36,40,33,41,42,40,35,34,33,30,36,39,20,42,34,29,40,40,36,9,35,36,30,38,39,36,36,29,39,16,41,45,34,37,37,28,38,38,39,39,31,43,37,29,25,37,44,31,44,33,35,34,45,44,20,38,31,17,32,39,38,32,19,27,49,41,34,36,31,26,37,37,39,35,36,32,27,33,38,32,35,31,43,38,38,34,39,30,37,34,32,27,24,29,42,23,31,35,44,39,40,40,34,28,25,32,38,43,35,33,37,39,44,41,27,35,27,34,38,40,40,38,36,31,41,39,37,32,40,31,40,36,35,34,42,35,38,29,42,37,37,41,25,9,28,35,44,33,38,36,37,34,40,37,42,38,34,40,40,34,33,35,37,40,34,31,37,28,30,31,36,29,29,41,33,32,34,54,35,40,37,29,37,39,38,40,41,25,45,35,33,28,34,38,19,31,36,21,37,31,40,35,39,43,45,30,42,28,26,31,46,25,35,42,34,41,33,41,35,33,35,28,41,38,37,25,34,31,36,37,38,38,37,31,42,39,32,33,33,34,38,38,25,39,46,41,38,34,37,41,39,23,36,44,33,6,33,40,33,31,27,38,42,38,41,33,31,51,36,44,36,34,38,30,35,37,32,36,37,43,31,33,37,39,53,33,35,38,34,35,35,36,26,30,16,38,35,33,38,29,32,33,36,39,46,39,37,35,35,33,38,41,31,37,29,45,35,30,35,35,37,13,39,39,38,34,33,42,37,42,29,35,36,38,37,35,30,42,38,36,39,36,25,36,27,38,34,41,41,33,34,41,31,36,29,39,36,33,34,37,21,38,32,36,29,39,29,36,19,46,32,37,45,40,41,17,33,41,35,41,39,42,31,36,37,37,38,36,35,33,36,32,9,30,45,39,29,39,22,34,43,38,41,43,32,28,35,31,44,37,43,37,35,41,40,36,46,39,35,35,42,38,43,31,35,39,36,33,37,38,35,38,36,35,41,38,33,34,45,42,34,36,37,22,33,38,38,42,39,38,41,33,30,29,38,34,41,35,37,43,36,40,33,30,36,42,41,27,42,35,40,30,40,38,40,40,30,34,40,29,42,43,35,29,41,34,41,36,42,35,35,41,45,28,33,34,35,31,36,36,33,45,31,36,36,37,36,43,36,29,39,33,43,35,47,35,38,44,49,38,43,37,32,39,36,42,38,45,23,37,41,41,35,40,36,31,31,33,40,38,37,29,38,34,36,32,36,30,37,37,41,41,36,41,34,33,36,35,34,37,33,34,41,35,24,44,38,34,37,41,35,38,41,39,21,24,37,39,34,44,31,30,39,42,36,34,44,41,42,44,31,43,42,34,32,35,36,39,39,41,38,39,41,39,35,37,8,37,36,40,36,40,39,39,47,35,35,34,33,29,26,25,35,35,45,37,37,34,14,30,35,28,39,38,37,41,32,34,39,40,21,27,28,34,37,46,42,41,42,39,45,38,35,44,33,33,44,51,35,36,35,33,47,40,39,31,32,9,43,36,38,43,22,30,38,37,38,45,38,40,39,34,38,41,43,41,41,38,42,32,40,43,29,31,37,43,45,37,34,37,35,41,38,41,35,38,34,44,35,31,37,34,37,32,30,14,37,38,40,45,38,42,32,36,43,46,41,33,33,37,31,39,36,38,43,39,33,38,46,34,36,45,13,34,40,28,40,41,39,38,34,38,37,33,42,38,40,39,23,42,34,39,34,35,46,36,43,39,31,36,32,39,32,34,43,32,25,33,35,33,41,38,32,40,28,32,38,34,42,36,30,41,45,33,35,26,40,38,36,47,33,38,41,35,42,36,32,33,45,39,41,41,28,40,36,39,36,22,34,41,32,39,24,27,34,31,35,37,28,25,34,30,42,33,34,36,33,21,43,27,33,44,40,41,31,36,35,42,39,22,36,46,35,38,36,42,41,30,36,40,42,38,43,26,36,32,40,24,33,39,43,33,37,39,38,40,38,44,41,32,41,35,35,24,25,30,29,42,41,39,40,37,39,30,11,28,34,42,43,13,47,38,40,37,36,40,36,31,41,32,38,38,35,37,43,37,32,36,46,32,36,39,47,41,38,32,40,32,35,41,44,41,35,29,20,20,23,36,35,34,32,37,35,28,29,27,31,36,37,30,41,39,32,31,26,42,16,35,31,34,39,37,31,37,35,35,40,40,32,38,39,30,33,35,35,26,37,28,33,40,34,33,32,40,35,33,40,43,36,30,42,36,27,41,29,39,28,39,27,18,37,43,38,13,39,39,42,41,36,32,40,39,45,49,38,31,35,33,43,37,35,36,32,32,40,24,31,37,39,40,23,38,37,22,16,28,26,34,37,44,35,30,34,16,35,38,41,31,38,37,37,31,39,26,37)
VarExplained_mRNA_SumofSquares=c(214,544,171,70,40,75,179,1616,272,30,36,38,182,72,123,52,111,102,39,245,185,674,429,150,98,36,98,31,69,63,82,100,36,106,84,94,425,104,32,47,16,220,51,49,213,65,93,76,172,63,272,371,50,187,143,154,41,169,20,60,30,64,23,170,28,25,50,20,88,140,31,76,49,28,20,61,87,33,79,23,28,46,178,81,124,42,11,22,422,74,177,39,23,104,68,34,62,15,29,24,30,64,15,218,26,29,263,10,32,11,27,22,144,37,55,73,96,80,88,122,129,88,22,189,23,81,45,58,210,44,72,13,76,84,70,204,26,27,164,180,16,72,25,15,80,151,15,18,25,14,52,41,120,166,127,92,103,77,57,35,25,36,39,24,35,58,21,63,39,11,11,34,26,167,240,73,18,40,23,36,86,74,472,67,36,117,13,147,925,34,88,74,33,14,54,10,12,35,35,15,31,36,87,38,55,1366,32,29,42,23,26,185,29,50,633,151,23,48,49,51,25,27,230,32,20,40,162,23,24,68,37,24,9,81,36,59,42,20,83,125,50,31,12,85,121,104,21,38,32,24,1693,64,57,865,72,74,65,38,16,61,159,67,18,126,16,35,28,28,12,24,101,28,38,19,13,30,17,49,26,111,41,10,11,63,23,145,74,43,38,12,37,23,12,23,44,22,219,95,31,21,53,55,28,41,16,16,22,51,41,18,69,17,66,25,35,17,48,58,193,63,73,10,85,41,59,38,56,23,631,25,18,88,38,38,303,19,58,132,130,42,42,49,645,15,28,69,24,108,24,43,21,20,29,14,123,267,14,120,28,56,27,72,39,201,1294,35,31,42,83,60,10,30,47,549,47,20,36,20,39,128,13,22,49,130,221,40,34,14,58,47,25,84,60,10,15,60,257,21,31,27,31,49,57,68,21,9,32,20,24,69,58,43,30,131,13,164,13,90,58,8,37,61,30,34,83,140,100,50,24,69,11,116,11,102,87,21,8,61,20,16,42,8,24,115,28,60,79,63,52,19,55,115,21,27,5,27,51,39,40,230,134,69,71,48,46,26,67,97,38,10,23,61,26,59,120,58,11,26,68,98,20,41,82,35,10,11,35,39,35,53,10,37,32,61,15,84,21,74,208,17,36,7,27,21,13,57,26,68,16,70,34,15,25,38,27,66,42,78,43,299,17,70,61,18,87,17,50,16,163,25,47,27,30,47,73,47,19,163,258,22,22,11,101,20,34,24,18,100,1,49,34,24,25,18,29,60,166,16,17,46,18,259,758,36,137,38,49,292,83,73,198,18,21,105,51,21,59,45,22,113,48,86,56,134,163,96,53,23,16,14,128,33,33,51,155,103,40,240,15,98,32,11,24,44,17,35,149,56,10,92,31,35,140,35,28,25,33,148,83,29,62,17,27,25,100,176,45,122,32,35,173,234,18,27,3,35,87,36,20,87,19,53,36,35,101,20,18,78,135,24,20,44,19,52,15,41,11,25,20,26,29,51,45,114,67,4,8,27,39,37,1095,36,33,54,26,48,67,61,47,16,15,18,40,15,79,56,66,117,14,84,41,28,37,83,67,30,141,73,57,63,49,39,75,74,25,14,27,55,20,31,9,44,35,38,66,11,25,30,20,18,314,266,50,22,35,27,21,84,17,27,24,36,44,51,20,19,134,16,54,26,15,12,50,23,130,66,12,19,56,190,10,25,27,82,89,24,290,8,49,30,18,66,10,119,92,16,58,43,32,88,87,71,28,24,58,267,21,39,82,58,88,32,40,35,33,3,41,29,16,20,10,39,53,88,23,24,29,493,38,51,70,48,63,61,116,49,32,19,82,17,68,48,30,21,26,47,14,20,21,29,32,27,120,63,64,39,16,32,41,182,37,13,18,87,53,109,20,51,547,26,22,44,29,68,38,61,57,57,41,40,44,61,63,17,59,106,31,49,71,62,42,26,7,49,31,67,25,25,19,47,46,166,135,113,173,23,22,40,43,36,21,117,10,14,18,17,26,101,79,33,14,37,20,151,63,61,71,18,5,41,24,1311,35,24,57,13,14,20,19,32,12,22,68,18,32,39,14,20,28,16,57,33,75,22,43,58,26,13,10,306,46,22,33,50,27,21,46,40,71,80,97,20,25,10,48,66,37,58,34,15,35,80,17,142,26,59,34,11,7,28,23,157,50,85,34,68,45,51,54,58,16,32,46,19,16,258,113,36,28,43,80,36,33,28,7,46,16,54,60,29,76,29,153,12,22,40,55,1903,37,34,48,13,27,56,6,22,29,14,42,84,18,125,98,84,40,47,72,25,21,13,19,74,9,25,31,36,18,46,13,18,48,60,53,160,30,37,34,31,40,107,63,60,23,54,26,22,59,35,91,52,59,51,144,111,33,10,42,14,21,23,60,34,100,29,53,23,50,42,35,1,492,99,117,52,34,16,18,57,165,23,46,26,8,77,56,35,48,59,19,53,61,36,45,10,14,15,16,24,12,35,69,24,26,46,30,769,36,127,30,55,50,55,16,27,116,20,21,85,36,39,44,78,46,20,44,8,41,260,17,42,16,11,92,18,44,13,34,129,20,16,31,26,11,34,10,12,32,47,58,225,56,219,85,62,35,33,82,45,8,51,42,27,23,155,119,23,24,34,62,48,21,22,49,10,25,17,49,27,50,39,25,27,90,16,45,186,28,8,33,178,28,13,13,27,16,54,23,20,99,26,58,13,16,37,100,26,278,18,54,24,338,26,87,90,68,19,35,47,20,28,81,12,49,31,37,19,36,45,46,49,10,119,19,211,112,17,20,40,9,52,29,40,61,30,24,21,17,124,69,42,28,60,13,124,14,8,68,48,96,75,41,41,34,221,31,55,517,35,75,30,43,27,80,23,17,31,45,20,46,15,21,38,26,20,24,34,24,63,60,18,51,29,18,1068,18,71,13,55,30,27,49,27,120,38,44,48,18,310,48,45,36,21,8,54,47,39,87,153,34,103,37,15,20,110,35,59,48,20,29,19,91,17,28,78,38,8,20,19,57,130,37,124,25,47,24,36,47,10,25,41,35,99,2,40,86,54,23,17,36,64,27,13,145,10,124,23,86,22,73,48,27,21,29,34,123,38,92,118,63,67,33,40,48,31,17,27,82,51,129,130,16,14,261,6,21,12,36,38,36,39,172,25,15,17,8,151,35,211,28,24,15,47,13,27,31,83,93,43,56,21,56,17,70,29,29,31,19,77,18,13,31,694,48,47,32,59,14,15,40,146,25,33,12,20,72,26,31,48,47,70,85,117,13,76,44,35,41,134,172,37,46,111,135,13,10,61,99,17,25,28,98,27,23,46,56,25,23,31,46,46,10,25,49,29,13,50,12,220,68,9,94,96,36,92,18,15,26,15,75,19,51,56,82,106,40,16,32,12,35,12,47,27,50,20,38,19,40,52,37,37,53,45,81,34,18,44,48,52,14,13,15,85,49,71,30,549,51,18,218,240,56,30,23,88,68,66,64,44,14,35,67,14,25,63,59,30,64,53,29,74,88,18,61,72,49,34,49,220,34,74,172,19,40,41,30,15,72,1172,63,39,18,29,15,21,42,9,50,126,201,152,68,59,44,13,54,41,65,82,182,28,4,54,80,28,36,12,27,26,20,35,30,25,81,18,75,1,29,24,279,23,17,129,15,9,27,53,40,35,21,92,41,40,49,12,37,169,75,25,63,25,72,18,20,24,209,84,16,27,15,21,20,28,125,23,35,69,110,15,23,59,124,27,73,30,19,20,11,15,87,21,54,44,25,13,53,1176,34,19,162,24,13,28,46,16,39,28,40,9,30,42,46,96,27,54,13,55,70,247,16,16,50,37,13,28,19,33,14,55,71,35,219,31,36,77,25,47,96,39,46,108,56,18,21,82,15,39,76,23,37,14,45,41,185,16,30,149,12,89,38,35,46,70,62,60,23,31,38,26,26,39,31,139,7,31,84,33,18,7,26,14,47,56,71,37,34,22,15,26,48,20,62,34,31,31,30,47,46,27,20,42,72,14,19,45,65,32,235,192,11,64,237,10,424,19,96,35,31,22,21,34,27,22,39,9,27,10,26,22,15,24,25,49,36,14,195,38,13,32,20,77,3,90,77,39,48,24,112,71,17,15,24,17,70,20,25,3,450,22,33,34,43,91,15,20,99,31,20,18,42,32,22,11,33,57,46,43,50,43,28,32,64,49,18,31,29,17,60,17,17,101,28,35,110,94,28,45,35,23,68,27,12,49,22,31,40,16,50,29,71,26,19,20,42,23,76,47,51,135,13,32,26,12,62,21,19,166,17,40,50,30,35,42,54,34,25,123,23,25,19,453,40,46,35,605,599,12,30,19,63,17,14,190,43,18,23,38,26,38,14,35,11,11,29,60,12,21,68,120,96,94,59,13,30,34,3,9,27,12,1076,46,38,30,27,35,109,71,44,56,15,92,13,20,71,19,23,20,95,24,45,112,22,72,15,43,465,23,100,45,5,33,25,31,29,38,41,39,23,38,65,112,33,134,25,11,16,34,19,22,28,56,13,63,45,21,37,40,17,31,11,8,37,73,78,77,30,44,103,37,9,82,27,8,26,33,12,80,14,22,29,46,25,48,58,5,10,30,21,20,16,260,13,25,38,28,42,19,32,123,18,33,32,59,131,15,35,15,47,40,42,22,32,9,18,47,32,54,12,49,54,9,20,68,12,33,53,73,54,46,51,23,41,34,14,27,26,16,20,73,48,34,20,50,67,41,100,11,43,16,42,59,30,13,10,58,29,81,60,23,96,20,111,60,66,86,36,94,144,27,50,57,21,37,15,134,9,17,38,128,15,25,111,44,2,57,69,10,24,465,39,7,54,31,11,62,9,82,75,64,12,21,92,15,64,44,5,41,195,18,18,5,68,19,19,20,25,60,25,22,13,120,100,33,28,22,68,16,20,29,137,63,621,45,20,60,20,40,31,30,28,138,37,26,9,179,84,15,83,53,11,44,30,87,74,47,10,26,41,34,25,38,109,8,52,42,67,57,47,33,29,23,18,104,21,24,18,80,101,28,50,28,206,33,23,300,11,14,16,34,33,17,11,11,65,169,506,15,10,43,60,8,25,19,38,56,244,36,26,35,41,537,43,61,22,23,87,23,38,27,48,97,15,16,19,40,30,10,35,12,21,25,95,53,26,372,15,27,30,40,32,43,34,20,51,16,23,75,85,26,29,19,72,12,113,21,52,10,52,78,15,33,27,36,39,24,28,25,25,71,46,85,19,40,27,43,32,26,25,10,61,41,50,26,22,25,125,56,25,18,66,59,55,32,32,85,24,31,26,185,18,14,31,31,78,17,24,17,35,227,9,54,24,26,46,12,12,22,29,192,25,8,25,46,13,23,26,51,41,27,34,35,105,12,22,22,16,22,30,117,14,23,2,11,46,71,49,7,327,41,15,124,16,43,48,38,26,44,32,34,21,20,29,30,21,19,7,17,27,203,13,30,15,13,21,39,19,39,62,16,64,55,38,20,13,41,21,49,12,51,66,27,46,55,4,34,15,22,19,54,40,40,26,265,79,37,50,266,52,149,1227,16,41,33,38,12,43,98,29,37,91,33,17,49,17,21,51,816,35,15,235,34,41,15,28,16,18,13,55,39,23,4,46,148,24,19,15,11,33,42,60,63,42,260,24,23,79,30,19,14,227,11,35,13,40,16,75,37,22,17,32,47,65,17,17,43,19,13,10,11,62,160,31,58,21,95,49,19,74,60,49,137,15,11,6,58,42,38,98,83,27,27,55,33,295,170,26,42,177,62,41,25,19,15,39,35,25,17,16,49,20,56,882,12,26,27,145,41,65,54,38,8,101,59,10,65,38,45,18,56,36,52,35,52,16,58,53,18,38,44,54,92,79,30,20,19,97,25,37,64,102,91,17,22,46,102,23,33,55,25,2,47,76,23,19,62,603,18,26,19,97,62,74,13,44,91,26,63,46,47,108,35,11,20,17,20,52,21,30,41,45,22,107,48,13,80,29,18,43,29,27,45,39,17,114,544,59,23,21,55,19,25,33,110,18,65,32,21,90,90,89,17,33,41,24,36,23,14,32,16,22,32,50,22,75,46,32,28,56,44,27,97,41,9,26,15,16,6,17,59,52,25,17,74,529,27,30,16,81,28,26,122,10,14,33,65,35,63,1143,44,26,57,17,102,30,107,31,29,342,20,26,17,23,114,16,26,28,37,53,58,15,24,75,12,30,35,22,21,28,23,38,42,114,112,40,13,52,63,73,35,13,83,30,133,50,72,32,38,23,22,125,30,16,54,52,13,7,45,319,18,35,20,38,44,105,25,19,74,14,24,43,39,14,131,273,34,23,1097,149,13,125,752,13,48,23,26,22,24,323,27,19,280,89,57,6,31,113,59,96,7,17,14,28,13,43,15,100,27,46,77,41,15,46,36,21,24,49,42,30,32,124,72,134,27,15,23,12,17,22,60,31,45,52,91,50,35,23,58,29,31,41,24,119,27,22,34,38,198,40,24,12,7,24,54,32,33,10,13,43,28,16,74,69,36,62,43,18,75,17,23,121,51,17,69,298,32,541,21,22,31,39,31,18,118,23,16,51,58,39,17,39,69,51,67,97,23,54,16,7,621,29,67,17,26,20,142,27,58,168,33,14,55,19,24,122,20,17,36,57,51,71,62,8,15,33,34,73,20,31,10,131,53,37,72,555,33,26,22,61,62,24,25,20,6,38,51,74,50,215,109,35,27,23,1,42,49,38,56,29,39,33,76,31,27,263,16,30,45,12,9,52,115,89,44,37,76,40,68,228,316,40,24,40,25,30,23,24,29,81,94,20,24,44,46,23,14,75,32,48,83,57,43,39,23,64,15,24,42,61,94,33,26,41,23,27,28,33,43,108,66,55,29,35,28,196,27,21,1560,48,47,228,50,29,16,7,80,27,73,29,176,40,22,37,24,128,14,31,34,52,40,24,341,31,32,78,16,16,126,17,15,51,39,13,16,39,22,54,39,24,129,30,64,19,459,135,49,56,661,15,25,64,15,13,18,19,83,109,18,126,6,23,71,18,59,21,56,21,41,23,146,44,47,12,55,63,27,26,15,129,31,19,51,31,34,30,20,154,32,30,24,61,16,28,35,21,52,113,76,29,15,42,27,179,47,51,1259,30,48,19,20,25,45,42,11,28,35,46,14,21,79,23,45,35,34,38,53,18,47,646,22,49,39,12,34,51,29,8,89,282,75,17,5,120,15,98,47,120,49,23,48,40,43,46,87,44,10,25,36,34,16,39,18,48,13,26,13,25,167,81,93,86,254,15,31,17,205,51,141,12,180,62,19,19,25,99,11,24,18,67,27,32,24,16,10,16,21,20,78,56,115,22,47,31,14,26,63,54,26,26,23,218,18,36,15,53,57,36,33,56,24,62,1058,21,47,82,140,19,16,25,33,22,39,233,56,26,44,24,29,22,19,36,44,77,28,43,23,30,27,64,25,114,40,33,40,41,607,26,25,36,18,17,22,60,4,27,51,69,43,40,225,31,61,24,10,27,11,44,93,14,35,1,34,41,19,42,100,72,106,111,172,351,145,22,31,44,59,26,25,26,8,68,38,27,41,44,16,30,63,369,53,32,21,23,141,20,87,90,55,17,30,28,30,210,60,28,40,35,50,15,22,48,28,46,82,107,203,39,22,39,51,783,99,11,50,18,20,28,52,20,53,76,8,50,62,29,2,25,73,61,45,11,14,18,113,45,41,36,69,12,90,13,38,25,4,126,33,44,22,12,18,40,17,50,29,17,22,134,15,93,20,33,116,43,31,147,51,9,32,11,42,135,12,15,35,67,48,48,21,426,22,31,45,53,57,23,8,15,200,42,52,13,88,11,25,20,96,21,92,34,69,10,21,77,97,77,47,43,26,80,16,24,11,37,21,68,44,76,29,3,33,33,24,35,11,17,17,59,402,16,17,18,84,116,11,85,50,68,47,37,76,24,16,85,68,53,52,64,51,59,13,19,12,58,49,23,10,50,31,29,189,16,45,53,82,35,212,47,45,92,10,44,26,38,10,27,26,24,10,10,40,31,259,32,33,24,27,111,53,44,23,16,81,16,54,35,9,73,40,21,71,30,121,9,75,16,14,113,24,109,35,22,29,389,181,133,71,992,32,91,16,21,17,29,19,15,14,22,63,31,645,53,20,11,273,19,28,32,25,19,21,66,22,22,15,75,24,26,16,34,34,62,26,41,33,35,81,79,46,27,628,11,87,37,202,37,31,45,24,40,39,49,38,62,21,212,36,19,1,23,27,224,78,66,37,12,43,19,63,25,53,32,31,81,49,224,92,35,18,51,58,26,150,72,8,36,11,108,42,8,66,52,128,33,122,30,320,29,82,42,32,19,21,159,32,24,13,20,22,16,58,31,33,3,43,51,60,34,54,31,93,9,38,186,75,50,80,35,61,93,16,31,47,79,206,16,22,30,204,29,14,50,98,73,31,59,15,47,9,15,24,137,24,21,83)
vioplot(VarExplained_mRNA_Genotype, VarExplained_mRNA_Diet, VarExplained_mRNA_Age, VarExplained_mRNA_Interaction, VarExplained_mRNA_Residual, col=c("deepskyblue","deepskyblue2","deepskyblue4","dodgerblue4","black"), ylim=c(0,100))

# RIGHT plot
VarExplained_Protein_Genotype=c(28,35,37,25,25,63,32,25,39,20,33,23,20,39,19,25,30,36,26,22,29,37,20,40,18,22,18,32,32,27,22,29,22,35,17,46,17,21,16,25,37,31,20,31,18,16,21,25,24,22,20,39,29,28,28,26,47,36,22,22,29,20,29,20,28,23,21,22,34,23,79,29,28,24,21,42,19,17,31,18,31,33,27,28,21,27,20,27,35,30,25,25,17,41,40,18,33,23,29,51,25,36,24,20,28,18,22,17,22,27,22,22,27,55,44,32,27,23,27,23,29,31,23,19,22,22,27,32,25,20,18,24,14,29,21,42,22,19,23,28,16,27,35,39,41,39,25,20,30,31,37,20,27,22,37,26,25,19,52,21,12,34,34,14,26,22,29,20,36,26,19,21,22,20,20,27,21,21,29,17,30,18,24,29,29,26,21,30,39,26,22,24,22,34,25,18,22,18,24,20,27,21,60,19,28,34,22,19,22,33,21,54,18,34,46,55,23,33,16,19,28,14,16,24,21,22,15,21,17,49,30,18,16,40,31,30,21,18,29,31,28,24,16,46,22,19,24,81,17,19,37,61,19,36,32,25,52,26,17,29,19,31,26,28,28,22,22,13,19,27,30,36,20,19,23,18,18,29,30,27,37,14,27,19,28,35,26,19,22,35,28,26,18,23,27,27,32,16,24,22,24,29,34,27,26,19,20,19,23,21,19,27,52,28,24,27,23,30,31,27,18,22,33,45,24,21,53,24,24,24,22,23,25,29,33,22,32,17,67,17,21,34,22,16,27,30,28,23,21,37,43,43,17,18,17,28,16,20,22,27,32,41,17,27,38,18,17,20,25,21,23,20,20,40,20,16,21,20,24,30,16,21,27,34,33,24,22,31,24,28,21,18,23,18,29,36,24,23,37,23,26,45,21,22,33,30,37,18,26,22,28,21,18,41,16,15,30,43,24,24,22,34,42,46,26,21,20,18,16,19,18,17,25,31,45,24,24,24,15,31,36,22,39,25,39,26,27,18,21,23,37,28,20,24,25,18,18,25,19,31,26,18,23,30,28,23,29,20,21,32,28,17,64,41,20,24,19,28,22,34,27,18,23,21,28,20,27,21,47,37,20,25,19,25,18,32,22,17,30,19,23,24,27,26,17,19,14,18,30,29,24,15,18,41,26,24,18,37,44,34,25,19,20,30,27,20,21,17,21,21,28,18,28,27,17,23,26,21,24,22,27,18,26,39,19,19,20,28,79,28,23,26,24,18,36,18,58,18,24,24,19,49,25,21,22,23,29,31,21,16,16,19,24,25,21,18,28,24,22,18,16,27,24,41,19,18,35,42,34,19,90,32,47,20,15,30,17,26,16,31,27,17,19,20,20,31,19,23,17,19,16,14,16,25,22,22,20,45,29,51,27,18,26,22,37,17,18,25,22,16,17,27,20,21,19,35,28,14,18,23,38,48,21,17,29,18,18,24,36,51,33,21,25,24,19,26,24,35,20,22,24,13,22,24,19,14,23,29,41,21,17,30,22,39,22,23,23,20,26,21,18,21,25,19,24,16,21,27,19,35,24,20,20,35,26,17,35,29,23,34,23,28,19,18,25,19,17,17,28,34,30,48,16,21,16,15,20,36,21,31,31,33,38,17,21,18,29,21,18,25,19,27,39,19,18,20,17,18,33,19,29,15,31,16,35,13,18,22,23,24,31,42,34,40,25,19,39,22,25,36,18,27,24,19,27,22,17,20,29,17,22,26,22,17,31,36,41,24,35,15,19,25,25,30,37,16,26,19,24,22,13,27,27,30,23,25,21,39,15,28,53,29,22,24,21,28,37,21,18,52,20,16,25,19,25,28,24,23,29,26,21,18,52,42,28,27,16,24,31,21,20,27,22,43,25,16,26,30,29,14,33,12,19,17,28,23,33,32,27,21,16,24,22,24,24,22,20,22,23,16,23,34,23,18,19,21,24,22,19,9,16,24,13,35,18,19,33,17,32,28,15,17,22,22,25,29,19,41,40,33,21,22,33,20,22,23,39,21,30,25,15,32,21,19,21,19,50,12,21,20,27,18,23,21,23,19,25,19,23,28,14,22,28,13,18,29,37,30,34,15,19,34,19,18,12,20,36,25,19,39,38,26,30,27,14,57,24,22,31,32,18,20,29,50,29,19,14,32,17,59,28,80,33,26,22,30,23,27,14,25,18,32,20,25,18,39,17,28,14,28,46,36,21,24,35,42,22,26,17,16,38,33,34,22,17,21,37,68,17,27,25,17,21,26,25,21,31,20,29,39,17,52,28,17,16,13,23,16,45,26,21,20,14,18,23,27,19,22,23,21,26,32,34,21,28,28,35,24,21,46,38,25,27,82,18,16,17,50,29,17,14,17,22,25,19,18,30,20,59,41,36,17,35,30,21,23,43,18,21,19,41,20,23,34,24,27,24,31,39,19,36,24,22,31,27,31,18,42,19,20,25,15,34,23,23,17,22,17,24,55,17,17,24,23,21,24,32,26,34,24,17,26,21,23,17,25,12,22,25,27,19,22,43,44,24,22,20,12,30,14,19,17,31,18,31,29,18,24,20,21,16,23,36,28,18,34,41,55,25,16,22,20,16,60,60,36,25,17,21,39,50,47,27,27,21,26,27,16,26,20,30,15,14,24,20,30,18,29,26,17,30,32,23,29,21,28,15,19,20,21,27,17,17,28,25,35,29,22,21,33,26,23,18,38,64,18,22,22,38,26,19,24,17,29,17,14,35,21,32,16,23,26,20,22,17,16,29,28,17,29,24,19,35,35,21,30,22,26,24,25,18,15,18,20,24,23,20,69,27,75,21,26,19,20,19,20,21,24,28,33,19,24,21,18,26,24,26,14,28,18,22,23,33,26,19,19,32,21,17,35,17,15,37,20,14,25,38,15,30,34,23,18,15,23,20,18,17,60,19,30,51,16,28,21,17,17,29,32,17,31,29,57,54,34,37,56,18,22,40,22,31,25,16,22,27,21,49,34,40,34,26,20,13,28,28,19,20,15,21,18,22,19,18,35,18,24,20,26,22,14,20,19,25,31,18,47,28,24,27,36,21,19,26,22,26,18,20,18,31,30,15,24,22,27,16,18,43,17,12,27,29,23,21,22,21,20,15,19,22,20,30,22,19,20,20,20,18,31,20,33,78,33,61,24,21,22,16,32,33,26,22,39,19,25,15,19,27,23,26,16,20,20,23,25,21,28,16,21,23,24,24,62,21,27,23,36,18,23,20,26,26,16,22,20,21,24,20,54,15,16,16,29,19,22,18,20,40,12,20,28,25,18,29,23,21,24,20,22,16,21,23,23,27,38,18,18,14,12,20,20,14,21,20,26,28,31,20,33,26,18,19,19,37,18,17,23,35,17,27,19,28,25,18,24,34,19,17,20,23,23,27,22,23,15,29,16,24,28,16,42,36,40,21,18,17,22,28,22,17,46,28,19,20,36,19,26,28,22,23,15,21,34,20,20,44,18,22,26,22,16,25,67,37,12,25,24,23,29,25,24,18,21,29,17,36,33,15,30,32,27,28,26,14,21,14,24,25,35,33,24,17,15,21,23,22,27,17,24,46,24,39,49,35,25,19,34,32,26,25,18,38,37,39,26,20,18,32,28,51,29,14,26,57,20,18,20,20,35,79,23,19,14,23,27,23,18,29,38,20,31,21,26,27,26,24,21,18,30,21,22,27,24,18,30,12,23,20,20,28,24,28,25,29,19,19,21,17,24,16,25,19,20,36,29,19,27,35,16,31,33,32,50,25,26,26,20,44,19,21,42,35,26,44,26,24,21,23,19,22,18,56,43,40,20,32,31,87,31,21,17,31,32,22,18,22,38,19,49,14,36,19,22,21,23,29,65,39,60,17,36,29,37,23,24,27,37,15,26,19,24,34,15,25,27,15,24,20,20,21,27,23,47,17,17,23,19,20,22,23,22,25,21,24,24,24,18,24,16,28,19,13,19,20,28,39,14,25,18,16,32,43,23,19,30,28,29,30,24,16,27,22,24,22,22,21,18,16,39,30,28,28,15,19,19,46,18,20,30,34,28,24,17,20,16,25,23,37,16,29,24,22,26,24,17,41,20,22,17,20,17,11,20,39,19,13,26,23,23,37,32,18,27,19,20,22,17,25,19,22,19,22,15,21,21,39,20,36,23,32,21,14,12,29,15,27,32,14,19,14,24,20,40,43,30,29,20,19,28,21,20,33,22,44,22,31,28,14,21,24,22,16,25,25,13,16,22,14,19,22,21,16,86,29,33,28,86,19,47,19,31,57,20,20,21,17,88,35,34,30,26,31,21,38,27,35,20,38,21,21,52,60,17,21,21,31,23,14,22,21,22,29,36,26,16,24,14,19,23,19,22,25,17,26,22,31,16,20,33,21,39,22,22,14,33,16,22,19,18,36,23,24,16,37,27,20,21,18,24,20,22,20,23,46,33,33,39,47,20,57,17,14,26,19,20,35,22,39,24,24,29,25,25,21,22,32,18,25,31,21,70,45,23,19,21,16,20,29,13,41,19,36,26,29,58,41,29,16,22,15,21,43,24,30,27,15,20,30,38,36,33,26,25,22,46,26,26,19,29,17,26,17,35,21,17,20,19,35,21,19,36,24,24,31,18,14,21,14,78,22,46,16,16,8,33,47,29,51,21,30,27,22,26,31,40,18,27,19,24,19,19,17,20,18,29,20,20,30,14,22,29,21,17,21,16,21,22,27,53,25,24,21,22,24,45,55,31,15,41,44,27,23,35,17,44,23,16,20,21,36,15,20,14,18,20,15,16,20,23,20,28,32,28,48,31,17,21,24,24,16,30,22,27,25,20,16,19,22,22,37,15,24,37,18,34,17,32,16,22,19,84,18,20,58,55,18,17,25,13,16,35,33,14,26,29,21,28,19,42,27,24,27,19,15,32,24,22,15,27,22,20,28,24,27,41,15,15,33,19,19,76,26,15,23,38,15,22,28,22,22,38,19,76,32,22,22,21,21,22,55,17,25,18,30,46,42,28,32,24,48,25,23,82,22,32,19,17,23,30,32,20,42,21,41,75,27,22,23,24,79,20,19,22,22,28,25,16,63,18,20,23,17,24,20,25,33,18,17,16,23,24,29,26,30,16,27,44,17,16,20,26,26,22,16,45,26,20,22,14,26,20,18,19,20,42,24,28,21,35,17,22,33,39,34,21,35,27,40,18,30,21,45,19,69,44,19,14,18,33,15,23,20,27,35,19,22,19,15,23,20,15,19,19,31,33,18,27,22,18,21,17,25,28,34,29,21,20,15,31,22,29,33,19,24,25,35,21,28,29,24,38,45,25,21,13,33,23,24,36,26,36,24,25,15,16,25,18,25,16,21,12,22,30,30,17,54,24,30,33,21,14,37,29,15,25,44,21,32,19,24,32,28,33,25,22,21,20,29,19,12,22,16,26,33,25,17,16,21,14,22,23,30,22,25,17,21,26,14,18,19,26,37,25,46,30,16,18,16,18,21,20,42,25,39,22,19,30,36,16,35,23,21,30,34,21,25,19,15,15,21,27,26,83,17,33,27,59,25,18,38,18,17,20,53,14,29,20,28,31,27,15,32,21,16,29,46,19,18,55,27,22,54,21,17,19,17,20,21,22,17,19,38,39,32,18,27,35,16,32,15,24,21,24,33,25,32,48,25,25,28,16,18,21,28,16,22,16,42,35,27,28,22,21,15,28,21,27,18,16,42,24,20,26,26,18,36,22,24,34,20,17,21,25,16,18,20,30,12,27,17,36,24,22,19,20,18,36,28,15,17,26,22,32,17,38,18,24,19,28,18,28,24,33,23,26,20,23,20,21,17,16,15,19,19,22,33,25,40,25,28,50,19,29,19,17,18,34,24,15,17,29,84,16,18,16,18,19,17,18,45,22,14,22,20,23,28,18,31,28,22,17,27,20,25,22,36,21,18,19,21,17,21,15,26,20,20,31,29,17,24,46,20,23,16,30,53,23,20,24,24,25,28,31,18,72,16,18,18,17,20,19,31,49,21,39,32,24,17,25,20,27,23,17,18,14,21,17,25,24,29,18,40,15,19,32,22,20,19,18,21,27,30,31,39,20,17,26,22,45,35,19,18,17,23,41,16,19,28,18,23,18,23,18,23,24,13,23,25,20,21,53,15,23,18,27,16,18,35,52,23,18,27,25,26,24,21,19,55,21,27,17,23,44,16,19,15,28,20,29,18,24,33,24,19,43,57,22,32,23,25,24,24,38,44,15,21,35,38,30,14,34,40,24,28,25,37,28,21,21,23,19,16,19,28,29,19,27,35,21,28,16,20,25,21,37,20,21,23,31,40,28,22,23,33,33,17,24,15,28,19,26,20,31,24,18,20,13,19,33,15,15,22,24,24,20,24,21,26,16,19,26,14,24,31,16,24,27,30,26,35,16,23,14,21,29,22,19,22,12,30,25,69,19,21,24,20,20,39,31,18,21,30,22,27,36,22,26,25,25,25,19,22,45,28,38,16,23,28,22,22,19,24,23,32,42,22,16,26,17,38,87,25,67,22,16,25,22,25,28,28,15,19,19,21,24,35,19,19,30,27,21,13,16,37,23,28,31,40,15,36,13,16,47,24,24,23,22,24,39,20,18,12,15,35,21,22,40,31,25,18,33,23,19,19,26,31,26,27,22,18,17,22,22,57,20,40,41,41,23,31,27,24,28,69,35,18,19,36,19,24,54,17,27,23,16,29,17,19,17,34,17,21,35,24,26,16,29,28,27,21,16,20,28,23,18,26,35,18,36,30,28,23,43,40,18,20,16,27,17,14,21,23,22,14,23,30,24,20,41,20,23,17,26,24,30,37,27,36,54,28,32,49,20,24,20,17,18,20,42,20,28,33,20,25,17,23,27,27,21,22,51,23,42,23,46,19,21,24,30,23,29,38,20,20,23,15,23,21,29,31,13,31,21,24,22,20,23,15,42,29,19,17,21,19,17,22,26,21,18,18,28,20,27,18,21,21,31,18,17,37,19,19,18,22,28,24,27,22,17,23,20,25,42,32,21,35,30,24,25,15,20,39,26,22,21,26,16,23,20,15,60,17,20,21,19,14,20,31,16,28,31,15,33,21,50,18,23,23,73,33,24,23,30,20,25,19,23,34,17,23,25,31,19,19,21,19,16,24,19,27,30,30,26,23,18,20,23,17,15,20,17,25,31,26,23,54,27,31,36,21,41,18,39,26,18,18,18,23,33,17,28,43,36,23,26,39,30,20,22,13,20,20,18,20,28,18,25,17,22,25,12,17,17,17,22,20,26,13,20,22,25,21,28,19,28,29,49,40,23,18,31,23,27,22,26,33,12,19,16,27,19,20,23,37,19,35,15,50,34,34,19,14,24,34,17,28,25,44,18,29,23,24,16,20,19,21,28,16,23,36,18,29,73,24,17,23,11,21,21,19,14,19,24,26,19,30,20,19,36,26,40,25,29,24,19,28,15,24,17,19,57,22,41,15,64,14,30,26,16,16,36,29,24,21,19,38,35,25,85,23,19,76,28,20,22,31,72,17,25,16,32,25,36,22,18,24,17,30,29,36,24,23,46,24,21,17,17,28,15,55,30,20,14,19,23,19,12,24,19,28,22,28,17,21,18,14,25,16,23,19,35,36,20,18,24,24,26,28,17,19,18,51,20,27,19,18,64,23,23,17,20,22,33,17,23,32,17,24,20,34,19,27,28,16,22,37,19,32,25,19,14,27,24,28,21,26,27,22,25,20,21,57,20,24,16,16,43,22,18,25,25,32,20,18,27,23,21,48,18,16,77,31,19,22,33,23,31,35,36,14,15,19,22,24,21,22,18,23,22,15,28,19,18,20,31,15,22,19,27,29,30,22,16,21,25,24,25,29,20,25,20,24,20,24,46,76,20,27,40,21,19,22,20,19,14,23,19,16,21,19,21,24,27,24,21,25,18,16,20,21,19,68,18,19,31,26,24,21,18,21,18,20,21,40,33,24,15,34,21,26,22,55,47,30,29,66,23,24,18,24,23,24,38,23,30,18,41,28,18,24,21,24,39,15,17,22,41,19,20,19,37,59,31,30,14,14,19,49,26,23,27,22,46,17,21,16,23,23,18,37,19,26,29,21,29,20,28,22,25,25,22,16,83,37,47,20,28,27,23,17,20,29,24,19,32,30,44,73,29,19,21,24,31,29,41,29,24,46,42,24,18,23,25,60,27,18,27,34,36,21,17,44,37,17,51,28,25,20,28,15,18,22,15,24,33,21,33,28,37,18,27,35,24,19,19,19,38,20,14,56,25,19,15,21,23,24,21,41,33,33,23,18,19,30,21,19,18,40,15,30,20,23,20,22,22,19,25,28,29,14,33,19,26,16,41,28,18,21,31,20,24,69,19,16,23,20,24,27,27,19,15,25,18,18,19,15,22,18,15,22,17,17,25,19,29,19,37,26,20,14,21,35,39,20,28,29,24,33,34,21,41,20,19,25,18,27,17,22,16,31,25,38,23)
VarExplained_Protein_Diet=c(35,7,34,12,17,4,17,6,19,14,19,22,40,10,45,2,1,22,29,3,11,20,42,11,34,5,0,12,2,17,4,16,9,27,1,3,13,2,0,35,7,11,0,0,0,0,2,35,3,3,8,10,1,19,20,8,15,24,1,5,6,23,5,23,1,0,2,0,32,2,0,0,0,0,0,3,5,16,20,1,1,4,19,14,5,0,0,13,26,4,11,0,3,1,0,0,10,2,9,6,1,5,3,8,14,5,4,5,0,4,1,0,25,5,9,8,11,1,4,10,8,15,0,8,3,14,4,11,10,1,7,2,0,2,0,5,0,7,8,23,0,2,22,0,18,5,2,2,24,2,3,0,0,3,2,1,2,0,1,0,1,10,1,1,14,5,2,6,2,0,1,1,3,6,4,2,0,0,4,1,18,31,6,7,2,0,0,0,5,2,5,0,0,2,0,7,1,0,0,2,1,1,1,2,0,22,0,0,1,1,15,0,5,6,4,7,1,7,0,2,1,1,25,0,25,6,0,2,1,3,27,1,4,1,2,27,2,2,3,1,0,2,1,0,0,1,9,2,2,0,8,0,1,17,12,2,7,2,12,6,7,6,2,4,6,8,0,2,0,0,11,12,4,0,5,0,0,10,3,2,8,6,1,1,26,2,4,0,1,1,3,1,0,5,5,13,19,0,0,3,0,1,0,1,8,3,1,3,0,10,5,1,0,2,1,0,3,0,15,1,1,5,0,19,9,6,17,5,5,0,7,34,3,6,0,1,0,18,6,8,1,4,3,2,1,10,0,15,0,18,1,13,1,3,5,17,0,6,6,2,3,5,0,2,13,0,2,4,2,3,0,0,0,34,0,0,5,1,0,6,0,1,0,4,18,2,7,5,1,2,1,4,0,1,4,15,7,0,3,4,3,4,3,7,15,14,5,2,6,0,3,3,21,0,0,1,6,2,0,1,1,0,5,1,2,1,0,4,2,7,1,5,0,0,15,0,4,5,1,0,7,2,0,3,14,3,7,4,1,5,1,13,0,3,12,7,1,3,2,5,4,0,5,14,4,0,2,2,0,0,2,1,2,1,6,3,1,2,2,3,0,1,8,3,0,0,0,1,0,3,1,0,0,5,2,5,3,5,4,1,1,13,1,10,0,0,1,3,6,11,1,1,2,3,2,2,16,0,0,1,2,1,7,0,1,2,6,0,0,0,2,2,6,6,1,5,1,2,26,1,1,1,4,1,24,0,2,3,0,0,3,1,1,0,0,1,0,0,1,1,0,0,0,0,3,3,8,2,5,6,0,0,0,1,1,1,2,0,1,3,0,0,5,0,4,0,26,0,6,4,0,1,1,0,0,1,0,18,1,7,19,0,9,0,1,0,1,13,0,1,4,0,0,17,0,1,2,2,4,2,0,1,2,1,8,0,5,2,2,1,1,2,0,0,22,7,1,1,1,0,0,0,0,1,1,0,1,5,3,1,1,2,1,3,0,3,0,3,1,0,1,16,0,5,10,0,1,4,0,2,0,0,0,9,0,0,1,2,6,2,0,1,0,0,20,0,0,4,0,2,2,0,1,2,0,1,1,3,6,1,0,0,0,0,1,0,1,4,4,2,0,0,0,1,1,0,3,0,1,2,20,1,0,3,5,1,0,7,10,0,1,1,0,4,2,0,1,25,4,7,12,0,3,0,6,3,0,0,36,0,2,0,23,0,4,1,3,9,8,0,3,10,2,1,3,4,1,1,0,2,0,1,0,16,6,1,0,2,5,4,4,0,3,9,2,0,1,1,1,1,0,19,9,0,4,1,0,2,0,7,1,0,5,1,2,0,0,1,1,0,2,0,2,4,2,2,0,2,2,9,0,0,0,1,15,0,0,0,0,1,1,1,1,4,0,1,1,4,1,1,1,1,0,4,7,3,1,18,5,3,2,5,1,1,9,0,0,6,4,2,8,5,1,2,1,0,2,3,0,0,1,2,1,3,0,3,0,0,2,7,1,0,2,0,0,1,14,4,0,0,0,0,0,4,1,2,8,0,0,3,0,22,4,3,7,0,0,2,0,0,0,0,11,0,4,1,0,0,0,0,1,0,0,2,15,12,1,0,2,0,0,0,0,0,0,0,0,1,0,2,3,3,7,0,0,0,4,0,0,4,1,3,2,3,1,4,3,2,1,0,0,0,0,9,2,4,0,1,1,4,0,12,0,1,2,1,0,2,0,3,2,0,0,0,0,13,1,0,3,0,1,2,14,0,1,2,2,0,0,2,7,0,0,1,0,0,1,8,4,20,2,1,4,0,1,0,2,0,0,5,9,3,4,0,7,18,1,7,3,1,3,4,1,3,2,1,3,1,0,0,0,4,0,0,0,6,0,11,4,0,1,0,1,7,0,0,0,0,1,0,0,3,8,0,1,1,1,0,14,5,0,5,2,10,0,1,1,0,0,2,12,0,0,8,6,1,0,2,0,0,10,0,3,2,1,1,6,0,5,9,20,0,3,3,3,1,15,0,0,0,1,0,2,1,9,0,0,0,6,0,1,9,4,3,4,0,0,1,1,0,0,1,0,0,5,0,4,1,0,1,0,0,0,0,2,0,3,11,0,0,0,0,0,0,0,0,10,0,0,1,0,5,0,10,0,0,0,3,6,0,0,0,2,0,2,5,0,0,1,0,1,1,0,3,0,0,2,1,1,0,0,0,1,16,4,1,0,5,3,0,4,3,2,5,0,6,0,0,0,1,1,1,0,2,1,3,0,1,2,2,0,0,0,1,0,0,1,1,0,11,1,3,0,10,2,1,0,1,1,4,2,0,1,0,1,1,3,0,5,0,5,1,0,0,0,1,1,0,3,1,0,9,2,0,3,13,1,0,7,1,0,1,0,3,1,5,22,2,0,2,1,4,1,0,0,6,6,0,0,0,0,0,5,1,0,1,0,2,0,0,1,0,3,0,0,0,1,0,0,5,0,1,2,19,0,11,4,3,9,0,0,0,7,2,2,3,0,1,8,2,3,0,3,11,0,1,1,0,1,0,2,0,1,1,4,0,1,0,0,1,2,1,0,3,6,1,1,1,5,8,2,10,13,0,3,0,0,3,2,1,0,16,0,3,0,1,0,0,1,2,3,0,0,0,1,0,0,0,1,1,10,0,1,1,0,2,0,1,2,4,2,1,0,0,4,2,0,0,1,0,1,5,2,0,0,0,0,1,4,1,2,1,0,0,0,6,0,3,2,3,2,4,1,9,1,1,3,1,1,2,7,2,1,1,2,2,2,1,0,0,0,3,1,0,2,0,6,0,4,0,9,0,1,1,0,3,0,0,2,2,0,0,7,5,2,0,2,1,2,0,1,1,0,1,0,0,1,2,1,0,2,0,2,0,7,0,0,0,4,0,2,0,0,2,4,14,0,5,0,0,1,1,1,0,1,3,1,1,1,3,11,10,4,0,0,0,0,0,0,3,5,1,1,3,0,0,20,7,3,0,10,1,0,0,0,0,0,0,1,3,0,3,2,3,0,8,1,4,1,2,1,0,1,8,0,1,0,5,4,2,0,2,4,1,0,2,3,6,1,0,1,7,1,0,0,2,1,33,7,1,0,2,0,1,1,0,3,2,5,2,16,0,0,6,2,2,0,0,0,5,0,1,0,2,4,0,10,3,1,0,2,0,0,0,0,4,1,15,1,1,0,0,0,0,0,0,0,0,3,2,2,0,0,5,1,0,1,0,13,1,2,1,0,1,1,1,3,6,3,0,1,0,0,0,0,2,1,6,0,8,1,0,1,1,2,0,7,0,0,1,0,0,5,41,0,1,0,0,7,0,4,7,3,0,0,4,0,2,0,0,0,1,14,5,6,9,0,1,2,4,4,0,0,18,0,6,0,12,0,7,0,0,6,15,0,15,0,1,1,1,0,4,0,2,0,0,1,0,0,0,0,0,2,0,0,0,1,0,7,0,2,0,2,1,0,0,0,0,0,0,0,0,4,0,2,1,4,2,1,1,6,1,3,4,6,4,2,5,0,14,1,4,0,0,0,1,0,4,0,0,0,0,1,0,1,0,2,25,0,0,1,4,0,4,4,0,0,0,5,1,0,2,7,1,1,2,4,2,0,0,0,0,1,4,0,2,15,0,1,7,12,2,4,1,9,3,0,1,0,0,2,1,3,1,16,0,0,7,2,2,2,1,11,1,0,2,0,5,0,5,0,0,0,5,0,0,0,0,1,0,1,0,0,1,4,0,0,0,7,3,1,2,3,1,0,1,0,2,2,5,1,0,0,2,0,0,1,0,0,1,2,1,8,1,1,0,3,1,4,0,1,3,6,0,1,0,2,3,0,1,1,0,0,0,0,0,2,0,2,2,0,0,2,1,2,1,6,2,1,0,15,1,6,5,0,2,0,3,1,2,0,1,0,2,0,1,0,0,0,1,5,0,14,2,1,0,1,2,2,2,2,0,6,2,0,0,0,1,2,1,2,1,3,0,0,0,10,1,0,2,4,4,0,1,0,1,0,1,0,0,2,0,0,8,0,0,2,0,0,3,14,6,9,0,2,1,7,0,2,0,1,2,11,2,1,1,5,0,3,1,0,2,0,1,0,10,0,2,2,0,1,1,0,0,0,0,2,2,0,0,2,0,5,3,0,0,1,1,0,3,2,0,5,0,0,0,0,0,0,0,2,0,0,0,0,1,0,2,0,0,0,6,1,4,1,0,11,0,0,4,0,1,4,0,4,6,0,1,2,0,0,2,1,20,0,0,1,0,1,0,0,1,0,1,5,0,1,5,1,0,2,1,1,4,7,0,0,1,0,0,6,3,0,9,0,0,2,16,0,0,0,4,1,1,2,0,2,2,0,0,2,0,1,6,4,0,2,4,0,0,0,5,1,0,15,0,1,0,0,2,2,6,0,1,0,1,1,2,3,0,0,0,0,0,0,0,0,0,0,1,0,0,0,6,1,1,2,0,0,1,1,0,0,1,1,4,0,20,0,1,0,1,4,2,0,8,0,3,1,4,1,2,0,3,2,0,0,10,8,1,0,0,0,0,2,0,0,3,1,2,1,0,4,0,0,2,0,1,0,0,1,1,2,0,0,15,0,1,10,1,3,3,1,1,4,3,1,0,0,3,5,2,0,1,4,4,1,0,7,2,0,2,0,1,0,0,1,0,6,0,4,0,1,4,0,0,1,3,0,1,0,3,0,5,0,0,0,0,3,0,0,3,0,0,2,0,2,0,1,2,1,0,7,0,3,9,1,2,37,2,0,0,0,2,4,1,0,1,2,1,1,0,3,3,3,0,0,2,0,2,1,0,3,1,1,1,0,2,0,2,4,0,1,0,0,2,3,0,1,0,0,0,5,0,0,3,2,1,0,1,0,2,5,3,0,32,0,0,1,3,0,2,0,1,0,0,3,0,0,0,3,0,9,2,2,0,5,2,0,0,1,0,10,15,1,0,1,2,0,0,0,0,1,0,1,2,0,2,0,7,8,0,0,2,10,1,1,0,0,1,0,0,0,0,0,0,0,1,4,1,2,1,0,0,1,2,0,1,3,0,5,0,1,2,7,1,0,0,6,7,1,0,0,0,0,1,1,1,0,0,25,0,0,1,5,0,0,1,0,3,0,0,0,0,1,13,3,1,0,2,0,1,3,0,2,23,2,0,3,3,0,10,5,10,5,0,3,0,0,0,0,4,1,0,1,1,0,0,8,0,0,0,1,2,1,2,0,0,1,0,0,2,0,9,0,0,0,4,3,0,6,1,0,10,0,3,0,1,1,0,0,2,0,1,0,0,0,0,0,5,3,0,2,2,1,0,2,3,0,13,0,2,2,4,3,3,1,3,2,0,1,0,10,0,2,2,0,1,2,1,0,5,1,1,1,2,0,0,1,2,0,1,2,0,0,0,1,2,5,0,14,9,1,0,3,2,5,0,4,1,0,12,1,0,0,2,0,0,0,0,1,3,0,1,0,1,0,1,0,1,0,4,0,3,1,1,1,0,0,6,1,0,4,0,2,0,0,0,1,0,1,14,1,0,0,1,4,1,0,13,1,1,1,2,0,0,1,0,0,0,3,0,2,0,0,1,1,0,1,1,0,3,1,4,0,3,0,1,0,0,2,4,0,3,3,2,9,3,0,2,0,2,6,9,3,1,1,6,9,2,0,1,0,1,0,0,38,0,8,0,1,9,0,0,6,2,0,4,1,1,4,0,0,3,1,2,0,0,0,2,1,6,1,1,0,0,3,4,1,5,0,0,3,0,2,3,0,4,0,0,1,0,1,1,4,5,2,0,0,2,0,0,0,0,0,0,8,11,0,0,0,3,0,0,5,0,0,0,3,1,9,6,2,0,0,6,0,1,4,1,0,19,0,8,0,2,1,7,1,0,1,4,2,3,1,0,3,0,0,0,1,3,3,0,0,0,0,0,0,0,0,0,0,0,0,2,1,2,6,1,0,3,1,0,0,0,1,2,3,2,2,1,0,0,1,0,4,1,1,2,0,0,0,1,0,0,8,0,5,0,0,0,12,0,0,0,0,2,0,0,5,0,7,2,0,1,1,2,0,1,0,0,0,0,0,5,0,0,1,2,0,0,14,0,3,0,0,9,4,8,1,0,1,0,4,0,3,17,4,0,0,2,0,0,0,0,1,0,1,3,0,0,4,1,1,1,2,0,1,0,0,1,0,0,4,2,1,2,0,1,1,0,1,5,6,3,2,2,8,0,0,9,2,0,0,0,4,0,1,0,1,5,1,1,0,1,0,4,4,0,0,0,1,0,0,0,0,0,3,12,0,1,3,0,3,0,0,1,9,5,2,0,0,0,0,0,0,2,0,0,0,0,13,0,0,2,0,1,1,2,1,7,28,5,2,0,1,0,0,1,10,5,10,1,3,14,0,1,2,4,9,1,0,0,0,2,0,1,1,1,0,0,0,0,2,4,0,3,0,0,3,5,0,3,2,5,1,0,0,0,4,0,3,0,0,0,0,2,0,0,0,0,1,4,3,2,0,0,1,0,0,5,2,0,0,2,10,0,0,0,5,1,3,1,1,0,0,0,13,1,0,2,1,1,4,5,0,0,1,0,0,0,0,0,0,0,11,3,0,8,0,1,0,1,2,7,1,0,10,2,1,0,2,2,0,2,4,1,0,2,7,0,10,0,2,0,1,0,2,3,2,0,0,6,4,1,1,0,0,0,3,0,0,11,0,0,1,0,0,0,3,0,1,0,0,16,1,0,0,1,1,0,1,0,0,0,1,1,1,6,0,0,0,2,0,1,2,1,0,0,0,2,1,0,0,18,1,0,1,1,0,0,0,0,2,0,0,6,19,1,4,3,0,3,0,2,0,0,1,10,3,0,2,0,5,0,2,0,1,0,0,0,1,0,0,1,0,1,0,3,0,0,2,3,1,2,0,0,2,1,2,0,0,0,0,1,0,1,2,1,6,1,6,2,19,0,1,2,0,0,3,1,2,0,7,0,1,1,0,0,0,1,0,0,0,1,9,1,1,2,2,0,8,0,9,1,1,0,7,0,0,9,2,0,0,4,1,0,0,1,1,0,0,2,0,2,0,0,20,3,1,0,3,0,0,3,3,1,0,0,2,2,0,0,1,1,0,0,7,0,8,1,4,0,0,1,1,0,0,4,0,11,2,1,0,3,0,3,1,1,2,9,19,6,0,0,0,0,2,0,1,2,2,2,0,9,1,0,1,0,0,0,52,0,4,0,0,0,2,0,3,26,0,0,5,9,1,0,1,1,2,0,7,0,0,1,0,1,4,0,1,0,1,0,1,2,1,1,0,1,2,0,4,1,0,2,0,3,1,2,5,0,0,1,0,3,8,1,0,4,4,0,1,4,0,13,3,0,0,10,4,0,0,9,7,3,15,2,1,2,1,2,1,0,0,3,0,0,2,1,0,0,0,0,0,1,0,1,6,0,0,1,4,1,2,0,0,1,0,0,1,0,0,0,1,0,3,1,2,7,1,0,2,2,1,1,0,0,5,4,2,11,2,1,1,0,1,0,0,1,1,0,0,9,0,0,1,1,2,0,0,1,0,0,7,1,0,1,4,1,0,4,6,1,3,0,1,0,7,5,0,0,2,0,0,0,1,3,2,0,6,8,0,0,4,17,0,4,1,1,0,0,0,1,1,3,9,3,0,3,1,1,0,1,32,4,0,0,0,0,1,5,1,0,3,2,0,0,3,1,0,12,3,0,0,0,1,0,0,8,0,11,1,23,17,5,0,2,0,1,4,7,3,1,2,4,14,0,0,5,0,2,5,0,0,2,0,0,1,3,1,2,0,0,0,7,1,0,6,0,0,2,0,0,6,7,1,0,0,1,0,5,7,0,2,2,1,0,0,1,1,0,1,4,0,1,0,0,2,6,0,0,0,1,0,1,1,0,0,2,3,0,0,3,0,0,1,0,0,3,1,0,0,0,0,1,0,1,1,0,0,1,0,5,1,0,2,0,0,0,5,0,0,4,2,0,1,0,0,1,4,1)
VarExplained_Protein_Age=c(1,0,0,1,0,0,2,0,7,0,1,1,5,1,0,0,5,0,2,1,0,0,2,0,1,0,7,3,0,2,0,0,1,1,0,0,6,4,0,0,1,0,0,1,0,0,2,0,0,0,6,1,1,1,1,2,0,0,1,0,0,0,0,1,0,0,2,4,1,11,0,1,0,1,1,2,0,9,2,4,6,1,0,2,1,1,1,0,1,3,0,3,4,5,0,0,2,0,0,3,4,0,0,1,0,0,2,8,1,2,2,8,2,1,2,1,7,4,0,3,7,0,16,4,0,8,3,3,0,5,5,0,0,15,12,1,2,12,6,3,0,0,0,0,1,3,3,0,1,4,7,4,2,4,2,0,3,1,2,0,0,5,7,2,4,1,4,7,5,0,1,0,1,1,5,3,3,2,2,0,7,0,8,2,1,0,0,0,4,3,9,7,0,0,5,9,0,1,3,2,0,0,0,0,0,0,4,1,4,4,2,0,7,2,11,2,0,0,1,2,13,0,6,2,1,0,2,1,0,1,2,0,0,1,0,0,1,0,0,2,0,3,0,0,4,1,1,0,1,1,2,0,0,1,3,1,0,0,6,2,17,3,0,5,1,16,2,4,1,0,1,2,8,0,3,0,4,0,3,5,2,2,0,0,2,0,1,6,1,9,3,8,2,1,3,6,2,1,0,1,1,11,3,0,0,2,0,18,2,6,5,0,0,0,2,0,3,1,4,5,1,1,3,1,1,0,0,6,9,1,1,1,2,7,1,1,0,0,0,0,1,0,4,2,0,2,1,1,0,0,1,0,1,0,8,0,0,1,3,5,2,3,0,8,1,0,12,4,2,0,0,0,5,0,0,1,4,1,1,6,0,0,0,6,0,0,0,5,0,6,1,16,7,2,2,2,0,0,2,2,3,1,8,2,0,2,0,0,4,3,4,6,2,0,2,1,5,4,1,2,2,0,1,0,1,0,1,0,1,2,5,1,0,0,0,4,6,3,2,0,2,2,0,3,0,1,0,1,1,1,1,0,0,2,0,9,2,2,0,1,2,1,5,0,6,1,3,0,1,1,1,10,0,1,0,1,0,0,2,1,2,0,1,1,0,3,0,0,0,1,0,0,1,2,0,1,1,0,5,0,2,1,2,0,0,0,6,2,4,0,2,0,0,0,1,2,7,1,1,13,1,0,0,0,2,0,2,0,0,0,0,0,5,3,4,0,0,4,3,0,2,1,4,0,7,0,1,3,0,3,5,0,1,0,0,1,0,0,0,1,2,0,11,0,1,1,0,0,4,5,7,0,3,4,1,0,4,0,0,1,0,0,3,0,10,3,1,1,1,1,0,2,2,0,4,0,0,3,1,2,2,0,1,0,0,2,0,4,0,3,2,0,7,1,0,0,2,3,1,2,1,0,0,1,0,0,2,2,4,0,2,0,1,0,0,5,11,4,0,3,0,2,0,0,1,0,0,5,6,3,4,13,3,1,2,6,0,7,1,1,0,2,3,7,11,0,0,7,1,7,1,1,1,5,2,1,0,8,3,5,1,1,1,2,0,10,2,2,0,2,0,2,5,6,1,11,2,5,5,1,1,4,0,0,0,3,0,4,0,0,6,2,3,2,7,0,0,9,0,1,4,16,1,1,8,3,0,1,7,0,1,11,0,1,3,0,7,3,4,1,2,0,2,1,1,0,5,0,2,4,0,0,1,4,1,0,5,0,3,1,4,6,6,5,5,16,2,0,2,1,4,0,0,2,5,15,1,3,0,1,0,0,5,1,0,0,1,1,3,0,1,0,4,0,1,0,3,14,0,1,6,0,5,0,1,2,1,0,18,1,5,1,1,0,0,6,1,1,10,3,0,0,1,0,0,3,0,1,9,4,0,3,7,0,0,8,4,0,11,2,1,0,0,2,0,0,5,7,1,2,3,5,0,2,1,0,1,2,1,0,3,3,0,0,0,1,0,2,0,0,0,15,0,6,0,0,3,0,0,0,2,0,0,1,0,3,4,0,1,7,0,1,0,6,7,1,5,0,0,0,0,3,0,0,2,0,1,0,6,0,1,0,0,6,3,1,1,0,4,0,1,0,0,3,1,5,0,6,0,0,0,1,1,2,0,0,6,1,2,2,3,0,1,0,0,0,3,0,0,0,0,3,4,1,5,1,1,0,2,1,4,0,0,4,1,0,0,5,1,2,1,3,1,0,3,2,0,0,0,0,1,2,0,0,2,0,0,1,4,0,6,0,4,1,0,4,1,1,1,0,2,1,9,1,0,1,0,3,1,0,4,0,6,2,0,3,3,0,0,7,6,0,0,0,3,0,0,1,0,1,4,5,3,16,2,9,3,7,0,0,5,2,0,0,0,5,0,0,0,0,1,0,4,0,0,0,6,0,1,0,0,4,3,3,7,1,1,0,7,8,1,2,2,8,6,1,0,3,1,2,0,3,0,6,0,0,3,0,2,5,1,2,1,0,0,1,6,0,0,0,0,0,0,1,0,5,0,1,7,7,0,1,2,0,8,0,1,1,1,0,0,11,0,1,3,1,2,0,3,0,0,0,3,1,4,2,0,1,1,2,0,1,7,0,0,3,1,0,3,8,2,1,3,1,2,2,0,0,1,7,0,2,0,6,1,0,0,0,6,1,3,0,5,3,1,0,3,15,0,0,1,0,6,1,0,6,4,0,1,1,6,1,1,3,1,0,0,0,4,0,3,2,0,2,0,0,7,3,3,3,0,0,2,0,0,0,1,0,10,0,5,0,6,0,0,0,0,2,6,0,1,0,1,8,4,0,1,0,3,4,0,1,0,1,0,0,0,0,4,1,0,0,0,13,0,4,4,5,0,2,2,2,0,7,0,5,3,1,1,1,1,2,0,0,0,1,1,0,0,1,0,0,2,0,0,2,1,0,2,1,7,3,1,1,0,0,2,0,0,2,0,3,6,0,2,0,1,0,0,1,0,1,0,2,1,1,1,1,0,0,0,0,1,9,0,0,2,1,0,0,1,0,0,3,3,2,1,2,1,0,0,10,2,9,11,3,0,0,3,1,0,3,0,7,2,3,1,3,4,3,2,8,3,1,3,0,1,1,3,0,0,8,0,0,2,2,0,1,0,7,1,1,1,0,1,3,1,0,0,7,0,0,1,3,0,1,1,3,0,4,1,0,0,3,3,14,3,7,3,6,0,2,2,0,0,1,0,3,4,0,1,0,2,0,0,0,0,2,9,8,1,7,0,0,0,7,0,0,0,0,3,1,0,0,1,1,0,2,0,0,0,1,0,2,0,1,0,0,0,1,5,0,0,2,0,4,4,0,14,0,6,10,1,1,2,1,2,4,0,4,1,6,0,0,0,2,0,4,2,1,1,0,0,2,4,0,0,0,0,0,3,1,2,0,0,1,1,2,0,4,0,0,9,0,0,0,0,1,0,1,4,2,5,1,0,1,0,3,1,4,6,0,2,0,2,0,0,4,12,1,2,9,0,5,1,1,5,0,0,2,7,0,1,0,13,14,1,5,0,0,3,0,0,5,0,0,0,1,1,7,1,1,0,1,3,0,0,11,3,4,2,0,0,0,10,0,0,3,1,1,1,2,2,3,6,0,4,3,4,0,1,1,6,1,1,0,1,0,1,0,1,6,1,4,0,1,17,0,0,0,0,0,1,11,0,1,9,0,0,5,0,0,0,2,0,4,0,0,3,0,4,0,0,0,0,0,4,0,1,0,0,1,1,2,0,6,0,9,0,16,0,1,4,6,1,1,1,2,6,0,5,5,0,0,4,1,1,1,6,1,0,0,0,2,0,0,4,1,2,0,1,1,9,1,0,1,0,6,2,2,5,0,0,1,0,0,0,0,0,1,4,0,2,0,13,0,0,1,0,2,0,2,1,2,3,0,0,0,0,1,8,6,0,7,0,1,0,1,0,2,0,0,0,1,4,0,0,3,0,0,0,1,0,2,2,0,0,0,0,0,3,3,2,2,0,0,2,5,1,3,0,3,0,1,12,2,3,2,0,0,13,3,1,1,3,0,2,5,0,1,13,0,3,0,0,1,0,1,1,0,0,0,1,1,4,6,0,0,1,5,0,0,2,2,0,4,0,0,0,1,0,0,0,1,0,0,0,1,5,0,2,0,1,2,1,2,4,2,0,0,2,13,2,2,1,2,0,2,4,0,2,4,10,1,1,10,2,0,0,1,1,1,0,0,7,0,1,1,0,1,4,0,0,1,5,0,3,1,0,2,2,0,0,2,0,2,0,0,0,11,0,0,0,0,1,0,3,0,2,0,1,0,4,0,0,0,0,7,4,0,0,0,0,1,1,1,0,1,0,0,0,1,0,0,3,2,0,1,5,0,1,2,2,1,4,0,3,1,1,1,0,1,0,0,3,0,3,0,0,8,0,1,0,2,1,1,0,0,0,3,6,0,0,1,0,2,0,5,0,1,0,0,0,0,0,0,0,0,0,3,4,2,3,0,7,1,4,0,3,2,5,12,0,1,1,0,0,0,0,1,0,1,0,1,1,1,0,0,0,0,5,5,0,1,1,0,3,3,0,1,0,10,11,0,0,1,1,0,2,0,1,5,0,2,2,1,0,0,0,2,0,8,1,0,0,1,1,1,2,0,1,1,0,1,1,1,0,0,8,0,1,0,0,2,0,0,0,13,0,7,0,1,0,1,0,0,2,3,6,2,3,1,4,0,0,3,1,1,1,0,1,1,4,0,2,1,0,1,3,0,0,4,1,0,1,0,0,2,0,0,1,1,0,0,0,4,0,1,5,0,1,0,0,8,2,0,0,1,0,0,1,3,3,3,2,0,0,1,0,1,0,4,5,0,0,0,0,2,0,1,4,0,3,4,0,5,0,8,0,0,1,0,1,1,2,1,8,0,1,1,0,1,4,0,0,0,1,0,1,2,0,2,1,2,0,3,0,2,1,0,2,0,4,1,0,3,2,2,0,0,0,2,1,4,1,0,1,7,0,2,1,0,6,7,0,0,0,12,1,1,0,2,1,2,0,1,4,6,2,0,5,0,0,3,1,0,0,3,0,1,0,1,1,0,0,5,1,1,0,1,0,0,2,0,0,2,2,0,5,0,4,0,0,2,5,1,1,2,0,2,0,3,0,0,2,1,1,1,0,7,1,0,1,2,2,0,0,0,1,0,0,1,1,2,0,1,2,5,0,0,0,4,0,8,0,1,1,0,0,1,0,1,0,0,4,0,0,0,3,0,0,0,2,0,1,1,1,2,2,2,3,0,0,4,3,1,1,0,0,2,6,2,5,0,17,1,2,1,0,0,0,2,0,1,1,0,2,13,0,1,1,0,0,0,4,0,0,1,0,2,1,1,2,0,8,1,0,0,2,0,5,0,0,0,0,0,1,0,0,0,7,0,2,24,3,9,0,6,1,0,0,0,3,0,1,1,5,1,2,3,2,3,2,1,0,0,0,0,0,2,6,0,0,0,1,0,0,1,0,1,1,1,3,1,1,4,1,2,2,1,0,2,0,0,0,4,1,0,5,10,3,0,14,0,12,0,0,4,1,1,0,0,0,0,2,1,2,0,12,4,4,1,2,1,4,8,1,0,0,0,1,0,2,0,2,0,0,1,0,2,0,0,1,0,0,2,1,0,0,0,2,0,4,0,1,0,0,0,1,0,0,21,8,7,3,0,6,0,0,1,2,1,0,3,5,1,0,3,0,0,1,6,0,9,0,0,1,0,0,5,3,1,1,1,0,0,8,0,0,0,5,0,1,1,1,3,1,2,0,0,0,5,1,2,0,0,4,2,0,1,2,0,4,2,1,3,2,3,0,6,0,1,2,0,0,0,6,0,0,1,2,3,8,0,0,2,1,6,4,1,1,2,1,0,1,0,0,0,0,0,0,0,0,0,0,0,4,1,1,6,0,2,0,0,2,0,7,2,12,0,2,1,1,1,0,2,0,0,3,1,3,0,0,0,1,1,1,0,10,0,2,4,3,1,0,0,4,3,0,3,4,1,0,0,0,3,9,1,0,1,2,0,0,0,1,0,0,1,0,2,2,5,3,8,2,0,1,3,12,1,0,0,0,2,0,3,0,1,0,12,5,3,5,0,0,4,9,0,1,2,0,1,5,1,1,9,0,1,2,3,0,0,2,0,0,0,2,9,3,4,0,2,1,6,1,1,0,0,2,1,11,0,0,2,1,0,0,15,0,0,0,0,1,0,0,0,0,0,2,2,9,3,1,2,0,0,1,0,12,1,1,0,0,1,1,2,1,0,0,0,1,1,1,0,0,0,1,2,10,0,0,2,18,0,0,1,0,0,0,0,1,0,0,2,0,0,10,0,6,2,0,2,4,0,0,5,1,1,4,7,0,0,1,1,0,2,0,0,0,0,1,3,2,0,2,0,0,2,0,0,0,2,0,1,2,0,3,0,0,1,0,0,0,2,3,16,3,0,4,0,0,0,0,0,5,1,0,10,0,0,1,0,2,0,0,0,5,0,0,0,0,1,0,2,0,13,0,0,2,0,0,6,2,0,0,4,1,1,0,0,3,0,0,1,5,3,0,0,0,0,0,0,1,0,0,0,1,0,1,4,0,0,3,0,2,0,1,2,0,0,0,1,1,1,1,2,1,2,0,2,0,0,2,1,0,0,3,3,1,1,0,1,12,1,2,9,1,1,4,1,3,1,1,0,3,0,5,5,0,1,2,2,0,5,0,0,3,1,0,1,0,0,0,2,3,3,0,0,0,1,1,4,2,0,3,2,1,5,1,1,0,1,3,0,2,4,0,1,0,0,0,0,0,6,5,2,20,1,13,1,1,1,0,0,1,0,0,3,0,2,6,0,0,1,3,1,0,0,1,0,1,0,0,0,0,0,0,4,1,1,0,1,1,0,0,0,10,1,0,5,2,1,9,2,1,13,7,6,3,4,0,1,1,0,0,0,2,10,0,0,7,9,0,1,0,0,1,4,2,5,0,2,0,0,1,0,0,0,1,1,0,1,0,2,1,0,3,3,0,0,5,1,0,3,4,13,0,1,0,1,8,6,1,1,3,0,0,4,0,1,1,5,3,0,6,0,1,0,0,0,4,1,0,0,0,0,2,2,0,5,1,0,0,1,4,0,4,0,1,2,0,0,0,2,0,0,7,1,1,9,1,1,7,2,0,1,0,0,3,1,7,1,3,6,0,3,0,1,1,0,7,0,2,1,3,0,20,0,7,0,0,0,0,2,3,1,0,1,1,0,3,0,0,4,0,0,2,0,0,1,0,1,0,0,1,3,1,1,2,1,1,6,3,0,0,0,0,2,0,7,0,0,1,0,3,1,0,0,1,2,3,5,0,8,0,0,0,2,3,5,0,0,0,0,3,2,0,8,1,0,0,0,0,0,4,5,1,19,0,1,5,0,2,0,0,1,0,0,1,3,0,2,0,0,1,0,0,0,0,0,2,4,1,1,0,3,0,3,0,0,8,0,0,1,8,0,0,0,0,1,5,1,0,0,0,2,1,1,0,0,4,0,2,0,2,0,1,0,16,3,0,0,0,0,1,1,1,2,0,0,1,2,2,3,0,0,0,0,1,6,0,0,3,0,2,0,9,5,2,2,2,5,0,4,0,1,1,0,1,1,5,0,2,2,0,0,0,0,1,0,2,24,0,2,2,2,4,1,6,0,2,5,2,0,0,3,1,1,2,12,3,4,1,0,0,4,3,1,2,0,1,4,0,3,4,1,1,0,1,0,5,6,1,3,2,5,1,0,0,1,0,0,1,0,0,0,0,0,9,1,2,0,0,0,0,0,1,1,0,0,4,0,0,0,3,0,0,1,4,1,1,0,1,7,0,0,1,6,0,0,2,0,1,1,0,0,0,8,0,1,2,5,1,4,3,4,2,0,0,0,1,1,0,2,0,0,4,0,1,4,0,0,0,1,1,3,1,1,1,0,0,0,5,3,2,0,1,0,0,0,0,0,0,1,2,2,1,8,0,2,0,0,0,2,1,18,4,3,7,1,0,0,2,0,0,1,1,7,0,16,0,0,1,8,0,0,5,3,3,1,5,1,1,0,1,0,2,0,2,10,2,0,0,0,5,3,0,1,2,0,0,2,1,3,2,0,0,1,0,1,2,0,2,0,1,1,5,1,0,1,1,0,7,3,12,3,0,0,0,0,0,1,1,0,0,0,0,2,0,2,10,3,0,0,2,6,1,1,3,0,3,2,2,0,1,1,1,3,1,1,7,3,2,0,1,2,2,4,0,0,3,14,0,2,0,1,0,1,0,3,2,1,1,4,2,0,1,2,2,0,0,7,1,0,2,0,0,3,3,0,0,2,2,0,0,0,7,1,1,0,1,0,2,2,2,0,0,2,0,3,1,2,0,12,2,0,0,0,0,0,1,0,0,1,0,3,2,1,0,5,0,0,0,12,0,1,1,0,0,0,4,0,1,0,1,0,16,0,0,0,0,0,0,0,0,9,6,0,0,3,3,0,0,0,2,0,0,0,3,1,1,0)
VarExplained_Protein_Interaction=c(21,32,13,31,31,15,21,35,15,32,21,25,18,24,17,36,36,21,18,38,27,26,18,22,28,36,41,28,30,23,39,25,29,24,41,23,30,33,37,18,29,34,37,30,34,39,33,20,37,41,33,26,31,27,27,33,22,17,40,32,30,25,33,30,29,35,35,32,18,35,12,40,37,36,33,21,44,26,26,35,27,33,30,28,41,40,42,33,24,35,31,39,38,24,33,33,26,41,26,19,35,30,31,37,28,41,32,29,33,42,38,33,21,18,25,33,30,31,38,26,30,27,26,38,30,31,33,30,33,37,37,37,39,28,30,21,32,24,33,23,43,34,20,33,23,23,33,38,22,34,25,37,35,39,26,42,42,38,22,38,43,21,29,41,26,31,28,35,26,37,41,36,37,36,36,40,35,38,32,37,20,24,28,27,35,38,32,26,20,35,30,34,41,27,34,33,39,42,35,33,34,39,19,34,34,26,38,37,32,32,27,20,35,29,18,20,42,26,41,37,37,39,33,43,23,38,34,41,47,24,23,38,37,31,32,22,32,35,30,31,27,39,39,26,32,44,30,8,43,40,23,21,40,24,28,38,18,39,31,30,29,28,33,31,35,29,35,38,32,33,28,27,32,35,35,37,33,26,39,30,25,35,38,42,22,35,28,34,43,32,34,31,35,34,33,24,18,38,35,42,36,28,33,26,35,34,38,28,32,33,38,34,23,34,33,40,39,31,26,40,34,39,38,17,36,41,15,34,28,41,33,23,38,28,33,39,29,36,13,46,39,29,28,36,31,29,33,23,43,22,28,21,36,36,35,28,38,34,38,36,37,24,34,28,21,40,34,46,43,35,30,38,32,13,31,37,37,41,32,36,38,38,38,28,24,39,47,36,34,28,38,31,34,39,34,26,34,40,29,33,40,24,28,41,22,26,27,40,36,32,36,38,29,30,40,42,36,25,42,33,34,33,30,24,38,37,41,35,41,43,34,43,36,36,21,31,35,40,35,37,26,35,29,35,23,32,36,33,44,39,31,27,39,36,32,26,45,36,39,20,35,41,37,26,33,31,34,32,38,34,34,39,15,35,38,35,36,34,38,31,27,36,32,37,26,39,37,37,25,33,43,36,38,34,38,28,32,43,31,40,39,26,41,28,42,37,38,43,30,26,36,37,41,27,33,33,37,32,24,23,31,39,48,31,41,37,37,36,40,37,40,39,24,37,41,35,32,35,26,45,40,35,36,29,24,41,41,37,11,31,35,36,36,37,34,43,17,38,43,34,32,27,33,42,35,37,36,34,35,34,33,42,36,39,38,39,34,41,32,35,42,35,34,28,33,41,24,28,26,29,4,34,27,36,37,33,40,26,35,30,32,38,31,43,39,33,33,38,38,34,37,38,35,30,32,39,31,25,35,21,37,43,36,44,30,43,36,36,33,46,43,34,30,36,31,30,30,41,42,35,27,26,43,36,25,38,41,36,29,22,28,32,32,35,39,31,35,26,35,37,34,27,39,29,29,39,33,32,26,35,37,34,34,21,36,42,33,32,36,38,35,38,36,38,33,35,41,42,48,28,38,43,27,25,31,32,31,39,41,27,30,31,40,36,35,43,37,37,30,29,35,24,36,34,37,39,38,29,36,28,24,20,34,28,33,40,40,40,38,36,35,28,31,44,36,39,31,26,31,29,26,42,26,44,29,36,37,45,20,30,32,25,21,28,39,34,29,38,37,33,33,30,28,34,42,33,38,39,35,38,31,36,41,32,31,22,32,29,27,36,54,36,33,28,28,43,36,34,34,39,44,27,34,30,38,36,46,21,38,38,22,33,31,38,36,36,29,39,33,25,29,39,33,34,36,31,40,40,30,28,37,36,21,30,32,36,36,30,29,36,39,37,33,29,35,33,37,35,28,42,39,35,43,39,33,38,30,20,39,38,31,35,37,32,36,36,37,32,40,35,28,27,43,39,37,38,38,37,41,41,46,32,38,26,39,39,33,45,33,37,37,45,35,39,34,28,33,29,29,34,34,36,33,35,36,44,26,36,37,34,38,28,37,38,41,33,26,34,35,40,37,37,36,37,39,40,34,40,31,31,39,34,35,36,36,28,30,36,33,36,41,29,33,40,40,41,26,27,34,29,25,33,31,31,40,22,37,38,29,34,36,38,32,23,35,42,35,33,39,19,31,10,31,26,45,27,37,40,33,39,36,32,34,33,36,33,34,38,38,31,25,28,43,37,32,23,36,30,37,39,26,24,19,35,37,32,30,14,41,27,32,34,35,35,35,38,33,33,31,24,41,26,26,49,39,48,37,36,33,30,32,35,29,39,36,29,41,37,35,41,36,34,27,26,30,29,26,38,42,29,29,38,36,9,39,32,33,20,32,40,33,38,36,26,40,38,28,37,21,26,30,38,27,34,40,29,28,43,37,37,26,41,35,27,37,30,36,37,32,34,28,44,31,27,35,31,41,30,36,34,31,47,39,34,36,40,28,36,39,16,28,38,31,29,37,32,28,35,35,27,34,34,35,29,32,38,43,27,30,40,37,27,25,29,33,31,41,42,35,46,45,40,34,38,26,34,33,35,39,45,38,44,30,28,40,31,29,16,29,37,39,41,39,19,20,29,34,41,41,26,26,21,30,28,36,32,36,42,31,40,37,38,32,35,39,29,38,29,36,41,26,37,31,30,36,34,36,41,32,38,32,38,42,27,29,28,36,39,38,33,37,33,40,31,17,38,33,35,34,34,36,44,39,37,43,40,34,38,26,37,34,44,40,42,33,45,31,32,38,28,38,40,32,26,33,26,34,26,38,33,41,35,39,40,24,37,36,16,30,13,35,39,38,35,44,34,34,37,35,28,40,30,40,32,31,27,32,35,35,41,39,30,31,32,28,34,25,43,40,31,41,37,28,32,41,43,30,37,41,29,34,37,36,44,45,35,39,20,32,36,24,40,35,42,35,33,34,35,40,40,29,22,21,28,30,17,40,35,28,43,35,39,35,33,32,39,26,36,27,31,31,46,45,27,32,34,32,33,39,42,38,31,41,23,34,34,35,32,35,41,40,40,34,27,39,25,33,30,32,21,39,32,29,38,45,48,37,41,31,37,44,24,31,30,48,32,26,44,51,28,35,40,42,34,40,39,41,32,38,31,28,35,43,37,34,42,39,28,40,25,10,37,20,32,37,33,41,28,30,30,42,28,38,37,45,36,35,34,34,38,37,41,36,36,39,40,38,34,30,31,37,17,37,32,37,26,40,39,38,31,31,38,36,33,33,30,36,19,43,41,40,31,38,31,41,37,26,42,43,38,30,36,36,33,38,37,38,35,33,36,30,33,33,30,33,37,40,38,32,31,38,31,37,34,37,35,38,33,35,42,34,42,31,40,39,35,24,38,33,38,28,33,36,39,31,31,39,41,39,40,41,40,37,40,32,43,38,33,37,22,27,19,31,42,39,38,38,33,36,23,29,43,39,29,39,36,23,32,29,36,40,27,37,37,27,38,36,37,36,36,31,13,29,43,28,27,34,33,37,44,34,35,31,33,28,30,37,32,29,30,33,35,36,42,34,33,31,22,30,35,37,41,34,39,36,38,36,20,26,34,34,19,30,39,34,36,36,31,31,35,23,30,30,33,37,38,34,35,22,27,43,31,24,34,32,35,29,27,9,40,36,39,38,29,32,33,31,21,40,29,43,36,41,39,45,38,36,36,34,35,32,33,42,32,38,32,37,38,28,33,31,36,38,45,36,42,39,27,41,28,34,38,31,29,43,35,32,34,32,26,35,22,40,36,39,41,23,36,35,27,29,37,26,11,43,32,39,39,35,40,19,23,21,40,38,35,6,30,40,38,36,27,27,35,32,23,41,22,35,28,36,39,34,29,35,15,27,14,45,30,38,33,37,30,29,21,42,37,42,39,34,33,39,36,41,43,42,37,42,38,39,23,38,44,35,38,35,31,30,29,38,37,41,33,28,44,37,42,31,36,37,33,32,37,28,35,32,33,35,32,29,39,42,33,25,30,25,30,41,32,31,30,38,39,40,34,39,26,33,33,27,41,43,34,24,38,36,25,33,30,38,37,38,40,37,35,33,39,31,33,33,38,31,35,28,38,36,35,42,35,36,36,30,33,42,33,30,34,29,31,36,34,37,38,34,36,36,37,33,42,34,36,32,38,23,37,35,33,29,38,35,44,25,39,28,33,32,36,41,35,36,31,27,30,33,38,41,40,40,22,33,40,26,33,30,35,35,39,32,39,39,39,40,37,40,36,32,37,41,33,36,7,37,36,41,7,37,33,44,34,16,32,38,33,43,7,24,29,34,34,32,31,21,38,28,44,28,36,39,22,16,31,30,41,26,34,33,32,34,40,40,30,40,39,36,38,34,34,38,36,30,38,28,37,34,41,35,33,40,24,36,33,47,31,38,37,38,43,35,35,38,45,21,32,41,34,44,31,38,33,36,37,25,35,30,26,26,33,23,40,40,35,36,33,35,37,21,32,37,36,30,29,41,39,34,33,30,34,42,13,28,30,37,41,38,35,33,34,32,34,19,38,29,23,25,34,33,35,39,35,26,46,22,34,33,37,30,27,28,33,32,30,31,20,36,30,39,35,39,34,35,26,27,36,42,39,32,35,39,30,40,34,33,39,49,36,37,11,39,25,43,34,40,34,25,40,23,32,38,38,37,38,31,28,37,35,38,38,38,40,40,37,46,31,34,39,24,39,46,33,40,41,41,42,37,30,35,22,30,40,33,30,42,18,19,33,40,27,27,39,44,31,38,27,36,41,40,30,33,45,38,43,30,34,37,38,39,36,40,34,25,37,25,27,37,37,33,28,36,37,43,33,38,40,40,37,30,39,33,48,38,32,29,30,37,35,42,39,41,7,36,39,19,24,32,38,33,41,38,31,36,35,33,32,38,36,40,23,31,34,37,39,39,34,42,36,40,35,29,38,34,41,33,33,35,34,35,35,39,12,32,48,38,30,39,37,36,26,39,27,40,12,24,37,37,34,39,27,23,40,36,45,40,23,29,34,32,28,24,29,34,7,40,34,39,41,42,30,31,30,24,29,24,11,36,30,33,43,11,39,46,37,35,38,38,33,17,40,35,40,41,32,32,30,35,34,39,30,41,36,30,31,32,38,33,22,41,43,29,37,34,39,43,27,37,39,28,46,29,35,38,33,42,25,39,26,39,32,38,39,31,26,28,33,31,32,28,32,34,34,25,43,14,26,39,40,38,33,38,39,40,38,20,35,32,33,37,27,21,39,30,36,34,31,34,43,39,35,37,36,37,32,31,32,28,40,37,27,36,35,32,41,34,36,26,42,28,26,46,26,22,39,30,40,33,32,32,23,33,30,33,40,37,39,32,35,33,37,42,38,39,38,28,39,23,22,34,37,41,38,24,36,34,26,27,38,36,35,32,32,35,29,34,37,32,42,27,40,38,35,35,35,31,25,38,34,39,47,39,37,37,33,42,35,39,41,38,43,40,26,24,38,24,34,41,36,44,36,38,37,32,36,25,39,32,36,36,39,30,39,34,31,21,42,33,33,35,36,40,36,39,7,35,28,30,19,40,34,20,37,37,35,21,40,35,35,33,26,32,40,28,37,41,34,18,40,39,22,33,35,25,37,41,36,37,37,35,37,38,40,32,31,27,39,32,21,31,34,39,37,37,26,32,29,34,26,41,38,34,36,33,40,28,38,40,39,31,31,32,42,38,31,37,33,35,34,38,34,26,37,31,35,39,26,32,37,30,35,38,39,38,30,32,31,35,29,44,38,43,31,36,34,45,37,39,36,39,40,37,32,32,33,39,30,33,39,39,34,39,25,32,27,39,24,40,39,37,37,42,36,39,40,32,40,27,32,33,37,35,25,38,33,44,41,38,31,35,45,35,41,8,39,37,42,37,36,42,44,25,35,39,30,40,40,32,32,27,33,30,42,29,32,35,38,28,37,47,38,38,35,34,43,33,39,34,28,33,38,30,25,36,28,39,33,22,32,33,36,32,29,28,34,40,11,41,46,35,40,36,36,27,17,42,35,33,32,45,40,36,24,39,36,36,38,36,35,41,29,32,43,30,38,39,31,34,36,40,39,41,35,33,29,29,40,43,35,36,26,34,28,39,37,29,30,42,29,36,31,40,35,38,39,31,27,43,33,35,32,32,21,39,39,39,33,44,42,13,25,40,35,33,33,37,32,34,40,25,35,31,41,29,27,39,38,47,38,37,37,42,33,29,34,34,28,23,34,29,36,34,33,47,34,29,43,38,31,27,38,38,30,29,29,34,45,31,31,35,37,35,28,41,32,27,33,44,32,31,37,37,38,40,36,35,27,33,33,34,31,31,35,38,41,33,30,31,43,39,29,35,38,40,23,41,30,42,38,42,29,43,38,41,33,31,34,40,42,32,36,34,32,46,36,37,44,32,29,37,39,25,41,36,42,29,34,32,34,34,43,31,38,14,45,38,35,35,28,31,26,38,40,35,41,37,30,36,35,38,38,30,38,35,25,37,31,31,27,34,34,38,34,35,36,27,27,40,27,40,40,30,5,33,18,32,36,31,40,37,40,32,35,44,28,41,34,32,40,36,29,32,27,44,41,25,40,34,39,34,33,33,39,32,30,41,26,34,36,42,26,39,42,38,40,30,35,33,29,32,41,35,35,45,37,36,40,27,29,35,42,39,42,33,39,21,38,24,25,31,44,33,33,35,34,14,30,39,39,29,42,39,19,38,28,38,41,38,40,37,41,37,42,37,33,33,38,42,29,28,26,36,34,33,37,41,38,33,23,38,33,27,30,35,27,30,33,32,39,26,42,31,32,43,39,39,36,32,46,39,25,35,37,44,42,25,32,30,36,33,23,36,25,24,39,22,31,40,39,36,26,43,36,25,36,35,41,34,26,30,35,42,19,39,23,33,24,33,34,38,32,36,36,28,39,36,42,41,38,31,28,36,35,30,32,44,50,40,36,41,27,36,41,37,36,36,35,37,36,35,37,38,37,38,31,45,37,36,33,37,37,28,33,33,37,41,38,31,32,31,40,38,41,30,25,32,37,33,34,35,37,36,32,26,36,34,36,37,37,39,39,42,19,37,40,36,41,42,44,21,34,36,27,39,31,35,24,37,35,38,12,29,40,36,36,36,35,41,37,28,33,36,35,35,35,29,37,35,45,36,35,33,35,32,37,37,37,37,40,38,49,39,46,35,30,31,28,22,34,21,25,32,29,33,24,39,33,35,35,39,35,45,33,28,26,42,37,29,35,38,37,37,35,41,36,38,33,41,31,43,38,41,37,32,47,28,36,42,30,42,36,35,36,35,40,36,44,30,23,29,38,33,30,36,28,37,35,32,39,38,39,34,36,33,31,27,36,34,37,24,24,34,43,38,38,26,42,31,32,30,31,41,33,39,41,41,39,42,30,38,31,29,37,34,11,43,38,39,43,41,44,43,39,41,32,35,36,29,38,35,32,38,34,32,37,37,34,32,38,37,45,39,18,37,22,33,16,41,33,36,41,35,30,23,35,37,36,24,30,33,7,30,35,11,36,37,38,35,11,38,38,38,34,35,34,40,38,39,40,36,29,28,34,32,28,25,36,38,35,31,39,28,32,40,37,35,34,46,33,41,40,39,42,34,45,31,39,40,35,40,40,45,29,32,22,36,38,32,38,29,32,31,37,22,28,26,38,36,18,38,42,37,32,42,34,41,35,32,34,33,35,22,42,27,32,14,37,29,38,32,37,40,36,40,19,29,31,32,28,38,40,35,37,22,36,31,43,40,23,39,40,31,37,27,35,39,34,40,35,24,32,34,12,31,44,33,34,29,30,30,29,41,41,40,36,37,35,31,33,30,33,43,37,37,40,38,30,48,27,51,29,34,29,35,36,36,34,29,26,29,31,35,39,35,37,39,24,14,35,36,30,40,36,44,39,40,39,45,35,35,35,31,39,41,35,36,35,36,43,44,40,31,40,15,37,32,36,39,41,36,30,41,38,31,35,28,33,33,27,30,40,33,33,24,20,32,38,14,39,31,39,32,32,34,34,41,21,40,33,33,37,34,35,34,27,47,36,30,31,38,40,39,24,24,36,27,36,44,40,23,31,41,34,41,29,38,39,38,44,39,37,31,44,36,32,39,31,29,27,38,33,28,43,35,9,25,22,36,37,31,35,32,40,27,43,38,30,20,22,13,29,45,36,34,34,31,28,32,38,24,32,31,37,35,27,17,34,41,34,27,31,41,35,27,22,42,13,27,38,36,29,46,41,38,37,34,24,37,27,31,30,33,35,32,39,35,32,29,31,35,40,20,35,44,37,40,39,38,32,24,28,26,34,40,34,28,31,36,33,26,34,32,45,40,39,41,36,37,32,29,30,41,23,39,35,40,16,34,38,42,31,40,34,11,37,38,36,36,38,29,33,35,35,32,41,35,33,37,34,34,33,40,35,37,37,40,32,43,40,26,39,37,35,32,29,41,30,26,31,27,31,38,26,32,40,33,33,34,39,36,43,26,37,29,34)
VarExplained_Protein_Residual=c(16,26,14,32,26,17,28,34,19,34,26,29,17,26,19,37,28,21,25,35,32,17,18,27,19,36,35,25,36,32,35,30,39,13,40,27,34,39,46,22,26,23,43,37,47,44,42,20,36,34,33,25,38,26,25,31,17,23,36,40,35,31,33,25,42,42,40,42,15,30,8,30,35,40,45,31,32,31,22,43,36,29,24,27,33,32,37,27,14,28,33,33,37,30,26,50,29,34,36,21,36,29,41,35,30,35,40,41,43,25,38,38,24,21,20,26,25,40,31,37,26,27,35,31,46,25,33,24,32,37,34,37,46,26,37,30,44,38,31,24,40,37,21,28,17,30,37,41,23,30,28,39,36,31,34,31,29,42,24,41,44,30,30,42,31,41,37,32,31,37,37,42,37,38,35,28,41,39,33,45,24,27,34,35,34,36,46,43,31,35,34,36,37,37,36,34,39,39,38,42,38,39,20,45,38,17,36,44,40,31,36,26,35,30,21,16,35,35,42,40,21,46,20,32,29,33,49,35,35,24,18,43,42,28,35,20,45,45,38,35,45,33,44,27,42,35,36,9,37,40,30,17,40,21,25,34,22,33,35,33,28,32,38,32,30,26,40,43,48,39,30,23,35,46,34,45,45,34,25,37,28,44,33,38,21,27,41,41,33,23,31,35,46,37,31,30,28,44,40,32,39,31,30,46,31,41,41,32,42,31,33,38,25,36,41,32,31,39,25,26,45,34,26,17,30,33,14,31,33,35,36,20,32,28,32,38,39,30,14,29,37,33,43,45,41,29,38,39,35,22,26,23,45,43,35,26,45,38,31,31,26,26,49,35,26,42,34,27,29,41,47,42,43,14,48,46,33,37,44,22,45,40,34,28,25,35,25,23,42,36,39,31,36,40,31,20,36,36,29,38,28,26,40,29,29,27,31,39,27,43,30,33,30,29,42,42,22,26,33,40,40,33,23,28,32,41,38,43,39,29,43,35,39,32,20,41,31,28,46,32,29,39,31,35,24,38,30,45,34,32,29,32,40,35,31,40,33,33,41,43,33,40,30,30,30,44,31,46,39,34,34,32,19,22,36,38,43,36,36,32,44,45,37,39,46,38,35,41,28,27,35,39,43,35,43,34,42,34,29,39,35,37,28,36,41,44,41,33,30,35,38,47,39,29,38,38,22,30,31,29,41,42,25,38,29,41,35,46,39,41,30,41,37,28,37,37,40,40,21,31,30,45,29,30,26,40,36,30,10,38,33,37,38,45,29,36,24,43,33,40,47,24,31,37,40,36,28,34,35,39,43,39,38,32,39,42,32,34,45,43,42,37,35,31,34,39,13,29,33,47,5,31,24,43,44,35,43,26,47,31,21,44,40,37,40,34,48,23,44,43,41,48,42,26,46,39,46,24,31,24,34,38,35,32,25,40,39,34,39,37,37,37,49,43,28,23,30,40,39,39,33,25,35,45,45,43,39,30,25,23,34,32,39,38,39,33,40,28,43,40,41,42,36,36,32,47,42,28,32,35,45,35,43,26,40,34,42,39,30,33,45,39,37,41,23,38,36,25,33,32,36,36,47,32,41,39,32,25,25,38,46,37,41,45,39,35,45,38,38,35,29,25,45,43,39,46,39,26,42,38,21,29,27,51,34,39,31,32,26,39,44,33,29,32,41,42,44,29,29,45,32,43,39,38,29,48,39,33,19,42,34,33,21,27,31,46,24,31,27,30,42,26,41,40,24,25,41,40,34,42,43,38,37,33,28,26,25,42,34,44,23,39,34,33,33,41,36,44,38,37,42,27,25,40,35,38,31,24,46,26,18,37,37,37,41,34,33,39,31,22,44,45,39,43,37,33,35,35,30,33,41,46,25,28,24,34,48,45,30,38,41,32,37,24,40,42,32,30,31,41,27,51,37,38,32,35,31,23,28,37,48,31,40,41,31,43,42,38,32,47,38,31,32,40,42,40,36,36,40,49,37,26,48,30,43,39,31,38,33,28,45,38,42,38,40,38,30,26,30,25,45,41,34,35,35,29,22,43,33,37,46,15,39,41,30,48,23,51,38,41,35,45,29,36,32,40,39,41,42,40,45,44,36,45,30,27,32,28,31,49,40,35,47,41,47,39,31,45,44,26,30,38,31,41,46,21,32,38,40,30,45,36,33,25,30,34,47,33,40,21,38,9,36,36,31,39,39,27,51,30,44,21,44,41,42,26,49,31,47,39,26,34,35,40,32,22,41,44,39,45,29,41,30,43,45,43,30,17,41,44,33,48,35,37,40,41,35,35,34,16,36,21,36,33,43,36,36,48,22,33,32,42,53,42,31,26,39,33,39,37,31,26,34,34,37,33,33,31,37,25,28,31,36,10,43,40,50,19,35,43,51,44,37,41,41,43,36,42,18,33,33,38,27,32,32,46,28,38,20,31,31,33,38,20,33,40,38,29,29,42,23,28,47,27,32,38,38,27,44,40,34,35,24,41,40,41,37,46,32,20,35,44,42,44,39,38,24,39,24,42,48,38,40,46,34,37,44,50,38,33,43,31,28,23,36,46,37,46,31,40,36,42,31,43,33,36,45,39,40,32,45,32,26,43,39,32,27,18,43,39,36,38,41,20,19,33,30,42,37,27,25,25,43,29,42,42,37,38,31,39,30,46,48,38,38,35,41,26,36,41,41,31,40,36,42,32,42,39,46,40,35,45,40,26,42,36,35,34,35,34,30,40,40,24,19,37,38,40,25,37,44,31,41,32,39,43,30,40,30,45,38,29,33,36,50,39,38,37,40,32,36,38,33,20,40,44,43,48,34,35,39,48,43,39,50,39,41,15,33,11,39,34,43,32,37,41,40,34,34,36,39,35,36,43,40,32,38,50,29,39,38,44,36,39,52,43,20,35,42,32,41,44,32,47,45,23,24,49,27,36,35,42,43,30,35,47,42,17,48,32,23,40,29,37,47,49,34,32,43,23,42,19,22,17,33,15,37,40,22,35,34,35,41,34,38,38,23,28,23,34,40,34,39,32,37,43,47,50,38,40,38,39,38,32,33,39,43,42,40,43,38,37,41,32,35,24,37,43,32,32,36,31,29,39,24,34,41,36,33,32,41,28,46,39,34,47,31,38,37,36,33,36,36,44,37,37,43,48,38,41,32,43,34,40,46,35,41,37,37,34,10,29,19,41,35,28,41,33,33,38,35,25,38,37,40,43,37,39,31,45,41,38,38,38,39,26,45,40,36,35,36,10,41,32,39,31,39,37,42,40,33,43,40,47,43,43,42,24,41,43,43,36,41,44,39,41,28,45,32,33,31,46,34,42,41,32,38,43,34,41,41,33,31,26,45,44,42,45,46,44,46,41,43,39,34,32,41,29,36,38,44,38,30,41,35,42,41,45,36,42,39,41,44,35,31,35,43,31,38,33,31,37,31,44,37,38,37,38,46,31,23,30,39,38,43,39,33,42,46,25,31,36,37,31,41,37,29,36,33,47,27,29,42,39,29,42,37,37,41,44,37,16,32,43,34,27,40,30,36,29,45,45,39,36,36,37,48,33,34,35,38,35,46,35,49,40,41,26,33,37,44,37,44,38,33,33,46,20,20,40,27,28,33,32,41,30,25,38,35,44,22,33,25,34,40,42,32,37,26,38,41,36,17,40,46,44,24,34,11,37,42,46,38,32,44,43,31,25,39,35,35,37,32,33,31,36,46,33,39,40,35,43,40,33,49,44,39,42,30,42,38,38,32,33,45,30,40,34,40,31,46,41,28,35,37,35,32,42,31,34,27,22,34,37,29,38,25,44,38,28,37,37,24,20,33,46,34,40,34,42,20,27,26,39,30,29,7,31,37,43,28,39,37,41,40,30,39,28,49,32,36,39,43,29,23,15,33,13,37,25,32,28,33,29,41,26,43,37,37,35,25,42,36,29,44,32,37,41,37,33,38,30,42,38,38,43,44,43,40,48,35,41,33,40,46,38,38,41,41,44,47,44,41,33,31,50,37,43,46,33,22,33,36,33,28,35,40,39,43,27,33,39,39,37,37,47,43,27,36,38,33,44,34,47,30,43,42,19,31,41,36,42,41,40,31,36,31,45,35,37,45,34,36,45,30,36,38,45,38,48,52,43,31,43,44,39,31,42,28,30,31,38,38,39,34,42,35,41,45,38,40,35,41,39,21,41,29,35,33,38,47,40,25,44,44,23,52,41,44,36,42,28,30,35,31,42,39,31,38,56,29,38,30,43,30,37,48,38,37,35,42,35,32,47,44,39,54,42,34,31,47,7,34,29,30,7,39,20,35,34,25,48,30,45,38,5,38,29,28,40,36,44,35,35,36,34,32,39,40,25,23,50,50,37,39,42,51,46,37,36,29,32,31,43,34,48,38,40,40,41,30,42,40,35,32,40,41,30,39,27,42,44,38,32,45,39,43,40,29,39,28,39,28,38,38,43,38,38,39,42,42,40,22,30,37,35,27,47,18,40,40,37,39,47,22,40,25,43,36,31,36,30,38,38,33,48,45,34,37,16,25,46,45,29,46,44,37,53,26,44,26,25,33,18,31,36,41,40,44,44,30,19,25,37,51,42,32,35,32,32,40,38,47,31,36,33,42,33,41,38,46,31,50,47,38,40,31,41,40,35,34,41,31,39,36,43,48,10,30,25,38,49,46,32,27,30,26,34,31,29,39,35,38,31,45,37,41,33,37,39,40,36,33,36,45,39,34,45,31,35,39,40,30,41,35,40,38,23,40,35,46,43,33,18,25,36,44,30,28,34,32,33,44,29,35,39,39,43,25,40,39,42,51,34,39,46,41,39,40,39,37,30,24,29,44,42,41,32,48,31,34,32,31,39,42,43,45,35,30,36,33,31,49,26,41,28,40,28,40,8,45,36,22,19,34,44,33,46,45,31,29,45,37,38,40,35,40,32,38,40,35,40,46,33,33,39,44,36,47,41,36,35,37,19,48,47,29,44,42,11,41,34,39,27,44,37,35,25,39,32,40,12,34,32,40,37,40,35,20,38,38,33,29,27,27,36,32,33,19,45,38,11,37,30,40,42,35,34,36,47,32,48,30,14,36,41,44,31,10,40,34,40,38,34,37,35,17,41,31,36,35,41,47,43,22,44,41,52,36,34,36,38,37,45,34,29,40,39,44,27,39,37,41,24,35,40,49,40,38,44,40,47,36,27,37,45,36,26,44,39,35,27,38,33,34,40,32,50,34,44,30,35,16,30,36,45,42,34,44,35,39,34,35,45,42,38,46,46,20,43,47,45,35,32,42,28,37,46,40,45,31,38,27,37,31,39,45,39,43,34,34,38,38,36,37,36,43,31,30,34,28,36,47,47,29,43,41,39,40,32,43,34,43,45,35,44,39,45,35,49,34,31,37,42,23,21,35,30,37,45,32,32,49,24,25,32,29,40,43,36,34,37,29,39,45,37,35,38,48,40,45,35,23,34,44,49,39,37,38,37,27,45,32,47,38,31,48,35,41,39,30,36,27,33,32,41,38,44,39,41,25,37,35,39,49,29,27,45,26,27,40,38,30,36,29,46,50,45,35,36,30,10,48,36,34,20,33,48,23,35,40,44,24,45,32,35,37,42,41,45,14,42,41,36,29,41,43,21,40,37,21,42,41,44,45,27,39,41,46,39,29,29,35,42,38,22,50,34,42,36,42,19,22,28,26,25,25,36,37,47,47,35,43,43,31,44,27,30,33,30,39,41,47,28,44,37,43,49,31,34,47,35,35,46,32,41,38,27,38,44,30,43,52,40,45,34,43,33,39,33,40,37,36,41,42,27,28,43,46,36,39,35,38,29,47,34,38,31,43,28,44,37,34,46,37,35,35,39,39,47,42,38,31,37,38,38,25,31,31,23,42,31,35,41,42,33,41,39,47,27,8,43,43,42,45,44,36,36,24,37,33,37,40,37,36,48,29,37,33,40,41,34,39,40,36,38,34,43,38,47,41,39,41,40,44,39,37,44,36,28,42,42,42,33,24,44,42,36,44,37,40,34,38,17,41,34,38,42,42,44,38,19,36,27,34,43,34,34,44,35,36,41,42,38,41,48,32,45,27,39,27,47,39,34,44,40,41,42,37,25,31,34,26,36,40,32,34,28,30,51,40,41,42,25,38,41,27,47,36,41,39,41,38,39,41,42,37,32,36,19,45,35,42,34,39,38,14,23,27,45,28,33,36,41,38,38,19,25,41,40,43,29,45,40,38,32,44,33,38,38,29,34,44,27,20,44,34,38,27,37,28,28,24,41,39,29,34,28,47,35,29,46,35,30,28,36,41,40,32,50,43,46,27,38,37,40,26,30,34,46,39,37,43,34,42,46,33,37,20,35,30,28,29,37,52,22,45,41,38,29,39,27,34,44,38,45,38,31,41,48,35,36,42,44,33,37,40,46,47,42,39,37,29,39,42,43,31,35,40,42,41,44,49,35,43,28,40,43,29,36,16,33,40,40,40,52,29,31,40,37,32,36,34,33,40,39,29,36,44,41,42,29,35,28,53,37,30,44,33,47,41,36,27,31,37,52,33,41,32,7,34,15,39,46,39,34,37,30,40,48,37,53,36,42,32,37,44,41,40,46,42,42,22,37,34,30,25,41,28,40,51,21,34,49,39,39,30,17,37,39,50,44,33,43,45,30,34,30,45,29,31,44,28,33,40,35,35,35,39,39,42,37,20,41,29,32,23,26,36,38,37,36,16,25,37,39,31,36,30,25,45,36,37,41,30,40,40,42,27,40,41,23,40,34,39,39,43,38,37,48,47,35,32,44,39,38,45,31,41,31,41,28,27,42,40,43,27,40,33,41,32,38,46,40,37,30,40,29,44,38,33,33,38,37,30,33,30,21,34,41,24,34,26,44,41,42,43,28,37,35,31,38,29,41,39,33,34,41,35,22,28,33,34,28,47,30,30,31,38,30,33,40,43,35,42,34,47,28,33,51,30,33,32,23,39,35,41,27,33,34,42,40,42,47,40,38,44,43,43,33,42,42,37,36,40,35,41,43,34,48,44,39,34,30,42,27,37,42,39,38,31,25,32,39,28,35,40,33,35,46,32,31,40,43,28,41,38,41,43,21,42,39,43,39,44,36,34,45,36,29,45,35,44,24,39,35,35,14,27,32,39,34,42,36,40,38,27,47,40,30,27,45,35,39,44,38,39,45,36,32,29,36,37,33,39,33,44,34,40,36,30,38,41,37,21,39,28,39,39,30,45,36,35,47,44,31,38,31,36,37,26,38,34,33,31,35,40,40,48,38,38,44,42,37,41,40,37,38,33,50,51,29,51,41,37,26,44,42,42,31,44,32,44,28,36,27,31,33,29,36,35,36,40,27,35,47,43,41,34,30,44,46,34,45,23,46,23,35,30,37,48,38,38,41,38,38,25,32,29,39,33,43,35,39,37,39,46,46,32,41,35,14,33,45,38,45,37,33,36,45,31,39,33,41,22,39,45,26,36,26,32,33,37,46,26,47,38,36,42,23,35,35,51,19,45,34,28,42,48,33,42,40,32,43,28,34,41,8,25,42,12,27,41,40,33,12,44,35,46,32,38,28,36,38,37,40,33,41,16,33,43,25,44,43,42,45,29,40,16,36,37,42,46,38,35,54,35,41,25,35,25,37,42,41,45,39,43,37,35,31,30,23,45,36,41,31,39,47,43,44,23,38,25,38,46,15,39,34,42,36,32,27,40,40,36,37,39,44,40,40,45,36,18,38,26,42,35,38,39,50,25,25,43,45,36,30,37,35,44,41,19,43,37,41,44,33,39,41,30,37,38,44,42,39,36,42,26,48,50,10,32,37,41,33,44,37,35,31,40,41,36,42,38,35,47,46,38,38,41,31,39,42,40,34,36,38,26,36,37,31,37,43,42,29,37,41,26,47,40,38,40,40,36,28,10,42,32,30,36,42,34,41,41,47,31,41,48,42,44,40,35,36,32,40,36,39,38,39,47,42,16,44,49,32,31,33,40,43,35,35,48,43,30,30,41,39,32,36,29,40,19,22,34,32,19,37,42,35,44,28,40,28,35,32,41,25,33,41,37,43,36,32,37,47,40,27,41,38,36,28,15,29,36,49,34,37,27,41,27,34,37,23,42,37,44,33,37,41,31,36,31,31,38,39,46,28,35,38,46,34,48,8,31,27,30,30,32,38,51,37,42,31,41,37,18,30,14,40,36,41,31,27,39,31,35,29,29,25,39,44,38,33,18,38,41,37,37,29,37,39,22,28,38,13,28,29,42,37,39,40,32,27,38,39,40,36,27,33,49,30,31,34,39,45,50,29,43,44,21,37,37,39,38,37,36,40,34,36,33,42,42,43,40,48,39,42,26,51,37,34,36,36,29,40,40,41,41,39,44,40,40,37,42,26,36,43,37,37,38,36,19,43,45,39,43,35,40,39,46,43,40,41,47,34,48,43,47,52,38,45,41,38,41,39,37,22,33,40,48,44,34,31,39,36,44,36,32,36,42,30,40,41,42,46,34,43,41,41,39,36,27,42)
VarExplained_Protein_SumofSquares=c(36,17,104,23,10,100,43,88,151,13,24,25,126,24,52,30,173,43,79,27,24,172,67,40,37,12,90,23,35,34,51,40,15,55,522,43,53,33,64,20,11,34,15,15,27,19,44,20,18,39,26,107,15,64,37,48,46,31,39,26,15,25,15,258,137,15,60,14,51,28,413,11,59,13,8,36,23,11,31,33,11,8,27,35,77,10,37,13,57,9,49,23,14,63,22,903,11,65,19,10,36,41,17,11,6,220,24,13,48,15,7,32,49,43,23,26,18,13,44,32,22,13,108,21,21,17,163,11,35,28,40,9,17,11,43,72,18,41,51,42,458,7,19,61,61,53,23,26,27,126,21,20,23,42,113,22,39,57,10,6,29,14,23,37,10,17,10,35,16,170,23,25,218,46,33,25,182,29,12,20,34,22,713,29,7,286,34,30,546,11,31,28,15,12,30,10,43,65,21,19,29,246,146,18,31,252,14,58,17,7,14,44,10,24,692,113,72,23,127,51,17,55,99,24,22,10,31,52,68,30,305,59,71,11,32,30,15,12,29,21,16,12,57,18,17,477,87,116,40,11,447,33,60,260,13,14,57,40,8,12,99,11,15,17,12,26,8,12,22,23,25,14,14,26,17,27,54,30,11,16,25,43,6,110,78,35,12,11,7,11,9,18,22,8,12,58,34,112,13,32,52,8,16,16,10,8,20,18,47,15,12,17,45,6,34,11,14,24,47,66,25,18,5,17,28,11,31,7,659,52,11,48,7,18,25,23,37,20,56,24,7,13,167,19,20,54,38,88,56,18,8,8,47,16,23,13,7,51,12,8,26,25,228,63,480,38,7,59,70,17,18,27,9,97,31,14,7,18,34,46,1085,21,21,50,42,20,22,5,17,21,11,28,33,46,22,19,99,10,16,8,9,39,18,11,5,11,14,53,9,15,11,16,12,103,23,13,8,24,41,24,18,24,12,12,19,22,54,22,15,15,153,8,68,21,40,43,9,9,66,50,26,20,24,62,12,56,22,14,25,23,7,38,55,21,34,12,11,9,36,159,12,12,16,13,8,14,9,10,82,53,4,27,18,11,35,18,19,30,15,38,41,16,23,16,25,12,16,19,61,11,20,78,102,12,16,20,19,15,11,320,15,9,11,7,37,31,13,30,24,27,11,120,51,7,7,11,23,20,34,258,25,16,32,30,36,67,10,22,25,57,14,89,27,13,29,19,9,675,102,27,14,17,7,8,14,13,13,6,642,31,14,15,15,61,16,16,36,16,83,31,32,30,71,147,20,13,9,40,320,35,183,12,49,26,21,19,10,28,18,20,53,7,17,12,33,30,50,6,8,67,370,9,60,88,25,17,27,62,290,30,22,19,20,38,5,11,19,82,12,12,17,39,14,9,9,94,25,20,23,12,37,105,16,34,41,23,72,44,50,15,17,8,21,20,86,8,19,160,45,33,9,27,29,839,15,22,9,7,10,27,56,12,22,16,14,8,24,7,32,42,25,101,44,27,13,1113,15,8,50,13,33,32,6,14,21,52,13,34,11,3,13,10,20,117,21,122,10,16,68,11,143,39,16,37,11,42,71,7,9,24,25,10,20,14,22,27,13,12,14,64,40,10,101,15,27,36,97,6,24,10,43,127,98,24,17,41,4,9,8,187,30,17,24,33,209,14,57,7,15,12,9,150,45,53,26,52,35,76,62,24,25,17,18,17,12,204,56,10,9,10,13,14,5,22,7,25,14,28,29,75,50,60,11,31,22,161,11,30,24,64,30,19,8,5,20,62,109,14,12,112,15,15,6,6,10,38,106,146,16,16,47,75,10,19,48,7,28,19,32,72,261,51,6,41,112,21,12,15,27,26,45,48,23,38,13,14,34,4,247,123,24,7,9,76,28,15,10,18,104,97,10,572,34,10,6,46,11,37,12,61,16,9,15,29,67,1160,59,31,11,30,26,9,19,37,20,11,65,19,317,11,22,16,31,44,153,9,69,10,33,28,16,1406,10,8,23,30,10,17,21,8,25,46,22,11,29,9,36,12,66,45,108,239,14,10,12,16,20,54,9,572,10,7,34,12,62,42,5,585,6,37,25,31,27,25,6,16,14,9,112,88,10,22,152,32,23,36,42,13,23,63,16,10,7,21,9,35,8,17,25,21,17,16,17,31,15,11,7,16,142,23,27,301,10,57,31,42,49,10,25,10,69,13,24,25,8,40,267,7,62,44,14,13,41,10,8,25,11,17,19,37,24,12,17,19,22,44,11,62,341,13,11,46,11,32,19,12,17,12,10,24,8,69,27,49,90,32,12,13,10,10,6,37,36,46,13,26,49,32,4,55,12,8,28,57,19,61,16,6,39,27,19,11,28,26,57,92,32,8,7,14,9,62,21,10,21,154,28,19,21,24,38,12,19,37,18,5,3,68,10,44,36,137,18,82,7,10,16,14,16,15,17,6,43,39,29,17,13,15,24,21,24,56,7,7,2,8,7,9,26,22,24,27,16,14,81,23,17,21,22,69,13,5,10,43,16,18,113,44,11,9,157,92,26,15,17,33,34,57,8,14,34,133,55,13,39,7,10,15,28,45,93,12,6,16,12,9,15,18,158,47,45,49,28,33,26,9,27,329,84,13,9,55,67,23,16,33,10,18,35,73,9,184,26,24,28,89,37,13,14,11,8,16,48,6,21,22,6,23,29,47,16,20,16,12,12,39,143,6,12,112,11,29,25,12,51,12,56,4,92,20,20,52,45,28,17,29,16,65,62,138,76,27,49,42,51,43,27,28,45,29,113,69,8,76,43,30,162,60,8,56,255,89,119,11,12,7,7,169,13,81,7,12,71,47,20,15,21,18,29,169,11,16,19,28,13,19,59,72,12,20,9,27,41,7,18,24,31,37,246,69,7,8,12,145,86,16,18,15,13,20,26,11,10,110,11,17,213,40,10,15,13,5,41,64,56,9,14,41,43,16,7,11,10,70,1112,20,9,11,10,777,13,25,62,32,10,44,29,201,13,67,20,11,15,16,5,9,12,68,24,147,14,48,53,22,9,19,13,55,46,372,7,23,22,78,24,10,9,17,10,27,14,7,66,19,54,12,14,15,13,24,13,11,423,10,39,24,3,9,16,11,9,12,16,106,35,14,13,17,31,22,7,7,10,17,160,26,13,9,59,17,31,9,6,11,10,27,31,51,38,15,32,12,39,98,39,9,20,26,38,52,10,9,49,336,315,10,7,16,7,7,121,786,10,8,14,42,20,38,15,25,10,15,114,1174,72,5,17,14,243,47,16,28,13,15,10,9,8,27,187,13,20,16,15,18,21,7,14,22,13,15,8,9,9,14,659,12,55,18,7,8,17,14,16,42,26,15,12,66,211,62,226,51,14,137,20,57,6,10,35,86,7,20,30,68,10,11,14,7,18,9,14,16,7,75,84,12,11,17,224,7,15,20,15,13,5,24,9,14,61,13,5,26,9,8,28,37,9,19,52,168,13,6,18,53,14,32,9,20,781,68,35,9,4,103,71,31,17,11,14,29,55,25,128,15,24,17,31,26,14,27,124,18,14,10,90,61,25,413,9,51,31,18,14,96,10,10,31,29,32,32,65,20,26,17,28,18,35,9,25,15,49,159,16,8,19,131,27,12,10,16,89,5,19,26,102,173,70,6,158,123,14,223,40,6,52,7,5,24,75,36,116,19,33,32,25,26,89,8,26,6,19,23,13,9,29,36,10,47,14,15,12,17,66,12,19,24,67,5,11,76,33,8,14,8,11,15,3,20,21,11,15,65,18,7,7,65,6,10,23,6,6,444,9,76,17,177,33,23,18,76,28,10,18,58,31,30,370,9,13,12,52,18,8,10,43,15,17,41,18,12,14,16,19,12,22,40,24,29,74,12,14,6,8,66,25,9,16,26,7,25,17,16,158,7,177,72,58,22,33,10,41,4,89,10,29,113,11,15,11,12,57,14,9,616,50,105,7,12,10,25,38,14,9,23,16,23,32,21,50,8,23,12,23,5,11,17,13,11,50,14,65,17,22,35,28,37,198,33,5,17,94,36,52,36,22,251,28,210,8,146,35,9,22,16,15,23,11,32,9,20,12,17,21,57,17,239,75,16,15,55,20,65,16,49,8,40,136,13,7,41,6,24,14,29,17,20,139,12,52,20,30,73,8,166,234,67,16,10,70,14,126,203,14,38,37,16,43,25,33,20,11,17,32,14,8,32,44,21,11,14,7,43,151,15,9,123,36,215,34,13,23,13,24,23,303,20,110,638,28,22,9,258,21,15,10,161,8,92,11,13,21,9,26,28,85,101,31,5,20,20,9,19,49,24,10,16,14,28,1133,378,20,56,46,77,172,33,28,19,26,28,5,25,45,54,18,142,15,12,24,12,11,75,48,17,8,18,20,45,30,15,25,50,24,17,57,8,29,30,38,23,22,57,12,13,20,30,43,27,16,65,18,13,16,77,9,10,621,35,25,27,17,67,13,38,184,6,22,24,9,17,10,11,27,35,28,70,17,12,17,42,14,14,1022,8,19,42,238,28,14,9,63,12,81,17,15,12,13,17,18,25,11,24,22,16,33,54,18,4,8,12,18,6,79,19,47,75,15,16,27,19,25,7,21,31,62,22,10,499,30,28,13,14,13,6,115,10,63,103,12,24,32,13,31,98,5,4,17,124,32,23,26,4,21,29,24,21,46,137,29,16,11,26,180,156,20,61,194,17,39,8,92,77,35,26,11,42,24,37,29,18,1083,45,13,18,131,70,31,241,8,16,28,10,59,12,15,38,103,17,76,12,143,14,286,17,91,36,16,13,12,20,37,6,170,17,32,602,21,23,10,33,200,37,9,166,19,30,32,253,41,14,11,44,28,9,9,8,158,14,1203,31,207,10,143,23,3,189,89,461,19,23,23,47,20,18,26,21,22,32,25,22,165,17,40,11,166,6,134,17,47,63,134,45,30,25,106,12,54,14,156,69,33,35,73,37,11,15,14,11,18,81,413,38,13,17,53,98,144,14,26,63,109,10,34,8,14,19,13,12,15,29,14,168,40,25,21,19,143,9,14,24,35,28,8,128,4,141,15,39,34,29,153,14,38,15,235,16,28,64,13,134,17,13,8,16,52,7,4,12,11,45,33,13,17,15,21,8,12,27,100,11,14,24,22,10,16,13,11,160,9,7,12,42,14,197,20,33,9,20,13,19,21,10,16,17,7,52,20,42,20,27,18,67,52,3,46,62,44,7,13,64,293,18,32,14,8,30,12,60,52,455,111,48,15,8,15,16,37,11,91,8,9,45,8,8,20,46,14,4,42,19,15,26,27,20,11,14,5,43,13,5,8,20,37,20,18,345,44,115,24,11,24,47,15,112,23,19,26,90,27,20,13,7,8,16,50,26,14,11,37,13,38,32,12,37,15,12,16,6,9,12,88,4,16,14,26,11,36,8,23,22,15,5,26,13,24,10,57,238,26,8,12,15,7,41,10,12,20,45,29,224,95,27,13,28,14,67,11,42,28,42,8,80,28,64,46,12,16,13,26,9,10,57,175,19,7,17,45,48,17,25,32,296,34,13,10,122,16,33,609,14,12,14,11,21,16,33,9,45,35,135,52,407,74,20,21,565,69,19,16,21,13,33,39,28,32,31,13,29,13,14,14,15,22,43,11,15,57,19,28,11,117,55,10,86,18,12,12,11,48,27,16,6,24,24,23,10,35,7,11,45,435,10,85,8,32,6,81,16,10,37,20,34,22,225,15,27,68,55,69,240,11,28,11,95,16,8,9,60,9,10,40,27,153,159,20,40,50,12,778,8,12,23,145,21,132,22,13,10,60,16,96,5,15,9,84,6,7,83,9,35,15,58,7,13,18,18,6,16,12,12,17,17,13,16,17,147,32,21,49,7,101,74,11,16,12,47,27,12,17,8,17,21,114,19,6,99,19,5,12,216,15,46,50,46,22,70,45,26,89,64,25,22,88,15,1038,66,37,103,18,8,31,11,22,14,18,43,10,106,25,26,67,39,32,199,15,19,38,25,9,12,10,19,12,99,8,15,14,15,15,8,12,30,18,24,4,69,61,9,5,16,7,18,24,29,156,99,256,20,21,11,9,6,14,15,257,11,75,17,16,22,12,86,16,7,11,11,12,48,33,7,32,10,109,60,70,12,70,71,32,41,36,6,86,18,733,42,79,13,171,25,8,7,13,37,13,11,96,12,39,15,41,48,21,138,895,12,10,94,8,8,18,9,23,9,9,54,12,290,8,13,37,26,12,203,17,7,66,43,43,24,10,17,18,35,11,6,18,33,47,210,22,244,20,13,77,32,94,40,15,216,33,31,1454,35,68,37,7,22,16,23,27,8,44,246,23,50,570,26,9,972,55,14,23,75,730,19,108,20,195,23,17,62,14,35,16,11,16,11,118,22,27,29,20,19,36,16,8,16,14,18,29,131,27,13,50,15,20,97,48,62,578,81,15,75,19,51,23,9,86,19,121,571,28,39,47,17,6,22,16,149,55,20,12,13,23,64,10,114,13,42,196,7,14,14,42,16,76,19,42,996,13,106,20,37,16,116,7,10,17,13,20,10,16,18,47,18,12,20,8,30,15,33,22,28,14,36,24,19,24,24,39,20,12,28,15,22,16,141,15,31,361,14,6,104,16,9,29,50,162,9,33,30,19,12,7,8,9,52,47,1475,75,79,20,206,14,18,10,8,27,10,24,25,55,28,20,16,27,18,130,23,7,16,101,31,37,207,6,116,10,211,28,11,12,19,11,12,21,8,14,26,25,14,19,13,22,21,11,18,9,15,42,16,41,11,107,10,49,34,26,32,12,30,10,29,24,32,16,19,22,31,19,12,96,20,43,49,24,18,41,126,72,42,30,32,16,35,118,23,16,19,44,12,4,414,12,25,123,10,21,11,9,6,135,7,19,20,14,8,16,8,28,16,19,47,10,13,25,34,11,2024,25,306,24,44,10,12,13,11,10,31,78,10,68,18,40,25,26,17,15,14,37,30,16,10,22,18,25,15,10,25,44,9,17,11,14,80,9,24,16,12,52,18,8,12,39,25,30,9,13,9,26,13,26,23,22,39,108,91,24,17,10,91,26,38,1229,29,72,21,75,34,128,25,27,13,24,13,11,252,13,19,17,23,12,9,88,81,18,7,19,7,34,1740,13,54,27,36,42,38,11,38,7,20,7,56,13,8,14,23,115,17,35,8,32,18,9,81,241,56,25,19,13,6,191,71,86,28,36,11,36,185,55,13,37,22,19,6,11,24,18,7,21,35,59,15,35,6,28,40,21,10,45,22,11,12,21,10,130,16,109,35,15,12,4,29,20,68,18,45,69,7,6,13,71,14,6,128,170,36,24,10,134,8,35,44,55,23,14,23,44,819,28,26,8,25,92,9,11,18,42,16,20,13,9,7,494,53,1412,19,19,127,10,14,55,6,24,23,89,14,12,67,19,102,23,8,295,1111,46,5,78,18,42,19,21,435,9,23,23,211,128,27,7,22,46,186,53,19,92,119,179,13,360,47,12,170,34,17,23,147,29,31,32,12,15,21,38,72,16,20,384,38,30,15,46,400,12,29,240,48,96,35,17,6,224,16,19,16,22,10,17,23,13,9,19,73,23,108,57,20,7,23,6,31,12,25,20,30,12,52,35,16,24,53,135,68,219,10,15,38,11,48,19,14,13,43,24,21,39,15,30,17,61,12,40,6,15,18,10,10,55,35,13,41,331,67,17,50,23,50,12,54,12,13,16,10,14,9,25,17,59,15,163,145,26,92,128,5,11,25,7,23,15,14,23,20,45,10,27,12,52,74,8,16,19,19,10,16,30,17,45,11,34,43,30,55,23,62,55,10,22,14,14,124,394,4,18,15,16,6,7,6,14,69,18,20,10,44,30,406,8,16,54,15,15,16,2191,14,17,12,20,21,50,8,161,12,61,16,66,12,20,55,9,11,11,202,117,36,27,18,13,34,39,37,8,27,26,35,5,53,35,6,60,42,15,13,15,37,22,14,10,7,26,10,12,16,25,14,7,48,15,13,254,23,12,44,15,16,31,29,22,282,26,25,34,112,24,12,296,38,15,25,58,10,28,51,91,21,20,14,12,14,54,14,53,24,27,25,22,119,27,146,8,13,13,26,38,34,40,6,67,193,59,93,80,214,115,6,39,11,11,8,4,11,91,18,28,11,104,432,230,23,12,49,21,14,6,9,8,10,34,113,23,27,29,12,21,30,18,27,20,25,9,19,47,33,34,19,55,160,15,21,9,61,62,17,160,15,16,7,22,8,11,7,161,12,18,23,51,32,24,19,11,16,26,18,26,26,9,15,33,21,156,6,25,15,12,113,12,430,9,140,147,13,8,124,8,17,6,26,19,27,16,137,12,255,17,35,1112,29,8,5,58,41,31,15,16,28,12,10,14,31,26,79,26,28,451,7,16,18,20,19,36,12,22,14,70,7,69,14,71,35,15,50,21,13,61,8,7,17,91,26,20,145,32,19,10,177,21,477,41,26,14,15)
vioplot(VarExplained_Protein_Genotype, VarExplained_Protein_Diet, VarExplained_Protein_Age, VarExplained_Protein_Interaction, VarExplained_Protein_Residual, col=c("orangered","firebrick1","firebrick3","darkred","black"), ylim=c(0,100))

#$$$$$$ #$$$$$$ #$$$$$$ #$$$$$$ #$$$$$$ #$$$$$$ #$$$$$$ #$$$$$$
#$$$$$$ #$$$$$$ #$$$$$$ #$$$$$$ #$$$$$$ #$$$$$$ #$$$$$$ #$$$$$$
# PANEL 2C 

library(LSD)

# LEFT plot (NB: load these Diet_FC variables for mRNA and protein above in Panel A section; exact same data are used)
Xaxis=rm.outlier(Diet_FC_Protein_All, fill=T)	# Replace huge outliers with NA
Yaxis=rm.outlier(Diet_FC_mRNA_All, fill=T)	# Replace huge outliers with NA
heatscatter(Xaxis, Yaxis, cex=0.6, ylim=c(-0.5,0.5), xlim=c(-0.5,0.5))
abline(lm(Yaxis~Xaxis), lty=1, lwd=1, col="black")
cor(Xaxis, Yaxis, method=c("pearson"), use="complete.obs")
signif(summary(lm(Yaxis~Xaxis))$coefficients[8], digits=2)

# RIGHT plot
mRNAdat_All=Top3772Data[1:3772,]
Protdat_All=Top3772Data[3773:dim(Top3772Data_CD)[1],]
age=Top3772Data_header[4,]
Age_Correlation_mRNA_All=NA
Age_Correlation_Protein_All=NA

for(i in 1:dim(mRNAdat_All)[1]) {
	Age_Correlation_mRNA_All[i]=cor(as.numeric(mRNAdat_All[i,]), as.numeric(age), method=c("spearman"), use="complete.obs")
}

for(i in 1:dim(Protdat_All)[1]) {
	Age_Correlation_Protein_All[i]=cor(as.numeric(Protdat_All[i,]), as.numeric(age), method=c("spearman"), use="complete.obs")
}

Xaxis=rm.outlier(Age_Correlation_Protein_All, fill=T)	# Replace outliers with NA
Yaxis=rm.outlier(Age_Correlation_mRNA_All, fill=T)	# Replace outliers with NA
heatscatter(Xaxis, Yaxis, cex=0.6, ylim=c(-0.5,0.5), xlim=c(-0.5,0.5))
abline(lm(Yaxis~Xaxis), lty=1, lwd=1, col="black")
cor(Xaxis, Yaxis, method=c("pearson"), use="complete.obs")
signif(summary(lm(Yaxis~Xaxis))$coefficients[8], digits=2)


#$$$$$$ #$$$$$$ #$$$$$$ #$$$$$$ #$$$$$$ #$$$$$$ #$$$$$$ #$$$$$$
#$$$$$$ #$$$$$$ #$$$$$$ #$$$$$$ #$$$$$$ #$$$$$$ #$$$$$$ #$$$$$$
# PANEL 2D 

library(LSD)

# Taking the average abundance for the 3772 matching genes
mRNA_Abundance=rowMeans(mRNAdat_All,na.rm=T)
Protein_Abundance=rowMeans(Protdat_All,na.rm=T)

Xaxis=rm.outlier(Protein_Abundance, fill=T)	# Replace huge outliers with NA
Yaxis=rm.outlier(mRNA_Abundance, fill=T)	# Replace huge outliers with NA
heatscatter(Xaxis, Yaxis, cex=0.6, ylim=c(0,15), xlim=c(4,22))
abline(lm(Yaxis~Xaxis), lty=1, lwd=1, col="black")
cor(Xaxis, Yaxis, method=c("pearson"), use="complete.obs")
signif(summary(lm(Yaxis~Xaxis))$coefficients[8], digits=2)


#$$$$$$ #$$$$$$ #$$$$$$ #$$$$$$ #$$$$$$ #$$$$$$ #$$$$$$ #$$$$$$
#$$$$$$ #$$$$$$ #$$$$$$ #$$$$$$ #$$$$$$ #$$$$$$ #$$$$$$ #$$$$$$
# PANEL 2E 
# Random correlations are using the real data from all 3772 paired mRNA and proteins, but shuffling gene labels
# "All" is for using all 275 individuals with data at both layers (mRNA, protein).

library(gplots)
Random_Individuals=NA
Protdat_All_Random_Individuals=sample(Protdat_All) # This shuffles columns so we're still comparing the right genes, but with the animal labels mixed up
mRNAdat_All_Random_Individuals=sample(mRNAdat_All) # This shuffles columns 

for(i in 1:dim(mRNAdat_All)[1]) {
	Random_Individuals[i]=cor(as.numeric(Protdat_All_Random_Individuals[i,]), as.numeric(mRNAdat_All_Random_Individuals[i,]), method=c("spearman"), use="complete.obs")
}

# This makes the correlations for all mRNA to protein (the file is ordered precisely so simply indexes can be used)
All=NA
for(i in 1:dim(mRNAdat_All)[1]) {
	All[i]=cor(as.numeric(Protdat_All[i,]), as.numeric(mRNAdat_All[i,]), method=c("spearman"), use="complete.obs")
}

plot(density(Random_Individuals, na.rm=T), xlim=c(-1,1), ylim=c(0, 6.2), col="black")
plotCI(density(All), col="lightblue", add=T, type="l")

signif(t.test(Random_Individuals,All, paired=TRUE)$p.value,digits=2) # or paired-false? should be true because it's the same gene before and after. 
shapiro.test(All)
shapiro.test(Random_Individuals)


#$$$$$$ #$$$$$$ #$$$$$$ #$$$$$$ #$$$$$$ #$$$$$$ #$$$$$$ #$$$$$$
#$$$$$$ #$$$$$$ #$$$$$$ #$$$$$$ #$$$$$$ #$$$$$$ #$$$$$$ #$$$$$$
# PANEL 2F 
# Random correlations are using the real data from all 3772 paired mRNA and proteins, but shuffling gene labels
# "All" is for using all 275 individuals with data at both layers (mRNA, protein).
# LowVar is comparing mRNA�protein correlations only for mRNAs with the lowest levels of variation
# HighVar is the converse, showing correlations only for mRNAs with the highest variation

# LEFT PANEL
# Load the mRNA and protein datasets from the previous panels
library(gplots)

# Now calculate the variance at the mRNA & protein level
Variance_mRNA=NA
for(i in 1:dim(mRNAdat_All)[1]) {
	Variance_mRNA[i]=var(as.numeric(mRNAdat_All[i,]), na.rm=T)
}

Variance_Prot=NA
for(i in 1:dim(Protdat_All)[1]) {
	Variance_Prot[i]=var(as.numeric(Protdat_All[i,]), na.rm=T)
}

Variance_mRNA_Top10pct=as.numeric(quantile(Variance_mRNA,prob=seq(0,1,length=11))[10])
Variance_Prot_Top10pct=as.numeric(quantile(Variance_Prot,prob=seq(0,1,length=11))[10])
Variance_mRNA_Bottom10pct=as.numeric(quantile(Variance_mRNA,prob=seq(0,1,length=11))[2])
Variance_Prot_Bottom10pct=as.numeric(quantile(Variance_Prot,prob=seq(0,1,length=11))[2])

HighVar_Correlation_mRNA_Protein=NA
LowVar_Correlation_mRNA_Protein=NA

for(i in 1:dim(mRNAdat_All)[1]) {
	if(Variance_mRNA[i]>=Variance_mRNA_Top10pct){
		HighVar_Correlation_mRNA_Protein[i]=cor(as.numeric(Protdat_All[i,]), as.numeric(mRNAdat_All[i,]), method=c("spearman"), use="complete.obs")
	}
	else
		HighVar_Correlation_mRNA_Protein[i]=NA
}

for(i in 1:dim(mRNAdat_All)[1]) {
	if(Variance_mRNA[i]<=Variance_mRNA_Bottom10pct){
		LowVar_Correlation_mRNA_Protein[i]=cor(as.numeric(Protdat_All[i,]), as.numeric(mRNAdat_All[i,]), method=c("spearman"), use="complete.obs")
	}
	else
		LowVar_Correlation_mRNA_Protein[i]=NA
}

plot(density(Random_Individuals, na.rm=T), xlim=c(-1,1), ylim=c(0, 6), col="black")
plotCI(density(All), col="lightblue", add=T, type="l")
plotCI(density(HighVar_Correlation_mRNA_Protein, na.rm=T), col="red", add=T, type="l")
plotCI(density(LowVar_Correlation_mRNA_Protein, na.rm=T), col="green", add=T, type="l")
signif(t.test(HighVar_Correlation_mRNA_Protein,LowVar_Correlation_mRNA_Protein, paired=FALSE)$p.value,digits=2) # 


# RIGHT PANEL
# NOTE: Calculating QTLs takes a significant amount of time and processing; see supporting documents for QTL analysis
# The code for calculating the LOD scores and p-values is in the file "b_Figure2_HelperFile_QTLs.r" which should be found alongside this document.
# Thus, a "shortcut" file is provided here which lists the LOD scores at the cis-location (closest marker) for all genes
# Note that the file contains all cis-QTL data, but we only consider the first 3511 which are ordered where mRNA and protein cis-QTLs were calculated
# For the same gene and aligned one-to-one. 

QTLData=read.table("b_Figure2_cisQTLs_All.csv", row.names=1, header=T, sep=",")

mRNA_cisQTL=NA
# The file is sorted to only take the 3511 genes with mRNA data and protein data that output QTLs
# We keep the information for all cis-eQTLs with LOD ��3 (canonical cutoff for significance)
for(i in 1:3511) {
	if(!is.na(as.numeric(as.character(QTLData$mRNA_All_LOD[i])))) {
		if(as.numeric(as.character(QTLData$mRNA_All_LOD[i])) >= 3.0) {
			mRNA_cisQTL[i]=as.numeric(as.character(QTLData$mRNA_All_LOD[i]))
		}
		else if (as.numeric(as.character(QTLData$mRNA_All_LOD[i])) <= 3.0) {
			mRNA_cisQTL[i]=NA
		}	
	}
}

KeepciseQTLs=which(!is.na(mRNA_cisQTL))
Genes_with_ciseQTL=as.character(QTLData$mRNA_Index[KeepciseQTLs])

mRNAdat_All=Top3772Data[1:3772,]
Protdat_All=Top3772Data[3773:dim(Top3772Data)[1],]
Correlation_has_ciseQTL=NA

# This looks up the genes with cis-eQTLs and only performs correlations for those ones
for(i in 1:dim(mRNAdat_All)[1]) {
	if(!is.na(match(rownames(mRNAdat_All)[i],Genes_with_ciseQTL))) { 
		Correlation_has_ciseQTL[i]=cor(as.numeric(mRNAdat_All[i,]), as.numeric(Protdat_All[i,]), method=c("spearman"), use="complete.obs")
	}
}

Prot_cisQTL=NA
# The file is sorted to only take the 3511 genes with mRNA data and protein data that output QTLs
# We keep the information for all cis-eQTLs with LOD ��3 (canonical cutoff for significance)
for(i in 1:3511) {
	if(!is.na(as.numeric(as.character(QTLData$Prot_All_LOD[i])))) {
		if(as.numeric(as.character(QTLData$Prot_All_LOD[i])) >= 3.0) {
			Prot_cisQTL[i]=as.numeric(as.character(QTLData$Prot_All_LOD[i]))
		}
		else if (as.numeric(as.character(QTLData$Prot_All_LOD[i])) <= 3.0) {
			Prot_cisQTL[i]=NA
		}	
	}
}

KeepcispQTLs=which(!is.na(Prot_cisQTL))
Genes_with_cispQTL=as.character(QTLData$mRNA_Index[KeepcispQTLs])

mRNAdat_All=Top3772Data[1:3772,]
Protdat_All=Top3772Data[3773:dim(Top3772Data)[1],]
Correlation_has_cispQTL=NA

# This looks up the genes with cis-eQTLs and only performs correlations for those ones
for(i in 1:dim(mRNAdat_All)[1]) {
	if(!is.na(match(rownames(mRNAdat_All)[i],Genes_with_cispQTL))) { 
		Correlation_has_cispQTL[i]=cor(as.numeric(mRNAdat_All[i,]), as.numeric(Protdat_All[i,]), method=c("spearman"), use="complete.obs")
	}
}


######
Any_cisQTL=NA
# The file is sorted to only take the 3511 genes with mRNA data and protein data that output QTLs
# We keep the information for all cis-eQTLs with LOD ��3 (canonical cutoff for significance)
for(i in 1:3511) {
	if(!is.na(as.numeric(as.character(QTLData$mRNA_All_LOD[i]))) && !is.na(as.numeric(as.character(QTLData$Prot_All_LOD[i])))) {
		if(as.numeric(as.character(QTLData$mRNA_All_LOD[i])) >= 3.0 && as.numeric(as.character(QTLData$Prot_All_LOD[i])) >= 3.0) {
			Any_cisQTL[i]=as.numeric(as.character(QTLData$mRNA_All_LOD[i]))
		}
		else if (as.numeric(as.character(QTLData$mRNA_All_LOD[i])) <= 3.0) {
			Any_cisQTL[i]=NA
		}	
	}
}

KeepcisQTLs=which(!is.na(Any_cisQTL))
Genes_with_cisQTL=as.character(QTLData$mRNA_Index[KeepcisQTLs])

mRNAdat_All=Top3772Data[1:3772,]
Protdat_All=Top3772Data[3773:dim(Top3772Data)[1],]
Correlation_has_cisQTL=NA

# This looks up the genes with cis-eQTLs and only performs correlations for those ones
for(i in 1:dim(mRNAdat_All)[1]) {
	if(!is.na(match(rownames(mRNAdat_All)[i],Genes_with_cisQTL))) { 
		Correlation_has_cisQTL[i]=cor(as.numeric(mRNAdat_All[i,]), as.numeric(Protdat_All[i,]), method=c("spearman"), use="complete.obs")
	}
}

## Then the final plot
plot(density(Random_Individuals, na.rm=T), xlim=c(-1,1), ylim=c(0, 6), col="black")
plotCI(density(Correlation_has_ciseQTL, na.rm=T), col="green", add=T, type="l") # Accepts any gene with a cis-eQTL
plotCI(density(Correlation_has_cispQTL, na.rm=T), col="blue", add=T, type="l") # Accepts any gene with a cis-pQTL
plotCI(density(Correlation_has_cisQTL, na.rm=T), col="grey", add=T, type="l") # Requires BOTH cis-pQTL and cis-eQTL for same gene


signif(t.test(Correlation_has_ciseQTL,Correlation_has_cispQTL, paired=FALSE)$p.value,digits=2) # or paired-false? should be true because it's the same gene before and after. 

#$$$$$$ #$$$$$$ #$$$$$$ #$$$$$$ #$$$$$$ #$$$$$$ #$$$$$$ #$$$$$$
#$$$$$$ #$$$$$$ #$$$$$$ #$$$$$$ #$$$$$$ #$$$$$$ #$$$$$$ #$$$$$$
# PANEL 2G 
# Random correlations are using the real data from all 3772 paired mRNA and proteins, but shuffling gene labels
# "All" is for using all 275 individuals with data at both layers (mRNA, protein).
# A list of all proteins in complexes is downloaded from CORUM: https://mips.helmholtz-muenchen.de/corum/download/allComplexes.txt.zip
# All subunits were taken from ALL complexes, so a gene listed as a CORUM complex member in human is also considered "in complex" in mouse
# All complex sizes, including dimers, are considered "in complex"
# The following genes 


mRNAdat_All=Top3772Data[1:3772,]
Protdat_All=Top3772Data[3773:dim(Top3772Data)[1],]
Correlation_is_Complex=NA
Correlation_is_Not_Complex=NA

setwd("~/Dropbox/Aging\ BXD\ Study/1\ _\ Manuscript\ -\ Evan\ Multi-Omic/")
CORUM_List=read.table("b_Figure2_CORUM_Details.csv", row.names=1, header=T, sep=",")


# This looks up the genes with complex membership and those  without, and splits them into groups
for(i in 1:dim(mRNAdat_All)[1]) {
	if(!is.na(match(rownames(mRNAdat_All)[i],CORUM_List$With_Complex_IDs))) { 
		Correlation_is_Complex[i]=cor(as.numeric(mRNAdat_All[i,]), as.numeric(Protdat_All[i,]), method=c("spearman"), use="complete.obs")
	}
	else {
		Correlation_is_Not_Complex[i]=cor(as.numeric(mRNAdat_All[i,]), as.numeric(Protdat_All[i,]), method=c("spearman"), use="complete.obs")
	}
}

plot(density(Random_Individuals, na.rm=T), xlim=c(-1,1), ylim=c(0, 6), col="black")
plotCI(density(All), col="lightblue", add=T, type="l")
plotCI(density(Correlation_is_Complex, na.rm=T), col="red", add=T, type="l")
plotCI(density(Correlation_is_Not_Complex, na.rm=T), col="green", add=T, type="l")

signif(t.test(Correlation_is_Complex,Correlation_is_Not_Complex, paired=FALSE)$p.value,digits=2) # or paired-false? should be true because it's the same gene before and after. 


#$$$$$$ #$$$$$$ #$$$$$$ #$$$$$$ #$$$$$$ #$$$$$$ #$$$$$$ #$$$$$$
#$$$$$$ #$$$$$$ #$$$$$$ #$$$$$$ #$$$$$$ #$$$$$$ #$$$$$$ #$$$$$$
# PANEL 2H 

library(dplyr)
library(gplots)
library(venneuler)

setwd("~/Dropbox/Aging\ BXD\ Study/1\ _\ Manuscript\ -\ Evan\ Multi-Omic/")

QTLData=read.table("b_Figure2_cisQTLs_All.csv", row.names=1, header=T, sep=",")

CD_mRNA_cisQTL=NA
CD_Prot_cisQTL=NA
Either_cisQTL=NA
	
# The file is sorted to only take the 3511 genes with mRNA data and protein data that output QTLs
for(i in 1:3511) {
	if(!is.na(as.numeric(as.character(QTLData$mRNA_CD_LOD[i])))) {
		if(as.numeric(as.character(QTLData$mRNA_CD_LOD[i])) >= 4.0) {
			CD_mRNA_cisQTL[i]=as.numeric(as.character(QTLData$mRNA_CD_LOD[i]))
		}
		else if (as.numeric(as.character(QTLData$mRNA_CD_LOD[i])) <= 4.0) {
			CD_mRNA_cisQTL[i]=NA
		}	
	}
}
		
for(i in 1:3511) {
	if(!is.na(as.numeric(as.character(QTLData$Prot_CD_LOD[i])))) {
		if(as.numeric(as.character(QTLData$Prot_CD_LOD[i])) >= 4.0) {
			CD_Prot_cisQTL[i]=as.numeric(as.character(QTLData$Prot_CD_LOD[i]))
		}
		else if (as.numeric(as.character(QTLData$Prot_CD_LOD[i])) <= 4.0) {
			CD_Prot_cisQTL[i]=NA
		}	
	}
}

for(i in 1:3511) {
	if(!is.na(as.numeric(as.character(QTLData$mRNA_CD_LOD[i]))) && !is.na(as.numeric(as.character(QTLData$Prot_CD_LOD[i])))) {
		if(as.numeric(as.character(QTLData$mRNA_CD_LOD[i])) >= 4.0 || as.numeric(as.character(QTLData$Prot_CD_LOD[i])) >= 4.0) {	
			Either_cisQTL[i]=as.numeric(as.character(QTLData$mRNA_CD_LOD[i])) # Whichever, it doesn't matter, this is just a placeholder
		}
	}
}

CD_mRNA_cisQTLcount=sum((!is.na(CD_mRNA_cisQTL))) # 117 is correct
CD_Prot_cisQTLcount=sum((!is.na(CD_Prot_cisQTL))) # 116 is correct
Either_cisQTLcount=sum((!is.na(Either_cisQTL)))

# Now count how many QTLs are in each part of the Venn diagram
CD_mRNA_cisQTLcount_unique=Either_cisQTLcount-CD_Prot_cisQTLcount 
CD_Prot_cisQTLcount_unique=Either_cisQTLcount-CD_mRNA_cisQTLcount
cisQTL_Overlap=Either_cisQTLcount-CD_mRNA_cisQTLcount_unique-CD_Prot_cisQTLcount_unique # 47 overlap

plot(venneuler(c(CD=CD_mRNA_cisQTLcount_unique, HFD=CD_Prot_cisQTLcount_unique, "CD&HFD"=cisQTL_Overlap)))
# Note that this plot flips and puts the largest Venn on the left; so the figure is flipped in illustrator since HFD i

# LEFT VENN

# Middle Diagram

KeepcisQTLs=which(!is.na(Either_cisQTL))

greycount=0;skybluecount=0;yellowcount=0;bluecount=0;redcount=0 # initialize the counters
y1=as.numeric(as.character(QTLData$mRNA_CD_LOD[KeepcisQTLs]))
y2=as.numeric(as.character(QTLData$Prot_CD_LOD[KeepcisQTLs]))


x1=rep(1, length=length(y1)) # make this a sequence the length of y1 and y2
x2=rep(2, length=length(y2)) # make this a sequence the length of y1 and y2

plotCI(NA,NA, xlim=c(.9,2.1), ylim=c(0,11), bty="L") ## This initializes the plot
for (i in 1:length(x1)) {
### First plot the lines (you can plot after, but it causes a small visual bug when lines are plotted on top of points)
# If loops are used to set the criteria for line coloration, easy to modify however you see fit
if(!is.na(y1[i]) & !is.na(y2[i])) {
	if(y1[i] >= 4.0 && y2[i] >= 4.0) {
 	 lines(x=c(x1[i],x2[i]), y=c(y1[i],y2[i]), lty="solid", col="grey") # No major change in LOD score
 	 greycount=greycount+1
 	 }
	else if(y1[i] >= 4.0 && y2[i] >= 2.0) {
 	 lines(x=c(x1[i],x2[i]), y=c(y1[i],y2[i]), lty="solid", col="skyblue")
 	 skybluecount=skybluecount+1
 	 }
	else if(y2[i] >= 4.0 && y1[i] >= 2.0) {
 	 lines(x=c(x1[i],x2[i]), y=c(y1[i],y2[i]), lty="solid", col="yellow")
 	 yellowcount=yellowcount+1
 	 } 	 
	else if(y1[i] >= 4.0 && y2[i] < 2.0) {
 	 lines(x=c(x1[i],x2[i]), y=c(y1[i],y2[i]), lty="solid", col="blue")
 	 bluecount=bluecount+1
 	 } 	  	 
	else if(y2[i] >= 4.0 && y1[i] < 2.0) {
 	 lines(x=c(x1[i],x2[i]), y=c(y1[i],y2[i]), lty="solid", col="red")
 	 redcount=redcount+1
 	 } 	  	 
}
 	
# now plot the actual points
  plotCI(x1[i],y1[i],lwd=2,pch=21,pt.bg="grey",col="black", add=T)
  plotCI(x2[i],y2[i],lwd=2,pch=21,pt.bg="grey",col="black", add=T)
}

greycount;skybluecount;yellowcount;bluecount;redcount

# Right Venn
plot(venneuler(c(mRNA=bluecount, Protein=redcount, "mRNA&Protein"=(greycount+skybluecount+yellowcount))))


#$$$$$$ #$$$$$$ #$$$$$$ #$$$$$$ #$$$$$$ #$$$$$$ #$$$$$$ #$$$$$$
#$$$$$$ #$$$$$$ #$$$$$$ #$$$$$$ #$$$$$$ #$$$$$$ #$$$$$$ #$$$$$$
# PANEL 2I

library(dplyr)
library(gplots)
library(venneuler)

# LEFT VENN
setwd("~/Dropbox/Aging\ BXD\ Study/1\ _\ Manuscript\ -\ Evan\ Multi-Omic/")

QTLData=read.table("b_Figure2_cisQTLs_All.csv", row.names=1, header=T, sep=",")

CD_mRNA_cisQTL=NA
HF_mRNA_cisQTL=NA
Either_mRNA_cisQTL=NA
	
# The file is sorted to only take the 3511 genes with mRNA data and protein data that output QTLs
for(i in 1:3511) {
	if(!is.na(as.numeric(as.character(QTLData$mRNA_CD_LOD[i])))) {
		if(as.numeric(as.character(QTLData$mRNA_CD_LOD[i])) >= 4.0) {
			CD_mRNA_cisQTL[i]=as.numeric(as.character(QTLData$mRNA_CD_LOD[i]))
		}
		else if (as.numeric(as.character(QTLData$mRNA_CD_LOD[i])) <= 4.0) {
			CD_mRNA_cisQTL[i]=NA
		}	
	}
}
		
for(i in 1:3511) {
	if(!is.na(as.numeric(as.character(QTLData$mRNA_HF_LOD[i])))) {
		if(as.numeric(as.character(QTLData$mRNA_HF_LOD[i])) >= 4.0) {
			HF_mRNA_cisQTL[i]=as.numeric(as.character(QTLData$mRNA_HF_LOD[i]))
		}
		else if (as.numeric(as.character(QTLData$mRNA_HF_LOD[i])) <= 4.0) {
			HF_mRNA_cisQTL[i]=NA
		}	
	}
}

for(i in 1:3511) {
	if(!is.na(as.numeric(as.character(QTLData$mRNA_CD_LOD[i]))) && !is.na(as.numeric(as.character(QTLData$mRNA_HF_LOD[i])))) {
		if(as.numeric(as.character(QTLData$mRNA_CD_LOD[i])) >= 4.0 || as.numeric(as.character(QTLData$mRNA_HF_LOD[i])) >= 4.0) {	
			Either_mRNA_cisQTL[i]=as.numeric(as.character(QTLData$mRNA_CD_LOD[i])) # Whichever, it doesn't matter, this is just a placeholder
		}
	}
}

CD_mRNA_cisQTLcount=sum((!is.na(CD_mRNA_cisQTL)))
HFD_mRNA_cisQTLcount=sum((!is.na(HF_mRNA_cisQTL)))
Either_mRNA_cisQTLcount=sum((!is.na(Either_mRNA_cisQTL)))

# Now count how many QTLs are in each part of the Venn diagram
CD_mRNA_cisQTLcount_unique=Either_mRNA_cisQTLcount-HFD_mRNA_cisQTLcount
HF_mRNA_cisQTLcount_unique=Either_mRNA_cisQTLcount-CD_mRNA_cisQTLcount
mRNA_cisQTL_Overlap=Either_mRNA_cisQTLcount-CD_mRNA_cisQTLcount_unique-HF_mRNA_cisQTLcount_unique

plot(venneuler(c(CD=CD_mRNA_cisQTLcount_unique, HFD=HF_mRNA_cisQTLcount_unique, "CD&HFD"=mRNA_cisQTL_Overlap)))
# Note that this plot flips and puts the largest Venn on the left; so the figure is flipped in illustrator since HFD is larger, for visual consistency

# Middle Diagram

# We want the indexes of all genes that had cis-eQTLs in either diet:

KeepcisQTLs=which(!is.na(Either_mRNA_cisQTL))

greycount=0;greencount=0;redcount=0;bluecount=0;skybluecount=0 # initialize the counters
y1=as.numeric(as.character(QTLData$mRNA_CD_LOD[KeepcisQTLs]))
y2=as.numeric(as.character(QTLData$mRNA_HF_LOD[KeepcisQTLs]))
x1=rep(1, length=length(y1)) # make this a sequence the length of y1 and y2
x2=rep(2, length=length(y2)) # make this a sequence the length of y1 and y2

plotCI(NA,NA, xlim=c(.9,2.1), ylim=c(0,11), bty="L") ## This initializes the plot
for (i in 1:length(x1)) {
### First plot the lines (you can plot after, but it causes a small visual bug when lines are plotted on top of points)
# If loops are used to set the criteria for line coloration, easy to modify however you see fit
if(!is.na(y1[i]) & !is.na(y2[i])) {
	if(y1[i] >= 4.0 && y2[i] >= 4.0) {
 	 lines(x=c(x1[i],x2[i]), y=c(y1[i],y2[i]), lty="solid", col="grey") # No major change in LOD score
 	 greycount=greycount+1
 	 }
	else if(y1[i] >= 4.0 && y2[i] >= 2.0) {
 	 lines(x=c(x1[i],x2[i]), y=c(y1[i],y2[i]), lty="solid", col="red")
 	 redcount=redcount+1
 	 }
	else if(y2[i] >= 4.0 && y1[i] >= 2.0) {
 	 lines(x=c(x1[i],x2[i]), y=c(y1[i],y2[i]), lty="solid", col="green")
 	 greencount=greencount+1
 	 } 	 
	else if(y1[i] >= 4.0 && y2[i] < 2.0) {
 	 lines(x=c(x1[i],x2[i]), y=c(y1[i],y2[i]), lty="solid", col="skyblue")
 	 skybluecount=skybluecount+1
 	 } 	  	 
	else if(y2[i] >= 4.0 && y1[i] < 2.0) {
 	 lines(x=c(x1[i],x2[i]), y=c(y1[i],y2[i]), lty="solid", col="blue")
 	 bluecount=bluecount+1
 	 } 	  	 
}
 	
# now plot the actual points
  plotCI(x1[i],y1[i],lwd=2,pch=21,pt.bg="grey",col="black", add=T)
  plotCI(x2[i],y2[i],lwd=2,pch=21,pt.bg="grey",col="black", add=T)
}

greycount;redcount;greencount;skybluecount;bluecount

# Right Venn

plot(venneuler(c(CD=skybluecount, HFD=bluecount, "CD&HFD"=(greycount+redcount+greencount))))
# Note that this plot flips and puts the largest Venn on the left; so the figure is flipped in illustrator since HFD is larger, for visual consistency

#$$$$$$ #$$$$$$ #$$$$$$ #$$$$$$ #$$$$$$ #$$$$$$ #$$$$$$ #$$$$$$
#$$$$$$ #$$$$$$ #$$$$$$ #$$$$$$ #$$$$$$ #$$$$$$ #$$$$$$ #$$$$$$
# SUPPLEMENTAL PANEL 2A 

library(outliers)
library(ggplot2)
library(gplots)

setwd("~/Dropbox/Aging\ BXD\ Study/1\ _\ Manuscript\ -\ Evan\ Multi-Omic/")

mRNAdat_young=Top3772Data_young[1:3772,]
mRNAdat_old=Top3772Data_old[1:3772,]

Protdat_young=Top3772Data_young[3773:dim(Top3772Data_young)[1],]
Protdat_old=Top3772Data_old[3773:dim(Top3772Data_old)[1],]

## Plots LEFT panel
Age_FC_mRNA_All=rowMeans(mRNAdat_young,na.rm=T)-rowMeans(mRNAdat_old,na.rm=T)
Age_p_mRNA_All=NA

for(i in 1:length(Age_FC_mRNA_All)) {
	Age_p_mRNA_All[i]=t.test(mRNAdat_young[i,], mRNAdat_old[i,])$p.value
}
Age_padjust_mRNA_All=p.adjust(Age_p_mRNA_All,method="BH")

log2FC_rm=rm.outlier(Age_FC_mRNA_All, fill=T)	# Replace huge outliers with NA
pvalue=rm.outlier(Age_p_mRNA_All,fill=T) # Replace huge outliers with NA
neglog10p=-log(pvalue, 10)
plotCI(log2FC_rm, neglog10p, bty="L", pch=21, col="blue", pt.bg="black", ylim=c(0,22), xlim=c(-1.1,1.1), las=1)

length(which(Age_padjust_mRNA_All<0.05))		# Sets Bonferroni-corrected p-value as cutoff
length(which(Age_p_mRNA_All<0.05)) # Counts number of peptides affected at the nominal p-value

## Plots RIGHT panel
Age_FC_Protein_All=rowMeans(Protdat_young,na.rm=T)-rowMeans(Protdat_old,na.rm=T)
Age_p_Protein_All=NA

for(i in 1:length(Age_FC_Protein_All)) {
	Age_p_Protein_All[i]=t.test(Protdat_young[i,], Protdat_old[i,])$p.value
}
Age_padjust_Protein_All=p.adjust(Age_p_Protein_All,method="BH")

log2FC_rm=rm.outlier(Age_FC_Protein_All, fill=T)	# Replace huge outliers with NA
pvalue=rm.outlier(Age_p_Protein_All,fill=T) # Replace huge outliers with NA
neglog10p=-log(pvalue, 10)
plotCI(log2FC_rm, neglog10p, bty="L", pch=21, col="red", pt.bg="black", ylim=c(0,22), xlim=c(-1.1,1.1), las=1)

length(which(Age_padjust_Protein_All<0.05))		# Sets Bonferroni-corrected p-value as cutoff
length(which(Age_p_Protein_All<0.05)) # Counts number of peptides affected at the nominal p-value


#$$$$$$ #$$$$$$ #$$$$$$ #$$$$$$ #$$$$$$ #$$$$$$ #$$$$$$ #$$$$$$
#$$$$$$ #$$$$$$ #$$$$$$ #$$$$$$ #$$$$$$ #$$$$$$ #$$$$$$ #$$$$$$
# SUPPLEMENTAL PANEL 2B
library(vioplot)

VarExplained_Metab_Genotype=c(14.83,12.64,11.59,17.95,11,13.76,64.51,15.73,15.39,14.98,13.64,13.45,18.72,18.68,16.57,16.58,24.83,14.57,12.82,15.15,16.53,19.74,15.58,13.12,13.31,19.73,14.35,16.31,18.66,13.39,10.64,13.82,14.79,9.36,16.06,10.62,13.38,9.35,8.85,14.14,16.83,8.05,19.47,15.05,14.17,35.13,15.66,18.7,19.84,14.33,17.2,18.18,14.05,16.94,11.76,14.56,15.55,13.72,15.33,13.08,18.38,13.98,17.2,10.3,14.32,40.45,16.43,11.99,11.9,10.96,11.98,16.02,14.95,15.16,7.48,17.45,11.51,22.41,9.29,9.21,11.91,12.58,28.18,10.23,14.78,15.48,15.55,16.24,13.79,18.36,10.25,18.98,67.84,11.29,12.94,10.46,25.87,14.65,13.05,11.76,14.84,14.75,12.03,11.31,13.45,15.5,17.87,26.12,12.36,12.88,23.31,11.38,24.14,12.14,12.02,19.08,10.52,18.59,13.96,30.76,14.57,14.79,11.97,10.97,19.95,14.72,14.48,14.73,11.18,16.58,22.12,11.87,20.11,12.27,32.89,19.41,14.83,18.46,22.74,16.94,14.59,11.76,13.91,11.7,11.38,12.16,14.76,16.41,9.65,16.1,13.87,9.63,8.32,16.71,16.33,13.12,8.11,21.47,18.37,16.05,14.87,12.32,12.71,18.04,16.13,15.52,14.19,16.78,13.22,8.64,15.38,13.08,12.1,11.72,20.01,11.69,13.91,12.89,14.67,9.33,16.34,22.23,8.31,14.1,11.49,11.63,13.9,12.19,15.05,15.35,14.48,14.82,14.24,10.46,13.98,12.2,12.58,9.28,10.22,12.93,11.66,11.87,12.05,12.83,8.14,35.56,18.68,15.36,13.52,13.09,12.18,12,17.34,18.84,10.04,18.97,18.61,10.17,17.68,17.14,13.76,9.65,13.77,23.52,15.16,23.04,12.02,11.13,9.35,12.99,20.23,8.8,14.07,29.35,16.2,33.19,8.15,9.38,17.08,11.48,11.91,8.56,10.56,12.35,12.37,18.14,11.68,10.23,14.53,14.21,14.93,23.88,11.79,13.52,13.8,9.16,11.01,11.87,18.66,13.18,16.32,9.78,11.91,10.87,12.64,11.89,10.88,10.95,21.41,18.51,17.44,19.94,11.36,9.8,18.48,9.22,15.05,14.75,20.27,16.45,13.22,14.69,10.84,16.15,14.25,12.65,11.12,12.88,14.87,15.16,15.23,13.27,15.76,11.62,9.6,12.9,12.86,16.72,10.61,12.29,11.06,10.12,15.88,14.99,14.92,14.43,12.07,8.88,13.59,10.83,14.62,12.45,11.78,9.47,8.48,19.51,14.42,10.43,13.53,18.12,18.25,13.24,9.79,12.8,17.66,9.62,11.13,17.47,15.91,13.21,9.42,14.83,11.35,11.96,10.54,10.12,9.68,10.76,12.03,13.54,10.11,11.3,10.84,12.43,13.58,11.59,11.25,11.65,9.51,10.98,11.78,11.92,6.71,16.9,15.39,17.02,11.99,26.39,15.27,16.27,78.46,17.85,18.53,13.04,16.58,21.07,23.81,19.21,21.82,11.69,25.78,12.21,15.81,12.48,19.08,15.26,18.06,14.87,14.1,16.12,26.25,13.16,12.22,18.12,16.09,10.95,12.11,8.29,13.01,13.14,12.56,11.68,10.51,12.68,15.21,26.09,13.02,12.23,24.61,12.87,12.68,60.17,14.15,21.15,22.09,19.86,12.54,16.75,6.72,10.83,8.38,16.49,10.83,16.08,15.3,11.64,16.71,15.89,11.05,17.69,65.89,8.9,15.63,16.75,10.59,10.45,13.84,19.33,14.98,13.6,12.09,19.63,24.81,16.26,19.54,26.37,15.14,18.12,10.35,24,14.38,39.11,14.82,18.44,20.7,12.44,14.42,17.12,11.63,16.23,17.97,11.38,14.61,22.1,80.9,19.78,12.44,10.8,34.18,15.07,17.43,23.24,19.43,15.01,23.22,17.04,16.36,13.24,23.59,17.45,19,33.81,18.44,16.29,12.34,13.66,23.04,29.66,17.06,16.76,17.7,29.61,8.86,31.25,13.82,21.57,30.77,21,15.2,14.64,15.55,16.31,18.19,18.34,11.61,23.11,8.94,17.04,12.65,15.22,25.59,12.95,20.4,22.84,12.25,41.18,18.6,15.9,16.04,20.75,24.83,18.12,31.24,10.33,16.33,17.6,13.69,13.75,16.23,16.77,55.39,14.04,17.64,18.04,12.12,18.84,13.38,15.79,10.08,12.71,18.05,32.04,21.55,14.84,20.01,16.87,13.67,31.65,18.36,16.62,20.87,27.37,13.61,14.59,14.65,18.17,19.81,10.57,14.62,11.98,14,17.78,23.05,16.82,16.63,14.43,14.09,17.12,18.52,17.8,20.26,13.86,18.34,19.16,14.96,15.68,13.91,13.67,15.03,18.94,20.19,16.38,22.56,19.85,17.4,10.42,12.47,23.78,16.14,15.11,13.75,9.45,14.32,15.03,13.44,13.16,16.02,12.54,12.19,12.65,19.75,63.18,11,15.49,16.01,15.8,12.97,17.88,11.1,19.36,14.59,18.33,16.69,13.97,13.94,11.65,12.83,17.21,22.56,13.08,29.28,15.24,17.49,14.27,19.98,16.29,16.44,16.87,18.04,23.36,11.56,14.59,12.26,10.27,13.59,15.11,15.12,14.8,10.67,15.09,18.45,11.46,12.8,17.01,20.27,10.89,9.4,13.74,19.08,14.64,23.79,14.78,14.33,14.41,13.61,15.23,15.57,17.62,12.8,15.89,19.54,15.39,15.15,11.93,14.64,18.1,16.27,11.22,22.47,13.68,20.72,26.94,20.39,8.83,22.91,14.23,18.94,25.13,15.46,27.15,15.24,12.37,13.07,11.37,12.4,15.26,33.51,17.92,20.28,17.89,10.35,13.39,17.29,13.66,13.97,13.02,12.91,14.93,14.59,12.29,13.58,13.67,19.72,16.75,19.51,19.02,15.23,10.15,13,11.59,13.44,20.6,17.01,14.76,20.11,12.29,13.47,13.61,13.98,13.53,12.92,11.24,11.94,15.18,18.6,17.51,18.36,11.06,13.73,11.63,12.2,14.32,24.18,11.2,12.03,12.1,14.27,18.89,13.58,11.95,12.74,13.15,14.22,12.68,14.83,15.43,11.37,14.89,18.38,19.02,14.57,13.61,17.01,14.09,15.84,11.99,16.25,22.14,13.75,17.49,16,13.53,15.23,18,15.34,11.38)
VarExplained_Metab_Diet=c(2.63,3.57,1.69,5.42,0.02,5.64,1.22,0.05,0.46,0.07,8.52,0.48,0.97,0.4,0.49,0.4,0.08,23.35,0.17,0.04,0.65,0,0.01,9.63,2.91,1.75,0.49,1.46,0.56,0.56,1.05,10.83,1.01,1.5,0.01,0.01,3.14,0.09,0.18,0.95,0.74,2.41,1.5,0.38,0.24,0.01,1.73,3.24,2.98,7.66,1.12,1.62,0.05,0.11,0.03,0,5.18,12.06,6.94,23.48,3.53,0.35,0.05,0.54,2.03,0.93,0.16,9.63,18.96,12.76,0.36,0.02,0,0,3.97,0.09,3.17,0.98,0.03,3.22,10.17,21.86,0.99,17.38,35.6,0.73,0.5,0.06,0,2.25,2.55,0.38,0.25,0.41,0.49,0.92,0.17,5.87,0.02,7.33,1.66,0.19,8.86,1.26,0.81,0.28,2.41,0.03,14.07,1.48,1.45,0.69,0.01,1.11,1.9,2.48,3.44,0.15,1.73,0.03,0.44,30.11,0.04,3.13,0.16,0.01,9.8,6.89,7.99,0.51,10.95,0.71,0.03,0.87,0,0.11,1.86,0.1,0.66,0,2.87,0,6.45,0.23,6.67,4.59,0.6,1.28,1.51,0.02,2.26,22.14,3.94,16.44,1.95,1.66,1.44,8.95,1.01,0.16,0,1.86,4.43,8.4,6.16,5.58,0.66,0.07,2.65,11.41,3.52,1.53,22.38,6.19,0.08,5.54,3.15,0,17.03,8.93,0.69,1.88,10.65,0.3,4.48,7.05,1.4,1.72,0.14,0,0.03,0.02,0.37,16.63,0.55,0.7,5.66,14.19,23.29,1.65,7.42,0.01,47.47,21.48,1.17,0.27,0.1,1.83,0.74,0.03,15.44,10.44,0.06,7.8,7.12,11.34,0,7.64,0.11,6.25,0.13,0.14,0.01,0.01,0.01,0,17.62,10.15,0.04,19.97,3.46,0.02,22.76,0.28,0.25,1.79,9.33,34.51,1.55,12.86,9.7,6.37,0.12,0.56,6.11,4.59,0.02,5.84,6.05,0.56,0.06,5.8,1.86,0.81,0.2,4.95,1.53,2.91,1.26,0.19,0.59,10.02,7.3,4.97,1.76,0.36,2.85,6.6,0,1.88,1.77,1.82,2.4,19.78,0.01,17.92,1.02,2.08,0.22,6.07,0.15,0.71,0.34,0.68,0.41,0,6.28,0.4,2.9,0.62,1.5,0.27,0.34,0.01,1.13,6.65,0.18,0.34,0.08,0.23,1.04,0.11,0.02,1.22,2.04,0.27,2.98,24.14,3.93,0.66,0.34,12.93,4.66,20.36,27.38,0.05,1.11,0.21,0.03,0.1,0,3.15,0.07,0,0.04,1.28,0,0,0.97,0.23,7.71,0.28,3.09,5.29,0.02,0.06,1.12,9.52,7.45,6.17,3.97,0.26,0.61,0.03,0.62,0.3,5.63,0.79,6.06,0.26,0,0.04,2.69,1.53,10.03,9.06,1.49,1.37,0.03,16.51,1.12,0.19,0.55,0.02,0.65,0.57,0.39,0.11,0.04,0.25,0.3,36.16,0.27,0.03,0.92,2.32,0.65,17.87,0.11,0.18,3.5,0.15,1.12,0.08,0.96,0.24,0.49,0.24,0.02,0.15,6.37,0.91,0.01,0.02,0.68,0.96,0.13,1.4,0.37,0.01,0.02,0.46,0.85,5.77,3.19,1.5,0.01,3.19,0.02,0.2,0.74,8.31,1,29.15,31.35,0,1.78,0.99,0.19,3.98,1.85,1.42,0.11,22.24,26.86,14.2,0.26,0.48,3.7,0,3.45,10.8,1.17,1.09,1.46,1.98,0.67,35.81,1.06,11.86,22.82,0.99,0.76,39.77,0.02,6.17,15.65,3.17,2.86,0.01,0.61,6.49,0.35,1.55,0.04,0.9,0,0.12,0.04,38.38,0.75,2.07,4.66,0.97,10.54,29.37,6.17,1.23,9.72,2.28,2.62,0.13,0.15,0.1,15.75,2.36,0.61,0.02,0.26,2.23,0.74,9.76,0.65,0.62,2.76,22.1,0.02,0,28.5,0.22,5.95,1.15,0.68,15.04,12.84,7.47,2.38,10.33,6.54,0.03,22.07,0.19,3.14,0.83,2.9,0.06,0,0.53,2.2,0.93,0,0.59,4.09,0.16,0.02,6.44,2.91,1.57,1.55,1.28,2.36,0.2,1.99,2.3,1.79,3.36,11.65,9.68,2.23,2.89,21.15,0.01,0.75,0.62,16.52,0.66,5.21,0.8,0.24,0.03,11.05,0.23,15.17,8.54,1.66,0.24,4.03,3.89,0.53,1.05,18.8,18.77,0.79,0.22,5.7,0.34,1.12,19.09,5.74,5.06,0.15,2.34,5.54,9.66,2.92,9.91,0.15,1.52,1.54,0.1,0.05,0,0.41,0.01,10.43,1.2,31.48,4.34,0.14,5.26,5.57,4.11,0.97,33.68,15.08,0.71,1.3,0.01,56.23,19.16,0.35,0.09,1.12,2.09,1.7,0.01,22.39,5.79,3.24,7.55,6.27,19.75,0.04,0.54,0.14,0,1.56,1.41,0.57,0,0.01,0.4,28.91,21.21,0.03,12.26,7.82,0.25,21.36,3.21,0.74,27.31,0.54,6.1,42.25,1.02,6.25,0.01,7.09,7.23,6.22,4.5,4.46,1.36,6.93,0.06,6.28,0.04,7.54,0.14,6.48,2.75,0.8,7.52,0.2,4.86,0.19,1.3,25.27,0.02,0.19,5.81,9.89,2.62,0.51,11.23,0.06,6.07,0.26,2.49,1.76,3.44,0,2.57,0.15,17.22,15.73,3.93,1.69,3.27,0.57,0.66,0.04,2.82,6.1,4.24,4.68,9.46,1.72,0.53,12.42,0.19,0.29,0.22,0.3,8.68,1.84,0.29,0.55,7.51,24.51,1.07,3.25,0.03,7.67,2.77,0.5,4.3,13.35,24.31,2.64,0.07,2.28,2.98,0,0.22,0.14,6.7,0.1,0.06,1.25,0.73,2.07,2.26,0.96,0.17,0.87,9.57,0.06,7.48,2,4.16,1.35,0.2,1.06,37.07,26.04,0.33,0,9.34,20.91,26.28,3.36,18.43,1.01,1.81,0.62,1.61,1.47,1.73,0,1.82,0.14,16.33,0,7.27,22.1,0.74,0,0.32,4.08,5.63,0.52,6.43,8.56,4.82)
VarExplained_Metab_Age=c(0.24,0.02,0,0.42,0.09,0.11,0.1,1.84,1.3,0.02,0.3,0.09,2.28,0.83,0.04,0.21,0,0.24,0.55,1.89,0.05,0.13,0.05,0.01,0.13,0.44,0,0.2,0,1.07,1.57,0.18,0.23,0.78,0.88,0.01,0.11,0.01,0.04,0,0.03,0.43,0.48,0.66,0.44,0.05,0.23,0.71,0.94,0.01,0.55,0,1.63,0.26,0.06,1.24,2.79,0.19,0.11,0.56,0.07,0.59,0.12,1.25,1.76,0.04,1.57,0.48,0.47,0.49,0.22,0.45,0.03,0.33,0.14,0.66,0.04,0.15,0.03,0.13,0.04,0,0.01,0.02,1.77,2.62,0.45,2.89,0.63,0.34,0.84,0.02,0.02,4.69,0.01,0.26,0.46,0.59,0.36,0.3,0.02,1.37,1.06,0.66,0.92,0.63,2.96,0.01,0.04,1.21,0.15,3.76,0,0.1,0.13,0.25,0.01,1.48,0.43,0.1,0.57,0.72,0.2,0.37,1.21,0.09,0.14,0.29,0.38,0.06,0.37,0.09,0.32,0.02,2.73,0.08,0.05,3.02,0.04,0.52,1.25,0.33,0.61,4.93,0,0.19,0.02,1.46,0.19,0.09,2.07,0.17,2.34,1.67,0.09,0.02,0.04,0.62,0.42,0,0.09,0.04,0.01,0.11,0.28,0.08,0.54,0.6,0.94,0.05,0.05,0,0.06,0.18,1.97,0.02,0.04,0.62,0,0.31,0.01,0.01,0.05,0.82,0.03,0.12,0.58,0.06,0.21,0.28,0.13,0.01,0.02,0.54,0.09,2.09,0.08,0.2,0.5,0,0.02,0.25,0.07,0.09,6.19,0.59,1.44,0.21,0.18,0.69,0.01,0.3,1.27,0.49,0.08,0.04,0.03,0.45,0.23,0.42,0.2,5.52,10.73,1.92,0,3.62,0.26,0.6,0.19,0.28,0.59,4.53,0.19,1.85,0.15,3.02,0.94,0.17,1.6,0.67,0,0.16,1.52,1.25,0.06,0.18,0.04,0.62,0.01,0.32,0.02,0.01,0.23,5.81,0.82,0.47,0.88,1.04,0.17,2.6,0.03,0.06,0.02,1.41,0.13,0.17,0.19,0.01,0,0.01,0.05,0.99,0.5,0.04,1.13,0.12,0.18,0.01,0.13,0.07,0.54,0.09,0.46,0.05,1.79,1.04,0.12,1.13,0.04,0.15,0.03,0,0.03,1.17,1,0.04,0.09,0.12,0.97,1.51,2.03,0.13,0.1,0,0.26,0.53,0.08,1.4,0.59,1.27,1.28,2.49,4.43,1.66,0.43,0.05,0.56,0.05,0.01,0.25,0.97,0.74,0.03,0.11,0.03,0.35,0,2.3,1.63,0.07,0.37,0.01,0.19,0,0.12,0.12,0.88,0.17,0.05,0.32,0.59,0.23,4.45,0.34,0.46,0.61,0.02,0.04,0.04,0.14,0,0.32,0.01,0.64,0.01,0.57,0.03,0.56,0.25,0.06,0.15,2.04,1.88,0.03,0.4,4.22,0.78,1.74,0.3,0,0.08,0,1.18,1.16,0.17,0.3,0.01,0.16,0,0.32,0.21,0.35,0.15,0.27,1.62,0.43,0.16,0.05,1.07,0.02,0.03,0.04,0.25,0.08,0.44,0.32,0.22,0.2,1.68,0,0.1,0.47,0.38,2.92,2.18,0.23,0.9,0.64,0.46,0.18,0,5.52,0.42,1.06,0.06,0,0.28,0.14,0.29,5.34,0.19,1.4,1.1,0.07,0.02,0.07,0.79,0.02,0.36,0.7,0,0.14,0.2,0.23,0.32,1.01,0.1,0.03,0,0.17,0,0.06,0.13,1.81,2.05,0.01,0.61,0.7,0.16,2.17,1.13,0.42,0.03,0.89,0.01,4.14,0.15,1.21,0.82,0.34,2.74,3.3,0.78,0.09,0.46,0.33,0.09,0.03,0.12,1.05,3.49,0,0,1.39,0.16,4.54,0,0.03,0.56,1.22,0.08,0.1,0.02,0.8,0.04,0,0,0.61,0.3,0.27,0.06,2.25,0.1,0.72,0.1,0.03,0,0.63,0.13,0.91,0.03,0.05,3.09,2.63,0.2,2.85,0.14,0.63,0.05,1.38,0.33,0.22,0.31,0,4.89,0,0.01,0.01,0.59,0.6,3.43,0.72,1.2,1.39,0.34,0.24,0,0.14,0.02,3.85,1.78,0,0.2,0.35,0,1,0.02,2.17,3.07,0,0.02,0.01,0.15,0.43,1.07,2.27,1.11,0.5,0.91,3.53,0.12,0.09,0.71,0.15,0.36,0.07,0.32,0.65,1.2,0.08,0.76,0.91,0.89,0.04,1.31,0,0.02,1.69,0.33,0.11,0.64,0.7,0,0.66,1.9,0.07,0.25,0.8,0.25,0.04,0.02,0.44,0,0.06,0.13,0.33,1.51,0.15,0,0,1.98,0.15,0.03,0.79,0.27,0.02,0.08,0,0.21,0.43,0,0.4,0.93,0.02,0,0,0,0.35,5.78,0.6,1.78,0,0.08,0.2,0.89,0.03,0.04,0.02,0.9,0.83,0.07,0.11,0.16,0.23,0.03,0.08,0.02,0.02,0.06,0.63,0.2,0,0,0.8,0.08,0.41,0.19,0.98,2.42,0.2,0.07,0.23,0.01,1.76,2.82,0.34,0.02,0.09,0.37,0.1,1.14,0.95,1.23,0.68,3.35,3.01,1.26,0.41,0,0.52,1.69,2.15,0.47,0.93,1.88,0.2,0.5,0.02,0.15,0.01,2.01,0.42,0.51,0.64,1.28,1.46,0.81,0.54,0.25,0.1,0.41,0.01,1.56,1.56,0.47,2.31,0.48,3.52,2.8,1.97,1.11,0.76,0.13,0.55,0.14,0.23,1.23,1.61,2.51,1.19,1.02,1.19,1.46,0.11,0.31,0.07,0.07,1.97,3.16,0.02,1.5,0.01,0.71,0.16,0.14,0.55,0.14,0.16,0.2,0.43,1.49,3.9,4.49,4.14,0.83,0.04,0.47,6.08,0.86,5.34,0.21,0.27,0.16,1.03,1.62,0.04,2.28,3.85,0.13,0.01,0,0.47,0.02,0.47,0.01,5.45,0.38,0.04,0.43,3.34,0.8,3.47,0.06,0.76,0.22,0.37)
VarExplained_Metab_Interaction=c(26.3,28.09,25.75,22.19,22.6,23.42,11.21,20.38,21.83,22.77,22.39,22.5,22.55,22.05,33.72,18.61,25.06,17.42,21.41,29.37,24.23,31.21,29.04,24.31,25.24,22.59,30.26,30.68,27.91,22.8,22.31,26.47,23.22,23.93,26.86,25.73,28.25,27.16,25.79,25.1,26.67,23.12,24.01,24.17,30.2,25.19,26.89,25.02,27.95,25.71,22.15,23.11,28.48,24.67,24.35,24.08,23.11,19.25,21.41,18.63,25.64,20.36,27.77,23.42,23.97,17.98,29.68,27.3,19.3,20.37,23.31,27.24,23.49,30.02,25.07,23.58,21.5,17.32,26.14,23.92,17.91,22.31,17.37,20.09,17.33,28.78,28.3,25.78,23.55,23.37,22.83,24.15,10.41,30.69,17.28,21.8,20.53,26.68,24.88,23.54,23.58,27.35,23.61,19.39,24.41,28.82,24.32,26.68,17,25.32,22.92,23.98,28.45,29.02,23.91,24.86,19.02,25.31,25.32,15.84,23.57,14.12,20.9,26.29,25.83,21.15,21.91,26.81,19.22,19.81,25.52,23.63,29.2,22.91,21.3,28.98,29.6,21.33,25.49,21.59,28,23.13,22.72,27.4,25.67,20.72,23.46,21.05,26.14,24.83,26.01,18.21,21.38,27.14,40.82,24.39,24.94,26.07,19.95,23.64,26.16,21.65,25.33,20.08,23.47,25.81,23.23,26.99,25.02,22.5,20.71,24.54,23.72,24.66,22.55,29.97,26.78,19,20.34,24.66,34.16,22.47,21.77,26.15,29.68,28.35,24.66,24.93,19.7,21.95,22.33,26.84,25.23,21.63,23.02,24.4,26.06,21.76,20.58,30.78,25.93,29.32,13.44,17.6,27.98,21.87,22.98,24.67,29.8,25.25,22.83,26.29,23.2,20.09,28.4,17.79,26.81,32.56,27.71,20.9,20.93,25.79,22.3,24.63,19.62,19.29,19.96,25.48,25.01,21.24,18.25,26.66,23.2,22.08,26.8,19.04,20.46,17.94,26.02,22.59,23.93,22.4,22.58,23.75,22.41,23.15,21.82,21.49,21.56,22.94,23.73,22.76,25.1,22.07,22.74,24.68,23.32,26.5,22.95,25.84,25.46,24.58,24.11,23.27,21.42,27.98,25.66,25.61,27.77,28.82,24.28,25.23,24.74,19.46,24.93,19.64,23.96,27.53,23.37,22.61,19.64,25.51,23.91,24.76,24.83,19.96,27.64,23.97,26.72,22.82,26.57,24.33,26.69,21.86,23.82,26.33,24.08,23.9,26.57,24.89,31.33,21.28,29.09,22.68,28.87,27.15,25.89,20.91,22.22,26.94,21.56,19.17,25.02,19.99,20.01,23.4,29.35,21.95,22.85,28.81,25.75,23.48,22.1,24,22.95,23.68,23.41,28.03,24.76,28.39,24.61,18.44,25.1,21.82,27.5,27.22,25.42,22.24,23.39,22.83,25.16,25.51,25.58,25.09,24.58,25.73,25.84,23.15,26.12,23.88,25.83,25.07,24.62,20.4,28.42,26.19,21.8,21.52,23.42,26.82,5.4,20.55,24.43,30.04,36.9,19.08,19.64,22.48,30.96,21.57,26.93,15.86,22.97,28.14,25.7,29.14,26.41,23.96,31.23,22.04,16.46,29.52,24.96,29.58,21.3,22.52,22,17.02,25.53,26.02,29.59,19.27,21.94,19.39,19.96,25,17.33,24.76,23.65,24.89,29.09,17.1,28.86,22.21,25.85,24.34,31.53,32.28,17.59,22.07,25.54,19.44,20.87,15.01,14.47,23.17,23.56,26.27,26.99,20.97,9.5,22.39,29.6,22.44,17.25,23.26,31.14,31.34,20.55,33.04,23.38,21.85,23.09,24.69,24.72,15.44,26.78,18.32,22.81,20.28,22.02,17.68,23.15,15.73,29.81,19.78,21.58,25.82,19.93,28.92,19.89,16.8,25.28,21.96,5.35,27.59,22.73,26.91,18.8,19.75,22.38,29.92,23.53,26.03,38.95,15.54,25.32,19.2,24.43,29.64,21.04,25.81,18.59,24.07,19.55,19.34,20.12,22.52,22.75,24.01,27.12,20.71,21.93,23.57,23.65,18.51,18.77,23.35,18.39,21.66,26.2,24.42,26.56,18.22,21.75,20.13,21.53,22.45,20.75,22.3,19.62,27.05,20.28,33.72,24.07,17.72,31.63,42.33,23.97,25.98,19.61,20.34,21.06,21.79,26.04,24.06,22.36,22.31,22.09,19.64,13.97,26.34,26.45,21.35,29.62,20.06,21.93,21.25,23.81,25.39,24.82,20.48,46.17,28.11,18.63,17.41,29.55,19.1,23.45,28.07,23.06,21.45,27.47,28.93,30.67,27.96,20.59,29.16,24.04,24.59,18.73,24.28,22.06,23.22,21.85,35.73,29.08,25.56,25.19,26.86,28.7,25.22,28.64,24.18,27.11,31.85,27.67,32.47,30.54,22.77,27.08,20.58,23.82,26.26,22.64,22.71,19.4,25.04,28.48,24,24.22,23.67,32.69,20.41,27.35,34.96,26.69,33.37,10.14,19.43,30.95,10.94,22.49,31.38,26.79,28.15,24.47,25.14,21.65,21.02,31.06,19.27,22.02,26.72,23.38,22.79,23.76,36.43,26.82,23.92,23.17,27.63,20.39,17.5,22.86,24.52,26.81,25.96,21.06,21.34,31.73,26.34,27.46,24.19,16.26,24.59,31.83,28.46,26.89,19.78,29.88,15.14,35.15,25.96,21,21.77,18.39,25.09,19.74,23.14,18.26,22.79,24.8,28.21,23.46,20.51,33.44,25.75,18.77,20.92,22.16,24.36,25,22.64,25.47,21.03,26.14,28.6,29.62,33.28,22.61,18.95,26.27,24.55,24.09,21.92,20.93,24.45,25.41,26.8,24.43,27.12,22.17,27.2,26.68,24.78,21.01,22.78,20.12,31.2,24.47,25.72,25.13,21.06,20.74,23,26.9,23.71,26.94,23.39,19.14,23.36,25.01,35.49,21.79,28.06,29.61,23.54,23.67,19.12,23.5,25.42,28.05,19.81,41.89,21.93,20.19,23,27.2,21.15,22.78,24.21,25.41,21.45,21.18,30.93,25.4,23.07,26.57,23.96,24.98,21.03,19.23,22.56,23.13,17.88,22.22,23.25,24.86,24.01,20.92,21.46,28.72,23.42,27.81,24.69,26.16,26.16,25.71,25.46,28.23,24.24,25.84,22.74,25.22,22.84,21.12,25.47,22.72,17.57,23.5,24.27,24.76,21.67,23.07,21.87)
VarExplained_Metab_Residual=c(56.01,55.69,60.97,54.03,66.29,57.08,22.96,62,61.02,62.17,55.14,63.49,55.49,58.04,49.17,64.2,50.03,44.42,65.05,53.55,58.54,48.92,55.32,52.93,58.42,55.49,54.9,51.35,52.87,62.18,64.43,48.7,60.76,64.43,56.2,63.63,55.11,63.38,65.14,59.81,55.73,66,54.54,59.75,54.95,39.62,55.48,52.33,48.3,52.28,58.98,57.1,55.79,58.02,63.8,60.12,53.37,54.78,56.21,44.24,52.38,64.72,54.85,64.49,57.92,40.6,52.16,50.6,49.37,55.43,64.12,56.27,61.52,54.49,63.34,58.22,63.77,59.14,64.5,63.52,59.97,43.25,53.44,52.28,30.52,52.39,55.21,55.04,62.03,55.67,63.53,56.48,21.48,52.93,69.27,66.56,52.97,52.22,61.69,57.07,59.9,56.34,54.43,67.38,60.41,54.78,52.45,47.15,56.54,59.1,52.17,60.19,47.4,57.64,62.05,53.33,67.02,54.47,58.55,53.28,60.85,40.26,66.9,59.24,52.85,64.04,53.66,51.28,61.24,63.03,41.05,63.69,50.34,63.92,43.08,51.42,53.67,57.1,51.06,60.95,53.3,64.79,56.31,55.74,56.28,62.34,61.15,59.8,62.51,58.96,55.79,49.84,64.02,38.04,40.81,60.81,65.48,42.89,60.24,60.14,58.89,64.14,57.52,53.36,53.97,53,61.38,55.57,58.17,57.4,60.35,60.85,41.73,57.25,55.39,52.78,56.12,67.48,47.96,56.78,48.79,53.42,59.22,58.62,54.32,52.85,59.46,61.1,64.9,62.41,63.03,58.3,60.13,50.74,62.36,60.6,55.62,54.56,45.42,54.63,54.98,58.55,26.96,48,56.53,41.71,56.8,57.94,55.75,60.93,49.55,50.97,58.13,52.77,54.36,51.85,54.55,49.17,54.27,55.29,64.98,58.9,53.19,49.93,65.21,54.05,50.15,52.65,65.4,45.52,57.48,60,39.78,46.44,56.6,42.95,61.12,38,53.74,52.41,54.46,62.52,65.22,62.09,59.05,53.95,66.45,61.82,57.85,61.97,61.25,47.54,61.02,57.78,62.44,60.73,63.26,57.67,56.96,58.2,57.6,55.56,56.66,59.48,64.06,59.59,60.43,56.82,50.82,50.78,56.47,52.02,61,50.92,55.46,53.1,59.79,55.63,56.01,54.8,66.46,58.99,64.46,58.35,58.72,66.35,54.84,61.63,55.47,61.25,56.67,62.13,57.17,65.34,64.45,54.09,62.78,58.91,61.76,61.08,54.54,68.36,54.92,61.1,53.91,57.62,58.99,44.66,59.67,60.3,62.2,52.96,54.1,48.52,43.69,56.99,54.56,67.37,63.58,52.72,55.03,59.4,68.01,63.09,59.31,65.07,65.45,52.19,56.72,58.09,57.89,66.45,60.27,60.93,61.82,62.48,62.9,57.31,57.08,57.14,60.17,62.69,58.52,62.12,60.76,61.77,57.26,64.37,58.27,64.74,62.39,62.66,65.97,60.53,46.15,47.17,64.69,50.15,61.03,40.34,14.87,59.38,54.61,56.86,45.47,55.06,55.39,56.47,46.88,66.48,46.92,35.77,59.77,58.19,54.13,52.98,54.88,43.14,54.56,61.35,53.58,56.82,61.56,51.95,60.04,65.86,65.24,74.4,60.39,60.67,51.45,68.1,67.29,67.83,63.72,47.63,69.3,61.41,49.69,62.23,58.12,21.8,55.77,47.94,46.69,54.06,55.02,47.14,75.21,66.71,65.34,50.24,66.88,38.7,38.81,65.19,57.67,56.71,61.48,52.03,22.58,65.88,53.56,38.49,45.27,52.02,53.97,48.83,60.4,52.67,61.08,47.58,50.72,57.73,53.97,55.2,57.31,27.73,65.78,43.68,40.78,42.16,61.15,24.24,47.42,61.6,47.75,53.19,65.43,52.66,60.4,64.91,59.73,53.51,13.7,47.59,64.67,60.96,46.15,26.46,56.69,41.47,51.6,57.9,26.82,37.72,52.07,66.3,42.14,49.57,53.85,40.24,62.81,58.14,52.2,60.1,56.22,47.77,59.37,55.79,54.35,39.82,68.53,43.75,59.72,37.82,50.43,55.04,37.61,63.21,52.24,55.86,54.47,47.68,53.71,49.26,67.15,49.55,59.93,61.53,32.68,59.76,53.09,39.97,60.58,38.18,49.63,40.6,57.74,50.97,55.23,60.73,43.3,67.72,52.72,51.9,61.03,62.36,59.55,61.72,24.85,58.7,52.72,56.92,56.13,57.49,53.04,53.14,63.86,55.17,34.2,47.47,31.32,56.09,44.84,64.06,51.56,46.28,54.89,55.28,45.01,50.93,43.59,47.51,51.96,51.35,54.45,55.87,59.89,58.85,48.35,39.08,53.4,59.59,55.46,49.44,55.39,37.57,49.35,50.21,50.13,57.67,46.59,46.95,53.7,42.56,58.24,50.65,52.56,58.08,52.04,62.34,53.2,53.23,47.63,65.6,36.41,46.04,54.97,55.6,56.44,62.33,52.02,30.82,44,50.84,54.48,53.94,21.43,48.75,46.96,25.64,65.36,50.25,55.22,56.03,40.09,51.19,63.8,51.63,48.08,42.25,60.33,58.75,62.54,65.55,61.85,44.6,44.28,62.4,45.76,56.72,33.13,46.82,56.23,46.9,48.89,56.9,38.63,51.25,55.91,31.65,59.58,59.21,27.87,59.19,46.78,56.71,55.3,57.27,45.26,68.89,47.59,54.88,51.72,66.88,65.74,60.14,51.22,61.88,51.41,59.44,60.06,48.09,59.91,59.06,50.78,55.24,42.79,63.07,56.98,53.48,48.72,62.13,56.03,46.64,56.28,53.7,47.65,50.03,53.22,48.53,52.87,63.12,50.97,46.42,43.89,46.47,57.29,42.77,57.75,59.43,64.21,57.97,53.54,54.26,39.99,49.31,57.64,50.28,52.34,60.7,55.71,63.49,64.53,52.99,57.89,57.54,55.12,54.84,41.66,61.14,51.89,47.18,50.9,49.92,53.42,60.41,47.47,43.8,59.4,52.71,51.2,62.34,37.69,65.5,66.12,54.73,55.55,65.24,61.56,63.82,59.87,60.95,59.13,50.84,55.24,56.14,59.45,56.49,59.33,56.6,50.75,61.9,62.95,32.91,36.99,51.46,60.7,49.37,45.23,38.83,53.53,44.45,54.72,58.03,59.58,53.49,54.3,53.79,57.19,59.86,56.99,46.37,58.93,52.45,40.14,51.61,63.1,61.28,55.63,53.1,59.43,53.14,52.81,61.55)
VarExplained_Metab_SumofSquares=c(96.6,96.8,48.4,79.5,111,186.2,333.9,78.2,198.5,96.6,51.3,69.7,130.1,123.3,154.9,248.3,427.4,97.4,57.7,146.8,54.8,81.3,184.8,309,48.1,79.5,135.9,121.6,147.7,170.3,13.9,76.3,316.7,105.9,133.9,29.6,53.4,76.5,61.1,84.7,125,74.3,111,164.9,162.2,203.6,152,214.5,289.7,57.5,238.8,66,34.6,28.7,292.4,40.2,162.8,108.6,67.1,282.7,533.8,141.2,169.7,8.5,198.4,169,193.7,124.3,96.7,42.8,188.3,418.3,322.9,107.5,42.1,117.5,56.6,262.6,96.3,40.9,63.6,199.9,330.4,100.5,180.2,148.6,97.6,118.5,534.1,90.4,78.1,88.5,498.3,190.9,85.4,311.6,128.8,436.3,153.7,1845.2,225.8,67.7,115,30.8,106.3,165.5,151.4,305.6,109,187.7,76.2,125.6,540.9,138.3,76.5,129.1,190,206,424.2,332.3,118.1,115.5,165,243.2,99.1,60.3,98.3,224.4,184.3,39.7,398.3,566.2,201.9,48.7,257.2,135.8,91.9,138.8,213.7,74.1,64,157.1,56.6,164.2,56.3,133.4,51,201.7,122,184.7,111.6,178.8,116,411.7,263,504,123.3,149.6,348.2,345.4,524.2,135.1,79,418.2,529.4,762.8,789.9,418.8,453.7,237.3,47.5,375.6,170,109.9,294.8,141.3,241.4,155.9,824.8,109.2,323.9,267.9,115.8,135.7,761.2,264.1,840.6,741.8,142.5,1057.9,49.2,25.2,102.9,628.3,98.2,144.8,103.3,187.1,427.6,217.7,341.4,178,2060.9,294,512.8,293,97.6,1020.1,116,110.4,217.2,225.7,105.9,102.4,187.2,126.4,48.1,180.4,4092.1,101.4,653.6,1037.7,607.5,153.5,641.4,162.9,336.6,255.6,341.9,154.6,443.9,483.1,212.5,241.4,184.9,327.2,211.6,438.2,140,415.2,100.9,97.4,218.7,34.2,72.1,577.9,53.7,131.3,91.3,84.3,311.2,1826.2,202.8,94.3,91.5,124,91.4,222.4,317.4,380.2,129.6,148.9,99.5,204.8,142.2,184.9,326.7,120.7,234.1,203.8,180.3,710.9,108.3,125.9,1038.4,270.1,754,204.2,161,235.8,114.6,344.8,109.3,120.9,96.7,58,136.5,111.3,214.9,1138.7,468.1,375.2,384.2,188.5,58.8,213.2,44.9,54.6,442.1,192.7,508,216.2,189,339.7,479.6,157.9,257.1,238.6,107.9,76.7,164.3,94.9,99.4,98.9,224.1,191.7,197.6,185.7,112.1,341.4,200.3,84.3,455.4,317,135,639.2,218.5,261.4,380.5,1125,348.1,269.9,279.1,200.2,144.1,222.2,163.3,753.9,1067.4,344.3,880.9,1168.1,86.7,915.5,1326.3,877.5,362.5,97.7,205.7,46.7,59.1,222.5,124.8,62.5,41.9,51.6,79.6,69.9,42.7,83.3,500.6,71.3,111,53.1,65,98.4,22.3,105.6,103.4,49.5,365.7,70.6,29.6,144.6,25.3,45.4,183.1,74.3,32.8,43.8,47.7,107.3,127.9,93.1,169,9.3,72.5,233.2,168.5,22.8,42.6,40.7,13.8,26.2,53.8,78,42.4,47.4,93.3,98.9,115.8,172,102.8,158,236.1,51.8,278.9,38.7,181,42.4,8.1,147.5,36.2,323.4,187.2,283.8,103.1,118.1,5.6,150.2,304.7,246.1,151.2,224.6,68.4,27.5,321.8,197.6,130.6,88.4,46.6,35.2,58.8,144.1,44.8,230.1,63.7,81,30.1,49.4,169.9,372.5,12.8,229.6,104.8,170.3,69.1,103.9,54.4,46.6,139.7,43.2,42.9,91.6,300.9,130,39.5,126.7,95.5,168.7,74.5,189.2,61.8,95.8,172.1,234.9,96.6,14,50.3,115.5,87.7,261.1,122,54.9,29.5,196.9,49.4,438.6,59.4,158.6,72.4,86.4,480.2,129.6,53.4,115.9,162.4,90.8,91.4,130.3,197.3,66.9,27.9,76.3,68,82.3,110.3,161.6,78.7,37,297.8,325.8,94.5,131.6,14.4,221.9,109.4,45,60.1,68.6,72.3,61.6,48.6,89.6,94.9,79.4,56.9,204.9,95.9,94.5,175.7,182.9,48.8,91.3,60.6,79,33.4,48.5,44.8,82.9,242.5,73.4,204.4,406.6,87.5,260.2,45.6,281.3,116.7,437.5,60.5,333.8,504.8,844.3,829.4,166.5,428.3,56.9,258.7,74.7,94.1,87.8,201,502.4,101.5,155.8,48.6,111.3,287.1,79.8,267.1,88.5,115.4,71,58.7,699.5,40.9,657.4,599.3,106.7,195.1,34.8,31.9,163.9,174,38.4,61.2,62.3,91.9,47.5,58.9,84.3,139.7,180.7,260.8,102,205,78.7,1355.4,105.3,184.3,250.2,140,759,57.1,93.2,154,133.9,57.3,85.7,127.6,76.9,103.7,169.3,913.6,244.9,120.4,210,260.9,54.5,112.3,285.3,116.2,240.4,16.9,52.4,122.9,536.6,70.2,329.7,208.7,119,98.1,101.1,235,183,147.3,31.3,63.1,146.2,118.2,59,58.7,185.8,406.9,104.4,55.7,24.3,39.9,207.5,1577.1,103.3,43.7,112,21.2,119.5,47.7,237.2,83.2,28.8,54,108,149.4,59.1,121.3,98.2,107.3,50.6,205.1,60.7,187.1,457.2,47.1,181.4,852.4,106,225.4,333.6,30.7,159.5,188.3,32,183.4,105,46.4,113.2,145.7,142.6,1131.4,509.9,141.4,405.8,142.8,527.1,74.1,91.1,57.3,593.1,111,58.9,135.1,541,289.4,58.7,562.9,272.6,93.7,63,84.9,150.3,45.5,67.2,158.6,275.3,127.3,160.4,459.3,91.1,100.2,75.8,623.8,90.5,248.6,311.3,121.6,272.5,633.5,172.2,261.4,128.1,154.8,107.4,266.1,134.9,91.3,344.8,442.4,241.9,231.2,219.5,132.1,256.8,237.1,247.9,169.7,38.3,110.7,133.4,185.2,303.5,220,211.2,218.3,193.6,121.4,108.5,222.7,260.2,267,71.3,247.3,61.5,108.3,170.3,152.4,51.9)

vioplot(VarExplained_Metab_Genotype, VarExplained_Metab_Diet, VarExplained_Metab_Age, VarExplained_Metab_Interaction, VarExplained_Metab_Residual, col=c("lightgreen","green","forestgreen","darkgreen","black"), ylim=c(0,100))


#$$$$$$ #$$$$$$ #$$$$$$ #$$$$$$ #$$$$$$ #$$$$$$ #$$$$$$ #$$$$$$
#$$$$$$ #$$$$$$ #$$$$$$ #$$$$$$ #$$$$$$ #$$$$$$ #$$$$$$ #$$$$$$
# SUPPLEMENTAL PANEL 2C
library(LSD)
library(outliers)

# LEFT PANEL
#Diet_FC_mRNA_All # Loaded from panel 2A
#Diet_FC_Protein_All # Loaded from panel 2A

Xaxis=rm.outlier(as.numeric(as.character(Diet_FC_Protein_All, fill=T)))	# Replace huge outliers with NA
Yaxis=rm.outlier(as.numeric(as.character(Diet_FC_mRNA_All, fill=T)))	# Replace huge outliers with NA
heatscatter(Xaxis, Yaxis, cex=0.6, ylim=c(-.6,.6), xlim=c(-.6,.6))
abline(lm(Yaxis~Xaxis), lty=1, lwd=1, col="black")
cor(Xaxis, Yaxis, method=c("pearson"), use="complete.obs")
signif(summary(lm(Yaxis~Xaxis))$coefficients[8], digits=2)

# RIGHT PANEL
#Age_Correlation_mRNA_All # Loaded from panel 2C
#Age_Correlation_Protein_All # Loaded from panel 2C

Xaxis=rm.outlier(as.numeric(as.character(Age_Correlation_Protein_All, fill=T)))	# Replace huge outliers with NA
Yaxis=rm.outlier(as.numeric(as.character(Age_Correlation_mRNA_All, fill=T)))	# Replace huge outliers with NA
heatscatter(Xaxis, Yaxis, cex=0.6, ylim=c(-.6,.6), xlim=c(-.6,.6))
abline(lm(Yaxis~Xaxis), lty=1, lwd=1, col="black")
cor(Xaxis, Yaxis, method=c("pearson"), use="complete.obs")
signif(summary(lm(Yaxis~Xaxis))$coefficients[8], digits=2)


#$$$$$$ #$$$$$$ #$$$$$$ #$$$$$$ #$$$$$$ #$$$$$$ #$$$$$$ #$$$$$$
#$$$$$$ #$$$$$$ #$$$$$$ #$$$$$$ #$$$$$$ #$$$$$$ #$$$$$$ #$$$$$$
# SUPPLEMENTAL PANEL 2D
# We calculate here the p-values for the correlations for the mRNAs in every decile of variation 
Variance_mRNA_Top10pct=as.numeric(quantile(Variance_mRNA,prob=seq(0,1,length=11))[10])
Top10Variance_Correlation_pvalue_mRNA_Protein=NA

for(i in 1:dim(mRNAdat_All)[1]) {
	if(Variance_mRNA[i]>=Variance_mRNA_Top10pct){
		Top10Variance_Correlation_pvalue_mRNA_Protein[i]=cor.test(as.numeric(Protdat_All[i,]),as.numeric(mRNAdat_All[i,]),method="spearman")$p.value[1]
	}
}

Variance_mRNA_10_20pct=as.numeric(quantile(Variance_mRNA,prob=seq(0,1,length=11))[9])
Top10_20Variance_Correlation_pvalue_mRNA_Protein=NA

for(i in 1:dim(mRNAdat_All)[1]) {
	if(Variance_mRNA[i]<Variance_mRNA_Top10pct && Variance_mRNA[i]>=Variance_mRNA_10_20pct){
		Top10_20Variance_Correlation_pvalue_mRNA_Protein[i]=cor.test(as.numeric(Protdat_All[i,]),as.numeric(mRNAdat_All[i,]),method="spearman")$p.value[1]
	}
}

Variance_mRNA_20_30pct=as.numeric(quantile(Variance_mRNA,prob=seq(0,1,length=11))[8])
Top20_30Variance_Correlation_pvalue_mRNA_Protein=NA

for(i in 1:dim(mRNAdat_All)[1]) {
	if(Variance_mRNA[i]<Variance_mRNA_10_20pct && Variance_mRNA[i]>=Variance_mRNA_20_30pct){
		Top20_30Variance_Correlation_pvalue_mRNA_Protein[i]=cor.test(as.numeric(Protdat_All[i,]),as.numeric(mRNAdat_All[i,]),method="spearman")$p.value[1]
	}
}

Variance_mRNA_30_40pct=as.numeric(quantile(Variance_mRNA,prob=seq(0,1,length=11))[7])
Top30_40Variance_Correlation_pvalue_mRNA_Protein=NA

for(i in 1:dim(mRNAdat_All)[1]) {
	if(Variance_mRNA[i]<Variance_mRNA_20_30pct && Variance_mRNA[i]>=Variance_mRNA_30_40pct){
		Top30_40Variance_Correlation_pvalue_mRNA_Protein[i]=cor.test(as.numeric(Protdat_All[i,]),as.numeric(mRNAdat_All[i,]),method="spearman")$p.value[1]
	}
}

Variance_mRNA_40_50pct=as.numeric(quantile(Variance_mRNA,prob=seq(0,1,length=11))[6])
Top40_50Variance_Correlation_pvalue_mRNA_Protein=NA

for(i in 1:dim(mRNAdat_All)[1]) {
	if(Variance_mRNA[i]<Variance_mRNA_30_40pct && Variance_mRNA[i]>=Variance_mRNA_40_50pct){
		Top40_50Variance_Correlation_pvalue_mRNA_Protein[i]=cor.test(as.numeric(Protdat_All[i,]),as.numeric(mRNAdat_All[i,]),method="spearman")$p.value[1]
	}
}

Variance_mRNA_50_60pct=as.numeric(quantile(Variance_mRNA,prob=seq(0,1,length=11))[5])
Top50_60Variance_Correlation_pvalue_mRNA_Protein=NA

for(i in 1:dim(mRNAdat_All)[1]) {
	if(Variance_mRNA[i]<Variance_mRNA_40_50pct && Variance_mRNA[i]>=Variance_mRNA_50_60pct){
		Top50_60Variance_Correlation_pvalue_mRNA_Protein[i]=cor.test(as.numeric(Protdat_All[i,]),as.numeric(mRNAdat_All[i,]),method="spearman")$p.value[1]
	}
}

Variance_mRNA_60_70pct=as.numeric(quantile(Variance_mRNA,prob=seq(0,1,length=11))[4])
Top60_70Variance_Correlation_pvalue_mRNA_Protein=NA

for(i in 1:dim(mRNAdat_All)[1]) {
	if(Variance_mRNA[i]<Variance_mRNA_50_60pct && Variance_mRNA[i]>=Variance_mRNA_60_70pct){
		Top60_70Variance_Correlation_pvalue_mRNA_Protein[i]=cor.test(as.numeric(Protdat_All[i,]),as.numeric(mRNAdat_All[i,]),method="spearman")$p.value[1]
	}
}

Variance_mRNA_70_80pct=as.numeric(quantile(Variance_mRNA,prob=seq(0,1,length=11))[3])
Top70_80Variance_Correlation_pvalue_mRNA_Protein=NA

for(i in 1:dim(mRNAdat_All)[1]) {
	if(Variance_mRNA[i]<Variance_mRNA_60_70pct && Variance_mRNA[i]>=Variance_mRNA_70_80pct){
		Top70_80Variance_Correlation_pvalue_mRNA_Protein[i]=cor.test(as.numeric(Protdat_All[i,]),as.numeric(mRNAdat_All[i,]),method="spearman")$p.value[1]
	}
}

Variance_mRNA_80_90pct=as.numeric(quantile(Variance_mRNA,prob=seq(0,1,length=11))[2])
Top80_90Variance_Correlation_pvalue_mRNA_Protein=NA

for(i in 1:dim(mRNAdat_All)[1]) {
	if(Variance_mRNA[i]<Variance_mRNA_70_80pct && Variance_mRNA[i]>=Variance_mRNA_80_90pct){
		Top80_90Variance_Correlation_pvalue_mRNA_Protein[i]=cor.test(as.numeric(Protdat_All[i,]),as.numeric(mRNAdat_All[i,]),method="spearman")$p.value[1]
	}
}

Variance_mRNA_90_100pct=as.numeric(quantile(Variance_mRNA,prob=seq(0,1,length=11))[1])
Top90_100Variance_Correlation_pvalue_mRNA_Protein=NA

for(i in 1:dim(mRNAdat_All)[1]) {
	if(Variance_mRNA[i]<Variance_mRNA_80_90pct && Variance_mRNA[i]>=Variance_mRNA_90_100pct){
		Top90_100Variance_Correlation_pvalue_mRNA_Protein[i]=cor.test(as.numeric(Protdat_All[i,]),as.numeric(mRNAdat_All[i,]),method="spearman")$p.value[1]
	}
}

# This last is all p-values for every correlation, regardless of bin, just for reference, not shown in the graph
All=NA
All_pval=NA
for(i in 1:dim(mRNAdat_All)[1]) {
		All[i]=cor(as.numeric(Protdat_All[i,]), as.numeric(mRNAdat_All[i,]), method=c("spearman"), use="complete.obs")
		All_pval[i]=cor.test(as.numeric(Protdat_All[i,]),as.numeric(mRNAdat_All[i,]),method="spearman")$p.value[1]
}

All_pval_ADJUSTED=p.adjust(All_pval,method="BH")
sum(All_pval_ADJUSTED<0.05,na.rm=T)/sum(!is.na(All_pval_ADJUSTED)) ## 32.8% of all mRNA�protein correlations have adj.p < 0.05

y10=sum(p.adjust(Top10Variance_Correlation_pvalue_mRNA_Protein,method="BH")<0.05,na.rm=T)/sum(!is.na(Top10Variance_Correlation_pvalue_mRNA_Protein))
y9=sum(p.adjust(Top10_20Variance_Correlation_pvalue_mRNA_Protein,method="BH")<0.05,na.rm=T)/sum(!is.na(Top10_20Variance_Correlation_pvalue_mRNA_Protein))
y8=sum(p.adjust(Top20_30Variance_Correlation_pvalue_mRNA_Protein,method="BH")<0.05,na.rm=T)/sum(!is.na(Top20_30Variance_Correlation_pvalue_mRNA_Protein))
y7=sum(p.adjust(Top30_40Variance_Correlation_pvalue_mRNA_Protein,method="BH")<0.05,na.rm=T)/sum(!is.na(Top20_30Variance_Correlation_pvalue_mRNA_Protein))
y6=sum(p.adjust(Top40_50Variance_Correlation_pvalue_mRNA_Protein,method="BH")<0.05,na.rm=T)/sum(!is.na(Top20_30Variance_Correlation_pvalue_mRNA_Protein))
y5=sum(p.adjust(Top50_60Variance_Correlation_pvalue_mRNA_Protein,method="BH")<0.05,na.rm=T)/sum(!is.na(Top20_30Variance_Correlation_pvalue_mRNA_Protein))
y4=sum(p.adjust(Top60_70Variance_Correlation_pvalue_mRNA_Protein,method="BH")<0.05,na.rm=T)/sum(!is.na(Top20_30Variance_Correlation_pvalue_mRNA_Protein))
y3=sum(p.adjust(Top70_80Variance_Correlation_pvalue_mRNA_Protein,method="BH")<0.05,na.rm=T)/sum(!is.na(Top20_30Variance_Correlation_pvalue_mRNA_Protein))
y2=sum(p.adjust(Top80_90Variance_Correlation_pvalue_mRNA_Protein,method="BH")<0.05,na.rm=T)/sum(!is.na(Top20_30Variance_Correlation_pvalue_mRNA_Protein))
y1=sum(p.adjust(Top90_100Variance_Correlation_pvalue_mRNA_Protein,method="BH")<0.05,na.rm=T)/sum(!is.na(Top20_30Variance_Correlation_pvalue_mRNA_Protein))

plot(NA,NA,ylim=c(0,1),xlim=c(0,10),bty="L")
rect(0,0,1,y1,col=rgb(0,25,0,max=255))
rect(1,0,2,y2,col=rgb(0,50,0,max=255))
rect(2,0,3,y3,col=rgb(0,75,0,max=255))
rect(3,0,4,y4,col=rgb(0,100,0,max=255))
rect(4,0,5,y5,col=rgb(0,125,0,max=255))
rect(5,0,6,y6,col=rgb(0,150,0,max=255))
rect(6,0,7,y7,col=rgb(0,175,0,max=255))
rect(7,0,8,y8,col=rgb(0,200,0,max=255))
rect(8,0,9,y9,col=rgb(0,225,0,max=255))
rect(9,0,10,y10,col=rgb(0,250,0,max=255))


#$$$$$$ #$$$$$$ #$$$$$$ #$$$$$$ #$$$$$$ #$$$$$$ #$$$$$$ #$$$$$$
#$$$$$$ #$$$$$$ #$$$$$$ #$$$$$$ #$$$$$$ #$$$$$$ #$$$$$$ #$$$$$$
# SUPPLEMENTAL PANEL 2E
# Need to load Random_Individuals and All from before again

Abundance_mRNA=NA
for(i in 1:dim(mRNAdat_All)[1]) {
	Abundance_mRNA[i]=mean(as.numeric(mRNAdat_All[i,]), na.rm=T)
}

Abundance_mRNA_Top10pct=as.numeric(quantile(Abundance_mRNA,prob=seq(0,1,length=11))[10])
Abundance_mRNA_Bottom10pct=as.numeric(quantile(Abundance_mRNA,prob=seq(0,1,length=11))[2])

HighAbund_Correlation_mRNA_Protein=NA
HighAbund_Correlation_mRNA_Protein_pval=NA
LowAbund_Correlation_mRNA_Protein=NA
LowAbund_Correlation_mRNA_Protein_pval=NA

for(i in 1:dim(mRNAdat_All)[1]) {
	if(Abundance_mRNA[i]>=Abundance_mRNA_Top10pct){
		HighAbund_Correlation_mRNA_Protein[i]=cor(as.numeric(Protdat_All[i,]), as.numeric(mRNAdat_All[i,]), method=c("spearman"), use="complete.obs")
		HighAbund_Correlation_mRNA_Protein_pval[i]=cor.test(as.numeric(Protdat_All[i,]),as.numeric(mRNAdat_All[i,]),method="spearman")$p.value[1]
	}
}

for(i in 1:dim(mRNAdat_All)[1]) {
	if(Abundance_mRNA[i]<=Abundance_mRNA_Bottom10pct){
		LowAbund_Correlation_mRNA_Protein[i]=cor(as.numeric(Protdat_All[i,]), as.numeric(mRNAdat_All[i,]), method=c("spearman"), use="complete.obs")
		LowAbund_Correlation_mRNA_Protein_pval[i]=cor.test(as.numeric(Protdat_All[i,]),as.numeric(mRNAdat_All[i,]),method="spearman")$p.value[1]		
	}
}

least_abundant=sum(p.adjust(LowAbund_Correlation_mRNA_Protein_pval,method="BH")<0.05,na.rm=T)/sum(!is.na(LowAbund_Correlation_mRNA_Protein_pval))
most_abundant=sum(p.adjust(HighAbund_Correlation_mRNA_Protein_pval,method="BH")<0.05,na.rm=T)/sum(!is.na(HighAbund_Correlation_mRNA_Protein_pval))


plot(density(Random_Individuals, na.rm=T), xlim=c(-1,1), ylim=c(0, 6), col="black")
plotCI(density(All), col="lightblue", add=T, type="l")
plotCI(density(HighAbund_Correlation_mRNA_Protein, na.rm=T), col="red", add=T, type="l")
plotCI(density(LowAbund_Correlation_mRNA_Protein, na.rm=T), col="green", add=T, type="l")
#plotCI(density(LowVar_Correlation_mRNA_Protein, na.rm=T), col="yellow", add=T, type="l") # Variance is a stronger predictor than abundance, slightly
#plotCI(density(HighVar_Correlation_mRNA_Protein, na.rm=T), col="skyblue", add=T, type="l")

signif(t.test(HighAbund_Correlation_mRNA_Protein,LowAbund_Correlation_mRNA_Protein, paired=FALSE)$p.value,digits=2) # 


#LowVar_Correlation_mRNA_Protein
#$$$$$$ #$$$$$$ #$$$$$$ #$$$$$$ #$$$$$$ #$$$$$$ #$$$$$$ #$$$$$$
#$$$$$$ #$$$$$$ #$$$$$$ #$$$$$$ #$$$$$$ #$$$$$$ #$$$$$$ #$$$$$$
# SUPPLEMENTAL PANEL 2F
x=Variance_mRNA # Loaded from  panel 2F code
y=Abundance_mRNA # Loaded from panel S2E code

xy=cbind(x,y)
x=x[complete.cases(xy)]
y=y[complete.cases(xy)]
x=rank(x, na.last="keep")
y=rank(y, na.last="keep")

plotCI(x,y,cex=0.2, pch=19, col="black", bty="L")
abline(lm(y~x))
rhoc=cor(x,y, method=c("spearman"), use="complete.obs")
pr = cor.test(x,y,method="spearman")$p.value[1]			# This outputs the p-value of the SPEARMAN correlation
rprint=substitute(paste(italic("rho"), " = ", rhoc), list(rhoc=signif(rhoc[1], digits=2)))	# Makes the r-value with italicized r
pletter=substitute(paste("   ", italic("p"), " = ", pr), list(pr=signif(pr[1], digits=2)))	
legend("topleft", bg='white', bty="n", cex=1, text.col="red", legend=c(as.expression(rprint), as.expression(pletter)))


#$$$$$$ #$$$$$$ #$$$$$$ #$$$$$$ #$$$$$$ #$$$$$$ #$$$$$$ #$$$$$$
#$$$$$$ #$$$$$$ #$$$$$$ #$$$$$$ #$$$$$$ #$$$$$$ #$$$$$$ #$$$$$$
# SUPPLEMENTAL PANEL 2G

setwd("~/Dropbox/Aging\ BXD\ Study/1\ _\ Manuscript\ -\ Evan\ Multi-Omic/")
CORUM_List=read.table("b_Figure2_CORUM_Details.csv", row.names=1, header=T, sep=",")

Complex_Size_1=1 # Set up the bins (somewhat arbitrarily)
Complex_Size_2=2
Complex_Size_3=3
Complex_Size_4=4
Complex_Size_5=5
Complex_Size_6=6
Complex_Size_10=10
Complex_Size_20=20

## Now for each bin, we create the list of genes that fit that size, then perform the correlation analysis for only those genes
Complex_Size_1_GeneList=NA
for(i in 1:dim(CORUM_List)[1]) {
	if(CORUM_List$ComplexSize[i]>=Complex_Size_1 && CORUM_List$ComplexSize[i]<Complex_Size_2){
		Complex_Size_1_GeneList[i]=CORUM_List$mRNA_ID[i]
	}
}
Complex_Size_1_GeneList=Complex_Size_1_GeneList[!is.na(Complex_Size_1_GeneList)]

Complex_Size_1_pValue=NA
for(i in 1:dim(mRNAdat_All)[1]) {
	if(!is.na(match(rownames(mRNAdat_All)[i],Complex_Size_1_GeneList))) { 
		Complex_Size_1_pValue[i]=cor.test(as.numeric(Protdat_All[i,]),as.numeric(mRNAdat_All[i,]),method="spearman")$p.value[1]
	}
}

## Complex size 2 (dimers only)
Complex_Size_2_GeneList=NA
for(i in 1:dim(CORUM_List)[1]) {
	if(CORUM_List$ComplexSize[i]>=Complex_Size_2 && CORUM_List$ComplexSize[i]<Complex_Size_3){
		Complex_Size_2_GeneList[i]=CORUM_List$mRNA_ID[i]
	}
}
Complex_Size_2_GeneList=Complex_Size_2_GeneList[!is.na(Complex_Size_2_GeneList)]

Complex_Size_2_pValue=NA
for(i in 1:dim(mRNAdat_All)[1]) {
	if(!is.na(match(rownames(mRNAdat_All)[i],Complex_Size_2_GeneList))) { 
		Complex_Size_2_pValue[i]=cor.test(as.numeric(Protdat_All[i,]),as.numeric(mRNAdat_All[i,]),method="spearman")$p.value[1]
	}
}

## Complex size 3
Complex_Size_3_GeneList=NA
for(i in 1:dim(CORUM_List)[1]) {
	if(CORUM_List$ComplexSize[i]>=Complex_Size_3 && CORUM_List$ComplexSize[i]<Complex_Size_4){
		Complex_Size_3_GeneList[i]=CORUM_List$mRNA_ID[i]
	}
}
Complex_Size_3_GeneList=Complex_Size_3_GeneList[!is.na(Complex_Size_3_GeneList)]

Complex_Size_3_pValue=NA
for(i in 1:dim(mRNAdat_All)[1]) {
	if(!is.na(match(rownames(mRNAdat_All)[i],Complex_Size_3_GeneList))) { 
		Complex_Size_3_pValue[i]=cor.test(as.numeric(Protdat_All[i,]),as.numeric(mRNAdat_All[i,]),method="spearman")$p.value[1]
	}
}

## Complex size 4
Complex_Size_4_GeneList=NA
for(i in 1:dim(CORUM_List)[1]) {
	if(CORUM_List$ComplexSize[i]>=Complex_Size_4 && CORUM_List$ComplexSize[i]<Complex_Size_5){
		Complex_Size_4_GeneList[i]=CORUM_List$mRNA_ID[i]
	}
}
Complex_Size_4_GeneList=Complex_Size_4_GeneList[!is.na(Complex_Size_4_GeneList)]

Complex_Size_4_pValue=NA
for(i in 1:dim(mRNAdat_All)[1]) {
	if(!is.na(match(rownames(mRNAdat_All)[i],Complex_Size_4_GeneList))) { 
		Complex_Size_4_pValue[i]=cor.test(as.numeric(Protdat_All[i,]),as.numeric(mRNAdat_All[i,]),method="spearman")$p.value[1]
	}
}

## Complex size 5
Complex_Size_5_GeneList=NA
for(i in 1:dim(CORUM_List)[1]) {
	if(CORUM_List$ComplexSize[i]>=Complex_Size_5 && CORUM_List$ComplexSize[i]<Complex_Size_6){
		Complex_Size_5_GeneList[i]=CORUM_List$mRNA_ID[i]
	}
}
Complex_Size_5_GeneList=Complex_Size_5_GeneList[!is.na(Complex_Size_5_GeneList)]

Complex_Size_5_pValue=NA
for(i in 1:dim(mRNAdat_All)[1]) {
	if(!is.na(match(rownames(mRNAdat_All)[i],Complex_Size_5_GeneList))) { 
		Complex_Size_5_pValue[i]=cor.test(as.numeric(Protdat_All[i,]),as.numeric(mRNAdat_All[i,]),method="spearman")$p.value[1]
	}
}

## Complex size 6 to 9
Complex_Size_6_GeneList=NA
for(i in 1:dim(CORUM_List)[1]) {
	if(CORUM_List$ComplexSize[i]>=Complex_Size_6 && CORUM_List$ComplexSize[i]<Complex_Size_10){
		Complex_Size_6_GeneList[i]=CORUM_List$mRNA_ID[i]
	}
}
Complex_Size_6_GeneList=Complex_Size_6_GeneList[!is.na(Complex_Size_6_GeneList)]

Complex_Size_6_pValue=NA
for(i in 1:dim(mRNAdat_All)[1]) {
	if(!is.na(match(rownames(mRNAdat_All)[i],Complex_Size_6_GeneList))) { 
		Complex_Size_6_pValue[i]=cor.test(as.numeric(Protdat_All[i,]),as.numeric(mRNAdat_All[i,]),method="spearman")$p.value[1]
	}
}

## Complex size 10 to 19
Complex_Size_10_GeneList=NA
for(i in 1:dim(CORUM_List)[1]) {
	if(CORUM_List$ComplexSize[i]>=Complex_Size_10 && CORUM_List$ComplexSize[i]<Complex_Size_20){
		Complex_Size_10_GeneList[i]=CORUM_List$mRNA_ID[i]
	}
}
Complex_Size_10_GeneList=Complex_Size_10_GeneList[!is.na(Complex_Size_10_GeneList)]

Complex_Size_10_pValue=NA
for(i in 1:dim(mRNAdat_All)[1]) {
	if(!is.na(match(rownames(mRNAdat_All)[i],Complex_Size_10_GeneList))) { 
		Complex_Size_10_pValue[i]=cor.test(as.numeric(Protdat_All[i,]),as.numeric(mRNAdat_All[i,]),method="spearman")$p.value[1]
	}
}

## Complex size � 20
Complex_Size_20_GeneList=NA
for(i in 1:dim(CORUM_List)[1]) {
	if(CORUM_List$ComplexSize[i]>=Complex_Size_20){
		Complex_Size_20_GeneList[i]=CORUM_List$mRNA_ID[i]
	}
}
Complex_Size_20_GeneList=Complex_Size_20_GeneList[!is.na(Complex_Size_20_GeneList)]

Complex_Size_20_pValue=NA
for(i in 1:dim(mRNAdat_All)[1]) {
	if(!is.na(match(rownames(mRNAdat_All)[i],Complex_Size_20_GeneList))) { 
		Complex_Size_20_pValue[i]=cor.test(as.numeric(Protdat_All[i,]),as.numeric(mRNAdat_All[i,]),method="spearman")$p.value[1]
	}
}

y8=sum(p.adjust(Complex_Size_20_pValue,method="BH")<0.05,na.rm=T)/sum(!is.na(Complex_Size_20_pValue))
y7=sum(p.adjust(Complex_Size_10_pValue,method="BH")<0.05,na.rm=T)/sum(!is.na(Complex_Size_10_pValue))
y6=sum(p.adjust(Complex_Size_6_pValue,method="BH")<0.05,na.rm=T)/sum(!is.na(Complex_Size_6_pValue))
y5=sum(p.adjust(Complex_Size_5_pValue,method="BH")<0.05,na.rm=T)/sum(!is.na(Complex_Size_5_pValue))
y4=sum(p.adjust(Complex_Size_4_pValue,method="BH")<0.05,na.rm=T)/sum(!is.na(Complex_Size_4_pValue))
y3=sum(p.adjust(Complex_Size_3_pValue,method="BH")<0.05,na.rm=T)/sum(!is.na(Complex_Size_3_pValue))
y2=sum(p.adjust(Complex_Size_2_pValue,method="BH")<0.05,na.rm=T)/sum(!is.na(Complex_Size_2_pValue))
y1=sum(p.adjust(Complex_Size_1_pValue,method="BH")<0.05,na.rm=T)/sum(!is.na(Complex_Size_1_pValue))

plot(NA,NA,ylim=c(0,1),xlim=c(0,11),bty="L")
rect(0,0,1,y1,col=rgb(0,25,0,max=255)) ## Complex size 1
rect(1,0,2,y2,col=rgb(0,50,0,max=255)) ## Complex size 2
rect(2,0,3,y3,col=rgb(0,75,0,max=255)) ## Complex size 3
rect(3,0,4,y4,col=rgb(0,100,0,max=255)) ## Complex size 4
rect(4,0,5,y5,col=rgb(0,125,0,max=255)) ## Complex size 5
rect(5,0,7,y6,col=rgb(0,150,0,max=255)) ## Complex size 6-9
rect(7,0,9,y7,col=rgb(0,175,0,max=255)) ## Complex size 10-19
rect(9,0,11,y8,col=rgb(0,200,0,max=255)) ## Complex size >= 20


#$$$$$$ #$$$$$$ #$$$$$$ #$$$$$$ #$$$$$$ #$$$$$$ #$$$$$$ #$$$$$$
#$$$$$$ #$$$$$$ #$$$$$$ #$$$$$$ #$$$$$$ #$$$$$$ #$$$$$$ #$$$$$$
# SUPPLEMENTAL PANEL 2H

# Drawn by hand to scale, using fixed points on LOD = 1, 2, 3, 4
# "True positives" used the actual data over a ratio of randomized input data for the QTL calculations
# Then at each LOD score cutoff, we took the # of true positive at ��[that LOD] divided by the # of cis-QTLs calculated at ��[that LOD] in the randomized data
# Randomized data meaning that the sample labels (mice) were shuffled with "sample", but still using the real measurement data.


#$$$$$$ #$$$$$$ #$$$$$$ #$$$$$$ #$$$$$$ #$$$$$$ #$$$$$$ #$$$$$$
#$$$$$$ #$$$$$$ #$$$$$$ #$$$$$$ #$$$$$$ #$$$$$$ #$$$$$$ #$$$$$$
# SUPPLEMENTAL PANEL 2I

setwd("~/Dropbox/Aging\ BXD\ Study/1\ _\ Manuscript\ -\ Evan\ Multi-Omic/")

QTLData=read.table("b_Figure2_cisQTLs_All.csv", row.names=1, header=T, sep=",")

CD_Prot_cisQTL=NA
HF_Prot_cisQTL=NA
Either_Prot_cisQTL=NA
	
# The file is sorted to only take the 3511 genes with Prot data and protein data that output QTLs
for(i in 1:3511) {
	if(!is.na(as.numeric(as.character(QTLData$Prot_CD_LOD[i])))) {
		if(as.numeric(as.character(QTLData$Prot_CD_LOD[i])) >= 4.0) {
			CD_Prot_cisQTL[i]=as.numeric(as.character(QTLData$Prot_CD_LOD[i]))
		}
		else if (as.numeric(as.character(QTLData$Prot_CD_LOD[i])) <= 4.0) {
			CD_Prot_cisQTL[i]=NA
		}	
	}
}
		
for(i in 1:3511) {
	if(!is.na(as.numeric(as.character(QTLData$Prot_HF_LOD[i])))) {
		if(as.numeric(as.character(QTLData$Prot_HF_LOD[i])) >= 4.0) {
			HF_Prot_cisQTL[i]=as.numeric(as.character(QTLData$Prot_HF_LOD[i]))
		}
		else if (as.numeric(as.character(QTLData$Prot_HF_LOD[i])) <= 4.0) {
			HF_Prot_cisQTL[i]=NA
		}	
	}
}

for(i in 1:3511) {
	if(!is.na(as.numeric(as.character(QTLData$Prot_CD_LOD[i]))) && !is.na(as.numeric(as.character(QTLData$Prot_HF_LOD[i])))) {
		if(as.numeric(as.character(QTLData$Prot_CD_LOD[i])) >= 4.0 || as.numeric(as.character(QTLData$Prot_HF_LOD[i])) >= 4.0) {	
			Either_Prot_cisQTL[i]=as.numeric(as.character(QTLData$Prot_CD_LOD[i])) # Whichever, it doesn't matter, this is just a placeholder
		}
	}
}

CD_Prot_cisQTLcount=sum((!is.na(CD_Prot_cisQTL)))
HFD_Prot_cisQTLcount=sum((!is.na(HF_Prot_cisQTL)))
Either_Prot_cisQTLcount=sum((!is.na(Either_Prot_cisQTL)))

# Now count how many QTLs are in each part of the Venn diagram
CD_Prot_cisQTLcount_unique=Either_Prot_cisQTLcount-HFD_Prot_cisQTLcount
HF_Prot_cisQTLcount_unique=Either_Prot_cisQTLcount-CD_Prot_cisQTLcount
Prot_cisQTL_Overlap=Either_Prot_cisQTLcount-CD_Prot_cisQTLcount_unique-HF_Prot_cisQTLcount_unique

plot(venneuler(c(CD=CD_Prot_cisQTLcount_unique, HFD=HF_Prot_cisQTLcount_unique, "CD&HFD"=Prot_cisQTL_Overlap)))

## Middle Slopegraph
KeepcispQTLs=which(!is.na(Either_Prot_cisQTL))

greycount=0;darkredcount=0;greencount=0;pinkcount=0;purplecount=0 # initialize the counters
y1=as.numeric(as.character(QTLData$Prot_CD_LOD[KeepcispQTLs]))
y2=as.numeric(as.character(QTLData$Prot_HF_LOD[KeepcispQTLs]))
x1=rep(1, length=length(y1)) # make this a sequence the length of y1 and y2
x2=rep(2, length=length(y2)) # make this a sequence the length of y1 and y2

plotCI(NA,NA, xlim=c(.9,2.1), ylim=c(0,11), bty="L") ## This initializes the plot
for (i in 1:length(x1)) {
### First plot the lines (you can plot after, but it causes a small visual bug when lines are plotted on top of points)
# If loops are used to set the criteria for line coloration, easy to modify however you see fit
if(!is.na(y1[i]) & !is.na(y2[i])) {
	if(y1[i] >= 4.0 && y2[i] >= 4.0) {
 	 lines(x=c(x1[i],x2[i]), y=c(y1[i],y2[i]), lty="solid", col="grey") # No major change in LOD score
 	 greycount=greycount+1
 	 }
	else if(y1[i] >= 4.0 && y2[i] >= 2.0) {
 	 lines(x=c(x1[i],x2[i]), y=c(y1[i],y2[i]), lty="solid", col="darkred")
 	 darkredcount=darkredcount+1
 	 }
	else if(y2[i] >= 4.0 && y1[i] >= 2.0) {
 	 lines(x=c(x1[i],x2[i]), y=c(y1[i],y2[i]), lty="solid", col="green")
 	 greencount=greencount+1
 	 } 	 
	else if(y1[i] >= 4.0 && y2[i] < 2.0) {
 	 lines(x=c(x1[i],x2[i]), y=c(y1[i],y2[i]), lty="solid", col="pink")
 	 pinkcount=pinkcount+1
 	 } 	  	 
	else if(y2[i] >= 4.0 && y1[i] < 2.0) {
 	 lines(x=c(x1[i],x2[i]), y=c(y1[i],y2[i]), lty="solid", col="purple")
 	 purplecount=purplecount+1
 	 } 	  	 
}
 	
# now plot the actual points
  plotCI(x1[i],y1[i],lwd=2,pch=21,pt.bg="grey",col="black", add=T)
  plotCI(x2[i],y2[i],lwd=2,pch=21,pt.bg="grey",col="black", add=T)
}

greycount;darkredcount;greencount;pinkcount;purplecount

# Right Venn

plot(venneuler(c(CD=pinkcount, HFD=purplecount, "CD&HFD"=(greycount+darkredcount+greencount))))
# Note that this plot flips and puts the largest Venn on the left; so the figure is flipped in illustrator since HFD is larger, for visual consistency


#$$$$$$ #$$$$$$ #$$$$$$ #$$$$$$ #$$$$$$ #$$$$$$ #$$$$$$ #$$$$$$
#$$$$$$ #$$$$$$ #$$$$$$ #$$$$$$ #$$$$$$ #$$$$$$ #$$$$$$ #$$$$$$
# SUPPLEMENTAL PANEL 2J
mRNAdat_young=Top3772Data_young[1:3772,]
Protdat_young=Top3772Data_young[3773:dim(Top3772Data_young)[1],]

# LEFT PANEL:
# RIGHT PANEL:
setwd("~/Dropbox/Aging\ BXD\ Study/1\ _\ Manuscript\ -\ Evan\ Multi-Omic/")

QTLData=read.table("b_Figure2_cisQTLs_All.csv", row.names=1, header=T, sep=",")

Young_Prot_cisQTL=NA
Young_mRNA_cisQTL=NA
Either_Young_cisQTL=NA
	
# The file is sorted to only take the 3511 genes with Prot data and protein data that output QTLs
for(i in 1:3511) {
	if(!is.na(as.numeric(as.character(QTLData$Prot_Young_LOD[i])))) {
		if(as.numeric(as.character(QTLData$Prot_Young_LOD[i])) >= 4.0) {
			Young_Prot_cisQTL[i]=as.numeric(as.character(QTLData$Prot_Young_LOD[i]))
		}
		else if (as.numeric(as.character(QTLData$Prot_Young_LOD[i])) <= 4.0) {
			Young_Prot_cisQTL[i]=NA
		}	
	}
}
		
for(i in 1:3511) {
	if(!is.na(as.numeric(as.character(QTLData$mRNA_Young_LOD[i])))) {
		if(as.numeric(as.character(QTLData$mRNA_Young_LOD[i])) >= 4.0) {
			Young_mRNA_cisQTL[i]=as.numeric(as.character(QTLData$mRNA_Young_LOD[i]))
		}
		else if (as.numeric(as.character(QTLData$mRNA_Young_LOD[i])) <= 4.0) {
			Young_mRNA_cisQTL[i]=NA
		}	
	}
}

for(i in 1:3511) {
	if(!is.na(as.numeric(as.character(QTLData$Prot_Young_LOD[i]))) && !is.na(as.numeric(as.character(QTLData$mRNA_Young_LOD[i])))) {
		if(as.numeric(as.character(QTLData$Prot_Young_LOD[i])) >= 4.0 || as.numeric(as.character(QTLData$mRNA_Young_LOD[i])) >= 4.0) {	
			Either_Young_cisQTL[i]=as.numeric(as.character(QTLData$Prot_Young_LOD[i])) # Whichever, it doesn't matter, this is just a placeholder
		}
	}
}

Young_Prot_cisQTLcount=sum((!is.na(Young_Prot_cisQTL)))
Young_mRNA_cisQTLcount=sum((!is.na(Young_mRNA_cisQTL)))
Either_Young_cisQTLcount=sum((!is.na(Either_Young_cisQTL)))

# Now count how many QTLs are in each part of the Venn diagram
Young_Prot_cisQTLcount_unique=Either_Young_cisQTLcount-Young_mRNA_cisQTLcount
Young_mRNA_cisQTLcount_unique=Either_Young_cisQTLcount-Young_Prot_cisQTLcount
Prot_cisQTL_Overlap=Either_Young_cisQTLcount-Young_Prot_cisQTLcount_unique-Young_mRNA_cisQTLcount_unique

## Now the Slopegraph
KeepcispQTLs=which(!is.na(Either_Young_cisQTL))

greycount=0;darkbluecount=0;yellowcount=0;skybluecount=0;redcount=0 # initialize the counters
y1=as.numeric(as.character(QTLData$Prot_Young_LOD[KeepcispQTLs]))
y2=as.numeric(as.character(QTLData$mRNA_Young_LOD[KeepcispQTLs]))
x1=rep(1, length=length(y1)) # make this a sequence the length of y1 and y2
x2=rep(2, length=length(y2)) # make this a sequence the length of y1 and y2

plotCI(NA,NA, xlim=c(.9,2.1), ylim=c(0,11), bty="L") ## This initializes the plot
for (i in 1:length(x1)) {
### First plot the lines (you can plot after, but it causes a small visual bug when lines are plotted on top of points)
# If loops are used to set the criteria for line coloration, easy to modify however you see fit
if(!is.na(y1[i]) & !is.na(y2[i])) {
	if(y1[i] >= 4.0 && y2[i] >= 4.0) {
 	 lines(x=c(x1[i],x2[i]), y=c(y1[i],y2[i]), lty="solid", col="grey") # No major change in LOD score
 	 greycount=greycount+1
 	 }
	else if(y1[i] >= 4.0 && y2[i] >= 2.0) {
 	 lines(x=c(x1[i],x2[i]), y=c(y1[i],y2[i]), lty="solid", col="darkblue")
 	 darkbluecount=darkbluecount+1
 	 }
	else if(y2[i] >= 4.0 && y1[i] >= 2.0) {
 	 lines(x=c(x1[i],x2[i]), y=c(y1[i],y2[i]), lty="solid", col="yellow")
 	 yellowcount=yellowcount+1
 	 } 	 
	else if(y1[i] >= 4.0 && y2[i] < 2.0) {
 	 lines(x=c(x1[i],x2[i]), y=c(y1[i],y2[i]), lty="solid", col="skyblue")
 	 skybluecount=skybluecount+1
 	 } 	  	 
	else if(y2[i] >= 4.0 && y1[i] < 2.0) {
 	 lines(x=c(x1[i],x2[i]), y=c(y1[i],y2[i]), lty="solid", col="red")
 	 redcount=redcount+1
 	 } 	  	 
}
 	
# now plot the actual points
  plotCI(x1[i],y1[i],lwd=2,pch=21,pt.bg="grey",col="black", add=T)
  plotCI(x2[i],y2[i],lwd=2,pch=21,pt.bg="grey",col="black", add=T)
}

greycount;darkbluecount;yellowcount;skybluecount;redcount


# RIGHT PANEL:
setwd("~/Dropbox/Aging\ BXD\ Study/1\ _\ Manuscript\ -\ Evan\ Multi-Omic/")

QTLData=read.table("b_Figure2_cisQTLs_All.csv", row.names=1, header=T, sep=",")

Young_Prot_cisQTL=NA
Old_Prot_cisQTL=NA
Either_Prot_cisQTL=NA
	
# The file is sorted to only take the 3511 genes with Prot data and protein data that output QTLs
for(i in 1:3511) {
	if(!is.na(as.numeric(as.character(QTLData$Prot_Young_LOD[i])))) {
		if(as.numeric(as.character(QTLData$Prot_Young_LOD[i])) >= 4.0) {
			Young_Prot_cisQTL[i]=as.numeric(as.character(QTLData$Prot_Young_LOD[i]))
		}
		else if (as.numeric(as.character(QTLData$Prot_Young_LOD[i])) <= 4.0) {
			Young_Prot_cisQTL[i]=NA
		}	
	}
}
		
for(i in 1:3511) {
	if(!is.na(as.numeric(as.character(QTLData$Prot_Old_LOD[i])))) {
		if(as.numeric(as.character(QTLData$Prot_Old_LOD[i])) >= 4.0) {
			Old_Prot_cisQTL[i]=as.numeric(as.character(QTLData$Prot_Old_LOD[i]))
		}
		else if (as.numeric(as.character(QTLData$Prot_Old_LOD[i])) <= 4.0) {
			Old_Prot_cisQTL[i]=NA
		}	
	}
}

for(i in 1:3511) {
	if(!is.na(as.numeric(as.character(QTLData$Prot_Young_LOD[i]))) && !is.na(as.numeric(as.character(QTLData$Prot_Old_LOD[i])))) {
		if(as.numeric(as.character(QTLData$Prot_Young_LOD[i])) >= 4.0 || as.numeric(as.character(QTLData$Prot_Old_LOD[i])) >= 4.0) {	
			Either_Prot_cisQTL[i]=as.numeric(as.character(QTLData$Prot_Young_LOD[i])) # Whichever, it doesn't matter, this is just a placeholder
		}
	}
}

Young_Prot_cisQTLcount=sum((!is.na(Young_Prot_cisQTL)))
HFD_Prot_cisQTLcount=sum((!is.na(Old_Prot_cisQTL)))
Either_Prot_cisQTLcount=sum((!is.na(Either_Prot_cisQTL)))

# Now count how many QTLs are in each part of the Venn diagram
Young_Prot_cisQTLcount_unique=Either_Prot_cisQTLcount-HFD_Prot_cisQTLcount
Old_Prot_cisQTLcount_unique=Either_Prot_cisQTLcount-Young_Prot_cisQTLcount
Prot_cisQTL_Overlap=Either_Prot_cisQTLcount-Young_Prot_cisQTLcount_unique-Old_Prot_cisQTLcount_unique

## Now the Slopegraph
KeepcispQTLs=which(!is.na(Either_Prot_cisQTL))

greycount=0;darkbluecount=0;yellowcount=0;pinkcount=0;purplecount=0 # initialize the counters
y1=as.numeric(as.character(QTLData$Prot_Young_LOD[KeepcispQTLs]))
y2=as.numeric(as.character(QTLData$Prot_Old_LOD[KeepcispQTLs]))
x1=rep(1, length=length(y1)) # make this a sequence the length of y1 and y2
x2=rep(2, length=length(y2)) # make this a sequence the length of y1 and y2

plotCI(NA,NA, xlim=c(.9,2.1), ylim=c(0,11), bty="L") ## This initializes the plot
for (i in 1:length(x1)) {
### First plot the lines (you can plot after, but it causes a small visual bug when lines are plotted on top of points)
# If loops are used to set the criteria for line coloration, easy to modify however you see fit
if(!is.na(y1[i]) & !is.na(y2[i])) {
	if(y1[i] >= 4.0 && y2[i] >= 4.0) {
 	 lines(x=c(x1[i],x2[i]), y=c(y1[i],y2[i]), lty="solid", col="grey") # No major change in LOD score
 	 greycount=greycount+1
 	 }
	else if(y1[i] >= 4.0 && y2[i] >= 2.0) {
 	 lines(x=c(x1[i],x2[i]), y=c(y1[i],y2[i]), lty="solid", col="darkblue")
 	 darkbluecount=darkbluecount+1
 	 }
	else if(y2[i] >= 4.0 && y1[i] >= 2.0) {
 	 lines(x=c(x1[i],x2[i]), y=c(y1[i],y2[i]), lty="solid", col="yellow")
 	 yellowcount=yellowcount+1
 	 } 	 
	else if(y1[i] >= 4.0 && y2[i] < 2.0) {
 	 lines(x=c(x1[i],x2[i]), y=c(y1[i],y2[i]), lty="solid", col="pink")
 	 pinkcount=pinkcount+1
 	 } 	  	 
	else if(y2[i] >= 4.0 && y1[i] < 2.0) {
 	 lines(x=c(x1[i],x2[i]), y=c(y1[i],y2[i]), lty="solid", col="purple")
 	 purplecount=purplecount+1
 	 } 	  	 
}
 	
# now plot the actual points
  plotCI(x1[i],y1[i],lwd=2,pch=21,pt.bg="grey",col="black", add=T)
  plotCI(x2[i],y2[i],lwd=2,pch=21,pt.bg="grey",col="black", add=T)
}

greycount;darkbluecount;yellowcount;pinkcount;purplecount
```
