## Supplementary material for "Multi-Omic Profiling of the Liver Across Diets and Age in a Diverse Mouse Population": All code used to generate figures: b_Figure3_Plotting_Code_vF.r

```
################################################################################  
# First priority is loading the multi-omics dataset, which will be used throughout
## This is the INDIVIDUAL data
################################################################################# 

setwd("~/Dropbox/Aging\ BXD\ Study/1\ _\ Manuscript\ -\ Evan\ Multi-Omic/")

AllData_header=read.table("aTableS2_ALL_DATA.csv", nrows=8, row.names=1, header=T, sep=",")
AllData_header=AllData_header[,3:ncol(AllData_header)]
AllData=read.table("aTableS2_ALL_DATA.csv", row.names=1, header=T,  na.strings="", sep=",", skip=8, stringsAsFactors=FALSE)
AllData=AllData[1:nrow(AllData),3:ncol(AllData)]

AllData_CD=matrix(NA, nrow=nrow(AllData), ncol=length(AllData))
for(i in 1:length(AllData)) {
	if(as.character(AllData_header["Diet",i])=="CD") {   AllData_CD[,i]=as.numeric(as.character(AllData[,i]))	}
} 
rownames(AllData_CD)=rownames(AllData)

AllData_HF=matrix(NA, nrow=nrow(AllData), ncol=length(AllData))
for(i in 1:length(AllData)) {
	if(as.character(AllData_header["Diet",i])=="HF") {   AllData_HF[,i]=as.numeric(as.character(AllData[,i]))	}
}
rownames(AllData_HF)=rownames(AllData)

AllData_young=matrix(NA, nrow=nrow(AllData), ncol=length(AllData))
for(i in 1:length(AllData)) {
	if(as.character(AllData_header["Age",i])<=420) {	AllData_young[,i]=as.numeric(as.character(AllData[,i]))	}
}
rownames(AllData_young)=rownames(AllData)

AllData_old=matrix(NA, nrow=nrow(AllData), ncol=length(AllData))
for(i in 1:length(AllData)) {
	if(as.character(AllData_header["Age",i])>=420) {	AllData_old[,i]=as.numeric(as.character(AllData[,i]))	}
}
rownames(AllData_old)=rownames(AllData)

################################################################################  
# FIGURE THREE
################################################################################# 

#$$$$$$ #$$$$$$ #$$$$$$ #$$$$$$ #$$$$$$ #$$$$$$ #$$$$$$ #$$$$$$
#$$$$$$ #$$$$$$ #$$$$$$ #$$$$$$ #$$$$$$ #$$$$$$ #$$$$$$ #$$$$$$
# PANEL 3A

# This is the output from DAVID

#$$$$$$ #$$$$$$ #$$$$$$ #$$$$$$ #$$$$$$ #$$$$$$ #$$$$$$ #$$$$$$
#$$$$$$ #$$$$$$ #$$$$$$ #$$$$$$ #$$$$$$ #$$$$$$ #$$$$$$ #$$$$$$
# PANEL 3B

# This is the output from GSEA


#$$$$$$ #$$$$$$ #$$$$$$ #$$$$$$ #$$$$$$ #$$$$$$ #$$$$$$ #$$$$$$
#$$$$$$ #$$$$$$ #$$$$$$ #$$$$$$ #$$$$$$ #$$$$$$ #$$$$$$ #$$$$$$
# PANEL 3C

# Remember, need to load the data at the top.

# LEFT PART
groupA=as.numeric(AllData_young["mRNA_20690_Ctsd",])
groupB=as.numeric(AllData_old["mRNA_20690_Ctsd",])
stripchart(groupA, at=0.7, pch=21, cex=1.2, bg="grey", method="jitter", xlim=c(0.6, 1.3), las=1, ylim=c(min(c(groupA,groupB), na.rm=T), max(c(groupA,groupB), na.rm=T)), vertical=T)
stripchart(groupB, at=1.0, pch=21, cex=1.2, bg="black", method="jitter", add=T, vertical=T)
	segments(0.65, quantile(groupA, na.rm=T)[2], 0.75, quantile(groupA, na.rm=T)[2], col="red", lwd=3)	# first quantile bar for first cohort
	segments(0.6, median(groupA, na.rm=T), 0.8, median(groupA, na.rm=T), col="red", lwd=3)	# median bar for first cohort
	segments(0.65, quantile(groupA, na.rm=T)[4], 0.75, quantile(groupA, na.rm=T)[4], col="red", lwd=3)	# first quantile bar for first cohort
	segments(0.7,quantile(groupA, na.rm=T)[2],0.7,quantile(groupA, na.rm=T)[4], col="red", lwd=3)
	segments(.95, quantile(groupB, na.rm=T)[2], 1.05, quantile(groupB, na.rm=T)[2], col="red", lwd=3)	# first quantile bar for second cohort	
	segments(.9, median(groupB, na.rm=T), 1.1, median(groupB, na.rm=T), col="red", lwd=3)	# median bar for second cohort, etc
	segments(.95, quantile(groupB, na.rm=T)[4], 1.05, quantile(groupB, na.rm=T)[4], col="red", lwd=3)	# first quantile bar for second cohort
	segments(1.0,quantile(groupB, na.rm=T)[2],1.0,quantile(groupB, na.rm=T)[4], col="red", lwd=3)
# legend	
pr = signif(t.test(groupA,groupB, paired=FALSE)$p.value,digits=2)			# This outputs the p-value of the SPEARMAN correlation but does not go below e-16 and I can't figure out the fix
pletter=substitute(paste("   ", italic("p"), " = ", pr), list(pr=signif(pr[1], digits=2)))	# Makes the r-value with italicized r

for (i in 1:1) {	# Why does this not work if it's not in a for loop??
	if (mean(groupA, na.rm=T)-mean(groupB, na.rm=T) >= 0) { # This calculates positive fold change
		fc=round(2^(mean(groupA, na.rm=T)-mean(groupB, na.rm=T)),2); legendposition="topright"
	} else if (mean(groupA, na.rm=T)-mean(groupB, na.rm=T) <= 0) { # This makes it into negative fold change, but SYMMETRICAL (i.e. -0.5 is the same as +2.0 but it's hard to tell)
		fc=round(-1*(2^(mean(groupB, na.rm=T)-mean(groupA, na.rm=T))),2); legendposition="topleft"
	} else {	fc = 0; legendposition="topleft"}
}
foldchange=paste("FC = ", fc)
legend(legendposition, bg='white', bty="n", cex=1, text.col="red", legend=c(foldchange, as.expression(pletter)))

### RIGHT PART
groupA=as.numeric(AllData_young["Prot_13918_Ctsd",])
groupB=as.numeric(AllData_old["Prot_13918_Ctsd",])
stripchart(groupA, at=0.7, pch=21, cex=1.2, bg="grey", method="jitter", xlim=c(0.6, 1.3), las=1, ylim=c(min(c(groupA,groupB), na.rm=T), max(c(groupA,groupB), na.rm=T)), vertical=T)
stripchart(groupB, at=1.0, pch=21, cex=1.2, bg="black", method="jitter", add=T, vertical=T)
	segments(0.65, quantile(groupA, na.rm=T)[2], 0.75, quantile(groupA, na.rm=T)[2], col="red", lwd=3)	# first quantile bar for first cohort
	segments(0.6, median(groupA, na.rm=T), 0.8, median(groupA, na.rm=T), col="red", lwd=3)	# median bar for first cohort
	segments(0.65, quantile(groupA, na.rm=T)[4], 0.75, quantile(groupA, na.rm=T)[4], col="red", lwd=3)	# first quantile bar for first cohort
	segments(0.7,quantile(groupA, na.rm=T)[2],0.7,quantile(groupA, na.rm=T)[4], col="red", lwd=3)
	segments(.95, quantile(groupB, na.rm=T)[2], 1.05, quantile(groupB, na.rm=T)[2], col="red", lwd=3)	# first quantile bar for second cohort	
	segments(.9, median(groupB, na.rm=T), 1.1, median(groupB, na.rm=T), col="red", lwd=3)	# median bar for second cohort, etc
	segments(.95, quantile(groupB, na.rm=T)[4], 1.05, quantile(groupB, na.rm=T)[4], col="red", lwd=3)	# first quantile bar for second cohort
	segments(1.0,quantile(groupB, na.rm=T)[2],1.0,quantile(groupB, na.rm=T)[4], col="red", lwd=3)
# legend	
pr = signif(t.test(groupA,groupB, paired=FALSE)$p.value,digits=2)			# This outputs the p-value of the SPEARMAN correlation but does not go below e-16 and I can't figure out the fix
pletter=substitute(paste("   ", italic("p"), " = ", pr), list(pr=signif(pr[1], digits=2)))	# Makes the r-value with italicized r

for (i in 1:1) {	# Why does this not work if it's not in a for loop??
	if (mean(groupA, na.rm=T)-mean(groupB, na.rm=T) >= 0) { # This calculates positive fold change
		fc=round(2^(mean(groupA, na.rm=T)-mean(groupB, na.rm=T)),2); legendposition="topright"
	} else if (mean(groupA, na.rm=T)-mean(groupB, na.rm=T) <= 0) { # This makes it into negative fold change, but SYMMETRICAL (i.e. -0.5 is the same as +2.0 but it's hard to tell)
		fc=round(-1*(2^(mean(groupB, na.rm=T)-mean(groupA, na.rm=T))),2); legendposition="topleft"
	} else {	fc = 0; legendposition="topleft"}
}
foldchange=paste("FC = ", fc)
legend(legendposition, bg='white', bty="n", cex=1, text.col="red", legend=c(foldchange, as.expression(pletter)))


#$$$$$$ #$$$$$$ #$$$$$$ #$$$$$$ #$$$$$$ #$$$$$$ #$$$$$$ #$$$$$$
#$$$$$$ #$$$$$$ #$$$$$$ #$$$$$$ #$$$$$$ #$$$$$$ #$$$$$$ #$$$$$$
# PANEL 3D

library(gplots)

# LEFT PANEL
yCD=as.numeric(AllData_CD['mRNA_20690_Ctsd',])
xCD=as.numeric(AllData_CD['Pheno_AnimalDeathAge',])
yHF=as.numeric(AllData_HF['mRNA_20690_Ctsd',])
xHF=as.numeric(AllData_HF['Pheno_AnimalDeathAge',])

plotCI(xCD,yCD,cex=1.1, pch=19, col="lightblue", bty="L", ylab=" [log2]", xlab=" [log2]", main="Title", las=1, xlim=c(100,900), ylim=c(min(c(yCD,yHF), na.rm=T)-0.01*min(c(yCD, yHF), na.rm=T), max(c(yCD,yHF), na.rm=T)+0.01*max(c(yCD,yHF), na.rm=T)))

abline(lm(yCD~xCD), col="lightblue")

rhoc=cor(xCD,yCD, method=c("pearson"), use="complete.obs")
pr = cor.test(xCD,yCD,method="pearson")$p.value[1]			
rprint=substitute(paste(italic("r"), " = ", rhoc), list(rhoc=signif(rhoc[1], digits=2)))	
pletter=substitute(paste("   ", italic("p"), " = ", pr), list(pr=signif(pr[1], digits=2)))
legend("topleft", bg='white', bty="n", cex=1, text.col="lightblue", legend=c(as.expression(rprint), as.expression(pletter)))

plotCI(xHF,yHF,cex=1.1, pch=19, col="darkblue", bty="L", ylab=" [log2]", xlab=" [log2]", main="Title", las=1, add=T, ylim=c(min(c(yHF,yHF), na.rm=T)-0.01*min(c(yHF, yHF), na.rm=T), max(c(yHF,yHF), na.rm=T)+0.01*max(c(yHF,yHF), na.rm=T)), xlim=c(min(c(xHF,xHF), na.rm=T)-0.01*min(c(xHF, xHF), na.rm=T), max(c(xHF,xHF), na.rm=T)+0.01*max(c(xHF,xHF), na.rm=T)))
abline(lm(yHF~xHF), col="darkblue")
rhoc=cor(xHF,yHF, method=c("pearson"), use="complete.obs")
pr = cor.test(xHF,yHF,method="pearson")$p.value[1]				
rprint=substitute(paste(italic("r"), " = ", rhoc), list(rhoc=signif(rhoc[1], digits=2)))	
pletter=substitute(paste("   ", italic("p"), " = ", pr), list(pr=signif(pr[1], digits=2)))	
legend("bottomright", bg='white', bty="n", cex=1, text.col="darkblue", legend=c(as.expression(rprint), as.expression(pletter)))

# RIGHT PANEL

yCD=as.numeric(AllData_CD['Prot_13918_Ctsd',])
xCD=as.numeric(AllData_CD['Pheno_AnimalDeathAge',])
yHF=as.numeric(AllData_HF['Prot_13918_Ctsd',])
xHF=as.numeric(AllData_HF['Pheno_AnimalDeathAge',])

plotCI(xCD,yCD,cex=1.1, pch=19, col="orange", bty="L", ylab=" [log2]", xlab=" [log2]", main="Title", las=1, xlim=c(100,900), ylim=c(min(c(yCD,yHF), na.rm=T)-0.01*min(c(yCD, yHF), na.rm=T), max(c(yCD,yHF), na.rm=T)+0.01*max(c(yCD,yHF), na.rm=T)))

abline(lm(yCD~xCD), col="orange")

rhoc=cor(xCD,yCD, method=c("pearson"), use="complete.obs")
pr = cor.test(xCD,yCD,method="pearson")$p.value[1]	
rprint=substitute(paste(italic("r"), " = ", rhoc), list(rhoc=signif(rhoc[1], digits=2)))	
pletter=substitute(paste("   ", italic("p"), " = ", pr), list(pr=signif(pr[1], digits=2)))
legend("topleft", bg='white', bty="n", cex=1, text.col="orange", legend=c(as.expression(rprint), as.expression(pletter)))

plotCI(xHF,yHF,cex=1.1, pch=19, col="red", bty="L", ylab=" [log2]", xlab=" [log2]", main="Title", las=1, add=T, ylim=c(min(c(yHF,yHF), na.rm=T)-0.01*min(c(yHF, yHF), na.rm=T), max(c(yHF,yHF), na.rm=T)+0.01*max(c(yHF,yHF), na.rm=T)), xlim=c(min(c(xHF,xHF), na.rm=T)-0.01*min(c(xHF, xHF), na.rm=T), max(c(xHF,xHF), na.rm=T)+0.01*max(c(xHF,xHF), na.rm=T)))
abline(lm(yHF~xHF), col="red")
rhoc=cor(xHF,yHF, method=c("pearson"), use="complete.obs")
pr = cor.test(xHF,yHF,method="pearson")$p.value[1]			
rprint=substitute(paste(italic("r"), " = ", rhoc), list(rhoc=signif(rhoc[1], digits=2)))
pletter=substitute(paste("   ", italic("p"), " = ", pr), list(pr=signif(pr[1], digits=2)))
legend("bottomright", bg='white', bty="n", cex=1, text.col="red", legend=c(as.expression(rprint), as.expression(pletter)))

#$$$$$$ #$$$$$$ #$$$$$$ #$$$$$$ #$$$$$$ #$$$$$$ #$$$$$$ #$$$$$$
#$$$$$$ #$$$$$$ #$$$$$$ #$$$$$$ #$$$$$$ #$$$$$$ #$$$$$$ #$$$$$$
# PANEL 3E & 3F (both are plotted here) & exported to your active directory
library(xlsx)
library(openxlsx)
library(survival)
library(survminer)
library(tidyverse)

setwd("~/Dropbox/Aging\ BXD\ Study/1\ _\ Manuscript\ -\ Evan\ Multi-Omic/")

# Kaplan-Meier graph using data from Table S5

ctsd_trial1 <- read.xlsx("./aTableS5_Ctsd_Celegans_Lifespan.xlsx",sheet = 2) %>% 
  as_tibble() %>%
  rename(Strain = `Strain./.genotype`,Time = `Time.of.death.(days)`)

ctsd_trial2 <- read.xlsx("./aTableS5_Ctsd_Celegans_Lifespan.xlsx",sheet = 3) %>% 
  as_tibble() %>%
  rename(Strain = `Strain./.genotype`,Time = `Time.of.death.(days)`)

  lifespans <- list("ctsd_trial1" = ctsd_trial1,"ctsd_trial2" = ctsd_trial2)


walk2(lifespans, names(lifespans), ~{
  fit <- survfit(Surv(Time, 1 - Censored) ~ Strain + RNAi, data = .x)
  out_ls <- ggsurvplot(fit = fit,data = .x,conf.int = FALSE,surv.median.line = "hv",palette = c("#482173","#2e6f8e","#29af7f","#b0cc31"))
  plot <- out_ls$plot + 
    coord_cartesian(xlim = c(0,62)) +
    scale_x_continuous(breaks = c(0,20,40,60)) +
    labs(x = "Animal age [days]", title = paste0("Experiment: ",.y)) +
    theme(legend.key.width = unit(1.5, "cm"), legend.position = "right")
  plot %>% ggsave(filename = paste0("./lifespan_",.y,".pdf"),width = 10, height = 6)
})
```
