## Supplementary material for "Multi-Omic Profiling of the Liver Across Diets and Age in a Diverse Mouse Population": All code used to generate figures: b_Figure4_Plotting_Code_vF.r

```
################################################################################  
# First priority is loading the multi-omics dataset, which will be used throughout
## This is the INDIVIDUAL data
################################################################################# 

setwd("~/Dropbox/Aging\ BXD\ Study/1\ _\ Manuscript\ -\ Evan\ Multi-Omic/")

AllData_header=read.table("aTableS2_ALL_DATA.csv", nrows=8, row.names=1, header=T, sep=",")
AllData_header=AllData_header[,3:ncol(AllData_header)]
AllData=read.table("aTableS2_ALL_DATA.csv", row.names=1, header=T,  na.strings="", sep=",", skip=8, stringsAsFactors=FALSE)
AllData=AllData[1:nrow(AllData),3:ncol(AllData)]

AllData_CD=matrix(NA, nrow=nrow(AllData), ncol=length(AllData))
for(i in 1:length(AllData)) {
	if(as.character(AllData_header["Diet",i])=="CD") {   AllData_CD[,i]=as.numeric(as.character(AllData[,i]))	}
} 
rownames(AllData_CD)=rownames(AllData)

AllData_HF=matrix(NA, nrow=nrow(AllData), ncol=length(AllData))
for(i in 1:length(AllData)) {
	if(as.character(AllData_header["Diet",i])=="HF") {   AllData_HF[,i]=as.numeric(as.character(AllData[,i]))	}
}
rownames(AllData_HF)=rownames(AllData)

AllData_young=matrix(NA, nrow=nrow(AllData), ncol=length(AllData))
for(i in 1:length(AllData)) {
	if(as.character(AllData_header["Age",i])<=420) {	AllData_young[,i]=as.numeric(as.character(AllData[,i]))	}
}
rownames(AllData_young)=rownames(AllData)

AllData_old=matrix(NA, nrow=nrow(AllData), ncol=length(AllData))
for(i in 1:length(AllData)) {
	if(as.character(AllData_header["Age",i])>=420) {	AllData_old[,i]=as.numeric(as.character(AllData[,i]))	}
}
rownames(AllData_old)=rownames(AllData)

################################################################################  
# FIGURE FOUR
################################################################################# 

library(imsbInfer)
library(WGCNA)
library(RColorBrewer)
library(ggbiplot)
library(outliers)
library(igraph)
source("~/Dropbox/Aging\ BXD\ Study/1\ _\ Manuscript\ -\ Evan\ Multi-Omic/b_helpFunctions_unilities.R")
source("~/Dropbox/Aging\ BXD\ Study/1\ _\ Manuscript\ -\ Evan\ Multi-Omic/b_helpFunctions.R")

#$$$$$$ #$$$$$$ #$$$$$$ #$$$$$$ #$$$$$$ #$$$$$$ #$$$$$$ #$$$$$$
#$$$$$$ #$$$$$$ #$$$$$$ #$$$$$$ #$$$$$$ #$$$$$$ #$$$$$$ #$$$$$$
# PANEL 4A
# Note, legend is hand-drawn, and the mt-DNA genes are turned to triangles and highlighted in red by hand in Illustrator.
### LEFT PANEL

pep1=AllData
GeneSet=read.csv(file="~/Dropbox/Aging\ BXD\ Study/1\ _\ Manuscript\ -\ Evan\ Multi-Omic/b_HelpNetworks_ALL_NETWORKS_CANON_mRNA_Prot_ALIGNED.csv", header=T, stringsAsFactors = FALSE)

target.cats <- c("mRNA_KEGG_OXPHOS_NoATPase")
GeneSet <- GeneSet[which(GeneSet$Description %in% target.cats),] 
GeneSet1<-GeneSet

# PURELY for visualization, two genes are dropped because they distort the graph. For total edge counts and node counts, they are included
GeneSet1 <- GeneSet1[-17,] # Drops Cox6a2
GeneSet1 <- GeneSet1[-21,] # Drops Cox7a2l; this gene also has a huge cis-QTL DNA variant that makes it a bad candidate for correlations in the BXDs

# FIRST, load the "CorrGraph" function at the end of this file (scroll, or look for "CORRGRAPH.FIND")
CorrGraph(pep1, GeneSet1, 0.0001, colors=c("black","red","green", "yellow","lightblue")) # Input data, input annotation, and p-value cutoff for edges

#########
### RIGHT PANEL
pep1=AllData
GeneSet=read.csv(file="~/Dropbox/Aging\ BXD\ Study/1\ _\ Manuscript\ -\ Evan\ Multi-Omic/b_HelpNetworks_ALL_NETWORKS_CANON_mRNA_Prot_ALIGNED.csv", header=T, stringsAsFactors = FALSE)

target.cats <- c("PROT_KEGG_OXPHOS_NoATPase")
GeneSet <- GeneSet[which(GeneSet$Description %in% target.cats),] 
GeneSet1<-GeneSet

# PURELY for visualization, two genes are dropped because they distort the graph. For total edge counts and node counts, they are included
GeneSet1 <- GeneSet1[-13,] # Drops COX17
GeneSet1 <- GeneSet1[-22,] # Drops COX7A2L; this gene also has a huge cis-QTL DNA variant that makes it a bad candidate for correlations in the BXDs
GeneSet1 <- GeneSet1[-22,] # Drops COX7B (NB: not a typo, since the drop is sequential and reordered)

CorrGraph(pep1, GeneSet1, 0.0001, colors=c("black","red","green", "yellow","lightblue")) # Input data, input annotation, and p-value cutoff for edges


#$$$$$$ #$$$$$$ #$$$$$$ #$$$$$$ #$$$$$$ #$$$$$$ #$$$$$$ #$$$$$$
#$$$$$$ #$$$$$$ #$$$$$$ #$$$$$$ #$$$$$$ #$$$$$$ #$$$$$$ #$$$$$$
# PANEL 4B
### LEFT PANEL

# We have to now generate all the correlation values from Figure 4A and store these in variables. 
pep1=AllData

# Now we store all correlation values for the networks so that we can make a density plot later
GeneSet=read.csv(file="~/Dropbox/Aging\ BXD\ Study/1\ _\ Manuscript\ -\ Evan\ Multi-Omic/b_HelpNetworks_ALL_NETWORKS_CANON_mRNA_Prot_ALIGNED.csv", header=T, stringsAsFactors = FALSE)
target.cats <- c("mRNA_KEGG_OXPHOS_NoATPase")
GeneSet <- GeneSet[which(GeneSet$Description %in% target.cats),] 
GeneSet1<-GeneSet
tmp = rownames(pep1)
pep1 = pep1[tmp,]
GeneSet1 = GeneSet1[GeneSet1[,3] != "",]	# The first column in the data file must match the third column in the "gene list" file!
rnams = as.character(rownames(pep1))
xx = getMatchingRows(GeneSet1[,3], rnams) # Get the row number of matching rows
GeneMatch = pep1[xx,]
GeneMatch = GeneMatch[order(rownames(GeneMatch)),] 
transform_GeneMatch=apply(t(GeneMatch), c(1,2), as.numeric) # If NAs are coerced in, then it will force everything into a character matrix
tmpr <- corAndPvalue(transform_GeneMatch,method="spearman")
tmp = tmpr$cor
diag(tmp) = 0
outputmatrix_mRNA_OXPHOS_ALL_CORRS=tmp[lower.tri(tmp)] 		

GeneSet=read.csv(file="~/Dropbox/Aging\ BXD\ Study/1\ _\ Manuscript\ -\ Evan\ Multi-Omic/b_HelpNetworks_ALL_NETWORKS_CANON_mRNA_Prot_ALIGNED.csv", header=T, stringsAsFactors = FALSE)
target.cats <- c("PROT_KEGG_OXPHOS_NoATPase")
GeneSet <- GeneSet[which(GeneSet$Description %in% target.cats),] 
GeneSet1<-GeneSet
tmp = rownames(pep1)
pep1 = pep1[tmp,]
GeneSet1 = GeneSet1[GeneSet1[,3] != "",]	# The first column in the data file must match the third column in the "gene list" file!
rnams = as.character(rownames(pep1))
xx = getMatchingRows(GeneSet1[,3], rnams) # Get the row number of matching rows
GeneMatch = pep1[xx,]
GeneMatch = GeneMatch[order(rownames(GeneMatch)),] 
transform_GeneMatch=apply(t(GeneMatch), c(1,2), as.numeric) # If NAs are coerced in, then it will force everything into a character matrix
tmpr <- corAndPvalue(transform_GeneMatch,method="spearman")
tmp = tmpr$cor
diag(tmp) = 0
outputmatrix_PROT_OXPHOS_ALL_CORRS=tmp[lower.tri(tmp)] 		

GeneSet=read.csv(file="~/Dropbox/Aging\ BXD\ Study/1\ _\ Manuscript\ -\ Evan\ Multi-Omic/b_HelpNetworks_ALL_NETWORKS_CANON_mRNA_Prot_ALIGNED.csv", header=T, stringsAsFactors = FALSE)
target.cats <- c("mRNA_KEGG_OXPHOS_NoATPase","PROT_KEGG_OXPHOS_NoATPase")
GeneSet <- GeneSet[which(GeneSet$Description %in% target.cats),] 
GeneSet1<-GeneSet
tmp = rownames(pep1)
pep1 = pep1[tmp,]
GeneSet1 = GeneSet1[GeneSet1[,3] != "",]	# The first column in the data file must match the third column in the "gene list" file!
rnams = as.character(rownames(pep1))
xx = getMatchingRows(GeneSet1[,3], rnams) # Get the row number of matching rows
GeneMatch = pep1[xx,]
GeneMatch = GeneMatch[order(rownames(GeneMatch)),] 
transform_GeneMatch=apply(t(GeneMatch), c(1,2), as.numeric) # If NAs are coerced in, then it will force everything into a character matrix
tmpr <- corAndPvalue(transform_GeneMatch,method="spearman")
tmp = tmpr$cor
diag(tmp) = 0
outputmatrix_Both_OXPHOS_ALL_CORRS=tmp[lower.tri(tmp)] 	
outputmatrix_Both_OXPHOS_ALL_CORRS2=outputmatrix_Both_OXPHOS_ALL_CORRS[!(outputmatrix_Both_OXPHOS_ALL_CORRS %in% outputmatrix_mRNA_OXPHOS_ALL_CORRS)]
outputmatrix_Both_OXPHOS_ALL_CORRS3=outputmatrix_Both_OXPHOS_ALL_CORRS2[!(outputmatrix_Both_OXPHOS_ALL_CORRS2 %in% outputmatrix_PROT_OXPHOS_ALL_CORRS)]
### NOTE: This "cross-layer" code has to be calculated last, and it suppresses the correlations that were found within protein and within mRNA, only retaining "across"

# Now we make the density plot
library(gplots)

plot(density(outputmatrix_Both_OXPHOS_ALL_CORRS3, na.rm=T), xlim=c(-1,1), ylim=c(0, 6.2), col="black")
plotCI(density(outputmatrix_mRNA_OXPHOS_ALL_CORRS), col="lightblue", add=T, type="l")
plotCI(density(outputmatrix_PROT_OXPHOS_ALL_CORRS), col="red", add=T, type="l")

### RIGHT PANEL
## Now we just need to show what the 'negative control' distribution is, and show what ALL correlations look like. So we just use the entire gene list for the paired 3772 genes with mRNA and protein data
GeneSet=read.csv(file="~/Dropbox/Aging\ BXD\ Study/1\ _\ Manuscript\ -\ Evan\ Multi-Omic/b_HelpNetworks_ALL_NETWORKS_CANON_mRNA_Prot_ALIGNED.csv", header=T, stringsAsFactors = FALSE)
target.cats <- c("mRNA_ALL")
GeneSet <- GeneSet[which(GeneSet$Description %in% target.cats),] 
GeneSet1<-GeneSet
tmp = rownames(pep1)
pep1 = pep1[tmp,]
GeneSet1 = GeneSet1[GeneSet1[,3] != "",]	# The first column in the data file must match the third column in the "gene list" file!
rnams = as.character(rownames(pep1))
xx = getMatchingRows(GeneSet1[,3], rnams) # Get the row number of matching rows
GeneMatch = pep1[xx,]
GeneMatch = GeneMatch[order(rownames(GeneMatch)),] 
transform_GeneMatch=apply(t(GeneMatch), c(1,2), as.numeric) # If NAs are coerced in, then it will force everything into a character matrix
tmpr <- corAndPvalue(transform_GeneMatch,method="spearman")
tmp = tmpr$cor
diag(tmp) = 0
outputmatrix_mRNA_ALL_CORRS=tmp[lower.tri(tmp)] 	

GeneSet=read.csv(file="~/Dropbox/Aging\ BXD\ Study/1\ _\ Manuscript\ -\ Evan\ Multi-Omic/b_HelpNetworks_ALL_NETWORKS_CANON_mRNA_Prot_ALIGNED.csv", header=T, stringsAsFactors = FALSE)
target.cats <- c("PROT_ALL")
GeneSet <- GeneSet[which(GeneSet$Description %in% target.cats),] 
GeneSet1<-GeneSet
tmp = rownames(pep1)
pep1 = pep1[tmp,]
GeneSet1 = GeneSet1[GeneSet1[,3] != "",]	# The first column in the data file must match the third column in the "gene list" file!
rnams = as.character(rownames(pep1))
xx = getMatchingRows(GeneSet1[,3], rnams) # Get the row number of matching rows
GeneMatch = pep1[xx,]
GeneMatch = GeneMatch[order(rownames(GeneMatch)),] 
transform_GeneMatch=apply(t(GeneMatch), c(1,2), as.numeric) # If NAs are coerced in, then it will force everything into a character matrix
tmpr <- corAndPvalue(transform_GeneMatch,method="spearman")
tmp = tmpr$cor
diag(tmp) = 0
outputmatrix_PROT_ALL_CORRS=tmp[lower.tri(tmp)] 	

GeneSet=read.csv(file="~/Dropbox/Aging\ BXD\ Study/1\ _\ Manuscript\ -\ Evan\ Multi-Omic/b_HelpNetworks_ALL_NETWORKS_CANON_mRNA_Prot_ALIGNED.csv", header=T, stringsAsFactors = FALSE)
target.cats <- c("mRNA_ALL","PROT_ALL")
GeneSet <- GeneSet[which(GeneSet$Description %in% target.cats),] 
GeneSet1<-GeneSet
tmp = rownames(pep1)
pep1 = pep1[tmp,]
GeneSet1 = GeneSet1[GeneSet1[,3] != "",]	# The first column in the data file must match the third column in the "gene list" file!
rnams = as.character(rownames(pep1))
xx = getMatchingRows(GeneSet1[,3], rnams) # Get the row number of matching rows
GeneMatch = pep1[xx,]
GeneMatch = GeneMatch[order(rownames(GeneMatch)),] 
transform_GeneMatch=apply(t(GeneMatch), c(1,2), as.numeric) # If NAs are coerced in, then it will force everything into a character matrix
tmpr <- corAndPvalue(transform_GeneMatch,method="spearman")
tmp = tmpr$cor
diag(tmp) = 0
outputmatrix_ALL_ALL_CORRS=tmp[lower.tri(tmp)] 
outputmatrix_ALL_ALL_CORRS2=outputmatrix_ALL_ALL_CORRS[!(outputmatrix_ALL_ALL_CORRS %in% outputmatrix_mRNA_ALL_CORRS)]
outputmatrix_ALL_ALL_CORRS3=outputmatrix_ALL_ALL_CORRS2[!(outputmatrix_ALL_ALL_CORRS2 %in% outputmatrix_PROT_ALL_CORRS)]

### NOTE: This "cross-layer" code has to be calculated last, and it suppresses the correlations that were found within protein and within mRNA, only retaining "across"

# Now we make the density plot
library(gplots)

plot(density(outputmatrix_ALL_ALL_CORRS3, na.rm=T), xlim=c(-1,1), ylim=c(0, 6.2), col="black")
plotCI(density(outputmatrix_mRNA_ALL_CORRS), col="lightblue", add=T, type="l")
plotCI(density(outputmatrix_PROT_ALL_CORRS), col="red", add=T, type="l")


#$$$$$$ #$$$$$$ #$$$$$$ #$$$$$$ #$$$$$$ #$$$$$$ #$$$$$$ #$$$$$$
#$$$$$$ #$$$$$$ #$$$$$$ #$$$$$$ #$$$$$$ #$$$$$$ #$$$$$$ #$$$$$$
# PANEL 4C

pep1=AllData
GeneSet=read.csv(file="~/Dropbox/Aging\ BXD\ Study/1\ _\ Manuscript\ -\ Evan\ Multi-Omic/b_HelpNetworks_ALL_NETWORKS_CANON_mRNA_Prot_ALIGNED.csv", header=T, stringsAsFactors = FALSE)

target.cats <- c("mRNA_REACTOME_CHOLESTEROL_BIOSYNTHESIS","PROT_REACTOME_CHOLESTEROL_BIOSYNTHESIS")
GeneSet <- GeneSet[which(GeneSet$Description %in% target.cats),] 
GeneSet1<-GeneSet

# PURELY for visualization, LBR protein is dropped. It is included in the edge and node counts.
GeneSet1 <- GeneSet1[-17,] # Drops LBR protein

# FIRST, load the "CorrGraph" function at the end of this file (scroll, or look for "CORRGRAPH.FIND")
CorrGraph(pep1, GeneSet1, 0.0001, colors=c("red","blue")) # Input data, input annotation, and p-value cutoff for edges


#$$$$$$ #$$$$$$ #$$$$$$ #$$$$$$ #$$$$$$ #$$$$$$ #$$$$$$ #$$$$$$
#$$$$$$ #$$$$$$ #$$$$$$ #$$$$$$ #$$$$$$ #$$$$$$ #$$$$$$ #$$$$$$
# PANEL 4D
pep1=AllData

GeneSet=read.csv(file="~/Dropbox/Aging\ BXD\ Study/1\ _\ Manuscript\ -\ Evan\ Multi-Omic/b_HelpNetworks_ALL_NETWORKS_CANON_mRNA_Prot_ALIGNED.csv", header=T, stringsAsFactors = FALSE)
target.cats <- c("mRNA_REACTOME_CHOLESTEROL_BIOSYNTHESIS")
GeneSet <- GeneSet[which(GeneSet$Description %in% target.cats),] 
GeneSet1<-GeneSet
tmp = rownames(pep1)
pep1 = pep1[tmp,]
GeneSet1 = GeneSet1[GeneSet1[,3] != "",]	# The first column in the data file must match the third column in the "gene list" file!
rnams = as.character(rownames(pep1))
xx = getMatchingRows(GeneSet1[,3], rnams) # Get the row number of matching rows
GeneMatch = pep1[xx,]
GeneMatch = GeneMatch[order(rownames(GeneMatch)),] 
transform_GeneMatch=apply(t(GeneMatch), c(1,2), as.numeric) # If NAs are coerced in, then it will force everything into a character matrix
tmpr <- corAndPvalue(transform_GeneMatch,method="spearman")
tmp = tmpr$cor
diag(tmp) = 0
outputmatrix_mRNA_Cholesterol_CORRS=tmp[lower.tri(tmp)] 	

GeneSet=read.csv(file="~/Dropbox/Aging\ BXD\ Study/1\ _\ Manuscript\ -\ Evan\ Multi-Omic/b_HelpNetworks_ALL_NETWORKS_CANON_mRNA_Prot_ALIGNED.csv", header=T, stringsAsFactors = FALSE)
target.cats <- c("PROT_REACTOME_CHOLESTEROL_BIOSYNTHESIS")
GeneSet <- GeneSet[which(GeneSet$Description %in% target.cats),] 
GeneSet1<-GeneSet
tmp = rownames(pep1)
pep1 = pep1[tmp,]
GeneSet1 = GeneSet1[GeneSet1[,3] != "",]	# The first column in the data file must match the third column in the "gene list" file!
rnams = as.character(rownames(pep1))
xx = getMatchingRows(GeneSet1[,3], rnams) # Get the row number of matching rows
GeneMatch = pep1[xx,]
GeneMatch = GeneMatch[order(rownames(GeneMatch)),] 
transform_GeneMatch=apply(t(GeneMatch), c(1,2), as.numeric) # If NAs are coerced in, then it will force everything into a character matrix
tmpr <- corAndPvalue(transform_GeneMatch,method="spearman")
tmp = tmpr$cor
diag(tmp) = 0
outputmatrix_PROT_Cholesterol_CORRS=tmp[lower.tri(tmp)] 	

GeneSet=read.csv(file="~/Dropbox/Aging\ BXD\ Study/1\ _\ Manuscript\ -\ Evan\ Multi-Omic/b_HelpNetworks_ALL_NETWORKS_CANON_mRNA_Prot_ALIGNED.csv", header=T, stringsAsFactors = FALSE)
target.cats <- c("mRNA_REACTOME_CHOLESTEROL_BIOSYNTHESIS","PROT_REACTOME_CHOLESTEROL_BIOSYNTHESIS")
GeneSet <- GeneSet[which(GeneSet$Description %in% target.cats),] 
GeneSet1<-GeneSet
tmp = rownames(pep1)
pep1 = pep1[tmp,]
GeneSet1 = GeneSet1[GeneSet1[,3] != "",]	# The first column in the data file must match the third column in the "gene list" file!
rnams = as.character(rownames(pep1))
xx = getMatchingRows(GeneSet1[,3], rnams) # Get the row number of matching rows
GeneMatch = pep1[xx,]
GeneMatch = GeneMatch[order(rownames(GeneMatch)),] 
transform_GeneMatch=apply(t(GeneMatch), c(1,2), as.numeric) # If NAs are coerced in, then it will force everything into a character matrix
tmpr <- corAndPvalue(transform_GeneMatch,method="spearman")
tmp = tmpr$cor
diag(tmp) = 0
outputmatrix_ALL_Cholesterol_CORRS=tmp[lower.tri(tmp)] 
outputmatrix_ALL_Cholesterol_CORRS2=outputmatrix_ALL_Cholesterol_CORRS[!(outputmatrix_ALL_Cholesterol_CORRS %in% outputmatrix_mRNA_Cholesterol_CORRS)]
outputmatrix_ALL_Cholesterol_CORRS3=outputmatrix_ALL_Cholesterol_CORRS2[!(outputmatrix_ALL_Cholesterol_CORRS2 %in% outputmatrix_PROT_Cholesterol_CORRS)]

### NOTE: This "cross-layer" code has to be calculated last, and it suppresses the correlations that were found within protein and within mRNA, only retaining "across"

# Now we make the density plot
library(gplots)

plot(density(outputmatrix_ALL_Cholesterol_CORRS3, na.rm=T), xlim=c(-1,1), ylim=c(0, 6), col="black")
plotCI(density(outputmatrix_mRNA_Cholesterol_CORRS), col="lightblue", add=T, type="l")
plotCI(density(outputmatrix_PROT_Cholesterol_CORRS), col="red", add=T, type="l")

#$$$$$$ #$$$$$$ #$$$$$$ #$$$$$$ #$$$$$$ #$$$$$$ #$$$$$$ #$$$$$$
#$$$$$$ #$$$$$$ #$$$$$$ #$$$$$$ #$$$$$$ #$$$$$$ #$$$$$$ #$$$$$$
# PANEL 4E

# LEFT PANEL (mRNA)
pep1=AllData
GeneSet=read.csv(file="~/Dropbox/Aging\ BXD\ Study/1\ _\ Manuscript\ -\ Evan\ Multi-Omic/b_HelpNetworks_ALL_NETWORKS_CANON_mRNA_Prot_ALIGNED.csv", header=T, stringsAsFactors = FALSE)

target.cats <- c("mRNA_REACTOME_CHOLESTEROL_BIOSYNTHESIS","mRNA_REACTOME_MITOCHONDRIAL_FATTY_ACID_BETA_OXIDATION")
GeneSet <- GeneSet[which(GeneSet$Description %in% target.cats),] 
GeneSet1<-GeneSet

# PURELY for visualization, LBR protein is dropped. It is included in the edge and node counts.
#GeneSet1 <- GeneSet1[-17,] # Drops LBR protein

# FIRST, load the "CorrGraph" function at the end of this file (scroll, or look for "CORRGRAPH.FIND")
CorrGraph(pep1, GeneSet1, 0.0001, colors=c("red","blue","orange")) # Input data, input annotation, and p-value cutoff for edges

# RIGHT PANEL (Protein)
pep1=AllData
GeneSet=read.csv(file="~/Dropbox/Aging\ BXD\ Study/1\ _\ Manuscript\ -\ Evan\ Multi-Omic/b_HelpNetworks_ALL_NETWORKS_CANON_mRNA_Prot_ALIGNED.csv", header=T, stringsAsFactors = FALSE)

target.cats <- c("PROT_REACTOME_CHOLESTEROL_BIOSYNTHESIS","PROT_REACTOME_MITOCHONDRIAL_FATTY_ACID_BETA_OXIDATION")
GeneSet <- GeneSet[which(GeneSet$Description %in% target.cats),] 
GeneSet1<-GeneSet

# PURELY for visualization, LBR protein is dropped. It is included in the edge and node counts.
#GeneSet1 <- GeneSet1[-17,] # Drops LBR protein

# FIRST, load the "CorrGraph" function at the end of this file (scroll, or look for "CORRGRAPH.FIND")
CorrGraph(pep1, GeneSet1, 0.0001, colors=c("red","blue","orange")) # Input data, input annotation, and p-value cutoff for edges


#$$$$$$ #$$$$$$ #$$$$$$ #$$$$$$ #$$$$$$ #$$$$$$ #$$$$$$ #$$$$$$
#$$$$$$ #$$$$$$ #$$$$$$ #$$$$$$ #$$$$$$ #$$$$$$ #$$$$$$ #$$$$$$
# PANEL 4F

## LEFT PANEL (mRNA)
pep1=AllData

GeneSet=read.csv(file="~/Dropbox/Aging\ BXD\ Study/1\ _\ Manuscript\ -\ Evan\ Multi-Omic/b_HelpNetworks_ALL_NETWORKS_CANON_mRNA_Prot_ALIGNED.csv", header=T, stringsAsFactors = FALSE)
target.cats <- c("mRNA_REACTOME_CHOLESTEROL_BIOSYNTHESIS")
GeneSet <- GeneSet[which(GeneSet$Description %in% target.cats),] 
GeneSet1<-GeneSet
tmp = rownames(pep1)
pep1 = pep1[tmp,]
GeneSet1 = GeneSet1[GeneSet1[,3] != "",]	# The first column in the data file must match the third column in the "gene list" file!
rnams = as.character(rownames(pep1))
xx = getMatchingRows(GeneSet1[,3], rnams) # Get the row number of matching rows
GeneMatch = pep1[xx,]
GeneMatch = GeneMatch[order(rownames(GeneMatch)),] 
transform_GeneMatch=apply(t(GeneMatch), c(1,2), as.numeric) # If NAs are coerced in, then it will force everything into a character matrix
tmpr <- corAndPvalue(transform_GeneMatch,method="spearman")
tmp = tmpr$cor
diag(tmp) = 0
outputmatrix_mRNA_Cholesterol_CORRS=tmp[lower.tri(tmp)] 	

GeneSet=read.csv(file="~/Dropbox/Aging\ BXD\ Study/1\ _\ Manuscript\ -\ Evan\ Multi-Omic/b_HelpNetworks_ALL_NETWORKS_CANON_mRNA_Prot_ALIGNED.csv", header=T, stringsAsFactors = FALSE)
target.cats <- c("mRNA_REACTOME_MITOCHONDRIAL_FATTY_ACID_BETA_OXIDATION")
GeneSet <- GeneSet[which(GeneSet$Description %in% target.cats),] 
GeneSet1<-GeneSet
tmp = rownames(pep1)
pep1 = pep1[tmp,]
GeneSet1 = GeneSet1[GeneSet1[,3] != "",]	# The first column in the data file must match the third column in the "gene list" file!
rnams = as.character(rownames(pep1))
xx = getMatchingRows(GeneSet1[,3], rnams) # Get the row number of matching rows
GeneMatch = pep1[xx,]
GeneMatch = GeneMatch[order(rownames(GeneMatch)),] 
transform_GeneMatch=apply(t(GeneMatch), c(1,2), as.numeric) # If NAs are coerced in, then it will force everything into a character matrix
tmpr <- corAndPvalue(transform_GeneMatch,method="spearman")
tmp = tmpr$cor
diag(tmp) = 0
outputmatrix_mRNA_Box_CORRS=tmp[lower.tri(tmp)] 	

GeneSet=read.csv(file="~/Dropbox/Aging\ BXD\ Study/1\ _\ Manuscript\ -\ Evan\ Multi-Omic/b_HelpNetworks_ALL_NETWORKS_CANON_mRNA_Prot_ALIGNED.csv", header=T, stringsAsFactors = FALSE)
target.cats <- c("mRNA_REACTOME_CHOLESTEROL_BIOSYNTHESIS","mRNA_REACTOME_MITOCHONDRIAL_FATTY_ACID_BETA_OXIDATION")
GeneSet <- GeneSet[which(GeneSet$Description %in% target.cats),] 
GeneSet1<-GeneSet
tmp = rownames(pep1)
pep1 = pep1[tmp,]
GeneSet1 = GeneSet1[GeneSet1[,3] != "",]	# The first column in the data file must match the third column in the "gene list" file!
rnams = as.character(rownames(pep1))
xx = getMatchingRows(GeneSet1[,3], rnams) # Get the row number of matching rows
GeneMatch = pep1[xx,]
GeneMatch = GeneMatch[order(rownames(GeneMatch)),] 
transform_GeneMatch=apply(t(GeneMatch), c(1,2), as.numeric) # If NAs are coerced in, then it will force everything into a character matrix
tmpr <- corAndPvalue(transform_GeneMatch,method="spearman")
tmp = tmpr$cor
diag(tmp) = 0
outputmatrix_mRNA_CholesterolBOX_CORRS=tmp[lower.tri(tmp)] 
outputmatrix_mRNA_CholesterolBOX_CORRS2=outputmatrix_mRNA_CholesterolBOX_CORRS[!(outputmatrix_mRNA_CholesterolBOX_CORRS %in% outputmatrix_mRNA_Cholesterol_CORRS)]
outputmatrix_mRNA_CholesterolBOX_CORRS3=outputmatrix_mRNA_CholesterolBOX_CORRS2[!(outputmatrix_mRNA_CholesterolBOX_CORRS2 %in% outputmatrix_mRNA_Box_CORRS)]

### NOTE: This "cross-layer" code has to be calculated last, and it suppresses the correlations that were found within protein and within mRNA, only retaining "across"

# Now we make the density plot
library(gplots)

plot(density(outputmatrix_mRNA_CholesterolBOX_CORRS3, na.rm=T), xlim=c(-1,1), ylim=c(0, 6), col="black")
plotCI(density(outputmatrix_mRNA_Cholesterol_CORRS), col="red", add=T, type="l")
plotCI(density(outputmatrix_mRNA_Box_CORRS), col="yellow", add=T, type="l")

## RIGHT PANEL (Protein)
pep1=AllData

GeneSet=read.csv(file="~/Dropbox/Aging\ BXD\ Study/1\ _\ Manuscript\ -\ Evan\ Multi-Omic/b_HelpNetworks_ALL_NETWORKS_CANON_mRNA_Prot_ALIGNED.csv", header=T, stringsAsFactors = FALSE)
target.cats <- c("PROT_REACTOME_CHOLESTEROL_BIOSYNTHESIS")
GeneSet <- GeneSet[which(GeneSet$Description %in% target.cats),] 
GeneSet1<-GeneSet
tmp = rownames(pep1)
pep1 = pep1[tmp,]
GeneSet1 = GeneSet1[GeneSet1[,3] != "",]	# The first column in the data file must match the third column in the "gene list" file!
rnams = as.character(rownames(pep1))
xx = getMatchingRows(GeneSet1[,3], rnams) # Get the row number of matching rows
GeneMatch = pep1[xx,]
GeneMatch = GeneMatch[order(rownames(GeneMatch)),] 
transform_GeneMatch=apply(t(GeneMatch), c(1,2), as.numeric) # If NAs are coerced in, then it will force everything into a character matrix
tmpr <- corAndPvalue(transform_GeneMatch,method="spearman")
tmp = tmpr$cor
diag(tmp) = 0
outputmatrix_PROT_Cholesterol_CORRS=tmp[lower.tri(tmp)] 	

GeneSet=read.csv(file="~/Dropbox/Aging\ BXD\ Study/1\ _\ Manuscript\ -\ Evan\ Multi-Omic/b_HelpNetworks_ALL_NETWORKS_CANON_mRNA_Prot_ALIGNED.csv", header=T, stringsAsFactors = FALSE)
target.cats <- c("PROT_REACTOME_MITOCHONDRIAL_FATTY_ACID_BETA_OXIDATION")
GeneSet <- GeneSet[which(GeneSet$Description %in% target.cats),] 
GeneSet1<-GeneSet
tmp = rownames(pep1)
pep1 = pep1[tmp,]
GeneSet1 = GeneSet1[GeneSet1[,3] != "",]	# The first column in the data file must match the third column in the "gene list" file!
rnams = as.character(rownames(pep1))
xx = getMatchingRows(GeneSet1[,3], rnams) # Get the row number of matching rows
GeneMatch = pep1[xx,]
GeneMatch = GeneMatch[order(rownames(GeneMatch)),] 
transform_GeneMatch=apply(t(GeneMatch), c(1,2), as.numeric) # If NAs are coerced in, then it will force everything into a character matrix
tmpr <- corAndPvalue(transform_GeneMatch,method="spearman")
tmp = tmpr$cor
diag(tmp) = 0
outputmatrix_PROT_Box_CORRS=tmp[lower.tri(tmp)] 	

GeneSet=read.csv(file="~/Dropbox/Aging\ BXD\ Study/1\ _\ Manuscript\ -\ Evan\ Multi-Omic/b_HelpNetworks_ALL_NETWORKS_CANON_mRNA_Prot_ALIGNED.csv", header=T, stringsAsFactors = FALSE)
target.cats <- c("PROT_REACTOME_CHOLESTEROL_BIOSYNTHESIS","PROT_REACTOME_MITOCHONDRIAL_FATTY_ACID_BETA_OXIDATION")
GeneSet <- GeneSet[which(GeneSet$Description %in% target.cats),] 
GeneSet1<-GeneSet
tmp = rownames(pep1)
pep1 = pep1[tmp,]
GeneSet1 = GeneSet1[GeneSet1[,3] != "",]	# The first column in the data file must match the third column in the "gene list" file!
rnams = as.character(rownames(pep1))
xx = getMatchingRows(GeneSet1[,3], rnams) # Get the row number of matching rows
GeneMatch = pep1[xx,]
GeneMatch = GeneMatch[order(rownames(GeneMatch)),] 
transform_GeneMatch=apply(t(GeneMatch), c(1,2), as.numeric) # If NAs are coerced in, then it will force everything into a character matrix
tmpr <- corAndPvalue(transform_GeneMatch,method="spearman")
tmp = tmpr$cor
diag(tmp) = 0
outputmatrix_PROT_CholesterolBOX_CORRS=tmp[lower.tri(tmp)] 
outputmatrix_PROT_CholesterolBOX_CORRS2=outputmatrix_PROT_CholesterolBOX_CORRS[!(outputmatrix_PROT_CholesterolBOX_CORRS %in% outputmatrix_PROT_Cholesterol_CORRS)]
outputmatrix_PROT_CholesterolBOX_CORRS3=outputmatrix_PROT_CholesterolBOX_CORRS2[!(outputmatrix_PROT_CholesterolBOX_CORRS2 %in% outputmatrix_PROT_Box_CORRS)]

### NOTE: This "cross-layer" code has to be calculated last, and it suppresses the correlations that were found within protein and within mRNA, only retaining "across"

# Now we make the density plot
library(gplots)

plot(density(outputmatrix_PROT_CholesterolBOX_CORRS3, na.rm=T), xlim=c(-1,1), ylim=c(0, 6), col="black")
plotCI(density(outputmatrix_PROT_Cholesterol_CORRS), col="red", add=T, type="l")
plotCI(density(outputmatrix_PROT_Box_CORRS), col="yellow", add=T, type="l")


#$$$$$$ #$$$$$$ #$$$$$$ #$$$$$$ #$$$$$$ #$$$$$$ #$$$$$$ #$$$$$$
#$$$$$$ #$$$$$$ #$$$$$$ #$$$$$$ #$$$$$$ #$$$$$$ #$$$$$$ #$$$$$$
# PANEL 4G

# These figures are output from GSEA using the mRNA and protein datasets and the selected gene sets
# The gene sets are manually entered in GSEA for target sets, using these sets:
# OXPHOS: Atp5a1, Atp5b, Atp5c1, Atp5d, Atp5e, Atp5f1, Atp5h, Atp5j, Atp5j2, Atp5l, Atp5o, Cox17, Cox4I1, Cox5a, Cox5b, Cox6a1, Cox6a2, Cox6b1, Cox6c, Cox7a1, Cox7a2, Cox7a2l, Cox7b, Cox7c, Cyc1, mt-ATP8, mt-CYTB, mt-ND1, mt-ND3, mt-ND5, Ndufa1, Ndufa10, Ndufa11, Ndufa2, Ndufa3, Ndufa4, Ndufa5, Ndufa6, Ndufa7, Ndufa8, Ndufa9, Ndufab1, Ndufb10, Ndufb2, Ndufb3, Ndufb4, Ndufb5, Ndufb6, Ndufb7, Ndufb8, Ndufb9, Ndufc2, Ndufs1, Ndufs2, Ndufs3, Ndufs4, Ndufs5, Ndufs6, Ndufs7, Ndufs8, Ndufv1, Ndufv2, Ndufv3, Sdha, Sdhb, Sdhc, Sdhd, Uqcr10, Uqcr11, Uqcrb, Uqcrc1, Uqcrc2, Uqcrfs1, Uqcrh, Uqcrq
# Cholesterol: Tm7sf2, Pmvk, Cyp51, Nsdhl, Hsd17b7, Ebp, Idi1, Sqle, Fdps, Sc4mol, Hmgcs1, Lss, Dhcr7, Mvk, Fdft1, Lbr, Dhcr24
# Using GSEA version 4, standalone download

#$$$$$$ #$$$$$$ #$$$$$$ #$$$$$$ #$$$$$$ #$$$$$$ #$$$$$$ #$$$$$$
#$$$$$$ #$$$$$$ #$$$$$$ #$$$$$$ #$$$$$$ #$$$$$$ #$$$$$$ #$$$$$$
# PANEL 4H
# Recall: need to load the AllData at top. 

# ggbiplot is on github:
# library(devtools)
# install_github("vqv/ggbiplot")

library(ggbiplot)
# LEFT PANEL: 
# OXPHOS protein data, separated by age cohort
# First let's go through all the animals' ages again and assign them as "young" or "old" again using the 419 day cutoff. This is for all 315 samples with protein data.
groupings_Age_Prot_category=NA
for(i in 1:315) {
	if (AllData_header[4,i]<=419) {
		groupings_Age_Prot_category[i]="young"
	}
	else
		groupings_Age_Prot_category[i]="old"
}

inputdat = rbind(AllData['Prot_13610_Atp5a1',],AllData['Prot_14033_Atp5b',],AllData['Prot_9644_Atp5c1',],AllData['Prot_6141_Atp5d',],AllData['Prot_17338_Atp5e',],AllData['Prot_6042_Atp5f1',],AllData['Prot_10960_Atp5h',],AllData['Prot_3917_Atp5j',],AllData['Prot_1953_Atp5j2',],AllData['Prot_13095_Atp5l',],AllData['Prot_9181_Atp5o',],AllData['Prot_7093_ATP8',],AllData['Prot_46625_Cox17',],AllData['Prot_1029_Cox4i1',],AllData['Prot_37952_Cox5a',],AllData['Prot_12175_Cox5b',],AllData['Prot_22888_Cox6a1',],AllData['Prot_57839_Cox6a2',],AllData['Prot_11344_Cox6b1',],AllData['Prot_1524_Cox6c',],AllData['Prot_19095_Cox7a1',],AllData['Prot_4783_Cox7a2',],AllData['Prot_49184_Cox7b',],AllData['Prot_9842_Cox7c',],AllData['Prot_5101_Cyc1',],AllData['Prot_2542_CYTB',],AllData['Prot_30795_ND1',],AllData['Prot_4384_ND3',],AllData['Prot_55212_ND5',],AllData['Prot_64922_Ndufa1',],AllData['Prot_38990_Ndufa10',],AllData['Prot_7917_Ndufa11',],AllData['Prot_12709_Ndufa2',],AllData['Prot_5697_Ndufa3',],AllData['Prot_7682_Ndufa4',],AllData['Prot_12590_Ndufa5',],AllData['Prot_65801_Ndufa6',],AllData['Prot_46413_Ndufa7',],AllData['Prot_15321_Ndufa8',],AllData['Prot_1461_Ndufa9',],AllData['Prot_37742_Ndufab1',],AllData['Prot_14614_Ndufb10',],AllData['Prot_69050_Ndufb2',],AllData['Prot_25796_Ndufb3',],AllData['Prot_68300_Ndufb4',],AllData['Prot_12279_Ndufb5',],AllData['Prot_26002_Ndufb6',],AllData['Prot_9767_Ndufb7',],AllData['Prot_19049_Ndufb8',],AllData['Prot_4236_Ndufb9',],AllData['Prot_80796_Ndufc2',],AllData['Prot_12857_Ndufs1',],AllData['Prot_60482_Ndufs2',],AllData['Prot_11334_Ndufs3',],AllData['Prot_10026_Ndufs4',],AllData['Prot_6163_Ndufs5',],AllData['Prot_6387_Ndufs6',],AllData['Prot_7803_Ndufs7',],AllData['Prot_3464_Ndufs8',],AllData['Prot_6244_Ndufv1',],AllData['Prot_10114_Ndufv2',],AllData['Prot_59191_Ndufv3',],AllData['Prot_10037_Sdha',],AllData['Prot_8734_Sdhb',],AllData['Prot_5796_Sdhc',],AllData['Prot_5833_Sdhd',],AllData['Prot_41297_Uqcr10',],AllData['Prot_8264_Uqcr11',],AllData['Prot_3263_Uqcrb',],AllData['Prot_9568_Uqcrc1',],AllData['Prot_33848_Uqcrc2',],AllData['Prot_11197_Uqcrfs1',],AllData['Prot_11590_Uqcrh',],AllData['Prot_5926_Uqcrq',])
inputdat2 <- inputdat[,colSums(is.na(inputdat))<1] # only take if NO NAs
fit=prcomp(t(inputdat2), scale=FALSE)
summary(fit)
ggbiplot(fit, groups = groupings_Age_Prot_category, ellipse = TRUE, circle = TRUE, scale=1, var.axes=FALSE, choices=c(1,2)) # choices is for which PCs


# RIGHT PANEL:
# Cholesterol protein data, separated by diet. Does not need to be a loop at all here, but is used as such for consistency.
groupings_Diet_Prot_category=NA
for(i in 1:315) {
	if (AllData_header[5,i]=="CD") {
		groupings_Diet_Prot_category[i]="CD"
	}
	else
		groupings_Diet_Prot_category[i]="HFD"
}

inputdat = rbind(AllData['Prot_14647_Cyp51',],AllData['Prot_100223_Dhcr24',],AllData['Prot_37671_Dhcr7',],AllData['Prot_5447_Ebp',],AllData['Prot_102677_Fdft1',],AllData['Prot_10379_Fdps',],AllData['Prot_10138_Hmgcs1',],AllData['Prot_14384_Hsd17b7',],AllData['Prot_10089_Idi1',],AllData['Prot_10962_Lss',],AllData['Prot_112250_Mvk',],AllData['Prot_13830_Nsdhl',],AllData['Prot_8077_Pmvk',],AllData['Prot_7063_Sc4mol',],AllData['Prot_12717_Sqle',])
inputdat2 <- inputdat[,colSums(is.na(inputdat))<1] # only take if NO NAs
fit=prcomp(t(inputdat2), scale=FALSE)
summary(fit)
ggbiplot(fit, groups = groupings_Diet_Prot_category, ellipse = TRUE, circle = TRUE, scale=1, var.axes=FALSE, choices=c(1,2)) # choices is for which PCs


#$$$$$$ #$$$$$$ #$$$$$$ #$$$$$$ #$$$$$$ #$$$$$$ #$$$$$$ #$$$$$$
#$$$$$$ #$$$$$$ #$$$$$$ #$$$$$$ #$$$$$$ #$$$$$$ #$$$$$$ #$$$$$$
# PANEL 4I
library(gplots)

# TOP PANEL (OXPHOS)
pep1=AllData_young

GeneSet=read.csv(file="~/Dropbox/Aging\ BXD\ Study/1\ _\ Manuscript\ -\ Evan\ Multi-Omic/b_HelpNetworks_ALL_NETWORKS_CANON_mRNA_Prot_ALIGNED.csv", header=T, stringsAsFactors = FALSE)
target.cats <- c("PROT_KEGG_OXPHOS_NoATPase")
GeneSet <- GeneSet[which(GeneSet$Description %in% target.cats),] 
GeneSet1<-GeneSet
tmp = rownames(pep1)
pep1 = pep1[tmp,]
GeneSet1 = GeneSet1[GeneSet1[,3] != "",]	# The first column in the data file must match the third column in the "gene list" file!
rnams = as.character(rownames(pep1))
xx = getMatchingRows(GeneSet1[,3], rnams) # Get the row number of matching rows
GeneMatch = pep1[xx,]
GeneMatch = GeneMatch[order(rownames(GeneMatch)),] 
transform_GeneMatch=apply(t(GeneMatch), c(1,2), as.numeric) # If NAs are coerced in, then it will force everything into a character matrix
tmpr <- corAndPvalue(transform_GeneMatch,method="spearman")
tmp = tmpr$cor
diag(tmp) = 0
outputmatrix_PROT_Oxphos_CORRS_YOUNG=tmp[lower.tri(tmp)] 	

pep1=AllData_old
GeneSet=read.csv(file="~/Dropbox/Aging\ BXD\ Study/1\ _\ Manuscript\ -\ Evan\ Multi-Omic/b_HelpNetworks_ALL_NETWORKS_CANON_mRNA_Prot_ALIGNED.csv", header=T, stringsAsFactors = FALSE)
target.cats <- c("PROT_KEGG_OXPHOS_NoATPase")
GeneSet <- GeneSet[which(GeneSet$Description %in% target.cats),] 
GeneSet1<-GeneSet
tmp = rownames(pep1)
pep1 = pep1[tmp,]
GeneSet1 = GeneSet1[GeneSet1[,3] != "",]	# The first column in the data file must match the third column in the "gene list" file!
rnams = as.character(rownames(pep1))
xx = getMatchingRows(GeneSet1[,3], rnams) # Get the row number of matching rows
GeneMatch = pep1[xx,]
GeneMatch = GeneMatch[order(rownames(GeneMatch)),] 
transform_GeneMatch=apply(t(GeneMatch), c(1,2), as.numeric) # If NAs are coerced in, then it will force everything into a character matrix
tmpr <- corAndPvalue(transform_GeneMatch,method="spearman")
tmp = tmpr$cor
diag(tmp) = 0
outputmatrix_PROT_Oxphos_CORRS_OLD=tmp[lower.tri(tmp)] 	

pep1=AllData
GeneSet=read.csv(file="~/Dropbox/Aging\ BXD\ Study/1\ _\ Manuscript\ -\ Evan\ Multi-Omic/b_HelpNetworks_ALL_NETWORKS_CANON_mRNA_Prot_ALIGNED.csv", header=T, stringsAsFactors = FALSE)
target.cats <- c("PROT_KEGG_OXPHOS_NoATPase")
GeneSet <- GeneSet[which(GeneSet$Description %in% target.cats),] 
GeneSet1<-GeneSet
tmp = rownames(pep1)
pep1 = pep1[tmp,]
GeneSet1 = GeneSet1[GeneSet1[,3] != "",]	# The first column in the data file must match the third column in the "gene list" file!
rnams = as.character(rownames(pep1))
xx = getMatchingRows(GeneSet1[,3], rnams) # Get the row number of matching rows
GeneMatch = pep1[xx,]
GeneMatch = GeneMatch[order(rownames(GeneMatch)),] 
transform_GeneMatch=apply(t(GeneMatch), c(1,2), as.numeric) # If NAs are coerced in, then it will force everything into a character matrix
tmpr <- corAndPvalue(transform_GeneMatch,method="spearman")
tmp = tmpr$cor
diag(tmp) = 0
outputmatrix_PROT_Oxphos_CORRS_ALLAGES=tmp[lower.tri(tmp)] 

plot(density(outputmatrix_PROT_Oxphos_CORRS_ALLAGES, na.rm=T), xlim=c(-1,1), ylim=c(0, 6), col="grey")
plotCI(density(outputmatrix_PROT_Oxphos_CORRS_YOUNG), col="pink", add=T, type="l")
plotCI(density(outputmatrix_PROT_Oxphos_CORRS_OLD), col="red", add=T, type="l")

# BOTTOM PANEL (CHOLESTEROL)
pep1=AllData_CD

GeneSet=read.csv(file="~/Dropbox/Aging\ BXD\ Study/1\ _\ Manuscript\ -\ Evan\ Multi-Omic/b_HelpNetworks_ALL_NETWORKS_CANON_mRNA_Prot_ALIGNED.csv", header=T, stringsAsFactors = FALSE)
target.cats <- c("PROT_REACTOME_CHOLESTEROL_BIOSYNTHESIS")
GeneSet <- GeneSet[which(GeneSet$Description %in% target.cats),] 
GeneSet1<-GeneSet
tmp = rownames(pep1)
pep1 = pep1[tmp,]
GeneSet1 = GeneSet1[GeneSet1[,3] != "",]	# The first column in the data file must match the third column in the "gene list" file!
rnams = as.character(rownames(pep1))
xx = getMatchingRows(GeneSet1[,3], rnams) # Get the row number of matching rows
GeneMatch = pep1[xx,]
GeneMatch = GeneMatch[order(rownames(GeneMatch)),] 
transform_GeneMatch=apply(t(GeneMatch), c(1,2), as.numeric) # If NAs are coerced in, then it will force everything into a character matrix
tmpr <- corAndPvalue(transform_GeneMatch,method="spearman")
tmp = tmpr$cor
diag(tmp) = 0
outputmatrix_PROT_Cholesterol_CORRS_CD=tmp[lower.tri(tmp)] 	

pep1=AllData_HF
GeneSet=read.csv(file="~/Dropbox/Aging\ BXD\ Study/1\ _\ Manuscript\ -\ Evan\ Multi-Omic/b_HelpNetworks_ALL_NETWORKS_CANON_mRNA_Prot_ALIGNED.csv", header=T, stringsAsFactors = FALSE)
target.cats <- c("PROT_REACTOME_CHOLESTEROL_BIOSYNTHESIS")
GeneSet <- GeneSet[which(GeneSet$Description %in% target.cats),] 
GeneSet1<-GeneSet
tmp = rownames(pep1)
pep1 = pep1[tmp,]
GeneSet1 = GeneSet1[GeneSet1[,3] != "",]	# The first column in the data file must match the third column in the "gene list" file!
rnams = as.character(rownames(pep1))
xx = getMatchingRows(GeneSet1[,3], rnams) # Get the row number of matching rows
GeneMatch = pep1[xx,]
GeneMatch = GeneMatch[order(rownames(GeneMatch)),] 
transform_GeneMatch=apply(t(GeneMatch), c(1,2), as.numeric) # If NAs are coerced in, then it will force everything into a character matrix
tmpr <- corAndPvalue(transform_GeneMatch,method="spearman")
tmp = tmpr$cor
diag(tmp) = 0
outputmatrix_PROT_Cholesterol_CORRS_HFD=tmp[lower.tri(tmp)] 	

pep1=AllData
GeneSet=read.csv(file="~/Dropbox/Aging\ BXD\ Study/1\ _\ Manuscript\ -\ Evan\ Multi-Omic/b_HelpNetworks_ALL_NETWORKS_CANON_mRNA_Prot_ALIGNED.csv", header=T, stringsAsFactors = FALSE)
target.cats <- c("PROT_REACTOME_CHOLESTEROL_BIOSYNTHESIS")
GeneSet <- GeneSet[which(GeneSet$Description %in% target.cats),] 
GeneSet1<-GeneSet
tmp = rownames(pep1)
pep1 = pep1[tmp,]
GeneSet1 = GeneSet1[GeneSet1[,3] != "",]	# The first column in the data file must match the third column in the "gene list" file!
rnams = as.character(rownames(pep1))
xx = getMatchingRows(GeneSet1[,3], rnams) # Get the row number of matching rows
GeneMatch = pep1[xx,]
GeneMatch = GeneMatch[order(rownames(GeneMatch)),] 
transform_GeneMatch=apply(t(GeneMatch), c(1,2), as.numeric) # If NAs are coerced in, then it will force everything into a character matrix
tmpr <- corAndPvalue(transform_GeneMatch,method="spearman")
tmp = tmpr$cor
diag(tmp) = 0
outputmatrix_PROT_Cholesterol_CORRS_ALLDIETS=tmp[lower.tri(tmp)] 

plot(density(outputmatrix_PROT_Cholesterol_CORRS_ALLDIETS, na.rm=T), xlim=c(-1,1), ylim=c(0, 6), col="green")
plotCI(density(outputmatrix_PROT_Cholesterol_CORRS_CD), col="grey", add=T, type="l")
plotCI(density(outputmatrix_PROT_Cholesterol_CORRS_HFD), col="black", add=T, type="l")

#$$$$$$ #$$$$$$ #$$$$$$ #$$$$$$ #$$$$$$ #$$$$$$ #$$$$$$ #$$$$$$
#$$$$$$ #$$$$$$ #$$$$$$ #$$$$$$ #$$$$$$ #$$$$$$ #$$$$$$ #$$$$$$
# PANEL S3A (Part of supplemental to Figure 4)

pep1=AllData
GeneSet=read.csv(file="~/Dropbox/Aging\ BXD\ Study/1\ _\ Manuscript\ -\ Evan\ Multi-Omic/b_HelpNetworks_ALL_NETWORKS_CANON_mRNA_Prot_ALIGNED.csv", header=T, stringsAsFactors = FALSE)

target.cats <- c("mRNA_KEGG_OXPHOS_NoATPase","PROT_KEGG_OXPHOS_NoATPase")
GeneSet <- GeneSet[which(GeneSet$Description %in% target.cats),] 
GeneSet1<-GeneSet

# PURELY for visualization, the same genes as in the original diagrams are dropped.
# For edge counts they are retained. 
GeneSet1 <- GeneSet1[-13,] # Drops COX17 (Protein)
GeneSet1 <- GeneSet1[-22,] # Drops COX7A2L (Protien); this gene also has a huge cis-QTL DNA variant that makes it a bad candidate for correlations in the BXDs
GeneSet1 <- GeneSet1[-22,] # Drops COX7B (Protein) (NB: not a typo, since the drop is sequential and reordered)
GeneSet1 <- GeneSet1[-89,] # Drops Cox6a2 (mRNA)
GeneSet1 <- GeneSet1[-93,] # Drops Cox7a2l (mRNA); this gene also has a huge cis-QTL DNA variant that makes it a bad candidate for correlations in the BXDs

# FIRST, load the "CorrGraph" function at the end of this file (scroll, or look for "CORRGRAPH.FIND")

CorrGraph(pep1, GeneSet1, 0.0001, colors=c("black","red","green", "yellow","lightblue")) # Input data, input annotation, and p-value cutoff for edges


#$$$$$$ #$$$$$$ #$$$$$$ #$$$$$$ #$$$$$$ #$$$$$$ #$$$$$$ #$$$$$$
#$$$$$$ #$$$$$$ #$$$$$$ #$$$$$$ #$$$$$$ #$$$$$$ #$$$$$$ #$$$$$$
# PANEL S3B (Part of supplemental to Figure 4)
# Recall: need to load the AllData at top. 

# ggbiplot is on github:
# library(devtools)
# install_github("vqv/ggbiplot")

library(ggbiplot)

# TOP LEFT PANEL:
# Real mRNA data for OXPHOS
inputdat=rbind(AllData['mRNA_41991_Atp5a1',],AllData['mRNA_28643_Atp5b',],AllData['mRNA_3843_Atp5c1',],AllData['mRNA_28087_Atp5d',],AllData['mRNA_6438_Atp5e',],AllData['mRNA_8587_Atp5f1',],AllData['mRNA_31896_Atp5h',],AllData['mRNA_38977_Atp5j',],AllData['mRNA_14556_Atp5j2',],AllData['mRNA_23777_Atp5l',],AllData['mRNA_39070_Atp5o',],AllData['mRNA_43304_mt-Atp8',],AllData['mRNA_38781_Cox17',],AllData['mRNA_22464_Cox4i1',],AllData['mRNA_23952_Cox5a',],AllData['mRNA_1069_Cox5b',],AllData['mRNA_13817_Cox6a1',],AllData['mRNA_20429_Cox6a2',],AllData['mRNA_18825_Cox6b1',],AllData['mRNA_36843_Cox6c',],AllData['mRNA_18772_Cox7a1',],AllData['mRNA_24254_Cox7a2',],AllData['mRNA_26322_Cox7b',],AllData['mRNA_34707_Cox7c',],AllData['mRNA_37117_Cyc1',],AllData['mRNA_43318_mt-Cytb',],AllData['mRNA_43289_mt-Nd1',],AllData['mRNA_43308_mt-Nd3',],AllData['mRNA_43315_mt-Nd5',],AllData['mRNA_25690_Ndufa1',],AllData['mRNA_1711_Ndufa10',],AllData['mRNA_40872_Ndufa11',],AllData['mRNA_41667_Ndufa2',],AllData['mRNA_17893_Ndufa3',],AllData['mRNA_15416_Ndufa4',],AllData['mRNA_15474_Ndufa5',],AllData['mRNA_37368_Ndufa6',],AllData['mRNA_40209_Ndufa7',],AllData['mRNA_3150_Ndufa8',],AllData['mRNA_16885_Ndufa9',],AllData['mRNA_20218_Ndufab1',],AllData['mRNA_39866_Ndufb10',],AllData['mRNA_15657_Ndufb2',],AllData['mRNA_349_Ndufb3',],AllData['mRNA_38775_Ndufb4',],AllData['mRNA_7498_Ndufb5',],AllData['mRNA_9926_Ndufb6',],AllData['mRNA_21951_Ndufb7',],AllData['mRNA_43101_Ndufb8',],AllData['mRNA_36936_Ndufb9',],AllData['mRNA_19648_Ndufc2',],AllData['mRNA_372_Ndufs1',],AllData['mRNA_2460_Ndufs2',],AllData['mRNA_5112_Ndufs3',],AllData['mRNA_33587_Ndufs4',],AllData['mRNA_10856_Ndufs5',],AllData['mRNA_34647_Ndufs6',],AllData['mRNA_28097_Ndufs7',],AllData['mRNA_42373_Ndufs8',],AllData['mRNA_42385_Ndufv1',],AllData['mRNA_40919_Ndufv2',],AllData['mRNA_40129_Ndufv3',],AllData['mRNA_34664_Sdha',],AllData['mRNA_11325_Sdhb',],AllData['mRNA_2443_Sdhc',],AllData['mRNA_23846_Sdhd',],AllData['mRNA_29537_Uqcr10',],AllData['mRNA_28126_Uqcr11',],AllData['mRNA_34575_Uqcrb',],AllData['mRNA_24662_Uqcrc1',],AllData['mRNA_20197_Uqcrc2',],AllData['mRNA_34201_Uqcrfs1',],AllData['mRNA_10648_Uqcrh',],AllData['mRNA_30021_Uqcrq',])
inputcategory=c("ComplexV","ComplexV","ComplexV","ComplexV","ComplexV","ComplexV","ComplexV","ComplexV","ComplexV","ComplexV","ComplexV","ComplexV","ComplexIV","ComplexIV","ComplexIV","ComplexIV","ComplexIV","ComplexIV","ComplexIV","ComplexIV","ComplexIV","ComplexIV","ComplexIV","ComplexIV","ComplexIV","ComplexIV","ComplexI","ComplexI","ComplexI","ComplexI","ComplexI","ComplexI","ComplexI","ComplexI","ComplexI","ComplexI","ComplexI","ComplexI","ComplexI","ComplexI","ComplexI","ComplexI","ComplexI","ComplexI","ComplexI","ComplexI","ComplexI","ComplexI","ComplexI","ComplexI","ComplexI","ComplexI","ComplexI","ComplexI","ComplexI","ComplexI","ComplexI","ComplexI","ComplexI","ComplexI","ComplexI","ComplexI","ComplexII","ComplexII","ComplexII","ComplexII","ComplexIII","ComplexIII","ComplexIII","ComplexIII","ComplexIII","ComplexIII","ComplexIII","ComplexIII")
inputdat2 <- inputdat[,colSums(is.na(inputdat))<1] # only take if NO NAs
fit=prcomp(inputdat2, scale=FALSE)
summary(fit)
ggbiplot(fit, groups = inputcategory, ellipse = TRUE, circle = TRUE, scale=1, var.axes=FALSE, choices=c(1,2)) # choices is for which PCs
# Note that separation of all complexes except CIV is also clear on using PC2,3 (Change choices to 2,3)

# TOP RIGHT PANEL: 
# Real protein data for OXPHOS
inputdat = rbind(AllData['Prot_13610_Atp5a1',],AllData['Prot_14033_Atp5b',],AllData['Prot_9644_Atp5c1',],AllData['Prot_6141_Atp5d',],AllData['Prot_17338_Atp5e',],AllData['Prot_6042_Atp5f1',],AllData['Prot_10960_Atp5h',],AllData['Prot_3917_Atp5j',],AllData['Prot_1953_Atp5j2',],AllData['Prot_13095_Atp5l',],AllData['Prot_9181_Atp5o',],AllData['Prot_7093_ATP8',],AllData['Prot_46625_Cox17',],AllData['Prot_1029_Cox4i1',],AllData['Prot_37952_Cox5a',],AllData['Prot_12175_Cox5b',],AllData['Prot_22888_Cox6a1',],AllData['Prot_57839_Cox6a2',],AllData['Prot_11344_Cox6b1',],AllData['Prot_1524_Cox6c',],AllData['Prot_19095_Cox7a1',],AllData['Prot_4783_Cox7a2',],AllData['Prot_49184_Cox7b',],AllData['Prot_9842_Cox7c',],AllData['Prot_5101_Cyc1',],AllData['Prot_2542_CYTB',],AllData['Prot_30795_ND1',],AllData['Prot_4384_ND3',],AllData['Prot_55212_ND5',],AllData['Prot_64922_Ndufa1',],AllData['Prot_38990_Ndufa10',],AllData['Prot_7917_Ndufa11',],AllData['Prot_12709_Ndufa2',],AllData['Prot_5697_Ndufa3',],AllData['Prot_7682_Ndufa4',],AllData['Prot_12590_Ndufa5',],AllData['Prot_65801_Ndufa6',],AllData['Prot_46413_Ndufa7',],AllData['Prot_15321_Ndufa8',],AllData['Prot_1461_Ndufa9',],AllData['Prot_37742_Ndufab1',],AllData['Prot_14614_Ndufb10',],AllData['Prot_69050_Ndufb2',],AllData['Prot_25796_Ndufb3',],AllData['Prot_68300_Ndufb4',],AllData['Prot_12279_Ndufb5',],AllData['Prot_26002_Ndufb6',],AllData['Prot_9767_Ndufb7',],AllData['Prot_19049_Ndufb8',],AllData['Prot_4236_Ndufb9',],AllData['Prot_80796_Ndufc2',],AllData['Prot_12857_Ndufs1',],AllData['Prot_60482_Ndufs2',],AllData['Prot_11334_Ndufs3',],AllData['Prot_10026_Ndufs4',],AllData['Prot_6163_Ndufs5',],AllData['Prot_6387_Ndufs6',],AllData['Prot_7803_Ndufs7',],AllData['Prot_3464_Ndufs8',],AllData['Prot_6244_Ndufv1',],AllData['Prot_10114_Ndufv2',],AllData['Prot_59191_Ndufv3',],AllData['Prot_10037_Sdha',],AllData['Prot_8734_Sdhb',],AllData['Prot_5796_Sdhc',],AllData['Prot_5833_Sdhd',],AllData['Prot_41297_Uqcr10',],AllData['Prot_8264_Uqcr11',],AllData['Prot_3263_Uqcrb',],AllData['Prot_9568_Uqcrc1',],AllData['Prot_33848_Uqcrc2',],AllData['Prot_11197_Uqcrfs1',],AllData['Prot_11590_Uqcrh',],AllData['Prot_5926_Uqcrq',])
inputcategory=c("ComplexV","ComplexV","ComplexV","ComplexV","ComplexV","ComplexV","ComplexV","ComplexV","ComplexV","ComplexV","ComplexV","ComplexV","ComplexIV","ComplexIV","ComplexIV","ComplexIV","ComplexIV","ComplexIV","ComplexIV","ComplexIV","ComplexIV","ComplexIV","ComplexIV","ComplexIV","ComplexIV","ComplexIV","ComplexI","ComplexI","ComplexI","ComplexI","ComplexI","ComplexI","ComplexI","ComplexI","ComplexI","ComplexI","ComplexI","ComplexI","ComplexI","ComplexI","ComplexI","ComplexI","ComplexI","ComplexI","ComplexI","ComplexI","ComplexI","ComplexI","ComplexI","ComplexI","ComplexI","ComplexI","ComplexI","ComplexI","ComplexI","ComplexI","ComplexI","ComplexI","ComplexI","ComplexI","ComplexI","ComplexI","ComplexII","ComplexII","ComplexII","ComplexII","ComplexIII","ComplexIII","ComplexIII","ComplexIII","ComplexIII","ComplexIII","ComplexIII","ComplexIII")
inputdat2 <- inputdat[,colSums(is.na(inputdat))<1] # only take if NO NAs
fit=prcomp(inputdat2, scale=FALSE)
summary(fit)
ggbiplot(fit, groups = inputcategory, ellipse = TRUE, circle = TRUE, scale=1, var.axes=FALSE, choices=c(1,2)) # choices is for which PCs
# Note that separation of all complexes except CIV is also clear on using PC2,3 (Change choices to 2,3)


# BOTTOM LEFT PANEL: 
# Real mRNA data for 74 randomly-selected genes
inputdat=rbind(AllData['mRNA_15078_Aak1',],AllData['mRNA_7166_Abcd3',],AllData['mRNA_35370_Abhd6',],AllData['mRNA_27839_Ado',],AllData['mRNA_23997_Adpgk',],AllData['mRNA_13928_Aldh2',],AllData['mRNA_21886_Ap1m1',],AllData['mRNA_15968_Aqp1',],AllData['mRNA_30190_Arf1',],AllData['mRNA_7968_Arhgef2',],AllData['mRNA_9798_Asph',],AllData['mRNA_34243_Bphl',],AllData['mRNA_29685_Cct4',],AllData['mRNA_30446_Cd68',],AllData['mRNA_3681_Chd6',],AllData['mRNA_37372_Cyp2d10',],AllData['mRNA_21876_Cyp4f18',],AllData['mRNA_2519_Dcaf8',],AllData['mRNA_29634_Ddc',],AllData['mRNA_5704_Ddrgk1',],AllData['mRNA_9492_Dhcr24',],AllData['mRNA_38287_Dnaja3',],AllData['mRNA_4672_Dync1i2',],AllData['mRNA_9545_Faah',],AllData['mRNA_22026_Farsa',],AllData['mRNA_14266_Fis1',],AllData['mRNA_1375_Fn1',],AllData['mRNA_310_Gls',],AllData['mRNA_8974_Gng5',],AllData['mRNA_42388_Gstp1',],AllData['mRNA_27781_H2afy2',],AllData['mRNA_32315_Hpcal1',],AllData['mRNA_8487_Hsd3b1',],AllData['mRNA_40338_Hspa1b',],AllData['mRNA_23402_Icam1',],AllData['mRNA_20890_Insr',],AllData['mRNA_42980_Lipa',],AllData['mRNA_27166_Man1a',],AllData['mRNA_2299_Mettl13',],AllData['mRNA_28473_Nap1l1',],AllData['mRNA_9927_Ndufb6',],AllData['mRNA_34647_Ndufs6',],AllData['mRNA_40920_Ndufv2',],AllData['mRNA_5679_Nop56',],AllData['mRNA_2636_Parp1',],AllData['mRNA_29512_Pes1',],AllData['mRNA_29701_Pex13',],AllData['mRNA_40729_Pex6',],AllData['mRNA_37958_Pi4ka',],AllData['mRNA_604_Plekha6',],AllData['mRNA_27778_Ppa1',],AllData['mRNA_15352_Ppfibp1',],AllData['mRNA_37614_Prph',],AllData['mRNA_41924_Psmg2',],AllData['mRNA_1616_Ptma',],AllData['mRNA_7639_Rab33b',],AllData['mRNA_35778_Rnase4',],AllData['mRNA_3782_Rpp38',],AllData['mRNA_27548_Rps12',],AllData['mRNA_39227_Rps6ka2',],AllData['mRNA_19799_Rrm1',],AllData['mRNA_2994_Rsu1',],AllData['mRNA_4301_Sh3glb2',],AllData['mRNA_17149_Slco1a1',],AllData['mRNA_33117_Slirp',],AllData['mRNA_41333_Snx2',],AllData['mRNA_41335_Snx24',],AllData['mRNA_15038_Suclg1',],AllData['mRNA_12382_Taok3',],AllData['mRNA_41756_Tcerg1',],AllData['mRNA_30248_Tmem11',],AllData['mRNA_9206_Tstd3',],AllData['mRNA_37521_Twf1',],AllData['mRNA_38399_Ube2v2',])
inputcategory=c("ComplexV","ComplexV","ComplexV","ComplexV","ComplexV","ComplexV","ComplexV","ComplexV","ComplexV","ComplexV","ComplexV","ComplexV","ComplexIV","ComplexIV","ComplexIV","ComplexIV","ComplexIV","ComplexIV","ComplexIV","ComplexIV","ComplexIV","ComplexIV","ComplexIV","ComplexIV","ComplexIV","ComplexIV","ComplexI","ComplexI","ComplexI","ComplexI","ComplexI","ComplexI","ComplexI","ComplexI","ComplexI","ComplexI","ComplexI","ComplexI","ComplexI","ComplexI","ComplexI","ComplexI","ComplexI","ComplexI","ComplexI","ComplexI","ComplexI","ComplexI","ComplexI","ComplexI","ComplexI","ComplexI","ComplexI","ComplexI","ComplexI","ComplexI","ComplexI","ComplexI","ComplexI","ComplexI","ComplexI","ComplexI","ComplexII","ComplexII","ComplexII","ComplexII","ComplexIII","ComplexIII","ComplexIII","ComplexIII","ComplexIII","ComplexIII","ComplexIII","ComplexIII")
inputdat2 <- inputdat[,colSums(is.na(inputdat))<1] # only take if NO NAs
fit=prcomp(inputdat2, scale=FALSE)
summary(fit)
ggbiplot(fit, groups = inputcategory, ellipse = TRUE, circle = TRUE, scale=1, var.axes=FALSE, choices=c(1,2)) # choices is for which PCs

# Real protein data for 74 randomly-selected genes
inputdat = rbind(AllData['Prot_9215_Pcbp1',],AllData['Prot_2982_Ugp2',],AllData['Prot_8961_Dak',],AllData['Prot_4805_Eftud2',],AllData['Prot_11744_Scarb2',],AllData['Prot_8065_Hsd17b12',],AllData['Prot_5851_Akr1a1',],AllData['Prot_13819_R3hcc1',],AllData['Prot_13411_Hbb-b1',],AllData['Prot_10170_Aldh1a7',],AllData['Prot_6428_Ahcy',],AllData['Prot_8970_Pecr',],AllData['Prot_20414_Slc25a12',],AllData['Prot_6425_Ugt2b5',],AllData['Prot_11939_Bphl',],AllData['Prot_8167_Aldh5a1',],AllData['Prot_639_Gsta3',],AllData['Prot_6426_Hnrnpk',],AllData['Prot_6807_Tpm3',],AllData['Prot_12888_Apoa1',],AllData['Prot_51905_Aqp1',],AllData['Prot_116056_Actg1',],AllData['Prot_17140_Tpp1',],AllData['Prot_3566_Cyp2c40',],AllData['Prot_7851_Vcp',],AllData['Prot_7606_Psmb6',],AllData['Prot_8531_Pdia5',],AllData['Prot_78869_Rad23b',],AllData['Prot_4601_Ddx17',],AllData['Prot_13599_Acsl1',],AllData['Prot_7827_Gstm6',],AllData['Prot_14569_Gpx1',],AllData['Prot_9181_Atp5o',],AllData['Prot_4037_Urad',],AllData['Prot_841_Sds',],AllData['Prot_52047_Lactb2',],AllData['Prot_11265_Hsp90b1',],AllData['Prot_56792_Ap2m1',],AllData['Prot_9155_Cct8',],AllData['Prot_8545_Acsm5',],AllData['Prot_13584_Phb',],AllData['Prot_739_Arpc5',],AllData['Prot_9760_Rps26',],AllData['Prot_8544_Hadh',],AllData['Prot_81575_Apoa1bp',],AllData['Prot_37007_Irgm1',],AllData['Prot_11461_Agmat',],AllData['Prot_723_Suclg1',],AllData['Prot_13416_Gstz1',],AllData['Prot_13349_Pygm',],AllData['Prot_8852_Agxt2',],AllData['Prot_553_Cmbl',],AllData['Prot_4044_Bdh1',],AllData['Prot_7862_Gbe1',],AllData['Prot_13546_Mat2b',],AllData['Prot_3874_Ces1c',],AllData['Prot_12897_Snrpd3',],AllData['Prot_13564_Hspe1',],AllData['Prot_3031_Qdpr',],AllData['Prot_4339_Pabpc1',],AllData['Prot_1991_Cnn3',],AllData['Prot_14740_Ppp2cb',],AllData['Prot_2433_Erp44',],AllData['Prot_3192_Aass',],AllData['Prot_9658_Rpl7a',],AllData['Prot_29020_Acadvl',],AllData['Prot_5349_Sec14l2',],AllData['Prot_11608_Fmo1',],AllData['Prot_2553_Ugt3a1',],AllData['Prot_6723_Tubb4b',],AllData['Prot_64141_Rpl35a',],AllData['Prot_13518_Glo1',],AllData['Prot_25003_Pycrl',],AllData['Prot_18468_Lsm3',],AllData['Prot_10816_Gulo',])
inputcategory=c("ComplexV","ComplexV","ComplexV","ComplexV","ComplexV","ComplexV","ComplexV","ComplexV","ComplexV","ComplexV","ComplexV","ComplexV","ComplexIV","ComplexIV","ComplexIV","ComplexIV","ComplexIV","ComplexIV","ComplexIV","ComplexIV","ComplexIV","ComplexIV","ComplexIV","ComplexIV","ComplexIV","ComplexIV","ComplexI","ComplexI","ComplexI","ComplexI","ComplexI","ComplexI","ComplexI","ComplexI","ComplexI","ComplexI","ComplexI","ComplexI","ComplexI","ComplexI","ComplexI","ComplexI","ComplexI","ComplexI","ComplexI","ComplexI","ComplexI","ComplexI","ComplexI","ComplexI","ComplexI","ComplexI","ComplexI","ComplexI","ComplexI","ComplexI","ComplexI","ComplexI","ComplexI","ComplexI","ComplexI","ComplexI","ComplexII","ComplexII","ComplexII","ComplexII","ComplexIII","ComplexIII","ComplexIII","ComplexIII","ComplexIII","ComplexIII","ComplexIII","ComplexIII","ComplexIII")
inputdat2 <- inputdat[,colSums(is.na(inputdat))<1] # only take if NO NAs
fit=prcomp(inputdat2, scale=FALSE)
summary(fit)
ggbiplot(fit, groups = inputcategory, ellipse = TRUE, circle = TRUE, scale=1, var.axes=FALSE, choices=c(1,2)) # choices is for which PCs


#$$$$$$ #$$$$$$ #$$$$$$ #$$$$$$ #$$$$$$ #$$$$$$ #$$$$$$ #$$$$$$
#$$$$$$ #$$$$$$ #$$$$$$ #$$$$$$ #$$$$$$ #$$$$$$ #$$$$$$ #$$$$$$
# PANEL S3C (Part of supplemental to Figure 4); note that this is a VERY large figure in terms of code

# TOP-LEFT (Ribosome)
pep1=AllData

GeneSet=read.csv(file="~/Dropbox/Aging\ BXD\ Study/1\ _\ Manuscript\ -\ Evan\ Multi-Omic/b_HelpNetworks_ALL_NETWORKS_CANON_mRNA_Prot_ALIGNED.csv", header=T, stringsAsFactors = FALSE)
target.cats <- c("mRNA_KEGG_RIBOSOME")
GeneSet <- GeneSet[which(GeneSet$Description %in% target.cats),] 
GeneSet1<-GeneSet
tmp = rownames(pep1)
pep1 = pep1[tmp,]
GeneSet1 = GeneSet1[GeneSet1[,3] != "",]	# The first column in the data file must match the third column in the "gene list" file!
rnams = as.character(rownames(pep1))
xx = getMatchingRows(GeneSet1[,3], rnams) # Get the row number of matching rows
GeneMatch = pep1[xx,]
GeneMatch = GeneMatch[order(rownames(GeneMatch)),] 
transform_GeneMatch=apply(t(GeneMatch), c(1,2), as.numeric) # If NAs are coerced in, then it will force everything into a character matrix
tmpr <- corAndPvalue(transform_GeneMatch,method="spearman")
tmp = tmpr$cor
diag(tmp) = 0
outputmatrix_mRNA_Ribosome_CORRS=tmp[lower.tri(tmp)] 	

GeneSet=read.csv(file="~/Dropbox/Aging\ BXD\ Study/1\ _\ Manuscript\ -\ Evan\ Multi-Omic/b_HelpNetworks_ALL_NETWORKS_CANON_mRNA_Prot_ALIGNED.csv", header=T, stringsAsFactors = FALSE)
target.cats <- c("PROT_KEGG_RIBOSOME")
GeneSet <- GeneSet[which(GeneSet$Description %in% target.cats),] 
GeneSet1<-GeneSet
tmp = rownames(pep1)
pep1 = pep1[tmp,]
GeneSet1 = GeneSet1[GeneSet1[,3] != "",]	# The first column in the data file must match the third column in the "gene list" file!
rnams = as.character(rownames(pep1))
xx = getMatchingRows(GeneSet1[,3], rnams) # Get the row number of matching rows
GeneMatch = pep1[xx,]
GeneMatch = GeneMatch[order(rownames(GeneMatch)),] 
transform_GeneMatch=apply(t(GeneMatch), c(1,2), as.numeric) # If NAs are coerced in, then it will force everything into a character matrix
tmpr <- corAndPvalue(transform_GeneMatch,method="spearman")
tmp = tmpr$cor
diag(tmp) = 0
outputmatrix_PROT_Ribosome_CORRS=tmp[lower.tri(tmp)] 	

GeneSet=read.csv(file="~/Dropbox/Aging\ BXD\ Study/1\ _\ Manuscript\ -\ Evan\ Multi-Omic/b_HelpNetworks_ALL_NETWORKS_CANON_mRNA_Prot_ALIGNED.csv", header=T, stringsAsFactors = FALSE)
target.cats <- c("mRNA_KEGG_RIBOSOME","PROT_KEGG_RIBOSOME")
GeneSet <- GeneSet[which(GeneSet$Description %in% target.cats),] 
GeneSet1<-GeneSet
tmp = rownames(pep1)
pep1 = pep1[tmp,]
GeneSet1 = GeneSet1[GeneSet1[,3] != "",]	# The first column in the data file must match the third column in the "gene list" file!
rnams = as.character(rownames(pep1))
xx = getMatchingRows(GeneSet1[,3], rnams) # Get the row number of matching rows
GeneMatch = pep1[xx,]
GeneMatch = GeneMatch[order(rownames(GeneMatch)),] 
transform_GeneMatch=apply(t(GeneMatch), c(1,2), as.numeric) # If NAs are coerced in, then it will force everything into a character matrix
tmpr <- corAndPvalue(transform_GeneMatch,method="spearman")
tmp = tmpr$cor
diag(tmp) = 0
outputmatrix_ALL_Ribosome_CORRS=tmp[lower.tri(tmp)] 
outputmatrix_ALL_Ribosome_CORRS2=outputmatrix_ALL_Ribosome_CORRS[!(outputmatrix_ALL_Ribosome_CORRS %in% outputmatrix_mRNA_Ribosome_CORRS)]
outputmatrix_ALL_Ribosome_CORRS3=outputmatrix_ALL_Ribosome_CORRS2[!(outputmatrix_ALL_Ribosome_CORRS2 %in% outputmatrix_PROT_Ribosome_CORRS)]

### NOTE: This "cross-layer" code has to be calculated last, and it suppresses the correlations that were found within protein and within mRNA, only retaining "across"

# Now we make the density plot
library(gplots)

plot(density(outputmatrix_ALL_Ribosome_CORRS3, na.rm=T), xlim=c(-1,1), ylim=c(0, 6), col="black")
plotCI(density(outputmatrix_mRNA_Ribosome_CORRS), col="blue", add=T, type="l")
plotCI(density(outputmatrix_PROT_Ribosome_CORRS), col="red", add=T, type="l")


# TOP-CENTER-LEFT (Mito Ribosome)
pep1=AllData

GeneSet=read.csv(file="~/Dropbox/Aging\ BXD\ Study/1\ _\ Manuscript\ -\ Evan\ Multi-Omic/b_HelpNetworks_ALL_NETWORKS_CANON_mRNA_Prot_ALIGNED.csv", header=T, stringsAsFactors = FALSE)
target.cats <- c("mRNA_REACTOME_MITOCHONDRIAL_TRANSLATION")
GeneSet <- GeneSet[which(GeneSet$Description %in% target.cats),] 
GeneSet1<-GeneSet
tmp = rownames(pep1)
pep1 = pep1[tmp,]
GeneSet1 = GeneSet1[GeneSet1[,3] != "",]	# The first column in the data file must match the third column in the "gene list" file!
rnams = as.character(rownames(pep1))
xx = getMatchingRows(GeneSet1[,3], rnams) # Get the row number of matching rows
GeneMatch = pep1[xx,]
GeneMatch = GeneMatch[order(rownames(GeneMatch)),] 
transform_GeneMatch=apply(t(GeneMatch), c(1,2), as.numeric) # If NAs are coerced in, then it will force everything into a character matrix
tmpr <- corAndPvalue(transform_GeneMatch,method="spearman")
tmp = tmpr$cor
diag(tmp) = 0
outputmatrix_mRNA_MitoRibosome_CORRS=tmp[lower.tri(tmp)] 	

GeneSet=read.csv(file="~/Dropbox/Aging\ BXD\ Study/1\ _\ Manuscript\ -\ Evan\ Multi-Omic/b_HelpNetworks_ALL_NETWORKS_CANON_mRNA_Prot_ALIGNED.csv", header=T, stringsAsFactors = FALSE)
target.cats <- c("PROT_REACTOME_MITOCHONDRIAL_TRANSLATION")
GeneSet <- GeneSet[which(GeneSet$Description %in% target.cats),] 
GeneSet1<-GeneSet
tmp = rownames(pep1)
pep1 = pep1[tmp,]
GeneSet1 = GeneSet1[GeneSet1[,3] != "",]	# The first column in the data file must match the third column in the "gene list" file!
rnams = as.character(rownames(pep1))
xx = getMatchingRows(GeneSet1[,3], rnams) # Get the row number of matching rows
GeneMatch = pep1[xx,]
GeneMatch = GeneMatch[order(rownames(GeneMatch)),] 
transform_GeneMatch=apply(t(GeneMatch), c(1,2), as.numeric) # If NAs are coerced in, then it will force everything into a character matrix
tmpr <- corAndPvalue(transform_GeneMatch,method="spearman")
tmp = tmpr$cor
diag(tmp) = 0
outputmatrix_PROT_MitoRibosome_CORRS=tmp[lower.tri(tmp)] 	

GeneSet=read.csv(file="~/Dropbox/Aging\ BXD\ Study/1\ _\ Manuscript\ -\ Evan\ Multi-Omic/b_HelpNetworks_ALL_NETWORKS_CANON_mRNA_Prot_ALIGNED.csv", header=T, stringsAsFactors = FALSE)
target.cats <- c("mRNA_REACTOME_MITOCHONDRIAL_TRANSLATION","PROT_REACTOME_MITOCHONDRIAL_TRANSLATION")
GeneSet <- GeneSet[which(GeneSet$Description %in% target.cats),] 
GeneSet1<-GeneSet
tmp = rownames(pep1)
pep1 = pep1[tmp,]
GeneSet1 = GeneSet1[GeneSet1[,3] != "",]	# The first column in the data file must match the third column in the "gene list" file!
rnams = as.character(rownames(pep1))
xx = getMatchingRows(GeneSet1[,3], rnams) # Get the row number of matching rows
GeneMatch = pep1[xx,]
GeneMatch = GeneMatch[order(rownames(GeneMatch)),] 
transform_GeneMatch=apply(t(GeneMatch), c(1,2), as.numeric) # If NAs are coerced in, then it will force everything into a character matrix
tmpr <- corAndPvalue(transform_GeneMatch,method="spearman")
tmp = tmpr$cor
diag(tmp) = 0
outputmatrix_ALL_MitoRibosome_CORRS=tmp[lower.tri(tmp)] 
outputmatrix_ALL_MitoRibosome_CORRS2=outputmatrix_ALL_MitoRibosome_CORRS[!(outputmatrix_ALL_MitoRibosome_CORRS %in% outputmatrix_mRNA_MitoRibosome_CORRS)]
outputmatrix_ALL_MitoRibosome_CORRS3=outputmatrix_ALL_MitoRibosome_CORRS2[!(outputmatrix_ALL_MitoRibosome_CORRS2 %in% outputmatrix_PROT_MitoRibosome_CORRS)]

### NOTE: This "cross-layer" code has to be calculated last, and it suppresses the correlations that were found within protein and within mRNA, only retaining "across"

# Now we make the density plot
library(gplots)

plot(density(outputmatrix_ALL_MitoRibosome_CORRS3, na.rm=T), xlim=c(-1,1), ylim=c(0, 6), col="black")
plotCI(density(outputmatrix_mRNA_MitoRibosome_CORRS), col="blue", add=T, type="l")
plotCI(density(outputmatrix_PROT_MitoRibosome_CORRS), col="red", add=T, type="l")

# TOP-CENTER-RIGHT (Proteasome)
pep1=AllData

GeneSet=read.csv(file="~/Dropbox/Aging\ BXD\ Study/1\ _\ Manuscript\ -\ Evan\ Multi-Omic/b_HelpNetworks_ALL_NETWORKS_CANON_mRNA_Prot_ALIGNED.csv", header=T, stringsAsFactors = FALSE)
target.cats <- c("mRNA_KEGG_PROTEASOME")
GeneSet <- GeneSet[which(GeneSet$Description %in% target.cats),] 
GeneSet1<-GeneSet
tmp = rownames(pep1)
pep1 = pep1[tmp,]
GeneSet1 = GeneSet1[GeneSet1[,3] != "",]	# The first column in the data file must match the third column in the "gene list" file!
rnams = as.character(rownames(pep1))
xx = getMatchingRows(GeneSet1[,3], rnams) # Get the row number of matching rows
GeneMatch = pep1[xx,]
GeneMatch = GeneMatch[order(rownames(GeneMatch)),] 
transform_GeneMatch=apply(t(GeneMatch), c(1,2), as.numeric) # If NAs are coerced in, then it will force everything into a character matrix
tmpr <- corAndPvalue(transform_GeneMatch,method="spearman")
tmp = tmpr$cor
diag(tmp) = 0
outputmatrix_mRNA_Proteasome_CORRS=tmp[lower.tri(tmp)] 	

GeneSet=read.csv(file="~/Dropbox/Aging\ BXD\ Study/1\ _\ Manuscript\ -\ Evan\ Multi-Omic/b_HelpNetworks_ALL_NETWORKS_CANON_mRNA_Prot_ALIGNED.csv", header=T, stringsAsFactors = FALSE)
target.cats <- c("PROT_KEGG_PROTEASOME")
GeneSet <- GeneSet[which(GeneSet$Description %in% target.cats),] 
GeneSet1<-GeneSet
tmp = rownames(pep1)
pep1 = pep1[tmp,]
GeneSet1 = GeneSet1[GeneSet1[,3] != "",]	# The first column in the data file must match the third column in the "gene list" file!
rnams = as.character(rownames(pep1))
xx = getMatchingRows(GeneSet1[,3], rnams) # Get the row number of matching rows
GeneMatch = pep1[xx,]
GeneMatch = GeneMatch[order(rownames(GeneMatch)),] 
transform_GeneMatch=apply(t(GeneMatch), c(1,2), as.numeric) # If NAs are coerced in, then it will force everything into a character matrix
tmpr <- corAndPvalue(transform_GeneMatch,method="spearman")
tmp = tmpr$cor
diag(tmp) = 0
outputmatrix_PROT_Proteasome_CORRS=tmp[lower.tri(tmp)] 	

GeneSet=read.csv(file="~/Dropbox/Aging\ BXD\ Study/1\ _\ Manuscript\ -\ Evan\ Multi-Omic/b_HelpNetworks_ALL_NETWORKS_CANON_mRNA_Prot_ALIGNED.csv", header=T, stringsAsFactors = FALSE)
target.cats <- c("mRNA_KEGG_PROTEASOME","PROT_KEGG_PROTEASOME")
GeneSet <- GeneSet[which(GeneSet$Description %in% target.cats),] 
GeneSet1<-GeneSet
tmp = rownames(pep1)
pep1 = pep1[tmp,]
GeneSet1 = GeneSet1[GeneSet1[,3] != "",]	# The first column in the data file must match the third column in the "gene list" file!
rnams = as.character(rownames(pep1))
xx = getMatchingRows(GeneSet1[,3], rnams) # Get the row number of matching rows
GeneMatch = pep1[xx,]
GeneMatch = GeneMatch[order(rownames(GeneMatch)),] 
transform_GeneMatch=apply(t(GeneMatch), c(1,2), as.numeric) # If NAs are coerced in, then it will force everything into a character matrix
tmpr <- corAndPvalue(transform_GeneMatch,method="spearman")
tmp = tmpr$cor
diag(tmp) = 0
outputmatrix_ALL_Proteasome_CORRS=tmp[lower.tri(tmp)] 
outputmatrix_ALL_Proteasome_CORRS2=outputmatrix_ALL_Proteasome_CORRS[!(outputmatrix_ALL_Proteasome_CORRS %in% outputmatrix_mRNA_Proteasome_CORRS)]
outputmatrix_ALL_Proteasome_CORRS3=outputmatrix_ALL_Proteasome_CORRS2[!(outputmatrix_ALL_Proteasome_CORRS2 %in% outputmatrix_PROT_Proteasome_CORRS)]

### NOTE: This "cross-layer" code has to be calculated last, and it suppresses the correlations that were found within protein and within mRNA, only retaining "across"

# Now we make the density plot
library(gplots)

plot(density(outputmatrix_ALL_Proteasome_CORRS3, na.rm=T), xlim=c(-1,1), ylim=c(0, 6), col="black")
plotCI(density(outputmatrix_mRNA_Proteasome_CORRS), col="blue", add=T, type="l")
plotCI(density(outputmatrix_PROT_Proteasome_CORRS), col="red", add=T, type="l")

# TOP-RIGHT (Random)

## This is the same as Figure 4B (left panel), code not repeated here; look above.

# BOTTOM-LEFT
pep1=AllData

GeneSet=read.csv(file="~/Dropbox/Aging\ BXD\ Study/1\ _\ Manuscript\ -\ Evan\ Multi-Omic/b_HelpNetworks_ALL_NETWORKS_CANON_mRNA_Prot_ALIGNED.csv", header=T, stringsAsFactors = FALSE)
target.cats <- c("mRNA_REACTOME_MITOCHONDRIAL_FATTY_ACID_BETA_OXIDATION")
GeneSet <- GeneSet[which(GeneSet$Description %in% target.cats),] 
GeneSet1<-GeneSet
tmp = rownames(pep1)
pep1 = pep1[tmp,]
GeneSet1 = GeneSet1[GeneSet1[,3] != "",]	# The first column in the data file must match the third column in the "gene list" file!
rnams = as.character(rownames(pep1))
xx = getMatchingRows(GeneSet1[,3], rnams) # Get the row number of matching rows
GeneMatch = pep1[xx,]
GeneMatch = GeneMatch[order(rownames(GeneMatch)),] 
transform_GeneMatch=apply(t(GeneMatch), c(1,2), as.numeric) # If NAs are coerced in, then it will force everything into a character matrix
tmpr <- corAndPvalue(transform_GeneMatch,method="spearman")
tmp = tmpr$cor
diag(tmp) = 0
outputmatrix_mRNA_BetaOxidation_CORRS=tmp[lower.tri(tmp)] 	

GeneSet=read.csv(file="~/Dropbox/Aging\ BXD\ Study/1\ _\ Manuscript\ -\ Evan\ Multi-Omic/b_HelpNetworks_ALL_NETWORKS_CANON_mRNA_Prot_ALIGNED.csv", header=T, stringsAsFactors = FALSE)
target.cats <- c("PROT_REACTOME_MITOCHONDRIAL_FATTY_ACID_BETA_OXIDATION")
GeneSet <- GeneSet[which(GeneSet$Description %in% target.cats),] 
GeneSet1<-GeneSet
tmp = rownames(pep1)
pep1 = pep1[tmp,]
GeneSet1 = GeneSet1[GeneSet1[,3] != "",]	# The first column in the data file must match the third column in the "gene list" file!
rnams = as.character(rownames(pep1))
xx = getMatchingRows(GeneSet1[,3], rnams) # Get the row number of matching rows
GeneMatch = pep1[xx,]
GeneMatch = GeneMatch[order(rownames(GeneMatch)),] 
transform_GeneMatch=apply(t(GeneMatch), c(1,2), as.numeric) # If NAs are coerced in, then it will force everything into a character matrix
tmpr <- corAndPvalue(transform_GeneMatch,method="spearman")
tmp = tmpr$cor
diag(tmp) = 0
outputmatrix_PROT_BetaOxidation_CORRS=tmp[lower.tri(tmp)] 	

GeneSet=read.csv(file="~/Dropbox/Aging\ BXD\ Study/1\ _\ Manuscript\ -\ Evan\ Multi-Omic/b_HelpNetworks_ALL_NETWORKS_CANON_mRNA_Prot_ALIGNED.csv", header=T, stringsAsFactors = FALSE)
target.cats <- c("mRNA_REACTOME_MITOCHONDRIAL_FATTY_ACID_BETA_OXIDATION","PROT_REACTOME_MITOCHONDRIAL_FATTY_ACID_BETA_OXIDATION")
GeneSet <- GeneSet[which(GeneSet$Description %in% target.cats),] 
GeneSet1<-GeneSet
tmp = rownames(pep1)
pep1 = pep1[tmp,]
GeneSet1 = GeneSet1[GeneSet1[,3] != "",]	# The first column in the data file must match the third column in the "gene list" file!
rnams = as.character(rownames(pep1))
xx = getMatchingRows(GeneSet1[,3], rnams) # Get the row number of matching rows
GeneMatch = pep1[xx,]
GeneMatch = GeneMatch[order(rownames(GeneMatch)),] 
transform_GeneMatch=apply(t(GeneMatch), c(1,2), as.numeric) # If NAs are coerced in, then it will force everything into a character matrix
tmpr <- corAndPvalue(transform_GeneMatch,method="spearman")
tmp = tmpr$cor
diag(tmp) = 0
outputmatrix_ALL_BetaOxidation_CORRS=tmp[lower.tri(tmp)] 
outputmatrix_ALL_BetaOxidation_CORRS2=outputmatrix_ALL_BetaOxidation_CORRS[!(outputmatrix_ALL_BetaOxidation_CORRS %in% outputmatrix_mRNA_BetaOxidation_CORRS)]
outputmatrix_ALL_BetaOxidation_CORRS3=outputmatrix_ALL_BetaOxidation_CORRS2[!(outputmatrix_ALL_BetaOxidation_CORRS2 %in% outputmatrix_PROT_BetaOxidation_CORRS)]

### NOTE: This "cross-layer" code has to be calculated last, and it suppresses the correlations that were found within protein and within mRNA, only retaining "across"

# Now we make the density plot
library(gplots)

plot(density(outputmatrix_ALL_BetaOxidation_CORRS3, na.rm=T), xlim=c(-1,1), ylim=c(0, 6), col="black")
plotCI(density(outputmatrix_mRNA_BetaOxidation_CORRS), col="blue", add=T, type="l")
plotCI(density(outputmatrix_PROT_BetaOxidation_CORRS), col="red", add=T, type="l")

# BOTTOM-CENTER-LEFT
pep1=AllData

GeneSet=read.csv(file="~/Dropbox/Aging\ BXD\ Study/1\ _\ Manuscript\ -\ Evan\ Multi-Omic/b_HelpNetworks_ALL_NETWORKS_CANON_mRNA_Prot_ALIGNED.csv", header=T, stringsAsFactors = FALSE)
target.cats <- c("mRNA_REACTOME_CITRIC_ACID_CYCLE_TCA_CYCLE")
GeneSet <- GeneSet[which(GeneSet$Description %in% target.cats),] 
GeneSet1<-GeneSet
tmp = rownames(pep1)
pep1 = pep1[tmp,]
GeneSet1 = GeneSet1[GeneSet1[,3] != "",]	# The first column in the data file must match the third column in the "gene list" file!
rnams = as.character(rownames(pep1))
xx = getMatchingRows(GeneSet1[,3], rnams) # Get the row number of matching rows
GeneMatch = pep1[xx,]
GeneMatch = GeneMatch[order(rownames(GeneMatch)),] 
transform_GeneMatch=apply(t(GeneMatch), c(1,2), as.numeric) # If NAs are coerced in, then it will force everything into a character matrix
tmpr <- corAndPvalue(transform_GeneMatch,method="spearman")
tmp = tmpr$cor
diag(tmp) = 0
outputmatrix_mRNA_TCACycle_CORRS=tmp[lower.tri(tmp)] 	

GeneSet=read.csv(file="~/Dropbox/Aging\ BXD\ Study/1\ _\ Manuscript\ -\ Evan\ Multi-Omic/b_HelpNetworks_ALL_NETWORKS_CANON_mRNA_Prot_ALIGNED.csv", header=T, stringsAsFactors = FALSE)
target.cats <- c("PROT_REACTOME_CITRIC_ACID_CYCLE_TCA_CYCLE")
GeneSet <- GeneSet[which(GeneSet$Description %in% target.cats),] 
GeneSet1<-GeneSet
tmp = rownames(pep1)
pep1 = pep1[tmp,]
GeneSet1 = GeneSet1[GeneSet1[,3] != "",]	# The first column in the data file must match the third column in the "gene list" file!
rnams = as.character(rownames(pep1))
xx = getMatchingRows(GeneSet1[,3], rnams) # Get the row number of matching rows
GeneMatch = pep1[xx,]
GeneMatch = GeneMatch[order(rownames(GeneMatch)),] 
transform_GeneMatch=apply(t(GeneMatch), c(1,2), as.numeric) # If NAs are coerced in, then it will force everything into a character matrix
tmpr <- corAndPvalue(transform_GeneMatch,method="spearman")
tmp = tmpr$cor
diag(tmp) = 0
outputmatrix_PROT_TCACycle_CORRS=tmp[lower.tri(tmp)] 	

GeneSet=read.csv(file="~/Dropbox/Aging\ BXD\ Study/1\ _\ Manuscript\ -\ Evan\ Multi-Omic/b_HelpNetworks_ALL_NETWORKS_CANON_mRNA_Prot_ALIGNED.csv", header=T, stringsAsFactors = FALSE)
target.cats <- c("mRNA_REACTOME_CITRIC_ACID_CYCLE_TCA_CYCLE","PROT_REACTOME_CITRIC_ACID_CYCLE_TCA_CYCLE")
GeneSet <- GeneSet[which(GeneSet$Description %in% target.cats),] 
GeneSet1<-GeneSet
tmp = rownames(pep1)
pep1 = pep1[tmp,]
GeneSet1 = GeneSet1[GeneSet1[,3] != "",]	# The first column in the data file must match the third column in the "gene list" file!
rnams = as.character(rownames(pep1))
xx = getMatchingRows(GeneSet1[,3], rnams) # Get the row number of matching rows
GeneMatch = pep1[xx,]
GeneMatch = GeneMatch[order(rownames(GeneMatch)),] 
transform_GeneMatch=apply(t(GeneMatch), c(1,2), as.numeric) # If NAs are coerced in, then it will force everything into a character matrix
tmpr <- corAndPvalue(transform_GeneMatch,method="spearman")
tmp = tmpr$cor
diag(tmp) = 0
outputmatrix_ALL_TCACycle_CORRS=tmp[lower.tri(tmp)] 
outputmatrix_ALL_TCACycle_CORRS2=outputmatrix_ALL_TCACycle_CORRS[!(outputmatrix_ALL_TCACycle_CORRS %in% outputmatrix_mRNA_TCACycle_CORRS)]
outputmatrix_ALL_TCACycle_CORRS3=outputmatrix_ALL_TCACycle_CORRS2[!(outputmatrix_ALL_TCACycle_CORRS2 %in% outputmatrix_PROT_TCACycle_CORRS)]

### NOTE: This "cross-layer" code has to be calculated last, and it suppresses the correlations that were found within protein and within mRNA, only retaining "across"

# Now we make the density plot
library(gplots)

plot(density(outputmatrix_ALL_TCACycle_CORRS3, na.rm=T), xlim=c(-1,1), ylim=c(0, 6), col="black")
plotCI(density(outputmatrix_mRNA_TCACycle_CORRS), col="blue", add=T, type="l")
plotCI(density(outputmatrix_PROT_TCACycle_CORRS), col="red", add=T, type="l")

# BOTTOM-CENTER-RIGHT
pep1=AllData

GeneSet=read.csv(file="~/Dropbox/Aging\ BXD\ Study/1\ _\ Manuscript\ -\ Evan\ Multi-Omic/b_HelpNetworks_ALL_NETWORKS_CANON_mRNA_Prot_ALIGNED.csv", header=T, stringsAsFactors = FALSE)
target.cats <- c("mRNA_KEGG_METABOLISM_OF_XENOBIOTICS_BY_CYTOCHROME_P450")
GeneSet <- GeneSet[which(GeneSet$Description %in% target.cats),] 
GeneSet1<-GeneSet
tmp = rownames(pep1)
pep1 = pep1[tmp,]
GeneSet1 = GeneSet1[GeneSet1[,3] != "",]	# The first column in the data file must match the third column in the "gene list" file!
rnams = as.character(rownames(pep1))
xx = getMatchingRows(GeneSet1[,3], rnams) # Get the row number of matching rows
GeneMatch = pep1[xx,]
GeneMatch = GeneMatch[order(rownames(GeneMatch)),] 
transform_GeneMatch=apply(t(GeneMatch), c(1,2), as.numeric) # If NAs are coerced in, then it will force everything into a character matrix
tmpr <- corAndPvalue(transform_GeneMatch,method="spearman")
tmp = tmpr$cor
diag(tmp) = 0
outputmatrix_mRNA_CYP450_CORRS=tmp[lower.tri(tmp)] 	

GeneSet=read.csv(file="~/Dropbox/Aging\ BXD\ Study/1\ _\ Manuscript\ -\ Evan\ Multi-Omic/b_HelpNetworks_ALL_NETWORKS_CANON_mRNA_Prot_ALIGNED.csv", header=T, stringsAsFactors = FALSE)
target.cats <- c("PROT_KEGG_METABOLISM_OF_XENOBIOTICS_BY_CYTOCHROME_P450")
GeneSet <- GeneSet[which(GeneSet$Description %in% target.cats),] 
GeneSet1<-GeneSet
tmp = rownames(pep1)
pep1 = pep1[tmp,]
GeneSet1 = GeneSet1[GeneSet1[,3] != "",]	# The first column in the data file must match the third column in the "gene list" file!
rnams = as.character(rownames(pep1))
xx = getMatchingRows(GeneSet1[,3], rnams) # Get the row number of matching rows
GeneMatch = pep1[xx,]
GeneMatch = GeneMatch[order(rownames(GeneMatch)),] 
transform_GeneMatch=apply(t(GeneMatch), c(1,2), as.numeric) # If NAs are coerced in, then it will force everything into a character matrix
tmpr <- corAndPvalue(transform_GeneMatch,method="spearman")
tmp = tmpr$cor
diag(tmp) = 0
outputmatrix_PROT_CYP450_CORRS=tmp[lower.tri(tmp)] 	

GeneSet=read.csv(file="~/Dropbox/Aging\ BXD\ Study/1\ _\ Manuscript\ -\ Evan\ Multi-Omic/b_HelpNetworks_ALL_NETWORKS_CANON_mRNA_Prot_ALIGNED.csv", header=T, stringsAsFactors = FALSE)
target.cats <- c("mRNA_KEGG_METABOLISM_OF_XENOBIOTICS_BY_CYTOCHROME_P450","PROT_KEGG_METABOLISM_OF_XENOBIOTICS_BY_CYTOCHROME_P450")
GeneSet <- GeneSet[which(GeneSet$Description %in% target.cats),] 
GeneSet1<-GeneSet
tmp = rownames(pep1)
pep1 = pep1[tmp,]
GeneSet1 = GeneSet1[GeneSet1[,3] != "",]	# The first column in the data file must match the third column in the "gene list" file!
rnams = as.character(rownames(pep1))
xx = getMatchingRows(GeneSet1[,3], rnams) # Get the row number of matching rows
GeneMatch = pep1[xx,]
GeneMatch = GeneMatch[order(rownames(GeneMatch)),] 
transform_GeneMatch=apply(t(GeneMatch), c(1,2), as.numeric) # If NAs are coerced in, then it will force everything into a character matrix
tmpr <- corAndPvalue(transform_GeneMatch,method="spearman")
tmp = tmpr$cor
diag(tmp) = 0
outputmatrix_ALL_CYP450_CORRS=tmp[lower.tri(tmp)] 
outputmatrix_ALL_CYP450_CORRS2=outputmatrix_ALL_CYP450_CORRS[!(outputmatrix_ALL_CYP450_CORRS %in% outputmatrix_mRNA_CYP450_CORRS)]
outputmatrix_ALL_CYP450_CORRS3=outputmatrix_ALL_CYP450_CORRS2[!(outputmatrix_ALL_CYP450_CORRS2 %in% outputmatrix_PROT_CYP450_CORRS)]

### NOTE: This "cross-layer" code has to be calculated last, and it suppresses the correlations that were found within protein and within mRNA, only retaining "across"

# Now we make the density plot
library(gplots)

plot(density(outputmatrix_ALL_CYP450_CORRS3, na.rm=T), xlim=c(-1,1), ylim=c(0, 6), col="black")
plotCI(density(outputmatrix_mRNA_CYP450_CORRS), col="blue", add=T, type="l")
plotCI(density(outputmatrix_PROT_CYP450_CORRS), col="red", add=T, type="l")

# BOTTOM-RIGHT
pep1=AllData

GeneSet=read.csv(file="~/Dropbox/Aging\ BXD\ Study/1\ _\ Manuscript\ -\ Evan\ Multi-Omic/b_HelpNetworks_ALL_NETWORKS_CANON_mRNA_Prot_ALIGNED.csv", header=T, stringsAsFactors = FALSE)
target.cats <- c("mRNA_REACTOME_BRANCHED_CHAIN_AMINO_ACID_CATABOLISM")
GeneSet <- GeneSet[which(GeneSet$Description %in% target.cats),] 
GeneSet1<-GeneSet
tmp = rownames(pep1)
pep1 = pep1[tmp,]
GeneSet1 = GeneSet1[GeneSet1[,3] != "",]	# The first column in the data file must match the third column in the "gene list" file!
rnams = as.character(rownames(pep1))
xx = getMatchingRows(GeneSet1[,3], rnams) # Get the row number of matching rows
GeneMatch = pep1[xx,]
GeneMatch = GeneMatch[order(rownames(GeneMatch)),] 
transform_GeneMatch=apply(t(GeneMatch), c(1,2), as.numeric) # If NAs are coerced in, then it will force everything into a character matrix
tmpr <- corAndPvalue(transform_GeneMatch,method="spearman")
tmp = tmpr$cor
diag(tmp) = 0
outputmatrix_mRNA_BCAAMetab_CORRS=tmp[lower.tri(tmp)] 	

GeneSet=read.csv(file="~/Dropbox/Aging\ BXD\ Study/1\ _\ Manuscript\ -\ Evan\ Multi-Omic/b_HelpNetworks_ALL_NETWORKS_CANON_mRNA_Prot_ALIGNED.csv", header=T, stringsAsFactors = FALSE)
target.cats <- c("PROT_REACTOME_BRANCHED_CHAIN_AMINO_ACID_CATABOLISM")
GeneSet <- GeneSet[which(GeneSet$Description %in% target.cats),] 
GeneSet1<-GeneSet
tmp = rownames(pep1)
pep1 = pep1[tmp,]
GeneSet1 = GeneSet1[GeneSet1[,3] != "",]	# The first column in the data file must match the third column in the "gene list" file!
rnams = as.character(rownames(pep1))
xx = getMatchingRows(GeneSet1[,3], rnams) # Get the row number of matching rows
GeneMatch = pep1[xx,]
GeneMatch = GeneMatch[order(rownames(GeneMatch)),] 
transform_GeneMatch=apply(t(GeneMatch), c(1,2), as.numeric) # If NAs are coerced in, then it will force everything into a character matrix
tmpr <- corAndPvalue(transform_GeneMatch,method="spearman")
tmp = tmpr$cor
diag(tmp) = 0
outputmatrix_PROT_BCAAMetab_CORRS=tmp[lower.tri(tmp)] 	

GeneSet=read.csv(file="~/Dropbox/Aging\ BXD\ Study/1\ _\ Manuscript\ -\ Evan\ Multi-Omic/b_HelpNetworks_ALL_NETWORKS_CANON_mRNA_Prot_ALIGNED.csv", header=T, stringsAsFactors = FALSE)
target.cats <- c("mRNA_REACTOME_BRANCHED_CHAIN_AMINO_ACID_CATABOLISM","PROT_REACTOME_BRANCHED_CHAIN_AMINO_ACID_CATABOLISM")
GeneSet <- GeneSet[which(GeneSet$Description %in% target.cats),] 
GeneSet1<-GeneSet
tmp = rownames(pep1)
pep1 = pep1[tmp,]
GeneSet1 = GeneSet1[GeneSet1[,3] != "",]	# The first column in the data file must match the third column in the "gene list" file!
rnams = as.character(rownames(pep1))
xx = getMatchingRows(GeneSet1[,3], rnams) # Get the row number of matching rows
GeneMatch = pep1[xx,]
GeneMatch = GeneMatch[order(rownames(GeneMatch)),] 
transform_GeneMatch=apply(t(GeneMatch), c(1,2), as.numeric) # If NAs are coerced in, then it will force everything into a character matrix
tmpr <- corAndPvalue(transform_GeneMatch,method="spearman")
tmp = tmpr$cor
diag(tmp) = 0
outputmatrix_ALL_BCAAMetab_CORRS=tmp[lower.tri(tmp)] 
outputmatrix_ALL_BCAAMetab_CORRS2=outputmatrix_ALL_BCAAMetab_CORRS[!(outputmatrix_ALL_BCAAMetab_CORRS %in% outputmatrix_mRNA_BCAAMetab_CORRS)]
outputmatrix_ALL_BCAAMetab_CORRS3=outputmatrix_ALL_BCAAMetab_CORRS2[!(outputmatrix_ALL_BCAAMetab_CORRS2 %in% outputmatrix_PROT_BCAAMetab_CORRS)]

### NOTE: This "cross-layer" code has to be calculated last, and it suppresses the correlations that were found within protein and within mRNA, only retaining "across"

# Now we make the density plot
library(gplots)

plot(density(outputmatrix_ALL_BCAAMetab_CORRS3, na.rm=T), xlim=c(-1,1), ylim=c(0, 6), col="black")
plotCI(density(outputmatrix_mRNA_BCAAMetab_CORRS), col="blue", add=T, type="l")
plotCI(density(outputmatrix_PROT_BCAAMetab_CORRS), col="red", add=T, type="l")


######################################################################################################################################################
######################################################################################################################################################
# CorrGraph function "CORRGRAPH.FIND"
######################################################################################################################################################
CorrGraph <- function(pep1, GeneSet1, cutofflevel, colors) {

tmp = rownames(pep1)
pep1 = pep1[tmp,]

GeneSet1 = GeneSet1[GeneSet1[,3] != "",]	# The first column in the data file must match the third column in the "gene list" file!
rnams = as.character(rownames(pep1))

xx = getMatchingRows(GeneSet1[,3], rnams) # Get the row number of matching rows
GeneMatch = pep1[xx,]
GeneMatch = GeneMatch[order(rownames(GeneMatch)),] 

siglevel = cutofflevel
transform_GeneMatch=apply(t(GeneMatch), c(1,2), as.numeric) # If NAs are coerced in, then it will force everything into a character matrix
tmpr <- corAndPvalue(transform_GeneMatch,method="spearman")
tmp = tmpr$cor
siz = dim(tmp)/2

diag(tmp) = 0
tmp[tmpr$p > siglevel] = 0 	# removes all correlation coefficients if their p-value is less than "siglevel" above.

# This sets the colors for the groups assigned in GeneList
mypaletteDiv = colors

# This makes sure the dots and colors correspond correctly
idx = match(rownames(tmp),as.character(GeneSet1[,3]))
graphnames = as.character(GeneSet1[idx,2])
colornames = as.numeric(GeneSet1[idx,5])

nedges = (sum(colSums(tmp !=0)) - sum(colSums(tmp == 1)))/2	# THIS COUNTS THE NUMBER OF EDGES, REMOVES SELF-EDGES, AND DIVIDES BY 2 (DOUBLED MATRIX)
nposedges = (sum(colSums(tmp >0)) - sum(colSums(tmp == 1)))/2	# THIS COUNTS THE NUMBER OF POSITIVE EDGES, REMOVES SELF-EDGES, AND DIVIDES BY 2 (DOUBLED MATRIX)
nnegedges = (sum(colSums(tmp <0)) - sum(colSums(tmp == 1)))/2	# THIS COUNTS THE NUMBER OF NEGATIVE EDGES, REMOVES SELF-EDGES, AND DIVIDES BY 2 (DOUBLED MATRIX)
nnodes = length(idx)
mean(tmpr$cor)		# Average correlation
mean(abs(tmpr$cor))	# Average absolute value of correlation

gr = plotGraph((tmp),names=graphnames, main = paste("P <", siglevel), vertcol=mypaletteDiv[colornames],repulserad=5e4)
plotGraph((tmp),names=graphnames, main = paste("P <", siglevel), vertcol=mypaletteDiv[colornames],repulserad=.6, nodesize=3.8, framecol="black", textcolor="black", myl = gr$myl)
legend(x=-1.4, y=1.2, bg='white', bty='n', legend=c(paste("Positive Edges = ", nposedges), paste("Negative Edges = ", nnegedges), paste("Node Count = ", nnodes)))

}
```
