## Supplementary material for "Multi-Omic Profiling of the Liver Across Diets and Age in a Diverse Mouse Population": All code used to generate figures: b_Figure5_Plotting_Code_vF.r

```
################################################################################  
# First priority is loading the multi-omics dataset, which will be used throughout
## This is the INDIVIDUAL data
################################################################################# 

setwd("~/Dropbox/Aging\ BXD\ Study/1\ _\ Manuscript\ -\ Evan\ Multi-Omic/")

AllData_header=read.table("aTableS2_ALL_DATA.csv", nrows=8, row.names=1, header=T, sep=",")
AllData_header=AllData_header[,3:ncol(AllData_header)]
AllData=read.table("aTableS2_ALL_DATA.csv", row.names=1, header=T,  na.strings="", sep=",", skip=8, stringsAsFactors=FALSE)
AllData=AllData[1:nrow(AllData),3:ncol(AllData)]

AllData_CD=matrix(NA, nrow=nrow(AllData), ncol=length(AllData))
for(i in 1:length(AllData)) {
	if(as.character(AllData_header["Diet",i])=="CD") {   AllData_CD[,i]=as.numeric(as.character(AllData[,i]))	}
} 
rownames(AllData_CD)=rownames(AllData)

AllData_HF=matrix(NA, nrow=nrow(AllData), ncol=length(AllData))
for(i in 1:length(AllData)) {
	if(as.character(AllData_header["Diet",i])=="HF") {   AllData_HF[,i]=as.numeric(as.character(AllData[,i]))	}
}
rownames(AllData_HF)=rownames(AllData)

AllData_young=matrix(NA, nrow=nrow(AllData), ncol=length(AllData))
for(i in 1:length(AllData)) {
	if(as.character(AllData_header["Age",i])<=420) {	AllData_young[,i]=as.numeric(as.character(AllData[,i]))	}
}
rownames(AllData_young)=rownames(AllData)

AllData_old=matrix(NA, nrow=nrow(AllData), ncol=length(AllData))
for(i in 1:length(AllData)) {
	if(as.character(AllData_header["Age",i])>=420) {	AllData_old[,i]=as.numeric(as.character(AllData[,i]))	}
}
rownames(AllData_old)=rownames(AllData)

################################################################################  
# FIGURE FIVE
################################################################################# 

library(imsbInfer)
library(WGCNA)
library(RColorBrewer)
library(ggbiplot)
library(outliers)
library(igraph)
source("~/Dropbox/Aging\ BXD\ Study/1\ _\ Manuscript\ -\ Evan\ Multi-Omic/b_helpFunctions_unilities.R")
source("~/Dropbox/Aging\ BXD\ Study/1\ _\ Manuscript\ -\ Evan\ Multi-Omic/b_helpFunctions.R")


#$$$$$$ #$$$$$$ #$$$$$$ #$$$$$$ #$$$$$$ #$$$$$$ #$$$$$$ #$$$$$$
#$$$$$$ #$$$$$$ #$$$$$$ #$$$$$$ #$$$$$$ #$$$$$$ #$$$$$$ #$$$$$$
# PANEL 5A
# Must load the above libraries and the "CorrGraph" function at the end of this file.

# LEFT PLOT
pep1=AllData_CD
GeneSet=read.csv(file="~/Dropbox/Aging\ BXD\ Study/1\ _\ Manuscript\ -\ Evan\ Multi-Omic/b_HelpNetworks_ALL_NETWORKS_CANON_mRNA_Prot_ALIGNED.csv", header=T, stringsAsFactors = FALSE)
target.cats <- c("PROT_REACTOME_CHOLESTEROL_BIOSYNTHESIS","PROT_REACTOME_CHOLESTEROL_BIOSYNTHESIS_NEWHITS")
GeneSet <- GeneSet[which(GeneSet$Description %in% target.cats),] 
GeneSet1<-GeneSet
CorrGraph(pep1, GeneSet1, 0.005, colors=c("green","red","green", "yellow","darkred")) 

# RIGHT PLOT
pep1=AllData_HF
GeneSet=read.csv(file="~/Dropbox/Aging\ BXD\ Study/1\ _\ Manuscript\ -\ Evan\ Multi-Omic/b_HelpNetworks_ALL_NETWORKS_CANON_mRNA_Prot_ALIGNED.csv", header=T, stringsAsFactors = FALSE)
target.cats <- c("PROT_REACTOME_CHOLESTEROL_BIOSYNTHESIS","PROT_REACTOME_CHOLESTEROL_BIOSYNTHESIS_NEWHITS")
GeneSet <- GeneSet[which(GeneSet$Description %in% target.cats),] 
GeneSet1<-GeneSet
CorrGraph(pep1, GeneSet1, 0.005, colors=c("green","red","green", "yellow","darkred")) 

#$$$$$$ #$$$$$$ #$$$$$$ #$$$$$$ #$$$$$$ #$$$$$$ #$$$$$$ #$$$$$$
#$$$$$$ #$$$$$$ #$$$$$$ #$$$$$$ #$$$$$$ #$$$$$$ #$$$$$$ #$$$$$$
# PANEL 5B

# LEFT (RDH11)
groupA=as.numeric(AllData_CD["Prot_12851_Rdh11",])
groupB=as.numeric(AllData_HF["Prot_12851_Rdh11",])
stripchart(groupA, at=0.7, pch=21, cex=1.2, bg="grey", method="jitter", xlim=c(0.6, 1.3), las=1, ylim=c(min(c(groupA,groupB), na.rm=T), max(c(groupA,groupB), na.rm=T)), vertical=T)
stripchart(groupB, at=1.0, pch=21, cex=1.2, bg="black", method="jitter", add=T, vertical=T)
	segments(0.65, quantile(groupA, na.rm=T)[2], 0.75, quantile(groupA, na.rm=T)[2], col="red", lwd=3)	# first quantile bar for first cohort
	segments(0.6, median(groupA, na.rm=T), 0.8, median(groupA, na.rm=T), col="red", lwd=3)	# median bar for first cohort
	segments(0.65, quantile(groupA, na.rm=T)[4], 0.75, quantile(groupA, na.rm=T)[4], col="red", lwd=3)	# first quantile bar for first cohort
	segments(0.7,quantile(groupA, na.rm=T)[2],0.7,quantile(groupA, na.rm=T)[4], col="red", lwd=3)
	segments(.95, quantile(groupB, na.rm=T)[2], 1.05, quantile(groupB, na.rm=T)[2], col="red", lwd=3)	# first quantile bar for second cohort	
	segments(.9, median(groupB, na.rm=T), 1.1, median(groupB, na.rm=T), col="red", lwd=3)	# median bar for second cohort, etc
	segments(.95, quantile(groupB, na.rm=T)[4], 1.05, quantile(groupB, na.rm=T)[4], col="red", lwd=3)	# first quantile bar for second cohort
	segments(1.0,quantile(groupB, na.rm=T)[2],1.0,quantile(groupB, na.rm=T)[4], col="red", lwd=3)
# legend	
pr = signif(t.test(groupA,groupB, paired=FALSE)$p.value,digits=2)			# This outputs the p-value of the SPEARMAN correlation but does not go below e-16 and I can't figure out the fix
pletter=substitute(paste("   ", italic("p"), " = ", pr), list(pr=signif(pr[1], digits=2)))	# Makes the r-value with italicized r

for (i in 1:1) {	# Why does this not work if it's not in a for loop??
	if (mean(groupA, na.rm=T)-mean(groupB, na.rm=T) >= 0) { # This calculates positive fold change
		fc=round(2^(mean(groupA, na.rm=T)-mean(groupB, na.rm=T)),2); legendposition="topright"
	} else if (mean(groupA, na.rm=T)-mean(groupB, na.rm=T) <= 0) { # This makes it into negative fold change, but SYMMETRICAL (i.e. -0.5 is the same as +2.0 but it's hard to tell)
		fc=round(-1*(2^(mean(groupB, na.rm=T)-mean(groupA, na.rm=T))),2); legendposition="topleft"
	} else {	fc = 0; legendposition="topleft"}
}
foldchange=paste("FC = ", fc)
legend(legendposition, bg='white', bty="n", cex=1, text.col="red", legend=c(foldchange, as.expression(pletter)))

# RIGHT (OBSCN)
groupA=as.numeric(AllData_CD["Prot_40085_Obscn",])
groupB=as.numeric(AllData_HF["Prot_40085_Obscn",])
stripchart(groupA, at=0.7, pch=21, cex=1.2, bg="grey", method="jitter", xlim=c(0.6, 1.3), las=1, ylim=c(min(c(groupA,groupB), na.rm=T), max(c(groupA,groupB), na.rm=T)), vertical=T)
stripchart(groupB, at=1.0, pch=21, cex=1.2, bg="black", method="jitter", add=T, vertical=T)
	segments(0.65, quantile(groupA, na.rm=T)[2], 0.75, quantile(groupA, na.rm=T)[2], col="red", lwd=3)	# first quantile bar for first cohort
	segments(0.6, median(groupA, na.rm=T), 0.8, median(groupA, na.rm=T), col="red", lwd=3)	# median bar for first cohort
	segments(0.65, quantile(groupA, na.rm=T)[4], 0.75, quantile(groupA, na.rm=T)[4], col="red", lwd=3)	# first quantile bar for first cohort
	segments(0.7,quantile(groupA, na.rm=T)[2],0.7,quantile(groupA, na.rm=T)[4], col="red", lwd=3)
	segments(.95, quantile(groupB, na.rm=T)[2], 1.05, quantile(groupB, na.rm=T)[2], col="red", lwd=3)	# first quantile bar for second cohort	
	segments(.9, median(groupB, na.rm=T), 1.1, median(groupB, na.rm=T), col="red", lwd=3)	# median bar for second cohort, etc
	segments(.95, quantile(groupB, na.rm=T)[4], 1.05, quantile(groupB, na.rm=T)[4], col="red", lwd=3)	# first quantile bar for second cohort
	segments(1.0,quantile(groupB, na.rm=T)[2],1.0,quantile(groupB, na.rm=T)[4], col="red", lwd=3)
# legend	
pr = signif(t.test(groupA,groupB, paired=FALSE)$p.value,digits=2)			# This outputs the p-value of the SPEARMAN correlation but does not go below e-16 and I can't figure out the fix
pletter=substitute(paste("   ", italic("p"), " = ", pr), list(pr=signif(pr[1], digits=2)))	# Makes the r-value with italicized r

for (i in 1:1) {	# Why does this not work if it's not in a for loop??
	if (mean(groupA, na.rm=T)-mean(groupB, na.rm=T) >= 0) { # This calculates positive fold change
		fc=round(2^(mean(groupA, na.rm=T)-mean(groupB, na.rm=T)),2); legendposition="topright"
	} else if (mean(groupA, na.rm=T)-mean(groupB, na.rm=T) <= 0) { # This makes it into negative fold change, but SYMMETRICAL (i.e. -0.5 is the same as +2.0 but it's hard to tell)
		fc=round(-1*(2^(mean(groupB, na.rm=T)-mean(groupA, na.rm=T))),2); legendposition="topleft"
	} else {	fc = 0; legendposition="topleft"}
}
foldchange=paste("FC = ", fc)
legend(legendposition, bg='white', bty="n", cex=1, text.col="red", legend=c(foldchange, as.expression(pletter)))


#$$$$$$ #$$$$$$ #$$$$$$ #$$$$$$ #$$$$$$ #$$$$$$ #$$$$$$ #$$$$$$
#$$$$$$ #$$$$$$ #$$$$$$ #$$$$$$ #$$$$$$ #$$$$$$ #$$$$$$ #$$$$$$
# PANEL 5C
# Stability-Prediction Plot
# Beforehand for all Stability–Prediction plots: on CRAN, install glmnet, questionr, and corpcor (w/ dependencies)
#
#install.packages("~/Documents/GitHub/StabilizedRegression-R/StabilizedRegression_1.0.tar.gz")
# Note that due to permutations, all the stability-prediction plots vary by up to around 5% per run, so a result that has a stability / prediction of 70% / 50% in one run might have e.g. 65% / 55% in the next.

library(StabilizedRegression)

## LEFT PANEL
# This is a reformatted version of the "AllData" file, with the top transcript per gene (taking the first one in the list) for all protein–mRNA matching genes
# THEN, as high numbers of reads of "0" distort correlations, any transcript with > 50 0's was discarded, so this mRNA file is somewhat smaller (3677 genes)
data=read.csv(file="~/Dropbox/Aging\ BXD\ Study/1\ _\ Manuscript\ -\ Evan\ Multi-Omic/b_Figure5_HelperFile_mRNAData.csv", header=T, stringsAsFactors = FALSE)
# The pathway info is the same as used for Figure 4, but now just matching based on gene symbol, so the other columns of metadata are stripped out. 
pathway_info=read.csv(file="~/Dropbox/Aging\ BXD\ Study/1\ _\ Manuscript\ -\ Evan\ Multi-Omic/b_Figure5_HelperFile_GeneSets.csv", header=T, stringsAsFactors = FALSE)

TargetPathway=pathway_info$REACTOME_CHOLESTEROL_BIOSYNTHESIS
TargetPathway=TargetPathway[TargetPathway != ""]
Xmat <- as.matrix(data[,-(1:3)]) # This drops the multiple header columns
env <- data[,2] ## This sets either diet or age as the key variable. 3 = age, 2 = diet (for these input files)
varnames <- colnames(Xmat)
A <- env ## Set variables for a single target gene against all others, and the controlling variable (env).
## now to compare against a full pathway and not just a target gene
X = Xmat[,!colnames(Xmat) %in% TargetPathway]
Y = Xmat[,colnames(Xmat) %in% TargetPathway]
Y=rowMeans(Y)

## Fix parameters
cores <- 3 # Can change depending on your computer
Sys.time()
pars <- list(m=6, B=5000, alpha_stab=0.1, alpha_pred=0.01, size_weight="linear", prescreen_size=50, use_resampling=FALSE,stab_test="exact",variable_importance="scaled_coefficient")

# Num reps sets minimum p-value (e.g. 100 == p = 0.01 increments, p = 10, p = 0.10 increments); this is what takes the most time.
fit_obj <- SRanalysis(X, Y, A, num_reps=100, pars_SR=pars, cores=cores)
plot(fit_obj, x_axis="SRpred", varnames=colnames(X), labels=TRUE)

# If desired, the actual values of the plot can be output to a table (can attach this to every stability-regression analysis; code only shown here)
write.table(fit_obj$results, "~/Desktop/REACTOME_CHOLESTEROL_BIOSYNTHESIS_DIET_mRNA.csv",sep=",")


## RIGHT PANEL
data=read.csv(file="~/Dropbox/Aging\ BXD\ Study/1\ _\ Manuscript\ -\ Evan\ Multi-Omic/b_Figure5_HelperFile_ProteinData.csv", header=T, stringsAsFactors = FALSE)
# The pathway info is the same as used for Figure 4, but now just matching based on gene symbol, so the other columns of metadata are stripped out. 
pathway_info=read.csv(file="~/Dropbox/Aging\ BXD\ Study/1\ _\ Manuscript\ -\ Evan\ Multi-Omic/b_Figure5_HelperFile_GeneSets.csv", header=T, stringsAsFactors = FALSE)

TargetPathway=pathway_info$REACTOME_CHOLESTEROL_BIOSYNTHESIS
TargetPathway=TargetPathway[TargetPathway != ""]
Xmat <- as.matrix(data[,-(1:3)]) # This drops the multiple header columns
env <- data[,2] ## This sets either diet or age as the key variable. 3 = age, 2 = diet (for these input files)
varnames <- colnames(Xmat)
A <- env ## Set variables for a single target gene against all others, and the controlling variable (env).
## now to compare against a full pathway and not just a target gene
X = Xmat[,!colnames(Xmat) %in% TargetPathway]
Y = Xmat[,colnames(Xmat) %in% TargetPathway]
Y=rowMeans(Y)

## Fix parameters
cores <- 3 # Can change depending on your computer
Sys.time()
pars <- list(m=6, B=5000, alpha_stab=0.1, alpha_pred=0.01, size_weight="linear", prescreen_size=50, use_resampling=FALSE,stab_test="exact",variable_importance="scaled_coefficient")

# Num reps sets minimum p-value (e.g. 100 == p = 0.01 increments, p = 10, p = 0.10 increments); this is what takes the most time.
fit_obj <- SRanalysis(X, Y, A, num_reps=100, pars_SR=pars, cores=cores)
plot(fit_obj, x_axis="SRpred", varnames=colnames(X), labels=TRUE)

write.table(fit_obj$results, "~/Desktop/REACTOME_CHOLESTEROL_BIOSYNTHESIS_DIET_PROT.csv",sep=",")

#$$$$$$ #$$$$$$ #$$$$$$ #$$$$$$ #$$$$$$ #$$$$$$ #$$$$$$ #$$$$$$
#$$$$$$ #$$$$$$ #$$$$$$ #$$$$$$ #$$$$$$ #$$$$$$ #$$$$$$ #$$$$$$
# PANEL 5D
library(StabilizedRegression)

# Stability-Prediction Plot

## LEFT PANEL
# This is a reformatted version of the "AllData" file, with the top transcript per gene (taking the first one in the list) for all protein–mRNA matching genes
# THEN, as high numbers of reads of "0" distort correlations, any transcript with > 50 0's was discarded, so this mRNA file is somewhat smaller (3677 genes)
data=read.csv(file="~/Dropbox/Aging\ BXD\ Study/1\ _\ Manuscript\ -\ Evan\ Multi-Omic/b_Figure5_HelperFile_mRNAData.csv", header=T, stringsAsFactors = FALSE)
# The pathway info is the same as used for Figure 4, but now just matching based on gene symbol, so the other columns of metadata are stripped out. 
pathway_info=read.csv(file="~/Dropbox/Aging\ BXD\ Study/1\ _\ Manuscript\ -\ Evan\ Multi-Omic/b_Figure5_HelperFile_GeneSets.csv", header=T, stringsAsFactors = FALSE)

TargetPathway=pathway_info$KEGG_METABOLISM_OF_XENOBIOTICS_BY_CYTOCHROME_P450
TargetPathway=TargetPathway[TargetPathway != ""]
Xmat <- as.matrix(data[,-(1:3)]) # This drops the multiple header columns
env <- data[,2] ## This sets either diet or age as the key variable. 3 = age, 2 = diet (for these input files)
varnames <- colnames(Xmat)
A <- env ## Set variables for a single target gene against all others, and the controlling variable (env).
## now to compare against a full pathway and not just a target gene
X = Xmat[,!colnames(Xmat) %in% TargetPathway]
Y = Xmat[,colnames(Xmat) %in% TargetPathway]
Y=rowMeans(Y)

## Fix parameters
cores <- 3 # Can change depending on your computer
Sys.time()
pars <- list(m=6, B=5000, alpha_stab=0.1, alpha_pred=0.01, size_weight="linear", prescreen_size=50, use_resampling=FALSE,stab_test="exact",variable_importance="scaled_coefficient")

# Num reps sets minimum p-value (e.g. 100 == p = 0.01 increments, p = 10, p = 0.10 increments); this is what takes the most time.
fit_obj <- SRanalysis(X, Y, A, num_reps=100, pars_SR=pars, cores=cores)
plot(fit_obj, x_axis="SRpred", varnames=colnames(X), labels=TRUE)

# If desired, the actual values of the plot can be output to a table (can attach this to every stability-regression analysis; code only shown here)
write.table(fit_obj$results, "~/Desktop/REACTOME_CYP450_DIET_mRNA.csv",sep=",")

## RIGHT PANEL
# This is a reformatted version of the "AllData" file, with the top transcript per gene (taking the first one in the list) for all protein–mRNA matching genes
# THEN, as high numbers of reads of "0" distort correlations, any transcript with > 50 0's was discarded, so this mRNA file is somewhat smaller (3677 genes)
data=read.csv(file="~/Dropbox/Aging\ BXD\ Study/1\ _\ Manuscript\ -\ Evan\ Multi-Omic/b_Figure5_HelperFile_mRNAData.csv", header=T, stringsAsFactors = FALSE)
# The pathway info is the same as used for Figure 4, but now just matching based on gene symbol, so the other columns of metadata are stripped out. 
pathway_info=read.csv(file="~/Dropbox/Aging\ BXD\ Study/1\ _\ Manuscript\ -\ Evan\ Multi-Omic/b_Figure5_HelperFile_GeneSets.csv", header=T, stringsAsFactors = FALSE)

TargetPathway=pathway_info$KEGG_METABOLISM_OF_XENOBIOTICS_BY_CYTOCHROME_P450
TargetPathway=TargetPathway[TargetPathway != ""]
Xmat <- as.matrix(data[,-(1:3)]) # This drops the multiple header columns
env <- data[,3] ## This sets either diet or age as the key variable. 3 = age, 2 = diet (for these input files)
varnames <- colnames(Xmat)
A <- env ## Set variables for a single target gene against all others, and the controlling variable (env).
## now to compare against a full pathway and not just a target gene
X = Xmat[,!colnames(Xmat) %in% TargetPathway]
Y = Xmat[,colnames(Xmat) %in% TargetPathway]
Y=rowMeans(Y)

## Fix parameters
cores <- 3 # Can change depending on your computer
Sys.time()
pars <- list(m=6, B=5000, alpha_stab=0.1, alpha_pred=0.01, size_weight="linear", prescreen_size=50, use_resampling=FALSE,stab_test="exact",variable_importance="scaled_coefficient")

# Num reps sets minimum p-value (e.g. 100 == p = 0.01 increments, p = 10, p = 0.10 increments); this is what takes the most time.
fit_obj <- SRanalysis(X, Y, A, num_reps=100, pars_SR=pars, cores=cores)
plot(fit_obj, x_axis="SRpred", varnames=colnames(X), labels=TRUE)

# If desired, the actual values of the plot can be output to a table (can attach this to every stability-regression analysis; code only shown here)
write.table(fit_obj$results, "~/Desktop/REACTOME_CYP450_AGE_mRNA.csv",sep=",")

#$$$$$$ #$$$$$$ #$$$$$$ #$$$$$$ #$$$$$$ #$$$$$$ #$$$$$$ #$$$$$$
#$$$$$$ #$$$$$$ #$$$$$$ #$$$$$$ #$$$$$$ #$$$$$$ #$$$$$$ #$$$$$$
# PANEL 5E

# Output from DAVID ( https://david.ncifcrf.gov/summary.jsp ) using hits from the CYP450 set:
#Ces1c,Ces1d,Ces1e,Ces1f,Ces1g,Ces2a,Cyp2a5,Cyp2c29,Cyp2c39,Cyp2c40,Cyp2c54,Cyp2f2,Gstm6,Gstm7,Prdx6,Cp,Hsd11b1,Mdh1,Nqo1,Nqo2,Sod1,Ugdh,Uqcrc1,Akr1c13,Hsd17b6,Rdh7,Akr1e1,Akr1a1,Hibadh,Sqrdl,Ndufa9,Sdhb,Cryl1,Pir,Bdh2,Aldh1b1,Dhcr24,Akr1c6,Cyp4v3,Gldc,Aldh1l1,Qdpr,Pecr,Akr1d1,L2hgdh,Blvrb,Creg1,Aadac,Afmid,Car3,Car5a,Ddah1,Echdc3,Eprs,Ftl1,Galm,Gcat,Hagh,Hrsp12,Inmt,Itih3,Macrod1,Mavs,Mybpc3,Myo1c,Nipsnap1,Nup155,Obscn,Otc,Park7,Pgrmc1,Phkb,Pon1,R3hcc1,Rgn,Selenbp1,Serpina3n,Serpinb1a,Tmem205,Ugt2b5,Upb1
# Against background of all 3772 genes in the mRNA and protein list

#$$$$$$ #$$$$$$ #$$$$$$ #$$$$$$ #$$$$$$ #$$$$$$ #$$$$$$ #$$$$$$
#$$$$$$ #$$$$$$ #$$$$$$ #$$$$$$ #$$$$$$ #$$$$$$ #$$$$$$ #$$$$$$
# PANEL 5F

# Stability-Prediction Plot
# Beforehand for all Stability–Prediction plots: on CRAN, install glmnet, questionr, and corpcor (w/ dependencies)
#
#install.packages("~/Documents/GitHub/StabilizedRegression-R/StabilizedRegression_1.0.tar.gz")
# Note that due to permutations, all the stability-prediction plots vary very slightly and cannot be PERFECTLY reproduced. However, there should be no more than a few percent of variance.

library(StabilizedRegression)

## LEFT PANEL
# This is a reformatted version of the "AllData" file, with the top transcript per gene (taking the first one in the list) for all protein–mRNA matching genes
# THEN, as high numbers of reads of "0" distort correlations, any transcript with > 50 0's was discarded, so this mRNA file is somewhat smaller (3677 genes)
data=read.csv(file="~/Dropbox/Aging\ BXD\ Study/1\ _\ Manuscript\ -\ Evan\ Multi-Omic/b_Figure5_HelperFile_mRNAData.csv", header=T, stringsAsFactors = FALSE)
# The pathway info is the same as used for Figure 4, but now just matching based on gene symbol, so the other columns of metadata are stripped out. 
pathway_info=read.csv(file="~/Dropbox/Aging\ BXD\ Study/1\ _\ Manuscript\ -\ Evan\ Multi-Omic/b_Figure5_HelperFile_GeneSets.csv", header=T, stringsAsFactors = FALSE)

TargetPathway=pathway_info$REACTOME_MITOCHONDRIAL_TRANSLATION
TargetPathway=TargetPathway[TargetPathway != ""]
Xmat <- as.matrix(data[,-(1:3)]) # This drops the multiple header columns
env <- data[,3] ## This sets either diet or age as the key variable. 3 = age, 2 = diet (for these input files)
varnames <- colnames(Xmat)
A <- env ## Set variables for a single target gene against all others, and the controlling variable (env).
## now to compare against a full pathway and not just a target gene
X = Xmat[,!colnames(Xmat) %in% TargetPathway]
Y = Xmat[,colnames(Xmat) %in% TargetPathway]
Y=rowMeans(Y)

## Fix parameters
cores <- 3 # Can change depending on your computer
Sys.time()
pars <- list(m=6, B=5000, alpha_stab=0.1, alpha_pred=0.01, size_weight="linear", prescreen_size=50, use_resampling=FALSE,stab_test="exact",variable_importance="scaled_coefficient")

# Num reps sets minimum p-value (e.g. 100 == p = 0.01 increments, p = 10, p = 0.10 increments); this is what takes the most time.
fit_obj <- SRanalysis(X, Y, A, num_reps=100, pars_SR=pars, cores=cores)
plot(fit_obj, x_axis="SRpred", varnames=colnames(X), labels=TRUE)

# If desired, the actual values of the plot can be output to a table (can attach this to every stability-regression analysis; code only shown here)
write.table(fit_obj$results, "~/Desktop/REACTOME_MITO_TRANSLATION_AGE_mRNA.csv",sep=",")


## RIGHT PANEL
data=read.csv(file="~/Dropbox/Aging\ BXD\ Study/1\ _\ Manuscript\ -\ Evan\ Multi-Omic/b_Figure5_HelperFile_ProteinData.csv", header=T, stringsAsFactors = FALSE)
# The pathway info is the same as used for Figure 4, but now just matching based on gene symbol, so the other columns of metadata are stripped out. 
pathway_info=read.csv(file="~/Dropbox/Aging\ BXD\ Study/1\ _\ Manuscript\ -\ Evan\ Multi-Omic/b_Figure5_HelperFile_GeneSets.csv", header=T, stringsAsFactors = FALSE)

TargetPathway=pathway_info$REACTOME_MITOCHONDRIAL_TRANSLATION
TargetPathway=TargetPathway[TargetPathway != ""]
Xmat <- as.matrix(data[,-(1:3)]) # This drops the multiple header columns
env <- data[,3] ## This sets either diet or age as the key variable. 3 = age, 2 = diet (for these input files)
varnames <- colnames(Xmat)
A <- env ## Set variables for a single target gene against all others, and the controlling variable (env).
## now to compare against a full pathway and not just a target gene
X = Xmat[,!colnames(Xmat) %in% TargetPathway]
Y = Xmat[,colnames(Xmat) %in% TargetPathway]
Y=rowMeans(Y)

## Fix parameters
cores <- 3 # Can change depending on your computer
Sys.time()
pars <- list(m=6, B=5000, alpha_stab=0.1, alpha_pred=0.01, size_weight="linear", prescreen_size=50, use_resampling=FALSE,stab_test="exact",variable_importance="scaled_coefficient")

# Num reps sets minimum p-value (e.g. 100 == p = 0.01 increments, p = 10, p = 0.10 increments); this is what takes the most time.
fit_obj <- SRanalysis(X, Y, A, num_reps=100, pars_SR=pars, cores=cores)
plot(fit_obj, x_axis="SRpred", varnames=colnames(X), labels=TRUE)

write.table(fit_obj$results, "~/Desktop/REACTOME_MITO_TRANSLATION_AGE_PROT.csv",sep=",")


#$$$$$$ #$$$$$$ #$$$$$$ #$$$$$$ #$$$$$$ #$$$$$$ #$$$$$$ #$$$$$$
#$$$$$$ #$$$$$$ #$$$$$$ #$$$$$$ #$$$$$$ #$$$$$$ #$$$$$$ #$$$$$$
# PANEL 5G

# Output from DAVID ( https://david.ncifcrf.gov/summary.jsp ) using hits from the Mitochondrial Translation (aka mito ribosome) set:
#Rpl11,Rps27l,Eif3i,Eif5a,Aco2,Aass,Atad3a,Cpt1b,Chchd2,Dld,Dlst,Echs1,Fam162a,Fis1,Got2,Gpx4,Grpel1,Hspe1,Hspa9,Hebp1,Hsd17b10,Lrpprc,Lonp1,Mdh2,Mcee,Mut,Mecr,Atp5e,Atp5f1,Atp5j,Atp5j2,Atp5o,Atp5s,Cox4i1,Cox5a,Cox6a1,Cox6b1,Cox6c,Cox7a2,Cox7c,Ndufa11,Ndufa13,Ndufa4,Ndufa7,Ndufb2,Ndufb4,Ndufb7,Ndufb8,Ndufb9,Ndufs4,Ndufs5,Ndufs6,Ndufs8,Sdha,Sdhb,Sdhd,Uqcr10,Uqcr11,Uqcrc1,Uqcrfs1,Uqcrq,Pmpca,Phb,Pccb,Psmb3,Ppa2,Pdhb,Pdha1,Slc25a3,Spryd4,Slirp,Txn2,Timm13,Timm17b,Timm8a1,Timm8b,Usmg5,Dap3,Ict1,Oxa1l,Tufm,Psma7,Psmb1,Psmb4,Psmb6,Apoc4,Calb2,Cap1,Dpm3,Hint1,Lpp,Mllt4,Nedd8,Nup62,Prrc2c,Selenbp1,Snrpd2,Ssr4,Tmed1,Ubr5
# Against background of all 3772 genes in the mRNA and protein list

#$$$$$$ #$$$$$$ #$$$$$$ #$$$$$$ #$$$$$$ #$$$$$$ #$$$$$$ #$$$$$$
#$$$$$$ #$$$$$$ #$$$$$$ #$$$$$$ #$$$$$$ #$$$$$$ #$$$$$$ #$$$$$$
# PANEL 5H

# Stability-Prediction Plot
library(StabilizedRegression)

## LEFT PANEL
# Body Weights, taken from Table S2 (all data) and aligned to the mRNA data "helper file" for this figure.
Pheno_BodyWeight=c(23.92,31.53,38.95,33.57,43.95,22.54,29.07,30.89,36.29,20.23,23.93,20.34,36.52,29.16,38.44,42.04,45.88,41.28,26.09,35.66,29.63,20.09,25.9,38.92,41.8,28.43,26.15,25.6,30.5,56.92,25.5,30.9,20.6,33.54,24.49,35.42,21.53,22.68,24.28,19.9,35.79,21.69,42.13,41.77,29.18,38.91,41.1,58.36,22.8,31.39,43.59,25.5,25.2,42.16,28.3,23.8,33.3,25.98,26.79,23.7,24.92,44.34,41.76,55.38,31.65,32.85,25.75,37.48,52.18,20.6,44.49,19.25,34.91,36.39,33.54,25,25.97,35.89,24.01,39.86,38.32,46.1,29.24,53.03,38.15,59.91,26.01,30.37,40.13,54.56,49.3,28.57,35.51,37.09,22.54,41.25,52.66,33.22,45.3,23.07,24.35,28.6,21.58,55.89,23.91,30.05,28.82,57.1,38.47,62.64,28.72,24.38,31.99,31.34,25.3,33.79,22.59,34.55,24.92,61.94,50.52,47.55,52.8,41.86,53.18,33.03,39.01,37.86,59.23,32.36,53.39,31.09,27.31,83.04,33.63,66.61,46.96,38.63,35.23,61.03,31.77,32.3,28.26,31.24,40.41,22.04,50.46,24.08,27.78,23.47,24.16,22.32,21.88,41.33,24.45,23.49,30.88,34.53,49.18,20.64,39.53,22.28,59.86,23.71,34.51,26.44,46.35,27.95,23.61,41.27,38.88,56.98,23.54,26.74,34.34,24.64,55.8,35.22,29.25,32.17,31.8,36.58,58.54,57.14,30.76,35.67,24.14,23.46,23.02,22.8,24.79,69.25,27.83,69.43,24.55,62.87,51.32,20.47,46.04,54.94,27.49,62.99,45.04,47.79,53.6,52.31,37.9,76.17,25.5,37.87,48.47,29.9,72.2,91.3,64.44,29.66,22.72,60.33,38.96,44.71,25.52,22.93,18.09,51.29,27,21.4,25.92,25.34,33.33,29.26,68.1,22.78,29.4,57.88,41.9,35.47,22.37,24.98,29.58,49.83,29.13,71.41,19.67,27.92,40.8,45.93,49.86,27.02,23.87,28.35,46.02,45.94,32.34,63.19,25.44,24.74,31.39,26.04,24.45,26.42,58.49,28,27.03,31.45,21.33,24.23,83.62,43.79,44.14,37.79,21.3,21.33,29.72,28.68,28.02,47.41,16.48,18.21,26.56,50.21,58.93,22.4,30.3,60.13,25.22,25.27,17.14,28.59,26.37,24.57,22.27)
Pheno_BodyWeight2=Pheno_BodyWeight # Ugly workaround to keep everything in matrix format. Data are averaged anyway so this does not impact results. 


# This is a reformatted version of the "AllData" file, with the top transcript per gene (taking the first one in the list) for all protein–mRNA matching genes
# THEN, as high numbers of reads of "0" distort correlations, any transcript with > 50 0's was discarded, so this mRNA file is somewhat smaller (3677 genes)
data=read.csv(file="~/Dropbox/Aging\ BXD\ Study/1\ _\ Manuscript\ -\ Evan\ Multi-Omic/b_Figure5_HelperFile_mRNAData.csv", header=T, stringsAsFactors = FALSE)
data=cbind(data,Pheno_BodyWeight) ## Body weight now added to the end of the file
data=cbind(data,Pheno_BodyWeight2) ## Body weight now added to the end of the file again; keeps everything in matrix format rather than vectorizing when target gene set n == 1 which results in a bug with some functions like rowMeans.

# The pathway info is the same as used for Figure 4, but now just matching based on gene symbol, so the other columns of metadata are stripped out. 
pathway_info=read.csv(file="~/Dropbox/Aging\ BXD\ Study/1\ _\ Manuscript\ -\ Evan\ Multi-Omic/b_Figure5_HelperFile_GeneSets.csv", header=T, stringsAsFactors = FALSE)

TargetPathway=pathway_info$BodyWeight
TargetPathway=TargetPathway[TargetPathway != ""]
Xmat <- as.matrix(data[,-(1:3)]) # This drops the multiple header columns
env <- data[,2] ## This sets either diet or age as the key variable. 3 = age, 2 = diet (for these input files)
varnames <- colnames(Xmat)
A <- env ## Set variables for a single target gene against all others, and the controlling variable (env).
## now to compare against a full pathway and not just a target gene
X = Xmat[,!colnames(Xmat) %in% TargetPathway]
Y = Xmat[,colnames(Xmat) %in% TargetPathway]

pars <- list(m=6, B=5000, alpha_stab=0.1, alpha_pred=0.01, size_weight="linear", prescreen_size=50, use_resampling=FALSE,stab_test="exact",variable_importance="scaled_coefficient")

# Num reps sets minimum p-value (e.g. 100 == p = 0.01 increments, p = 10, p = 0.10 increments); this is what takes the most time.
fit_obj <- SRanalysis(X, Y, A, num_reps=100, pars_SR=pars, cores=cores)
plot(fit_obj, x_axis="SRpred", varnames=colnames(X), labels=TRUE)

# If desired, the actual values of the plot can be output to a table (can attach this to every stability-regression analysis; code only shown here)
write.table(fit_obj$results, "~/Desktop/BodyWeight_DIET_mRNA.csv",sep=",")

### RIGHT PANEL:

# Body Weights, taken from Table S2 (all data) and aligned to the mRNA data "helper file" for this figure.
Pheno_BodyWeight=c(53.39,35.51,38.44,45.88,41.28,46.02,37.09,27.02,24.57,30.88,41.25,53.03,34.53,63.19,38.15,45.94,49.86,58.49,26.56,22.27,17.14,31.53,20.64,50.21,21.33,23.47,40.41,22.04,28.72,24.23,21.3,27,83.62,50.46,24.16,43.79,33.54,25.92,25.34,35.79,55.89,31.09,21.53,22.68,60.33,37.86,24.38,31.99,23.07,38.96,23.91,31.34,30.05,27.31,29.13,49.18,38.63,24.08,44.71,22.32,47.79,53.6,57.88,61.94,39.53,22.59,25.3,51.32,36.58,27.49,55.8,69.25,18.21,20.09,22.54,58.54,57.14,28.02,38.95,29.07,29.58,47.55,30.76,28.59,38.32,33.22,58.93,22.93,24.01,31.8,30.37,71.41,62.99,52.31,27.03,35.89,24.64,42.04,72.2,16.48,46.04,23.71,44.34,52.18,36.39,38.88,41.76,56.98,20.6,25.5,41.8,25.9,28.43,55.38,25,44.49,40.8,30.9,36.52,24.74,24.28,46.35,59.86,34.51,26.44,41.27,23.54,24.35,33.57,26.74,37.79,27.83,26.37,24.49,33.3,25.98,25.5,35.23,28,29.18,33.63,41.33,35.42,42.13,25.5,25.75,37.48,29.4,27.95,23.8,22.54,29.16,23.92,32.85,19.25,28.6,32.3,40.13,46.1,33.33,41.9,29.26,20.6,21.69,38.91,59.23,34.91,26.79,21.88,23.7,33.79,24.45,30.5,32.34,32.17,25.27,23.46,23.02,27.92,31.77,60.13,26.15,19.9,43.95,23.49,22.8,24.92,56.92,34.55,21.33,24.98,52.8,57.1,25.2,61.03,45.04,41.86,22.4,44.14,49.83,91.3,22.37,42.16,54.94,47.41,29.72,37.9,28.3,24.55,41.77,62.87,66.61,28.57,22.78,48.47,33.03,58.36,39.01,25.52,24.45,64.44,31.39,35.47,25.6,76.17,45.3,36.29,29.9,38.92,38.47,24.92,26.01,35.66,29.63,60.33,29.24,36.36,55.68,23.02,28.41,22.71,20.2,35.59,59.73,52.66,36.26,58.3,25.97,83.04,34.34,35.22,25.56,18.09,24.79,22.8,54.56,27.78,31.39,25.44,33.54,30.89,28.82,26.04,20.47,23.75,15.71,45.93,73.7,29.25,50.52,30.3,60.71,24.92,82.47,20.23,37.87,19.1,28.35,39.86,28.68,68.1,26.42,49.3,22.28,23.93,51.18,21.4,31.45,42.05,24.53,38.86,43.5,63.35,63.7,25,29.4,31.97,32.95,28.26,65.27,53.18,24.4,27.19,26.36,41.1,32.36,35.15,59.91,47.98,45.43,46.96,49.33,24.95,43.59,26.09)
Pheno_BodyWeight2=Pheno_BodyWeight # Ugly workaround to keep everything in matrix format. Data are averaged anyway so this does not impact results. 

data=read.csv(file="~/Dropbox/Aging\ BXD\ Study/1\ _\ Manuscript\ -\ Evan\ Multi-Omic/b_Figure5_HelperFile_ProteinData.csv", header=T, stringsAsFactors = FALSE)
data=cbind(data,Pheno_BodyWeight) ## Body weight now added to the end of the file
data=cbind(data,Pheno_BodyWeight2) ## Body weight now added to the end of the file

# The pathway info is the same as used for Figure 4, but now just matching based on gene symbol, so the other columns of metadata are stripped out. 
pathway_info=read.csv(file="~/Dropbox/Aging\ BXD\ Study/1\ _\ Manuscript\ -\ Evan\ Multi-Omic/b_Figure5_HelperFile_GeneSets.csv", header=T, stringsAsFactors = FALSE)

TargetPathway=pathway_info$BodyWeight
TargetPathway=TargetPathway[TargetPathway != ""]
Xmat <- as.matrix(data[,-(1:3)]) # This drops the multiple header columns
env <- data[,2] ## This sets either diet or age as the key variable. 3 = age, 2 = diet (for these input files)
varnames <- colnames(Xmat)
A <- env ## Set variables for a single target gene against all others, and the controlling variable (env).
## now to compare against a full pathway and not just a target gene
X = Xmat[,!colnames(Xmat) %in% TargetPathway]
Y = Xmat[,colnames(Xmat) %in% TargetPathway]
Y=rowMeans(Y)

## Fix parameters
cores <- 3 # Can change depending on your computer
Sys.time()
pars <- list(m=6, B=5000, alpha_stab=0.1, alpha_pred=0.01, size_weight="linear", prescreen_size=50, use_resampling=FALSE,stab_test="exact",variable_importance="scaled_coefficient")

# Num reps sets minimum p-value (e.g. 100 == p = 0.01 increments, p = 10, p = 0.10 increments); this is what takes the most time.
fit_obj <- SRanalysis(X, Y, A, num_reps=100, pars_SR=pars, cores=cores)
plot(fit_obj, x_axis="SRpred", varnames=colnames(X), labels=TRUE)

# If desired, the actual values of the plot can be output to a table (can attach this to every stability-regression analysis; code only shown here)
write.table(fit_obj$results, "~/Desktop/BodyWeight_DIET_Protein.csv",sep=",")

#$$$$$$ #$$$$$$ #$$$$$$ #$$$$$$ #$$$$$$ #$$$$$$ #$$$$$$ #$$$$$$
#$$$$$$ #$$$$$$ #$$$$$$ #$$$$$$ #$$$$$$ #$$$$$$ #$$$$$$ #$$$$$$
# PANEL 5I
library(StabilizedRegression)


# Stability-Prediction Plot
# LEFT PANEL:
# Separating based on COX7A2L allele rather than diet or age
cox7a2l_allele=c("D2","B6","D2","B6","D2","D2","B6","B6","B6","B6","B6","B6","B6","B6","B6","B6","D2","D2","D2","B6","B6","D2","B6","B6","B6","B6","B6","D2","B6","B6","D2","D2","D2","B6","B6","B6","D2","D2","D2","D2","B6","B6","D2","Heterozygous","D2","D2","D2","Heterozygous","B6","Heterozygous","Heterozygous","D2","D2","D2","B6","D2","B6","B6","B6","B6","D2","D2","B6","Heterozygous","B6","B6","D2","D2","D2","B6","B6","B6","D2","D2","B6","B6","D2","D2","D2","D2","B6","D2","B6","B6","B6","B6","D2","B6","D2","B6","B6","B6","B6","B6","B6","B6","B6","B6","B6","B6","B6","B6","B6","D2","D2","D2","B6","B6","D2","B6","B6","D2","D2","B6","B6","D2","D2","D2","D2","B6","B6","B6","Heterozygous","D2","Heterozygous","Heterozygous","D2","B6","B6","D2","B6","D2","D2","Heterozygous","D2","D2","B6","Heterozygous","B6","B6","D2","D2","D2","D2","B6","B6","B6","B6","B6","B6","B6","B6","B6","D2","B6","D2","D2","D2","D2","D2","Heterozygous","B6","D2","B6","D2","D2","Heterozygous","D2","D2","D2","B6","B6","D2","B6","B6","B6","D2","D2","B6","D2","B6","B6","B6","D2","D2","D2","B6","B6","D2","B6","D2","B6","B6","B6","B6","B6","D2","D2","D2","Heterozygous","D2","D2","B6","D2","D2","B6","D2","B6","B6","B6","Heterozygous","B6","Heterozygous","Heterozygous","B6","B6","B6","D2","D2","D2","D2","B6","B6","D2","Heterozygous","B6","B6","B6","D2","B6","B6","B6","B6","D2","D2","B6","B6","B6","B6","B6","D2","D2","D2","B6","B6","B6","B6","B6","B6","B6","B6","Heterozygous","D2","D2","D2","B6","B6","B6","D2","D2","B6","B6","B6","D2","B6","B6","Heterozygous","Heterozygous","B6","D2","B6","D2","B6","B6","B6","B6","D2","D2","B6","B6","B6","B6","B6","B6","B6","D2","D2","D2","B6","D2","D2")

# The pathway info is the same as used for Figure 4, but now just matching based on gene symbol, so the other columns of metadata are stripped out. 
data=read.csv(file="~/Dropbox/Aging\ BXD\ Study/1\ _\ Manuscript\ -\ Evan\ Multi-Omic/b_Figure5_HelperFile_mRNAData.csv", header=T, stringsAsFactors = FALSE)
data=cbind(cox7a2l_allele,data) ## COX7A2L allele added to the front of the file
#To_Drop=which(cox7a2l_allele=="Heterozygous") # If we decide to drop heterozygous
#data <- data[-To_Drop,]

# The pathway info is the same as used for Figure 4, but now just matching based on gene symbol, so the other columns of metadata are stripped out. 
pathway_info=read.csv(file="~/Dropbox/Aging\ BXD\ Study/1\ _\ Manuscript\ -\ Evan\ Multi-Omic/b_Figure5_HelperFile_GeneSets.csv", header=T, stringsAsFactors = FALSE)

TargetPathway=pathway_info$REACTOME_MITOCHONDRIAL_TRANSLATION
TargetPathway=TargetPathway[TargetPathway != ""]
Xmat <- as.matrix(data[,-(1:4)]) # This drops the multiple header columns, NOTE: Now we have four since COX7A2L was inserted. 
env <- data[,1] ## Now we set COX7A2L allele as the target environment
varnames <- colnames(Xmat)
A <- env ## Set variables for a single target gene against all others, and the controlling variable (env).
## now to compare against a full pathway and not just a target gene
X = Xmat[,!colnames(Xmat) %in% TargetPathway]
Y = Xmat[,colnames(Xmat) %in% TargetPathway]
Y=rowMeans(Y)

## Fix parameters
cores <- 3 # Can change depending on your computer
Sys.time()
pars <- list(m=6, B=5000, alpha_stab=0.1, alpha_pred=0.01, size_weight="linear", prescreen_size=50, use_resampling=FALSE,stab_test="exact",variable_importance="scaled_coefficient")

# Num reps sets minimum p-value (e.g. 100 == p = 0.01 increments, p = 10, p = 0.10 increments); this is what takes the most time.
fit_obj <- SRanalysis(X, Y, A, num_reps=100, pars_SR=pars, cores=cores)
plot(fit_obj, x_axis="SRpred", varnames=colnames(X), labels=TRUE)

# If desired, the actual values of the plot can be output to a table (can attach this to every stability-regression analysis; code only shown here)
write.table(fit_obj$results, "~/Desktop/MitoTranslation_COX7A2LAllele_mRNA.csv",sep=",")

# RIGHT PANEL:
# Separating based on COX7A2L allele rather than diet or age
cox7a2l_allele=c("B6","B6","B6","D2","D2","B6","B6","B6","D2","D2","B6","B6","D2","D2","B6","Heterozygous","B6","B6","B6","D2","D2","B6","D2","B6","B6","B6","B6","B6","B6","B6","B6","Heterozygous","Heterozygous","B6","B6","Heterozygous","B6","B6","B6","B6","D2","D2","D2","D2","D2","B6","D2","D2","B6","D2","D2","B6","D2","D2","D2","D2","Heterozygous","B6","D2","B6","D2","D2","D2","B6","Heterozygous","D2","B6","D2","B6","D2","D2","B6","D2","D2","D2","B6","D2","B6","D2","B6","B6","B6","D2","D2","B6","B6","B6","B6","D2","B6","B6","D2","D2","B6","B6","D2","B6","B6","Heterozygous","D2","D2","B6","D2","D2","D2","B6","B6","B6","B6","D2","B6","B6","B6","Heterozygous","B6","B6","B6","D2","B6","B6","D2","Heterozygous","D2","D2","D2","D2","D2","B6","B6","B6","D2","B6","B6","B6","B6","B6","D2","B6","B6","D2","D2","D2","B6","D2","B6","D2","D2","B6","D2","D2","B6","B6","D2","B6","B6","B6","D2","D2","D2","D2","D2","B6","D2","B6","D2","B6","D2","B6","B6","B6","D2","B6","B6","D2","D2","D2","B6","D2","B6","D2","B6","B6","D2","D2","D2","B6","D2","B6","D2","D2","B6","Heterozygous","B6","D2","B6","B6","D2","B6","B6","B6","Heterozygous","B6","D2","Heterozygous","B6","B6","D2","B6","B6","Heterozygous","B6","D2","B6","B6","Heterozygous","Heterozygous","Heterozygous","D2","D2","D2","B6","Heterozygous","B6","D2","B6","B6","B6","B6","B6","D2","D2","D2","B6","B6","B6","B6","B6","B6","D2","Heterozygous","B6","B6","B6","D2","B6","B6","B6","B6","Heterozygous","B6","D2","D2","B6","D2","B6","B6","B6","B6","D2","B6","B6","B6","B6","D2","B6","B6","B6","Heterozygous","B6","B6","B6","B6","D2","B6","B6","B6","D2","B6","D2","B6","B6","D2","B6","B6","B6","D2","B6","D2","Heterozygous","B6","B6","D2","D2","Heterozygous","D2","D2","B6","D2","D2","Heterozygous","Heterozygous","D2","B6","B6","D2","D2","B6","B6","B6","D2","B6","B6","D2","Heterozygous","D2")

data=read.csv(file="~/Dropbox/Aging\ BXD\ Study/1\ _\ Manuscript\ -\ Evan\ Multi-Omic/b_Figure5_HelperFile_ProteinData.csv", header=T, stringsAsFactors = FALSE)
data=cbind(cox7a2l_allele,data) ## COX7A2L allele added to the front of the file

# The pathway info is the same as used for Figure 4, but now just matching based on gene symbol, so the other columns of metadata are stripped out. 
pathway_info=read.csv(file="~/Dropbox/Aging\ BXD\ Study/1\ _\ Manuscript\ -\ Evan\ Multi-Omic/b_Figure5_HelperFile_GeneSets.csv", header=T, stringsAsFactors = FALSE)

TargetPathway=pathway_info$REACTOME_MITOCHONDRIAL_TRANSLATION
TargetPathway=TargetPathway[TargetPathway != ""]
Xmat <- as.matrix(data[,-(1:4)]) # This drops the multiple header columns, NOTE: Now we have four since COX7A2L was inserted. 
env <- data[,1] ## Now we set COX7A2L allele as the target environment
varnames <- colnames(Xmat)
A <- env ## Set variables for a single target gene against all others, and the controlling variable (env).
## now to compare against a full pathway and not just a target gene
X = Xmat[,!colnames(Xmat) %in% TargetPathway]
Y = Xmat[,colnames(Xmat) %in% TargetPathway]
Y=rowMeans(Y)

## Fix parameters
cores <- 3 # Can change depending on your computer
Sys.time()
pars <- list(m=6, B=5000, alpha_stab=0.1, alpha_pred=0.01, size_weight="linear", prescreen_size=50, use_resampling=FALSE,stab_test="exact",variable_importance="scaled_coefficient")

# Num reps sets minimum p-value (e.g. 100 == p = 0.01 increments, p = 10, p = 0.10 increments); this is what takes the most time.
fit_obj <- SRanalysis(X, Y, A, num_reps=100, pars_SR=pars, cores=cores)
plot(fit_obj, x_axis="SRpred", varnames=colnames(X), labels=TRUE)


# If desired, the actual values of the plot can be output to a table (can attach this to every stability-regression analysis; code only shown here)
write.table(fit_obj$results, "~/Desktop/MitoTranslation_COX7A2LAllele_Protein2.csv",sep=",")


#$$$$$$ #$$$$$$ #$$$$$$ #$$$$$$ #$$$$$$ #$$$$$$ #$$$$$$ #$$$$$$
#$$$$$$ #$$$$$$ #$$$$$$ #$$$$$$ #$$$$$$ #$$$$$$ #$$$$$$ #$$$$$$
# PANEL S4A (Supplemental to Figure 5)
# Must load the above libraries and the "CorrGraph" function at the end of this file.

# LEFT PLOT
pep1=AllData_CD
GeneSet=read.csv(file="~/Dropbox/Aging\ BXD\ Study/1\ _\ Manuscript\ -\ Evan\ Multi-Omic/b_HelpNetworks_ALL_NETWORKS_CANON_mRNA_Prot_ALIGNED.csv", header=T, stringsAsFactors = FALSE)
target.cats <- c("mRNA_REACTOME_CHOLESTEROL_BIOSYNTHESIS","mRNA_REACTOME_CHOLESTEROL_BIOSYNTHESIS_NEWHITS")
GeneSet <- GeneSet[which(GeneSet$Description %in% target.cats),] 
GeneSet1<-GeneSet
CorrGraph(pep1, GeneSet1, 0.0001, colors=c("lightblue","lightblue","green", "darkblue","darkblue")) 

# RIGHT PLOT
pep1=AllData_HF
GeneSet=read.csv(file="~/Dropbox/Aging\ BXD\ Study/1\ _\ Manuscript\ -\ Evan\ Multi-Omic/b_HelpNetworks_ALL_NETWORKS_CANON_mRNA_Prot_ALIGNED.csv", header=T, stringsAsFactors = FALSE)
target.cats <- c("mRNA_REACTOME_CHOLESTEROL_BIOSYNTHESIS","mRNA_REACTOME_CHOLESTEROL_BIOSYNTHESIS_NEWHITS")
GeneSet <- GeneSet[which(GeneSet$Description %in% target.cats),] 
GeneSet1<-GeneSet
CorrGraph(pep1, GeneSet1, 0.0001, colors=c("lightblue","lightblue","green", "darkblue","darkblue")) 


#$$$$$$ #$$$$$$ #$$$$$$ #$$$$$$ #$$$$$$ #$$$$$$ #$$$$$$ #$$$$$$
#$$$$$$ #$$$$$$ #$$$$$$ #$$$$$$ #$$$$$$ #$$$$$$ #$$$$$$ #$$$$$$
# PANEL S4B (Supplemental to Figure 5)
# Each of these four plots is a superimposition of 100x plots separating a randomized version of mitochondrial translation (top) or PPAR signalling (bottom) for the mRNA and protein input datasets. This is an empirical way to determine the approximate false positive rate. 


#$$$$$$ #$$$$$$ #$$$$$$ #$$$$$$ #$$$$$$ #$$$$$$ #$$$$$$ #$$$$$$
#$$$$$$ #$$$$$$ #$$$$$$ #$$$$$$ #$$$$$$ #$$$$$$ #$$$$$$ #$$$$$$
# PANEL S4C (Supplemental to Figure 5)
# Each of these four plots is a superimposition of 100x plots separating a randomiezd version of mitochondrial translation (top) or PPAR signalling (bottom) for the mRNA and protein input datasets. This is an empirical way to determine the approximate false positive rate. 

## LEFT PANEL
# This is a reformatted version of the "AllData" file, with the top transcript per gene (taking the first one in the list) for all protein–mRNA matching genes
# THEN, as high numbers of reads of "0" distort correlations, any transcript with > 50 0's was discarded, so this mRNA file is somewhat smaller (3677 genes)
data=read.csv(file="~/Dropbox/Aging\ BXD\ Study/1\ _\ Manuscript\ -\ Evan\ Multi-Omic/b_Figure5_HelperFile_mRNAData.csv", header=T, stringsAsFactors = FALSE)
# The pathway info is the same as used for Figure 4, but now just matching based on gene symbol, so the other columns of metadata are stripped out. 
pathway_info=read.csv(file="~/Dropbox/Aging\ BXD\ Study/1\ _\ Manuscript\ -\ Evan\ Multi-Omic/b_Figure5_HelperFile_GeneSets.csv", header=T, stringsAsFactors = FALSE)

TargetPathway=pathway_info$KEGG_PEROXISOME
TargetPathway=TargetPathway[TargetPathway != ""]
Xmat <- as.matrix(data[,-(1:3)]) # This drops the multiple header columns
env <- data[,2] ## This sets either diet or age as the key variable. 3 = age, 2 = diet (for these input files)
varnames <- colnames(Xmat)
A <- env ## Set variables for a single target gene against all others, and the controlling variable (env).
## now to compare against a full pathway and not just a target gene
X = Xmat[,!colnames(Xmat) %in% TargetPathway]
Y = Xmat[,colnames(Xmat) %in% TargetPathway]
Y=rowMeans(Y)

## Fix parameters
cores <- 3 # Can change depending on your computer
Sys.time()
pars <- list(m=6, B=5000, alpha_stab=0.1, alpha_pred=0.01, size_weight="linear", prescreen_size=50, use_resampling=FALSE,stab_test="exact",variable_importance="scaled_coefficient")

# Num reps sets minimum p-value (e.g. 100 == p = 0.01 increments, p = 10, p = 0.10 increments); this is what takes the most time.
fit_obj <- SRanalysis(X, Y, A, num_reps=100, pars_SR=pars, cores=cores)
plot(fit_obj, x_axis="SRpred", varnames=colnames(X), labels=TRUE)

# If desired, the actual values of the plot can be output to a table (can attach this to every stability-regression analysis; code only shown here)
write.table(fit_obj$results, "~/Desktop/Peroxisome_Diet_mRNA.csv",sep=",")


## RIGHT PANEL
data=read.csv(file="~/Dropbox/Aging\ BXD\ Study/1\ _\ Manuscript\ -\ Evan\ Multi-Omic/b_Figure5_HelperFile_mRNAData.csv", header=T, stringsAsFactors = FALSE)
# The pathway info is the same as used for Figure 4, but now just matching based on gene symbol, so the other columns of metadata are stripped out. 
pathway_info=read.csv(file="~/Dropbox/Aging\ BXD\ Study/1\ _\ Manuscript\ -\ Evan\ Multi-Omic/b_Figure5_HelperFile_GeneSets.csv", header=T, stringsAsFactors = FALSE)

TargetPathway=pathway_info$KEGG_PEROXISOME
TargetPathway=TargetPathway[TargetPathway != ""]
Xmat <- as.matrix(data[,-(1:3)]) # This drops the multiple header columns
env <- data[,3] ## This sets either diet or age as the key variable. 3 = age, 2 = diet (for these input files)
varnames <- colnames(Xmat)
A <- env ## Set variables for a single target gene against all others, and the controlling variable (env).
## now to compare against a full pathway and not just a target gene
X = Xmat[,!colnames(Xmat) %in% TargetPathway]
Y = Xmat[,colnames(Xmat) %in% TargetPathway]
Y=rowMeans(Y)

## Fix parameters
cores <- 3 # Can change depending on your computer
Sys.time()
pars <- list(m=6, B=5000, alpha_stab=0.1, alpha_pred=0.01, size_weight="linear", prescreen_size=50, use_resampling=FALSE,stab_test="exact",variable_importance="scaled_coefficient")

# Num reps sets minimum p-value (e.g. 100 == p = 0.01 increments, p = 10, p = 0.10 increments); this is what takes the most time.
fit_obj <- SRanalysis(X, Y, A, num_reps=100, pars_SR=pars, cores=cores)
plot(fit_obj, x_axis="SRpred", varnames=colnames(X), labels=TRUE)

write.table(fit_obj$results, "~/Desktop/Peroxisome_Age_mRNA.csv",sep=",")


#$$$$$$ #$$$$$$ #$$$$$$ #$$$$$$ #$$$$$$ #$$$$$$ #$$$$$$ #$$$$$$
#$$$$$$ #$$$$$$ #$$$$$$ #$$$$$$ #$$$$$$ #$$$$$$ #$$$$$$ #$$$$$$
# PANEL S4D (Supplemental to Figure 5)

## LEFT PANEL
# THEN, as high numbers of reads of "0" distort correlations, any transcript with > 50 0's was discarded, so this mRNA file is somewhat smaller (3677 genes)
data=read.csv(file="~/Dropbox/Aging\ BXD\ Study/1\ _\ Manuscript\ -\ Evan\ Multi-Omic/b_Figure5_HelperFile_mRNAData.csv", header=T, stringsAsFactors = FALSE)
# The pathway info is the same as used for Figure 4, but now just matching based on gene symbol, so the other columns of metadata are stripped out. 
pathway_info=read.csv(file="~/Dropbox/Aging\ BXD\ Study/1\ _\ Manuscript\ -\ Evan\ Multi-Omic/b_Figure5_HelperFile_GeneSets.csv", header=T, stringsAsFactors = FALSE)

TargetPathway=pathway_info$KEGG_PEROXISOME
TargetPathway=TargetPathway[TargetPathway != ""]
Xmat <- as.matrix(data[,-(1:3)]) # This drops the multiple header columns
env <- data[,2] ## This sets either diet or age as the key variable. 3 = age, 2 = diet (for these input files)
varnames <- colnames(Xmat)
A <- env ## Set variables for a single target gene against all others, and the controlling variable (env).
## now to compare against a full pathway and not just a target gene
X = Xmat[,!colnames(Xmat) %in% TargetPathway]
Y = Xmat[,colnames(Xmat) %in% TargetPathway]
Y=rowMeans(Y)

## Fix parameters
cores <- 3 # Can change depending on your computer
Sys.time()
pars <- list(m=6, B=5000, alpha_stab=0.1, alpha_pred=0.01, size_weight="linear", prescreen_size=50, use_resampling=FALSE,stab_test="exact",variable_importance="scaled_coefficient")

# Num reps sets minimum p-value (e.g. 100 == p = 0.01 increments, p = 10, p = 0.10 increments); this is what takes the most time.
fit_obj <- SRanalysis(X, Y, A, num_reps=100, pars_SR=pars, cores=cores)
plot(fit_obj, x_axis="SRpred", varnames=colnames(X), labels=TRUE)

# If desired, the actual values of the plot can be output to a table (can attach this to every stability-regression analysis; code only shown here)
write.table(fit_obj$results, "~/Desktop/CYP450_Diet_Protein.csv",sep=",")


## RIGHT PANEL
data=read.csv(file="~/Dropbox/Aging\ BXD\ Study/1\ _\ Manuscript\ -\ Evan\ Multi-Omic/b_Figure5_HelperFile_mRNAData.csv", header=T, stringsAsFactors = FALSE)
# The pathway info is the same as used for Figure 4, but now just matching based on gene symbol, so the other columns of metadata are stripped out. 
pathway_info=read.csv(file="~/Dropbox/Aging\ BXD\ Study/1\ _\ Manuscript\ -\ Evan\ Multi-Omic/b_Figure5_HelperFile_GeneSets.csv", header=T, stringsAsFactors = FALSE)

TargetPathway=pathway_info$KEGG_PEROXISOME
TargetPathway=TargetPathway[TargetPathway != ""]
Xmat <- as.matrix(data[,-(1:3)]) # This drops the multiple header columns
env <- data[,3] ## This sets either diet or age as the key variable. 3 = age, 2 = diet (for these input files)
varnames <- colnames(Xmat)
A <- env ## Set variables for a single target gene against all others, and the controlling variable (env).
## now to compare against a full pathway and not just a target gene
X = Xmat[,!colnames(Xmat) %in% TargetPathway]
Y = Xmat[,colnames(Xmat) %in% TargetPathway]
Y=rowMeans(Y)

## Fix parameters
cores <- 3 # Can change depending on your computer
Sys.time()
pars <- list(m=6, B=5000, alpha_stab=0.1, alpha_pred=0.01, size_weight="linear", prescreen_size=50, use_resampling=FALSE,stab_test="exact",variable_importance="scaled_coefficient")

# Num reps sets minimum p-value (e.g. 100 == p = 0.01 increments, p = 10, p = 0.10 increments); this is what takes the most time.
fit_obj <- SRanalysis(X, Y, A, num_reps=100, pars_SR=pars, cores=cores)
plot(fit_obj, x_axis="SRpred", varnames=colnames(X), labels=TRUE)

write.table(fit_obj$results, "~/Desktop/CYP450_Age_Protein.csv",sep=",")


#$$$$$$ #$$$$$$ #$$$$$$ #$$$$$$ #$$$$$$ #$$$$$$ #$$$$$$ #$$$$$$
#$$$$$$ #$$$$$$ #$$$$$$ #$$$$$$ #$$$$$$ #$$$$$$ #$$$$$$ #$$$$$$
# PANEL S4E (Supplemental to Figure 5)
library(StabilizedRegression)


# Stability-Prediction Plot
# LEFT PANEL:
# Separating based on COX7A2L allele rather than diet or age
hmgcs2_allele=c("B6","D2","B6","D2","B6","B6","B6","B6","B6","D2","D2","D2","B6","B6","D2","D2","B6","B6","B6","B6","B6","D2","D2","B6","D2","B6","B6","D2","D2","D2","B6","B6","B6","D2","D2","D2","B6","B6","B6","B6","D2","D2","B6","heterozygous","D2","D2","B6","heterozygous","B6","heterozygous","heterozygous","B6","D2","D2","D2","B6","D2","D2","D2","B6","B6","B6","D2","heterozygous","D2","B6","B6","B6","D2","D2","D2","B6","B6","D2","B6","B6","D2","B6","D2","B6","B6","D2","D2","D2","D2","D2","D2","B6","B6","D2","B6","D2","D2","D2","D2","D2","D2","B6","D2","B6","D2","D2","B6","D2","D2","B6","B6","B6","B6","D2","B6","B6","B6","D2","D2","D2","B6","B6","B6","B6","D2","B6","heterozygous","B6","heterozygous","heterozygous","B6","D2","D2","B6","B6","B6","B6","heterozygous","B6","B6","D2","heterozygous","D2","B6","D2","B6","B6","B6","D2","D2","D2","D2","D2","B6","B6","D2","D2","B6","B6","D2","B6","B6","B6","B6","heterozygous","D2","D2","D2","D2","B6","heterozygous","B6","B6","B6","B6","D2","B6","D2","B6","B6","B6","B6","D2","B6","D2","B6","B6","B6","B6","B6","B6","D2","D2","B6","B6","D2","B6","D2","B6","B6","B6","B6","D2","heterozygous","B6","B6","D2","D2","D2","D2","D2","B6","D2","D2","heterozygous","B6","heterozygous","heterozygous","B6","B6","B6","D2","D2","D2","D2","D2","D2","B6","heterozygous","B6","D2","D2","B6","D2","D2","D2","D2","D2","B6","D2","B6","D2","B6","B6","B6","D2","B6","D2","D2","D2","D2","D2","D2","D2","B6","heterozygous","B6","B6","B6","D2","D2","D2","D2","B6","D2","D2","D2","B6","B6","B6","heterozygous","heterozygous","B6","B6","D2","D2","D2","B6","B6","B6","B6","D2","D2","D2","D2","D2","B6","B6","D2","D2","B6","D2","B6","D2","B6")

# The pathway info is the same as used for Figure 4, but now just matching based on gene symbol, so the other columns of metadata are stripped out. 
data=read.csv(file="~/Dropbox/Aging\ BXD\ Study/1\ _\ Manuscript\ -\ Evan\ Multi-Omic/b_Figure5_HelperFile_mRNAData.csv", header=T, stringsAsFactors = FALSE)
data=cbind(hmgcs2_allele,data) ## COX7A2L allele added to the front of the file

# The pathway info is the same as used for Figure 4, but now just matching based on gene symbol, so the other columns of metadata are stripped out. 
pathway_info=read.csv(file="~/Dropbox/Aging\ BXD\ Study/1\ _\ Manuscript\ -\ Evan\ Multi-Omic/b_Figure5_HelperFile_GeneSets.csv", header=T, stringsAsFactors = FALSE)

TargetPathway=pathway_info$REACTOME_MITOCHONDRIAL_TRANSLATION
TargetPathway=TargetPathway[TargetPathway != ""]
Xmat <- as.matrix(data[,-(1:4)]) # This drops the multiple header columns, NOTE: Now we have four since COX7A2L was inserted. 
env <- data[,1] ## Now we set HMGCS2 allele as the target environment
varnames <- colnames(Xmat)
A <- env ## Set variables for a single target gene against all others, and the controlling variable (env).
## now to compare against a full pathway and not just a target gene
X = Xmat[,!colnames(Xmat) %in% TargetPathway]
Y = Xmat[,colnames(Xmat) %in% TargetPathway]
Y=rowMeans(Y)

## Fix parameters
cores <- 3 # Can change depending on your computer
Sys.time()
pars <- list(m=6, B=5000, alpha_stab=0.1, alpha_pred=0.01, size_weight="linear", prescreen_size=50, use_resampling=FALSE,stab_test="exact",variable_importance="scaled_coefficient")

# Num reps sets minimum p-value (e.g. 100 == p = 0.01 increments, p = 10, p = 0.10 increments); this is what takes the most time.
fit_obj <- SRanalysis(X, Y, A, num_reps=100, pars_SR=pars, cores=cores)
plot(fit_obj, x_axis="SRpred", varnames=colnames(X), labels=TRUE)

# If desired, the actual values of the plot can be output to a table (can attach this to every stability-regression analysis; code only shown here)
write.table(fit_obj$results, "~/Desktop/MitoTranslation_HMGCS2Allele_mRNA.csv",sep=",")

# RIGHT PANEL:
# Separating based on COX7A2L allele rather than diet or age
hmgcs2_allele=c("B6","D2","D2","B6","B6","B6","D2","D2","D2","B6","D2","D2","B6","B6","D2","heterozygous","D2","D2","D2","B6","B6","D2","B6","D2","B6","B6","D2","D2","B6","B6","D2","heterozygous","heterozygous","D2","B6","heterozygous","D2","D2","D2","D2","D2","B6","B6","B6","D2","D2","B6","B6","B6","D2","D2","D2","B6","B6","B6","B6","heterozygous","D2","D2","D2","D2","D2","D2","B6","heterozygous","B6","D2","B6","B6","B6","B6","D2","D2","D2","B6","B6","B6","B6","B6","B6","B6","B6","B6","D2","B6","B6","D2","D2","D2","D2","B6","D2","B6","D2","D2","B6","B6","D2","heterozygous","B6","D2","D2","B6","D2","D2","B6","D2","D2","D2","B6","D2","D2","B6","heterozygous","B6","D2","D2","B6","B6","D2","B6","heterozygous","D2","D2","B6","B6","B6","D2","D2","D2","B6","B6","B6","D2","D2","D2","B6","D2","D2","D2","B6","B6","D2","B6","D2","B6","B6","D2","B6","B6","D2","B6","B6","B6","B6","D2","B6","B6","D2","B6","B6","D2","B6","D2","D2","D2","B6","D2","D2","B6","D2","B6","D2","B6","B6","D2","D2","D2","D2","D2","B6","B6","B6","B6","D2","B6","B6","D2","B6","D2","D2","heterozygous","B6","D2","B6","D2","B6","D2","B6","B6","heterozygous","B6","D2","heterozygous","B6","D2","D2","D2","B6","heterozygous","B6","B6","D2","D2","heterozygous","heterozygous","heterozygous","B6","D2","D2","B6","heterozygous","D2","D2","B6","D2","B6","B6","B6","B6","B6","D2","B6","B6","D2","D2","D2","D2","D2","heterozygous","B6","D2","D2","D2","D2","D2","B6","D2","heterozygous","B6","B6","D2","D2","B6","B6","D2","D2","D2","B6","B6","B6","B6","D2","B6","D2","D2","D2","heterozygous","D2","D2","B6","B6","B6","D2","D2","D2","B6","D2","B6","B6","D2","B6","B6","D2","D2","D2","B6","B6","heterozygous","D2","D2","B6","B6","heterozygous","B6","D2","D2","B6","B6","heterozygous","heterozygous","B6","D2","B6","B6","B6","B6","D2","D2","D2","D2","D2","B6","heterozygous","B6")

data=read.csv(file="~/Dropbox/Aging\ BXD\ Study/1\ _\ Manuscript\ -\ Evan\ Multi-Omic/b_Figure5_HelperFile_ProteinData.csv", header=T, stringsAsFactors = FALSE)
data=cbind(hmgcs2_allele,data) ## COX7A2L allele added to the front of the file

# The pathway info is the same as used for Figure 4, but now just matching based on gene symbol, so the other columns of metadata are stripped out. 
pathway_info=read.csv(file="~/Dropbox/Aging\ BXD\ Study/1\ _\ Manuscript\ -\ Evan\ Multi-Omic/b_Figure5_HelperFile_GeneSets.csv", header=T, stringsAsFactors = FALSE)

TargetPathway=pathway_info$REACTOME_MITOCHONDRIAL_TRANSLATION
TargetPathway=TargetPathway[TargetPathway != ""]
Xmat <- as.matrix(data[,-(1:4)]) # This drops the multiple header columns, NOTE: Now we have four since COX7A2L was inserted. 
env <- data[,1] ## Now we set HMGCS2 allele as the target environment
varnames <- colnames(Xmat)
A <- env ## Set variables for a single target gene against all others, and the controlling variable (env).
## now to compare against a full pathway and not just a target gene
X = Xmat[,!colnames(Xmat) %in% TargetPathway]
Y = Xmat[,colnames(Xmat) %in% TargetPathway]
Y=rowMeans(Y)

## Fix parameters
cores <- 3 # Can change depending on your computer
Sys.time()
pars <- list(m=6, B=5000, alpha_stab=0.1, alpha_pred=0.01, size_weight="linear", prescreen_size=50, use_resampling=FALSE,stab_test="exact",variable_importance="scaled_coefficient")

# Num reps sets minimum p-value (e.g. 100 == p = 0.01 increments, p = 10, p = 0.10 increments); this is what takes the most time.
fit_obj <- SRanalysis(X, Y, A, num_reps=100, pars_SR=pars, cores=cores)
plot(fit_obj, x_axis="SRpred", varnames=colnames(X), labels=TRUE)


# If desired, the actual values of the plot can be output to a table (can attach this to every stability-regression analysis; code only shown here)
write.table(fit_obj$results, "~/Desktop/MitoTranslation_HMGCS2Allele_Protein2.csv",sep=",")


######################################################################################################################################################
######################################################################################################################################################
# CorrGraph function "CORRGRAPH.FIND"
######################################################################################################################################################
CorrGraph <- function(pep1, GeneSet1, cutofflevel, colors) {

tmp = rownames(pep1)
pep1 = pep1[tmp,]

GeneSet1 = GeneSet1[GeneSet1[,3] != "",]	# The first column in the data file must match the third column in the "gene list" file!
rnams = as.character(rownames(pep1))

xx = getMatchingRows(GeneSet1[,3], rnams) # Get the row number of matching rows
GeneMatch = pep1[xx,]
GeneMatch = GeneMatch[order(rownames(GeneMatch)),] 

siglevel = cutofflevel
transform_GeneMatch=apply(t(GeneMatch), c(1,2), as.numeric) # If NAs are coerced in, then it will force everything into a character matrix
tmpr <- corAndPvalue(transform_GeneMatch,method="spearman")
tmp = tmpr$cor
siz = dim(tmp)/2

diag(tmp) = 0
tmp[tmpr$p > siglevel] = 0 	# removes all correlation coefficients if their p-value is less than "siglevel" above.

# This sets the colors for the groups assigned in GeneList
mypaletteDiv = colors

# This makes sure the dots and colors correspond correctly
idx = match(rownames(tmp),as.character(GeneSet1[,3]))
graphnames = as.character(GeneSet1[idx,2])
colornames = as.numeric(GeneSet1[idx,5])

nedges = (sum(colSums(tmp !=0)) - sum(colSums(tmp == 1)))/2	# THIS COUNTS THE NUMBER OF EDGES, REMOVES SELF-EDGES, AND DIVIDES BY 2 (DOUBLED MATRIX)
nposedges = (sum(colSums(tmp >0)) - sum(colSums(tmp == 1)))/2	# THIS COUNTS THE NUMBER OF POSITIVE EDGES, REMOVES SELF-EDGES, AND DIVIDES BY 2 (DOUBLED MATRIX)
nnegedges = (sum(colSums(tmp <0)) - sum(colSums(tmp == 1)))/2	# THIS COUNTS THE NUMBER OF NEGATIVE EDGES, REMOVES SELF-EDGES, AND DIVIDES BY 2 (DOUBLED MATRIX)
nnodes = length(idx)
mean(tmpr$cor)		# Average correlation
mean(abs(tmpr$cor))	# Average absolute value of correlation

gr = plotGraph((tmp),names=graphnames, main = paste("P <", siglevel), vertcol=mypaletteDiv[colornames],repulserad=5e4)
plotGraph((tmp),names=graphnames, main = paste("P <", siglevel), vertcol=mypaletteDiv[colornames],repulserad=.6, nodesize=3.8, framecol="black", textcolor="black", myl = gr$myl)
legend(x=-1.4, y=1.2, bg='white', bty='n', legend=c(paste("Positive Edges = ", nposedges), paste("Negative Edges = ", nnegedges), paste("Node Count = ", nnodes)))

}
```
