## Supplementary material for "Multi-Omic Profiling of the Liver Across Diets and Age in a Diverse Mouse Population": All code used to generate figures: b_helpFunctions.R

```
# Note that this is used for plotting network graphs in R
#help functions
source("~/Dropbox/Aging\ BXD\ Study/1\ _\ Manuscript\ -\ Evan\ Multi-Omic/b_helpFunctions_unilities.R")

library(igraph)

getMatchingRows = function(uniprotnam , rnams){
  xx <- NULL
  for(i in 1:length(uniprotnam)){
    dd =  which(uniprotnam[i]  == rnams)
    xx <-c(xx,(dd))
  }
  xx <- xx[!is.na(xx)]
  return(xx)
}

plotHistCor2Cond = function(tmp,main=""){
  dow = 1:(dim(tmp)[1]/2)
  up = (dim(tmp)[1]/2+1):dim(tmp)[1]
  hist(uppertriang(tmp[dow,dow]),angle=-45,col=1,density=10,xlim=c(-1,1),main=main)
  hist(uppertriang(tmp[up,up]),add=TRUE,col=2,border=T,density=10)
  hist(uppertriang(tmp[dow,up]),add=TRUE,col=3,border=T,density=5,angle=30)
  legend("topleft",c("HFD","CD","HFD/CD"),col=c(1,2,3),lty=c(1,1,1))
}


selectDifferenceInteractions = function(tmp, diffthresh = 0.54){
  dow = 1:(dim(tmp)[1]/2)
  up = (dim(tmp)[1]/2+1):dim(tmp)[1]
  HDFg = tmp[up,up]
  CDg = tmp[dow,dow]
  R = cor(c(HDFg),c(CDg),method="spearman")
  plot(c(HDFg),c(CDg),xlab= "cor(HFD)", ylab="cor(CD)")
  legend("topleft",paste("R=",round(R,digits=2)))
  
  diffHDFCD = (HDFg - tmp[dow,dow])
  idx=diffHDFCD>diffthresh | diffHDFCD < -diffthresh
  points(HDFg[idx],CDg[idx],col=2)
  diffHDFCD[!idx] = 0
  return(diffHDFCD)
}

plotGraph <- function(matr, names, main = "cor(HFD)-cor(CD)",vertcol = NULL,deleteemptyvertexes = TRUE,cex=1,repulserad=1, framecol="white", nodesize=6, textcolor="black",myl = NULL, rescale=TRUE){
  graph.speed <- createGraph((matr),names)
  ww = E(graph.speed)$weight
  E(graph.speed)$color=E(graph.speed)$weight
  if(!is.null(vertcol)){   V(graph.speed)$color=vertcol }
  
  if(deleteemptyvertexes==TRUE){   graph.speed <- delete.vertices(graph.speed,which(degree(graph.speed)==0)) } # if null, then this deletes vertexes with no edges

  if(is.null(myl)){
    myl <- layout.fruchterman.reingold(graph.speed,weights=abs(ww))*repulserad
  }
  plotGraph2(graph.speed,myl,cex,cexdot,main=main, framecol, nodesize, textcolor, rescale)
  return(list(graph = graph.speed, myl = myl, rescale))
}

getDown = function(tmp){
  dow = 1:(dim(tmp)[1]/2)
  return(tmp[dow,dow])
}

getUp = function(tmp){
  up = (dim(tmp)[1]/2+1):dim(tmp)[1]
  return(tmp[up,up])
}


getDownUp = function(tmp){
  down = 1:(dim(tmp)[1]/2)
  up = (dim(tmp)[1]/2+1):dim(tmp)[1]
  return(tmp[down,up])
}

graphMeasures = function(gdiff){
  res = list(
    degree = centralization.degree(gdiff)$centralization,
    centralization = centralization.closeness(gdiff)$centralization,
    betweeness = centralization.betweenness(gdiff)$centralization
  )
  return(res)
}
```
