## Supplementary material for "Multi-Omic Profiling of the Liver Across Diets and Age in a Diverse Mouse Population": All code used to generate figures: b_helpFunctions_StabilityPredictionPlots.R

```
# Beforehand: on CRAN, install glmnet, questionr, and corpcor (w/ dependencies)

#
#install.packages("~/Documents/GitHub/StabilizedRegression-R/StabilizedRegression_1.0.tar.gz")

library(StabilizedRegression)


# To load the data:
#data=read.csv(file="~/Dropbox/Aging\ BXD\ Study/P\ _\ NIKLAS\ PFISTER\ COLLAB/1_R_INPUT_preprocessed_full_mRNA_data_v2_plusphenotypes.csv", header=T, stringsAsFactors = FALSE)
#data=read.csv(file="~/Dropbox/Aging\ BXD\ Study/P\ _\ NIKLAS\ PFISTER\ COLLAB/1_R_INPUT_preprocessed_top3900only_mRNA_data_v2.csv", header=T, stringsAsFactors = FALSE)
#data=read.csv(file="~/Dropbox/Aging\ BXD\ Study/P\ _\ NIKLAS\ PFISTER\ COLLAB/1_R_INPUT_preprocessed_top3900only_protein_data_v2_withphenotypes.csv", header=T, stringsAsFactors = FALSE)

# These are artificial "independent" variables created using strong cis-QTLs, one by one
#data=read.csv(file="~/Dropbox/Aging\ BXD\ Study/P\ _\ NIKLAS\ PFISTER\ COLLAB/1_R_INPUT_preprocessed_top3900only_protein_data_vTTEST_QTLs.csv", header=T, stringsAsFactors = FALSE)
#data=read.csv(file="~/Dropbox/Aging\ BXD\ Study/P\ _\ NIKLAS\ PFISTER\ COLLAB/1_R_INPUT_preprocessed_full_mRNA_data_v2_vTTEST_QTLS.csv", header=T, stringsAsFactors = FALSE)

# To load the data on laptop:
#data=read.csv(file="~/Dropbox/Aging\ BXD\ Study/P\ _\ NIKLAS\ PFISTER\ COLLAB/1_R_INPUT_preprocessed_full_mRNA_data_v2.csv", header=T, stringsAsFactors = FALSE)
#data=read.csv(file="~/Dropbox/Aging\ BXD\ Study/P\ _\ NIKLAS\ PFISTER\ COLLAB/1_R_INPUT_preprocessed_top3900only_mRNA_data_v2.csv", header=T, stringsAsFactors = FALSE)
#data=read.csv(file="~/Dropbox/Aging\ BXD\ Study/P\ _\ NIKLAS\ PFISTER\ COLLAB/1_R_INPUT_preprocessed_top3900only_protein_data_v2_Cox7a2l.csv", header=T, stringsAsFactors = FALSE)
data=read.csv(file="~/Dropbox/Aging\ BXD\ Study/P\ _\ NIKLAS\ PFISTER\ COLLAB/1_R_INPUT_preprocessed_full_mRNA_data_v2_Cox7a2l.csv", header=T, stringsAsFactors = FALSE)

# to load the Gene Set info
pathway_info=read.csv(file="~/Dropbox/Aging\ BXD\ Study/P\ _\ NIKLAS\ PFISTER\ COLLAB/1_R_INPUT_GeneSets_v2.csv", header=T, stringsAsFactors = FALSE)


Sys.time()
#pathway_list <- pathway_info$pathway_list_mrna
### SET THIS BY HAND
#TargetPathway=pathway_info$REACTOME_CHOLESTEROL_BIOSYNTHESIS_EXTENDED
TargetPathway=pathway_info$REACTOME_MITOCHONDRIAL_TRANSLATION
#TargetPathway=pathway_info$BODYWEIGHT


#### TEMPORARY FOR "LEAVE ONE OUT" TESTS
#TargetPathway=TargetPathway[-10] # Removes element N in the pathway (GGPS1)
#TargetPathway=TargetPathway[-10] # Removes element N in the pathway (LBR)
#TargetPathway=TargetPathway[-10] # Removes element N in the pathway (SOAT1)

TargetPathway=TargetPathway[TargetPathway != ""]

# Change to drop 1:2 if you don't have age in as a cofactor
Xmat <- as.matrix(data[,-(1:3)])
## This sets either diet or age as the key variable. 3 = age, 2 = diet
env <- data[,3]
varnames <- colnames(Xmat)

## Set variables for a single target gene against all others, and the controlling variable (env).
#X <- Xmat[,varnames != "Rdh11"]
#Y <- Xmat[,varnames == "Rdh11"]
A <- env

## now suppose we want to compare against a full pathway and not just a target gene
X = Xmat[,!colnames(Xmat) %in% TargetPathway]
Y = Xmat[,colnames(Xmat) %in% TargetPathway]
Y=rowMeans(Y)


#Yes i think your script looks correct. The issue is that i set some of the parameters very low to make it fast when testing. To get the analysis i did you would need to set the following:
#- prescreen_size=50 (in pars_SR)
#- B=5000 (in pars_SR)
#- num_reps=200
## This takes about 2 minutes PER REP on my home iMac for mRNA data

## Fix parameters
cores <- 3
Sys.time()
pars <- list(m=6,
 			 B=5000, # change from 5000
             alpha_stab=0.1,
             alpha_pred=0.01,
             size_weight="linear",
             prescreen_size=50, # change from 500
             use_resampling=FALSE,
             stab_test="exact",
             variable_importance="scaled_coefficient")

# Num reps sets minimum p-value (e.g. 100 == p = 0.01 increments, p = 10, p = 0.10 increments); this is what takes the most time.
fit_obj <- SRanalysis(X, Y, A, num_reps=100,
                      pars_SR=pars, cores=cores)

fit_obj$results$names <- colnames(X) # This adds the column names to the output matrix

## Plot with SRdiff on the x_axis
#
#
#plot(fit_obj, x_axis="SRdiff", varnames=colnames(X), labels=TRUE)

## Plot with SRpred on the x_axis
plot(fit_obj, x_axis="SRpred", varnames=colnames(X), labels=TRUE)

Sys.time()


write.table(fit_obj$results, "~/Desktop/Hmgcs2_Allele_mRNA_TargetofMitoTranslation_ProteinHit_100x.csv",sep=",")
```
