## Supplementary material for "Multi-Omic Profiling of the Liver Across Diets and Age in a Diverse Mouse Population": All code used to generate figures: b_helpFunctions_unilities.R

```
#computes fold 
computefold = function(ratio2){
  source("helpFunctions_unilities.R")
  cnam = sub("-","_",colnames(ratio2))
  bl = fixSampleNames(cnam)
  bl = split2table( bl , split="\\_" )
  bl = bl[, 2:3 ]
  
  colnames(ratio2)=apply(bl,1,paste,collapse="_")
  
  cd = ratio2[,bl[,2] == "CD"]
  hfd = ratio2[,bl[,2] == "HFD"]
  dim(cd)
  dim(hfd)
  
  cd = cd[,which(bl[bl[,2] == "CD",1] %in% bl[bl[,2] == "HFD",1])]
  dim(cd)
  dim(hfd)
  colnames(cd)
  colnames(hfd)
  sum(is.na(hfd))
  sum(is.na(cd))
  hfd[hfd==0] = NA
  cd[cd ==0] = NA
  
  range(hfd)
  range(cd)
  strainratio  = hfd - cd
  return(strainratio)
}


getMolfun <- function(pfilename){
    pgl <- read.csv(pfilename , stringsAsFactors=FALSE)
    gofunction <- pgl$GOMolecularFunction
    dd <- lapply(gofunction , function(x){d <- strsplit(x,";"); return(d[[1]][1])})
    dd <- lapply(dd , function(x){d <- strsplit(x,","); return(d[[1]][1])})
    tmp <- (unlist(dd))
    return(list(ids=pgl$Mapped.IDs, molfun=tmp))
}

rmNarows <- function(plev )
{
    x <- apply(plev,1,function(x){sum(is.na(x))})
    plev <- plev[-which(x>0),]
}

#reads data rm na and log2
prepareData <- function(plev){
   pln <- log2(plev)
   return(pln)
}

##
selectSignificant2 <- function(dd, siglevel = 0.7, minPcor=0)
    {
        tmpq5 <- apply(dd,c(2,3),quantile,1-siglevel)
        tmpq95 <- apply(dd,c(2,3),quantile,siglevel)
        diag(tmpq5) <- 0
        diag(tmpq95) <- 0
        
        indic <- (tmpq5 > minPcor | tmpq95 < -minPcor)

        tmpmedian <- apply(dd,c(2,3),median)
        tmpmedian[!indic] <- 0
        diag(tmpmedian)=0
        return(tmpmedian)
    }

selectSignificant <- function(dd, siglevel = 0.8, minPcor=0)
    {
        tmpq5 <- apply(dd,c(2,3),quantile,1-siglevel)
        tmpq95 <- apply(dd,c(2,3),quantile,siglevel)
        diag(tmpq5) <- 0
        diag(tmpq95) <- 0
        
        indic <- (tmpq5 > 0 | tmpq95 < 0)

        tmpmedian <- apply(dd,c(2,3),median)
        tmpmedian[!indic] <- 0
        diag(tmpmedian)=0
        indic <- (tmpmedian > minPcor | tmpmedian < -minPcor)
        tmpmedian[!indic] <- 0
        return(tmpmedian)

    }


createGraph <- function(adjMatrix,protnam){
    library(igraph)
    graph.speed=graph.adjacency(adjmatrix=adjMatrix, mode="undirected", weighted=TRUE)
    V(graph.speed)$name = protnam
    temp.degree=degree(graph.speed)
    pname <- protnam
    pname[temp.degree==0]=NA
    V(graph.speed)$label=pname
    return(graph.speed)
}

# This doesn't appear to be the one that we use
plotGraph <- function(graph,myl=NULL){
    vdeg <- degree(graph)
    library(RColorBrewer)
    mypalette<-brewer.pal(9,"BuGn")
    vcols <- mypalette[round(rank(vdeg-max(vdeg))/length(vdeg) * 7)+1]

    mypalette <- brewer.pal(9,"YlGnBu")
    weights <- E(graph)$weight+1
    ecols <- round(((weights/max(weights)*8)+1))
    range(ecols)
    ecols <- mypalette[ecols]
    
    x <- plot(graph, layout=myl,  vertex.size=3, vertex.frame.color="red",  edge.color= ecols,
              vertex.color=vcols,
              vertex.label.font=1,
              vertex.label.cex=0.7,
              vertex.label.color=1,
              rescale=rescale)
    return(x)
}

# This seems to be the primary plotting function called

plotGraph2 <- function(graph,myl,cex,main="",vertcol=NULL, framecol, nodesize, textcolor, rescale){
    vdeg <- degree(graph)

    library(RColorBrewer)
    mypalette<-brewer.pal(9,"BuGn")
    #vcols <- mypalette[round(rank(vdeg-max(vdeg))/length(vdeg) * 7)+1]

    mypalette <- brewer.pal(9,"RdBu")
    weights <- E(graph)$color
    weights[weights>0] <- sqrt(weights[weights>0])
    weights[weights<0] <- -1* sqrt(-1*weights[weights<0])
    weights <- weights+1
    ecols <- round(((weights/2 * 8)+1))
    ecols <- mypalette[ecols]
    
    x <- plot(graph, layout=myl,  vertex.size=nodesize, vertex.frame.color=framecol,
              edge.color= ecols,
              rescale=rescale,
  #            ylim=c(-1, 1),
  #            xlim=c(-1, 1),
  #            asp=1,
              vertex.label.font=1,
              vertex.label.cex=cex,
              vertex.label.color=textcolor,
              vertex.color = V(graph)$color , main=main)
    return(x)
}


computeBoost <- function(pln,ffunc,nr=250)
    {
        result1=ffunc(pln)
        bla <- dim(result1)
        dd <- array(0,dim=c(nr,bla[1],bla[2]))
        for(i in 1:nr){
            print(i)
            ps <- pln[sample(dim(pln)[1],replace=TRUE),]
            dd[i,,] <- ffunc(as.matrix(ps))
        }
        return(dd)
    }

computeBoost2 <- function(pln,g1,g2,ffunc,nr=250)
    {
        result1 <- ffunc(pln)
        bla <- dim(result1)
        dd <- array(0,dim=c(nr,bla[1],bla[2]))
        for(i in 1:nr){
            print(i)
            ps <- pln[c(
                g1[sample(length(g1),replace=TRUE)]
                ,g2[sample(length(g2),replace=TRUE)]
                ),]
            dd[i,,] <- ffunc(as.matrix(ps))
        }
        return(dd)
    }


plotGraphAnn <- function(graph,myl,vfc="white",family=1){
    library(RColorBrewer)

    mypalette <- brewer.pal(9,"RdBu")
    weights <- E(graph)$color
    weights[weights>0] <- (weights[weights>0])^0.2
    weights[weights<0] <- -1* (-1*weights[weights<0])^0.2
    weights <- weights+1
    ecols <- round(((weights/2 * 8)+1))
    ecols <- mypalette[ecols]
    
    x <- plot(graph, layout=myl,  vertex.size=6, vertex.frame.color=vfc, edge.width=4, edge.color= ecols,vertex.label=V(graph)$protname ,vertex.label.family=family,vertex.label.font=1 ,vertex.label.dist=0.4, vertex.label.cex=1 , vertex.label.color=1)
    return(x)
}

plotGraphAnn2 <- function(graph,myl,vfc="white",family=1,edge.width=NULL,vertex.label.cex=1,vertex.label.color=1){
  weights <- E(graph)$weight
  cols = rep("pink",length(weights))
#  cols[weights > 0] = "cornflowerblue"
  cols[weights > 0] = "red"
  x <- plot(graph, layout=myl,  vertex.size=6, vertex.frame.color=vfc, edge.width=edge.width,
            edge.color= cols,vertex.label=V(graph)$protname ,vertex.label.family=family,
            vertex.label.font=1 ,vertex.label.dist=0.4, vertex.label.cex=vertex.label.cex ,
            vertex.label.color=vertex.label.color)
  return(x)
}


#quantile normalization
myqnormalize <- function(pln){
    tmp <- normalize.quantiles(as.matrix(log(pln)))
    rownames(tmp) <- rownames(pln)
    colnames(tmp) <- colnames(pln)
    tmp <- scale(tmp)
    plns1 <- t(scale(t(tmp)))
}

#get Colors for sample
getColorsforSample <- function(pln){
    samplenames <- colnames(pln)
    cols <- rep("",length(samplenames))
    cols[grep("HFD",samplenames)] <- "gray"
    cols[grep("CD",samplenames)] <- "white"
    return(cols)
}

giveCols2names <- function(protnam,tt,ml){
    mcol <- rep("",length(protnam))
    for(i in 1:length(protnam))
        {
            x <- (tt[which(tt$ids==protnam[i]),2])
            if(length(x)==0)
                {
                    mcol[i] <- "white" 
                }
            else if(is.na(x)){
                mcol[i] <- "white"
            }
            else{
                #cat(protnam[i]," " ,ml[[x]],"\n")
                if(length(ml[[x]])==0)
                    {
                        mcol[i] <- "lightgray"
                    }
                else
                    {
                        mcol[i] <- ml[[x]]
                    }
            }
        }
    return(mcol)
}

fixSampleNames <- function(cnames)
{
    cnames<-sub("\\.","\\_",cnames)
    return(cnames)
}

splitcolnames <- function(cnames)
{
    cnamessplit <- strsplit(cnames,"\\_")
    colnamestable <- NULL
    for(i in cnamessplit)
        {
            colnamestable <- rbind(colnamestable,i)
        }
    colnames(colnamestable) <- c("Time","strain","replicate","condition")
    colnamestable <- as.data.frame(colnamestable)
    ftable(colnamestable$Time,colnamestable$strain,colnamestable$replicate,colnamestable$condition)
}


split2table <- function(protnames,split="\\||\\_")
{
    protnames[1:10]
    cnamessplit <- strsplit(protnames,split)
    protnam <- NULL
    for(i in cnamessplit)
        {
            protnam <- rbind(protnam,i)
        }
    return(protnam)
}
```
